# An integrated single-nucleus and spatial transcriptomics atlas reveals the molecular landscape of the human hippocampus

**DOI:** 10.1101/2024.04.26.590643

**Authors:** Jacqueline R. Thompson, Erik D. Nelson, Madhavi Tippani, Anthony D. Ramnauth, Heena R. Divecha, Ryan A. Miller, Nicholas J. Eagles, Elizabeth A. Pattie, Sang Ho Kwon, Svitlana V. Bach, Uma M. Kaipa, Jianing Yao, Christine Hou, Joel E. Kleinman, Leonardo Collado-Torres, Shizhong Han, Kristen R. Maynard, Thomas M. Hyde, Keri Martinowich, Stephanie C. Page, Stephanie C. Hicks

## Abstract

The hippocampus contains many unique cell types, which serve the structure’s specialized functions, including learning, memory and cognition. These cells have distinct spatial organization, morphology, physiology, and connectivity, highlighting the importance of transcriptome-wide profiling strategies that retain cytoarchitectural organization. Here, we generated spatially-resolved transcriptomics (SRT) and single-nucleus RNA-sequencing (snRNA-seq) data from adjacent tissue sections of the anterior human hippocampus in ten adult neurotypical donors to define molecular profiles for hippocampal cell types and spatial domains. Using non-negative matrix factorization (NMF) and label transfer, we integrated these data by defining gene expression patterns within the snRNA-seq data and inferring their expression in the SRT data. We identified NMF patterns that captured transcriptional variation across neuronal cell types and indicated that the response of excitatory and inhibitory postsynaptic specializations were prioritized in different SRT spatial domains. We used the NMF and label transfer approach to leverage existing rodent datasets, identifying patterns of activity-dependent transcription and subpopulations of dentate gyrus granule cells in our SRT dataset that may be predisposed to participate in learning and memory ensembles. Finally, we characterized the spatial organization of NMF patterns corresponding to non-*cornu ammonis* pyramidal neurons and identified snRNA-seq clusters mapping to distinct regions of the retrohippocampus, to three subiculum layers, and to a population of presubiculum neurons. To make this comprehensive molecular atlas accessible to the scientific community, both raw and processed data are freely available, including through interactive web applications.

## 1 Introduction

Spatio-molecular organization in neuronal tissue reflects patterns of cellular composition and circuit connectivity that underlie fundamental structure-function relationships in the brain. Mapping patterns of gene expression onto brain topography using spatially-resolved transcriptomics (SRT) has facilitated the ability to glean novel biological insights into the molecular mechanisms that link structure to function, and better understand how these relationships change over time, development or disease progression (1–3). In the human brain, these techniques have enabled molecular delineation of cortical laminae beyond classic histological definitions (4,5), identified novel cell types and their topographical organization in the noradrenergic locus coeruleus (6), and described differentiation trajectories at multiple gestational stages of the human fetal brain (7). These technologies have been deployed to better understand disease states in the brain, including mapping the local microenvironment of multiple sclerosis lesions (8,9) and infiltration patterns of malignant glioblastoma (10).

Comprehensive spatio-molecular mapping of the human hippocampus (HPC) is critical to understand how its unique organizational structure supports many fundamental biological processes (11,12). The HPC includes the dentate gyrus (DG) and the *cornu ammonis* (CA) regions, subdivided into CA1-4, each of which contains specialized cell types and distinct laminar organization. The organization of these specialized neuronal cell types into neuropil-enriched layers, including the DG molecular layer (ML), stratum lucidum (SL), and stratum radiatum (SR), support well-defined functions within the canonical HPC circuitry. The trisynaptic loop, which supports various features of learning, memory and the stress response, initiates with inputs from the entorhinal cortex (ENT), which traverse from DG to CA3 to CA1, culminating with a relay to the subiculum (SUB), the major output nucleus of the HPC (13,14). Output circuits from the SUB to numerous cortical and subcortical regions control important cognitive and motivated behaviors (11,15), and are implicated in multiple neuropsychiatric and neurodevelopmental disorders (16,17).

Defining the molecular composition of cell types that play specialized roles in HPC circuit function is a prerequisite to targeting their function for therapeutic interventions. However, available transcriptomic profiles generated using single-nucleus RNA-sequencing (snRNA-seq) from postmortem human HPC tissue (18–21) lack important spatial information and do not retain cytosolic or synaptic transcripts (22). Additionally, many existing transcriptomic datasets have focused specifically on the DG given its importance in development and aging (23–28), or have inconsistently sampled across HPC subregions, resulting in cellular composition differences between donors. To investigate gene expression at cellular resolution across the human HPC, we curated postmortem human tissue specimens with well-defined HPC neuroanatomy that systematically encompassed all subfields and sampled across the structure’s diverse longitudinal axis. We deployed a discovery-based experimental design using a well-validated platform to measure gene expression transcriptome-wide in a spatial context, and generated paired snRNA-seq data from adjacent tissue sections to investigate gene expression at cellular resolution. To maximize the utility and value of this data resource for the community we sourced HPC tissue from the same adult, neurotypical brain donors for which we recently provided comprehensive, paired SRT and snRNA-seq data in the human dorsolateral prefrontal cortex (dlPFC) (5).

We used spot-level deconvolution and non-negative matrix factorization (NMF) to integrate the SRT and snRNA-seq datasets, providing novel biological insights about the molecular organization of HPC cell types, cell states, and spatial domains in the human brain. We also deployed new computational strategies for overcoming inherent limitations of postmortem human tissue by incorporating functional molecular data in model organisms. Specifically, we used the human gene expression data to identify latent factors, then incorporated existing rodent datasets that feature information on circuit connectivity and neural activity induction to make predictions about axonal projection targets and likelihood of ensemble recruitment in spatially-defined cellular populations of the human HPC. The ability to infer functional roles for human cell types within the context of intact circuitry has profound potential for understanding how function in the human brain is disrupted in disease.

## 2 Results

### 2.1 Experimental design and overview of profiling human hippocampus

We obtained tissue blocks of postmortem human brain containing anterior HPC from the same 10 neurotypical adult brain donors used in our previous study investigating spatial gene expression in the dlPFC (**Figure 1A, Supplementary Table 1**) (5). We used histological staining to determine neuroanatomical orientation and inclusion of major subfields. Then, we used the 10x Genomics Visium Spatial Gene Expression (Visium-H&E) and 3’ Single Cell Gene Expression platforms to generate paired SRT maps and snRNA-seq data of the human HPC. For each donor, we first collected 1-2 100μm cryosections for snRNA-seq, and then mounted 10μm cryosections across multiple Visium capture areas to profile all major subfields (CA1-4, DG, SUB, **Supplementary Table 2**). We performed Visium-H&E for *n*=36 capture areas (2-5 per donor) (**Figure 1B-C, Extended Data Fig. 1**). We visualized cytoarchitecture from the H&E images and confirmed expected gene expression patterns of *SNAP25* and *MBP* across HPC structure (**Figure 1C**, **Extended Data Fig. 1**). We applied standard preprocessing and quality control workflows to all *n*=36 capture areas to remove low-quality spots (**Extended Data Fig. 2**, **Extended Data Fig. 3**). We retained 150,917 spots from *n*=36 capture areas, hereafter referred to as the SRT dataset.

**Figure 1.**
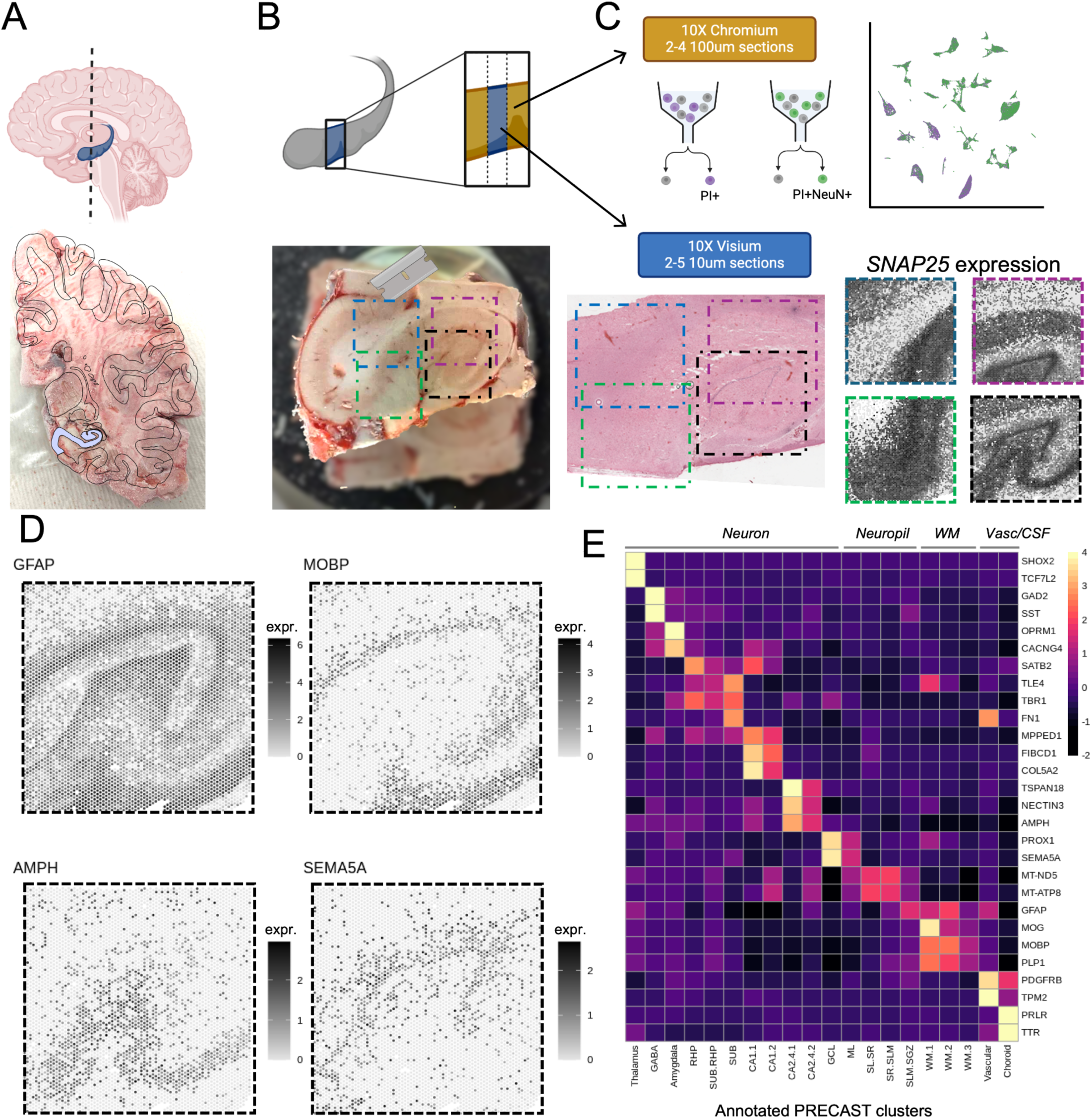
Experimental design to generate paired single-nucleus RNA-sequencing (snRNA-seq) and spatially-resolved transcriptomics (SRT) data in the human hippocampus. (**A**) Postmortem human tissue blocks containing the anterior hippocampus were dissected from 10 adult neurotypical brain donors. (**B**) Tissue blocks were scored and cryosectioned for snRNA-seq assays (gold), and placement on Visium slides (Visium-H&E, blue). (**C**) Top: Tissue sections (2-4 100μm cryosections per donor) from all ten donors were collected from the same tissue blocks for measurement with the 10x Genomics Chromium 3’ gene expression platform. For each donor, two samples were generated, one sorted based on propidium iodide (PI, purple) and the second sorted based on PI+ and NeuN+ (green). Replicate samples were collected from three donors for a total of *n*=26 total snRNA-seq libraries. Bottom: 10μm tissue sections from all ten donors were placed onto 2-5 capture areas to include the extent of the HPC (*n*=36 total capture areas), for measurement with the 10x Genomics Visium-H&E platform. Orientation was verified based on expression of known marker genes. (**D**) Canonical marker genes were identified as spatially variable genes using `nnSVG` (29). Spots are colored by log_2_ normalized counts. (**E**) SRT data was clustered using `PRECAST` (30) with *k*=18 and clusters were annotated (columns) based on expression of known marker genes (rows). Cluster groupings indicated at the top of the heatmap define which clusters contributed to the broad domains of Neuron, Neuropil, white matter (WM), and vascular/ cerebrospinal fluid cell-enriched (Vasc/CSF). RHP: retrohippocampus, SUB: subiculum, CA2.4: *cornu ammonis* (CA) regions 2 through 4 (CA2, CA3, CA4), GCL: dentate gyrus granule cell layer, ML: dentate gyrus molecular layer, SL: stratum lucidum, SR: stratum radiatum, SLM: stratum lacunosum-moleculare, SGZ: dentate gyrus subgranular zone. Hippocampal region abbreviations are also presented in **Supplementary Table 2**.

Following tissue collection for SRT experiments, we collected a second set of 1-2 100μm cryosections from each donor, which were pooled with the cryosections that were collected earlier for snRNA-seq (**Figure 1C, Methods 4.5**). We collected two populations of nuclei for snRNA-seq: a NeuN+ population to enrich for neurons, and a PI+ population. Following sequencing, we applied standard processing and quality control (QC) workflows to remove empty droplets, doublets, and poor-quality nuclei (**Extended Data Fig. 4**, **Extended Data Fig. 5**, **Methods 4.6**). After processing, we retained 75,411 high-quality nuclei across all 10 donors.

### 2.2 Molecular profiles for spatial domains in the human hippocampus using spatially-resolved transcriptomics

To define hippocampal tissue domains in SRT data, we used a data-driven, multi-sample workflow to generate spatially informed clusters. We employed `nnSVG` (29) to select 2000 spatially variable genes (SVGs) which were then used as input to generate unsupervised spatially-aware clusters with `PRECAST` (30) (**Extended Data Fig. 6A, Methods 4.4**, **Figure 1D, Supplementary Table 3**). `nnSVG` and `PRECAST` were chosen for their ability to improve on non-spatially aware feature selection and clustering methods and their computational efficiency (31,32).

We considered a range of clustering resolutions (*k*) and used Akaike Information Criterion (AIC) to focus our search on clusters generated with *k*=16, *k*=17, and *k*=18 (**Extended Data Fig. 6B**). These clusters corresponded with known hippocampal regions based on marker gene expression, and all resolutions identified a sparsely distributed GABAergic cluster (**Extended Data Fig. 6C**). In addition to probing the expression of known marker genes, we evaluated the fidelity of `PRECAST` clustering output to known anatomical organization by comparison to histological annotations of all *n*=36 capture areas (**Methods 4.4**, **Extended Data Fig. 7**). This helped us to select `PRECAST` *k*=18 clusters due to the presence of a single cluster that roughly mapped to both the stratum lacunosum-moleculare (SLM) and the subgranular zone (SGZ) of the dentate gyrus (**Extended Data Fig. 6D-E**).

`PRECAST` clusters were readily split into broad spatial domains of neuron cell-body rich regions (Neuron), neuropil-rich regions (Neuropil), white matter (WM), and vasculature and choroid plexus (Vasc/CSF) based on nuclei density and expression of key non-neuronal genes like *MOBP*, *TTR*, and *PDGFRB* (**Figure 1E)**. Neuronal `PRECAST` clusters have unique expression of canonical marker genes that agree with anatomical organization (**Figure 1E, Extended Data Fig. 6C**). Due to similar gene expression, clusters corresponding to CA2-4 and CA1 were collapsed into their respective domains (**Extended Data Fig. 8A-B**). Neuropil clusters were annotated based on a combination of gene expression and anatomical localization to the stratum lucidum (SL), stratum radiatum (SR), dentate gyrus molecular layer (ML), SLM, and SGZ (**Extended Data Fig. 8C-D**).

The final spatial domains (**Extended Data Fig. 9**, **Supplementary Table 2**) identify the dentate gyrus granule cell layer (GCL), the *cornu ammonis* (split into the CA1 and CA2-4), the subiculum (SUB), cortical neurons of other retrohippocampal regions (RHP), and a domain that appears to be transitional between the SUB and RHP (SUB.RHP). Our spatial domains further include neuropil regions ML, SL.SR, SR.SLM, and SLM.SGZ. We also assigned annotations for populations of cell types that are not traditionally associated with fixed HPC anatomical regions: GABA, vasculature (Vasc), and choroid plexus (CP). As the SLM and SGZ are comprised of distinct cell types and exhibit distinct functions, we expected these two domains would cluster independently. However, we reasoned that including a grouping that corresponded to the combined transcriptional variation across these two HPC domains provided us a better opportunity to elucidate gene expression differences present in these regions than if we utilized *k*=16 clusters, where 4825 of the 7820 SLM.SGZ spots belonged to cluster 5 (*n*=37003 spots total) and the remaining 38% of SLM.SGZ were distributed across many clusters. Further, the combinatorial nature of the SLM.SGZ spatial domain was consistent with other neuropil regions SL.SR and SR.SLM, where the *k*=18 PRECAST clusters were most accurately annotated to a combination of adjacent regions.

Within the RHP and SUB.RHP spatial domains, we identified spatially-restricted spots corresponding to thalamus (33,34) and amygdala (21,35,36) based on anatomy and gene expression (**Methods 4.4**, **Extended Data Fig. 10**). With the exception of one capture area that was entirely amygdala tissue, these spots were not very abundant and were present in only a few donors. We examined the ability of alternative clustering approaches to separate the SLM.SGZ cluster and to distinguish thalamus and amygdala. We explored `GraphST`, which utilizes graph-based autoencoders (**Extended Data Fig. 11**), and `BayesSpace`, which we used in previous studies (5,24) (**Extended Data Fig. 12**). We compared these strategies with the spatial domains annotated from `PRECAST` clusters and observed strong agreement in most domains. However, the distribution of the neuropil clusters did not correspond to the canonical organization of neuropil layers in the HPC, and neither thalamus- or amygdala-specific clusters were present (**Extended Data Fig. 11**, **Extended Data Fig. 12**, **Extended Data Fig. 13**). Spots corresponding to the thalamus and amygdala were removed from downstream differential expression (DE) analysis to focus on HPC and RHP transcriptomes.

Next, we asked if we could discern transcriptomic profiles for HPC spatial domains using DE analysis. We first pseudobulked data by capture area, aggregating all UMIs within all spots for each of the 16 spatial domains (**Figure 2B**). Principal components analysis (PCA) revealed that top components of variation stratified the pseudobulked data by spatial domain (**Figure 2C**, **Extended Data Fig. 14**) and that more variance was explained by spatial domain compared with other covariates such as capture area, donor, Visium slide, sex, or age (**Extended Data Fig. 15**).

**Figure 2.**
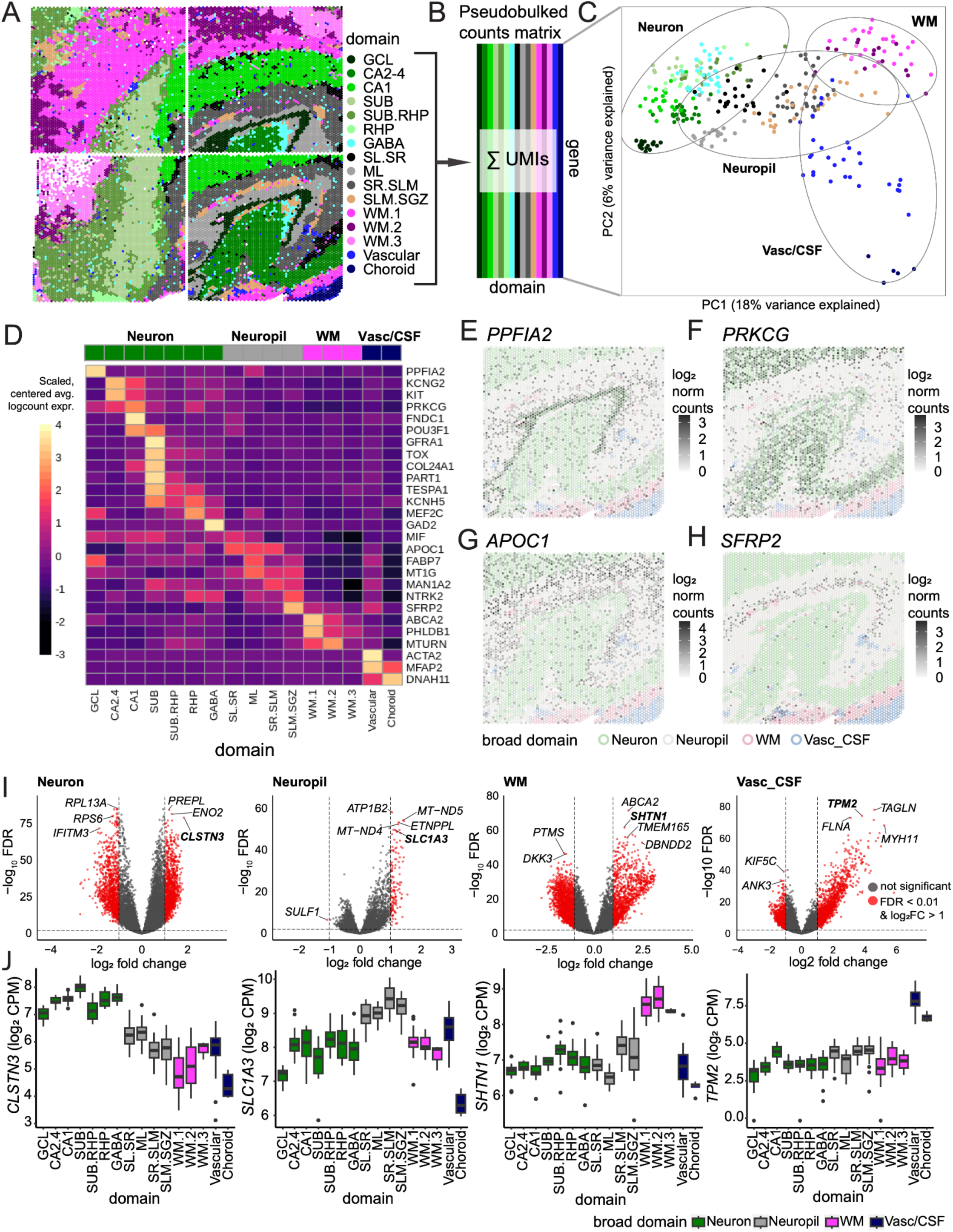
Spatial domain annotation and differential expression (DE) in the human hippocampus using spatially-resolved transcriptomics data. (**A**) Integrated and merged spot plot of four Visium capture areas from the same donor (Br3942) with spots colored by the 16 spatial domains annotated from *k*=18 `PRECAST` clusters. (CA: cornu ammonis) CA1.1/CA1.2 were collapsed to CA1 and CA2-4.1/CA2-4.2 were collapsed to CA2-4. See **Supplementary Table 2** for abbreviations. (**B**) Schematic illustrating pseudobulking approach, which collapses the spot-level to spatial domain-level data within each capture area, summing the total unique molecular identifiers (UMIs) for each new group. (**C**) First two principal components (PCs) of the pseudobulked samples with colors corresponding to spatial domains and labeled ovals representing four broad domains: neuron cell body-enriched (greens and light blue (GABA)), neuropil-enriched (greys), white matter (WM)-enriched (purples), vasculature- and cerebrospinal fluid cell-enriched (Vasc/CSF) (dark blue). (**D**) Heatmap showing differentially expressed gene (DEG) expression (rows) across the spatial domains (columns). Grouping across the top shows broad domain annotations. (**E**) Spot plot showing spatial expression of DEG *PPFIA2*, a known marker for the granular cell layer (GCL). Spots are filled by log_2_ normalized counts. Spot borders are colored by broad domain. (**F**) Spot plot showing expression of DEG *PRKCG*, which is known to be enriched in CA1-4 domains. Spots are filled by log_2_ normalized counts. Spot borders are colored by broad domain. (**G**) Spot plot showing expression of DEG *APOC1*, a known astrocyte cell marker which is enriched in molecular layer (ML), stratum lucidum(SL)-stratum radiatum (SR), and SR-stratum lacunosum moleculare (SLM) domains. Spots are filled by log_2_ normalized counts. Spot borders are colored by broad domain. (**H**) Spot plot showing expression of DEG *SFRP2*, which was specifically increased in the SLM-subgranular zone (SGZ) domain. Spots are filled by log_2_ normalized counts. Spot borders are colored by broad domain. (**I**) Volcano plots illustrating results from DE analysis for each broad-level domain, with log_2_ fold change on the *x*-axis and false discovery rate (FDR) adjusted, -log_10_ transformed *p*-values on the *y*-axis. Genes colored red pass both FDR and log_2_ fold change thresholds (FDR adjusted *p*-value < 0.01 and log_2_ fold change > 1). Top DEGs for each broad domain grouping are labeled. (**J**) Boxplots showing expression of DEGs for neuron-enriched regions (*CLSTN3*), neuropil-enriched regions (*SLC1A3*), white matter regions (*SHTN1*), and vascular/CSF regions (*TPM2*). Each data point represents a pseudobulked sample. Spatial domains are on the *x*-axis and normalized gene expression in log_2_ counts per million (cpm) is on the *y*-axis. Boxes are colored by broad cluster grouping.

To identify differentially expressed genes (DEGs) across the spatial domains, we employed a ‘layer-enriched’ linear mixed-effects modeling strategy to test for differences between one domain versus all others (adjusted for sex, age, and sequencing batch) as previously described (4,5) (**Extended Data Fig. 16**, **Extended Data Fig. 17**, **Supplementary Table 4**). We used the results from this enrichment model to identify DEGs for spatial domains (**Figure 2D**). We confirmed canonical marker genes for spatial domains, such as *PPFIA2* for GCL (**Figure 2E**), *PRKCG* for pyramidal neuron layers CA1-4 (**Figure 2F**), and *MOBP* for WM. We also identified several novel marker genes, particularly with respect to HPC neuropil-enriched domains. These included *APOC1*, enriched in ML, SR, and SL (**Figure 2G**), and *SFRP2*, which marks SLM/SGZ (**Figure 2H**).

Using the same modeling strategy, we asked if we could identify genes specific to broad domains (**Supplementary Table 5**). This analysis identified genes enriched in each of the 4 broad domains (**Figure 2I**), and revealed marker genes that were enriched across all spatial domains within each broad domain, including *CLSTN3* for neuron-rich areas, *SLC1A3* for neuropil-enriched domains, *SHTN1* for WM domains, and *TPM2* for vascular/CSF domains (**Figure 2J**). Altogether, these findings reveal widespread differences in spatial gene expression across the HPC corresponding with both discrete subregions and broad domains.

### 2.3 Molecular profiles for cell types in the human hippocampus using paired single-nucleus RNA-sequencing

We used snRNA-seq data from all ten donors to define human HPC cell types (**Figure 1C**). After QC and batch correction (**Methods 4.6, Extended Data Fig. 18**), we applied graph-based clustering and identified *k*=60 fine-grained clusters (**Extended Data Fig. 19**). Preliminary annotations of these clusters using known marker gene expression identified many non-neuronal cell types (3 CP, 4 vascular, 1 ependymal, 3 astrocyte, 2 oligodendrocyte, 2 oligodendrocyte precursor, and 3 immune cell clusters, **Extended Data Fig. 19F**) and many neuronal cell types (11 GABAergic, 1 Cajal Retzius, 5 GC, 7 HPC pyramidal, 12 RHP pyramidal, 5 amygdala, and 1 thalamus cluster, **Extended Data Fig. 20**). We next sought to expand our understanding of these cell types and improve cluster annotations by performing DE analysis using the pseudobulk approach (**Supplementary Table 6**). Informed by the significant results from DE analysis, we generated 24 cell types (**Figure 3A-B**) that did not exhibit donor-specific bias (**Extended Data Fig. 21**).

**Figure 3.**
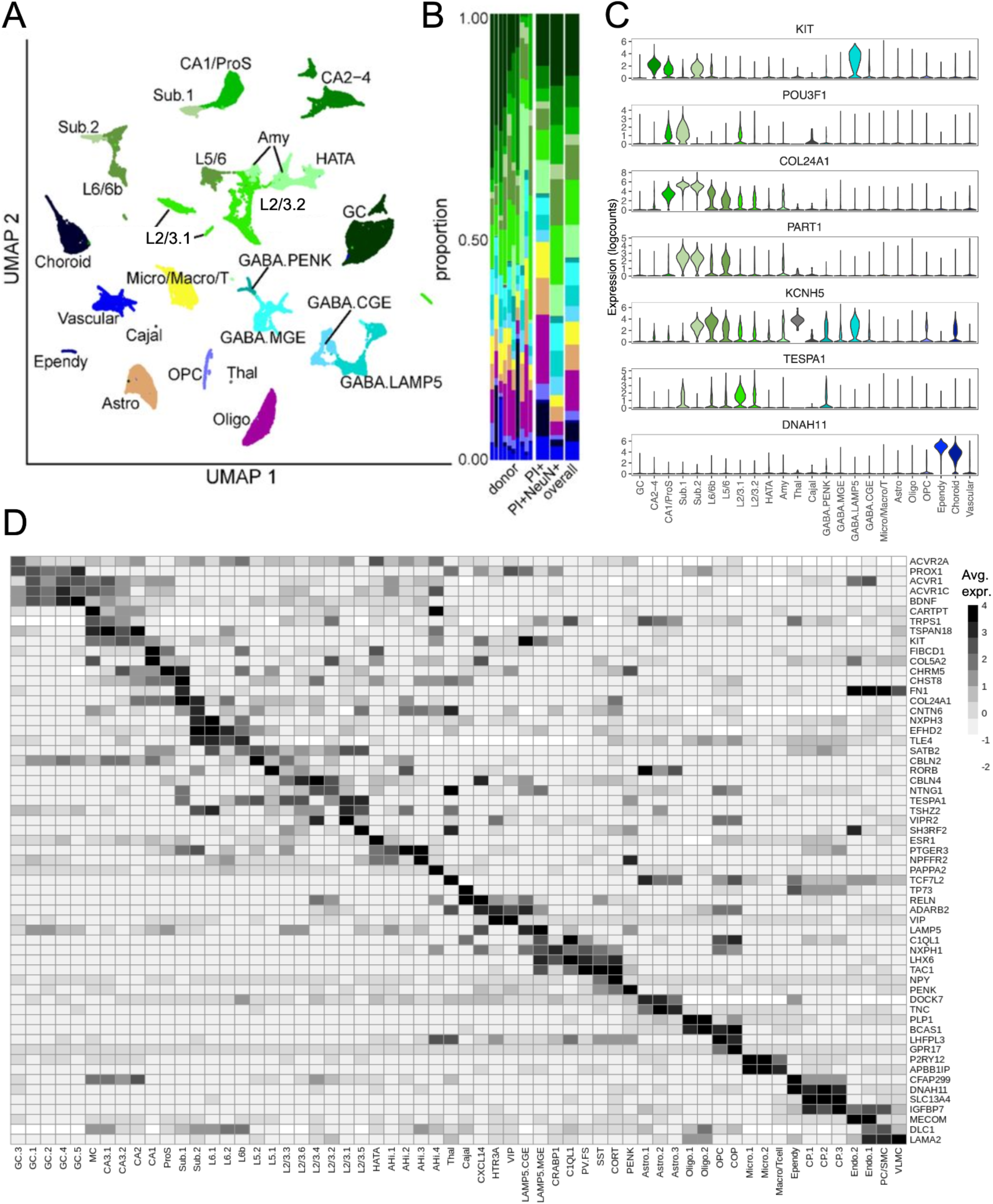
Cell type identification and differential expression (DE) in the human hippocampus using single-nucleus RNA-sequencing (snRNA-seq). (**A**) Uniform manifold approximation and projection (UMAP) representation of the snRNA-seq dataset. Individual nuclei are represented as points that are colored and labeled by cell type. GC: dentate gyrus granule cell, CA2-4: *cornu ammonis* (CA) regions 2 through 4 (CA2, CA3, CA4), CA1/ProS: CA1 and prosubiculum, Sub: subiculum, HATA: hippocampus-amygdala transition area, Amy: amygdala, Thal: thalamus, Cajal: Cajal-Retzius cells, GABA.PENK: *PENK* positive GABAergic neurons, GABA.MGE: medial ganglionic eminence-derived GABAergic neurons, GABA.LAMP5: *LAMP5* positive GABAergic neurons, GABA.CGE: central ganglionic eminence-derived GABAergic neurons, Micro/Macro/T: microglia and macrophages and T-cells, Astro: astrocytes, Oligo: oligodendrocytes, OPC: oligodendrocyte progenitor cells, Ependy: ependymal cells. Cell type abbreviations are also presented in **Supplementary Table 2**. (**B**) Left: Stacked bar plots of cell types indicated in (**A**) showing proportions of nuclei for each donor (columns). Right:Stacked bar plots of cell types indicated in (**A**), with columns indicating nuclei grouped by sort strategy (propidium iodide (PI)+ or PI+NeuN+), and across the overall dataset (all nuclei). (**C**) Violin plots showing log_2_ normalized expression (*y*-axis) of select significant genes identified with spatial domain differential expression (DE) analysis and *n*=60 snRNA-seq cluster DE analysis. Nuclei are grouped based on cell types (*x*-axis) for improved visibility and fill color also corresponds to cell type as indicated in (**A**). (**D**) Heatmap showing a selection of significant genes (*y*-axis) from snRNA-seq DE analysis across all *n*=60 clusters (columns). Heatmap is colored by mean log_2_ normalized counts. Additional cluster abbreviations not defined in (**A**) are AHi: amygdala-hippocampal region, CXCL14: *CXCL14* positive GABAergic neurons, HTR3A: *HTR3A* positive GABAergic neurons, VIP: *VIP* positive GABAergic neurons, CRABP1: *CRABP1* positive GABAergic neurons, C1QL1: *C1QL1* positive GABAergic neurons, PV.FS: *PVALB* positive fast-spiking GABAergic neurons, SST: *SST* positive GABAergic neurons, CORT: *CORT* positive GABAergic neurons, COP: committed oligodendrocyte precursor, CP: choroid plexus tissue, Endo: endothelial cells, PC/SMC: pericytes and smooth muscle cells, VLMC: vascular lepto-meningeal cells. Cell cluster abbreviations are also presented in **Supplementary Table 2**.

Using these differentially expressed genes (DEGs), we were further able to resolve minute differences between cell classes (**Figure 3D**), including distinguishing oligodendrocyte progenitor cells (OPCs) from committed oligodendrocyte precursors (COPs) (37) (**Extended Data Fig. 22A**). Some DEGs reflected specific cellular functions shared across cell types. For instance, both ependymal cells and CP cells showed enrichment of genes related to motile cilia, such as *DNAH11*, which circulate CSF through ventricles (**Extended Data Fig. 22A**). Although we were able to find genes that distinguished RHP clusters, our ability to make biological inferences from these DEGs was impaired by the limited characterization of HPC-proximal RHP pyramidal subtypes (**Extended Data Fig. 22B**). We were similarly restricted in our interpretation of the amygdala clusters, as the amygdala was not systematically targeted and was only present in a few donors (**Extended Data Fig. 22C**). Our results demonstrate the continuum of gene expression in HPC pyramidal neurons while still distinguishing gene expression that was unique to the SUB (**Extended Data Fig. 22D**). The GABAergic nuclei belonged to several different subtypes, highlighting the mosaic nature of the HPC inhibitory neuron population (**Extended Data Fig. 22E, F**). Intriguingly, we observe subsets of GCs that express distinct forms of activin receptors (*ACVR1*, *ACVR2A*, *ACVR1C*), suggesting a stable heterogeneity within the DG GCL during adulthood (**Extended Data Fig. 22G**). These results were further validated using an alternative analysis that implemented `scVI` (38,39) where we found consistent cell types (**Methods 4.6**, **Extended Data Fig. 24**, **Extended Data Fig. 25**).

We then leveraged our paired SRT data in two ways. First, to ascertain the similarity of snRNA-seq clusters with spatial domains detected with SRT, we correlated the gene-level enrichment model *t*-statistics for each of the snRNA-seq superfine clusters (**Supplementary Table 6**) with the *t*-statistics from the enrichment model of the SRT domains (**Supplementary Table 4**). These analyses were strongly correlated across many clusters and domains, indicating appropriate assignment of hippocampal nuclei despite lacking *a priori* spatial information (**Extended Data Fig. 23**). Gene expression profiles for some cell types and spatial domains overlapped strongly (e.g. GCs with GCL; CP cells with CP domain), while other domains contained multiple cell types (e.g., GABAergic cell types correlated strongly with the GABA domain). We then compared snRNA-seq DEGs with those identified between spatial domains. We find multiple DEGs significant in both data modalities, including canonical and novel markers of pyramidal cell types (**Figure 2D**, **Figure 3C**). We identify that *KIT*, a CA3 marker in SRT, is expressed in pyramidal cell types from CA3 and CA1 but also in a specific GABAergic population. *POU3F1* is expressed at similar levels in CA1 and Sub.1 clusters in snRNA-seq data, but is restricted to the CA1 in SRT. Additional genes that were expressed in the SUB spatial domain included *COL24A1* and *KCNH5*, which were expressed in the continuum of Sub.1, Sub.2, and RHP clusters, and *PART1* and *TESPA1*, which exhibited more cluster-specific expression in the Sub.1, Sub.2, and RHP clusters.

The HPC is implicated in many pathological conditions, but the specific cell types and spatial domains driving these associations remain unclear (16,40–42). To identify cell types and spatial domains associated with genetic risk for diseases and disorders, we used stratified linkage disequilibrium score regression (S-LDSC) (43–45) to calculate enrichment of heritability of multiple polygenic traits, including neuropsychiatric and non-psychiatric conditions (46–61). S-LDSC regression coefficients represent contribution of a given annotation (e.g. spatial domain or cell type) to heritability of a given trait (**Methods 4.7**). We performed S-LDSC regressions for each superfine cell type. We observed many expected associations, including enrichment of Alzheimer’s disease (AD) risk in microglia (47) (**Extended Data Fig. 26B**). We then performed the same analysis for each spatial domain, and observed enrichment of genetic risk for schizophrenia (SCZ) in the CA1 domain, and multiple disorders, including SCZ, in the RHP (**Extended Data Fig. 26A**). We thus sought additional integration strategies for our snRNA-seq and SRT data to enable better understanding of the diversity of HPC cellular populations and their spatial organization.

### 2.4 Subfield-specific changes in cell type composition using patterns shared between snRNA-seq and SRT data

We capitalized on the variation and the diversity of cellular populations in the snRNA-seq data (**Figure 3**) to further explore cell type composition across spatial domains using spot-level deconvolution algorithms (62). While many algorithms have been developed to predict cell type proportions within individual Visium spots using single cell reference data, these methods have not yet been comprehensively benchmarked across various brain regions or in heterogeneous tissues. To provide a robust spot deconvolution reference dataset for postmortem human anterior HPC, we generated data using the Visium Spatial Proteogenomics assay (Visium-SPG). Visium-SPG, which replaces H&E histology with immunofluorescence (IF) staining to label proteins of interest, was performed on tissue from two donors (one male and one female) with particularly clear anatomical orientation.

With Visium-SPG, we labeled cell type-specific proteins: NEUN (marking neurons), OLIG2 (marking oligodendrocytes), GFAP (marking astrocytes), and TMEM119 (marking microglia) (**Extended Data Fig. 27A**). Multispectral fluorescence imaging was performed followed by the standard Visium protocol to generate gene expression libraries from each tissue section, as previously described (5). Following QC (**Extended Data Fig. 27B-E**), we used `RcppML` for transferring spatial domain labels identified in our larger SRT dataset into the Visium-SPG data (**Extended Data Fig. 27F**). We then benchmarked `cell2location` (63), `RCTD` (64), and `Tangram` (65) on their ability to predict cell type composition at multiple snRNA-seq classification depths (**Methods 4.8, Extended Data Fig. 28**, **Extended Data Fig. 29).** Performance of different deconvolution tools was evaluated by comparing algorithm-predicted cell compositions to cell identities determined from fluorescence intensity in each IF channel, providing an orthogonal validation (**Methods 4.8, Extended Data Fig. 30, Extended Data Fig. 31, Extended Data Fig. 32, Extended Data Fig. 33, Extended Data Fig. 34, Extended Data Fig. 35**, **Extended Data Fig. 36**, **Extended Data Fig. 37**) similar to our previous work (5). Given the challenge of deconvolving transcriptionally fine cell types (**Extended Data Fig. 29**), these benchmark analyses confirmed `RCTD` showed the most consistent performance at both mid and fine resolution across all cell types and samples (**Extended Data Fig. 36**, **Extended Data Fig. 37**). We thus applied `RCTD` to the SRT dataset (**Figure 4A-B**, **Extended Data Fig. 29**, **Extended Data Fig. 37**).

**Figure 4.**
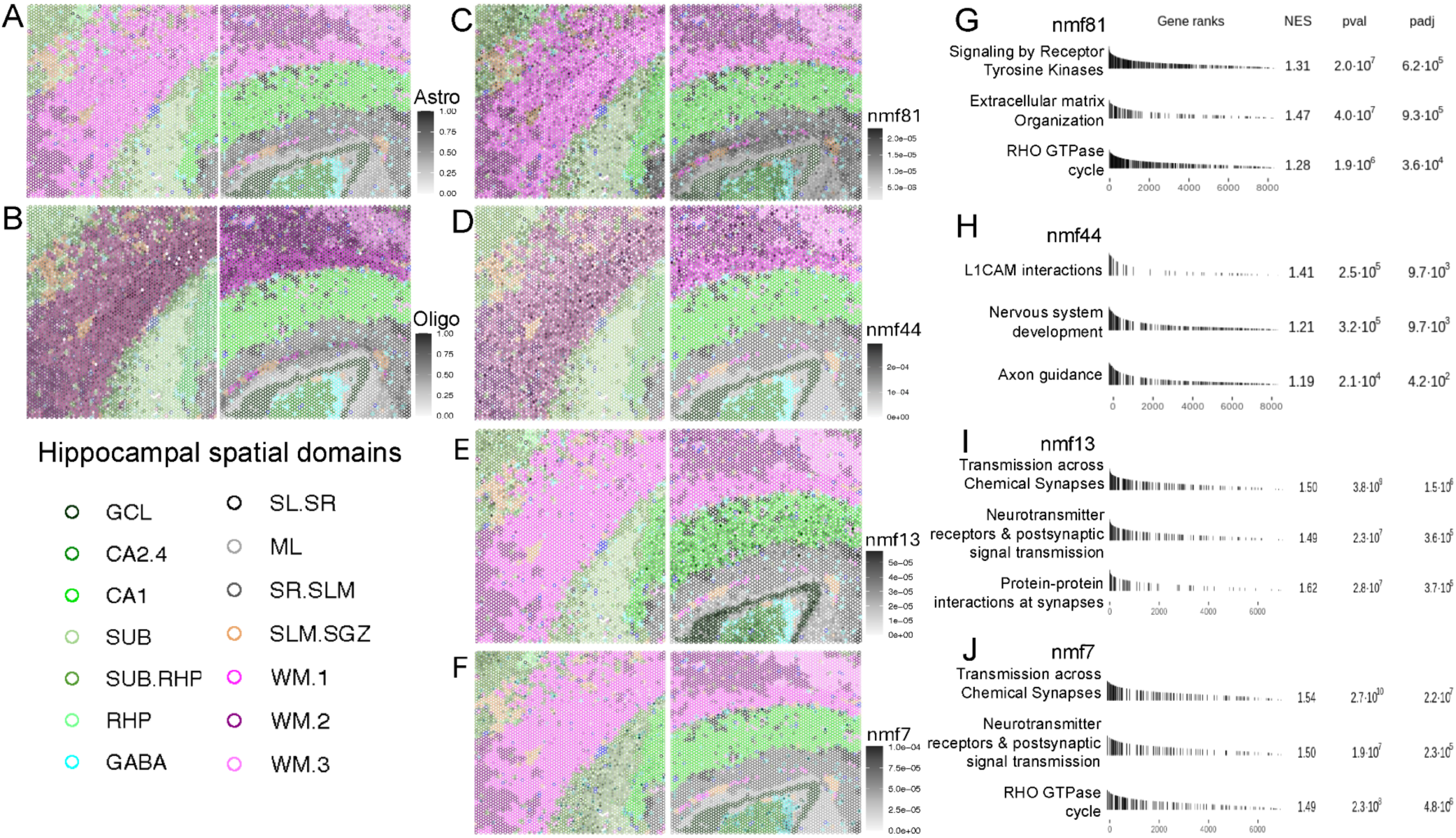
Non-negative matrix factorization (NMF) reveals cell type heterogeneity and biologically relevant pathways in hippocampal subfields compared to RCTD deconvolution results. (**A**) `RCTD` prediction of proportion of astrocytes (Astro, fill color) per Visium spot in example capture areas from donor Br3942. Spot borders are colored by spatial domain. See **Supplementary Table 2** for abbreviations. (**B**) `RCTD` prediction of proportion of oligodendrocytes (Oligo, fill color) per Visium spot in example capture areas from donor Br3942. Spot borders are colored by spatial domain. (**C**) Spot plots displaying spot-level weights (fill color) for NMF pattern nmf81 (specifically elevated in astrocyte snRNA-seq clusters) in example capture areas from donor Br3942. Spot borders are colored by spatial domain. (**D**) Spot plots displaying spot-level weights (fill color) for NMF pattern nmf44 (specially elevated in oligodendrocyte snRNA-seq clusters) in example capture areas from donor Br3942. Spot borders are colored by spatial domain. (**E**) Spot plots displaying spot-level weights (fill color) for NMF pattern nmf13 (elevated in neuronal snRNA-seq clusters) in example capture areas from donor Br3942. Spot borders are colored by spatial domain. (**F**) Spot plots displaying spot-level weights (fill color) for NMF pattern nmf7 (elevated in neuronal snRNA-seq clusters) in example capture areas from donor Br3942. Spot borders are colored by spatial domain. (**G**) Gene set enrichment analysis (GSEA) table for nmf81 showing that genes with stronger weights contributed to the significant enrichment of biological pathways (like “Extracellular matrix organization”) associated with astrocytes. NES: normalized enrichment score, pval: *p*-value, padj: FDR adjusted *p*-value. (**H**) GSEA table for nmf44 showing that genes with stronger weights contributed to the significant enrichment of biological pathways (like “L1CAM interactions” and “Axon guidance”) associated with oligodendrocytes. (**I**) GSEA table for nmf13 showing that genes with stronger weights contributed to the significant enrichment of biological pathways highly relevant to neuronal signaling. Investigation of the specific genes contributing to these terms indicate that transcriptional variation captured by nmf13 is highly relevant to excitatory postsynaptic response. (**J**) GSEA table for nmf7 showing that genes with stronger weights contributed to the significant enrichment of biological pathways highly relevant to neuronal signaling. Investigation of the specific genes contributing to these terms indicate that transcriptional variation captured by nmf7 is highly relevant to the structure and maintenance of inhibitory postsynaptic specializations.

While spot-level deconvolution methods are valuable, most methods rely on *a priori* classification of data into discrete clusters, thereby overlooking transcriptional heterogeneity within cell types. Coupling non-negative matrix factorization (NMF) with transfer learning provides an intuitive approach to identify continuous gene expression patterns that may be more functionally relevant (66). NMF decomposes high-dimensional data into *k* lower-dimensional latent spaces (67) **(Methods 4.9, Extended Data Fig. 38**). When applied to snRNA-seq data, latent factors represent distinct gene expression patterns that can define biological or technical variation. Patterns may be specific to cell types, cell states, or other biological processes (68,69). Transfer learning can map identified patterns onto independent datasets (69), allowing for discernment of patterns shared across species and data modalities.

Here, we used NMF to define gene expression patterns within the snRNA-seq data and then mathematically project the patterns to transfer the weights onto SRT data in order to probe the spatial organization of these patterns (**Extended Data Fig. 38**). Using the `RcppML` R package (67), we performed NMF on the normalized snRNA-seq counts matrix at rank *k*=100 to define 100 NMF patterns (**Extended Data Fig. 39**, **Extended Data Fig. 40A**, **Supplementary Table 9**). To enable comparisons between the contributions of specific genes and nuclei to each pattern, pattern weights are normalized such that both nuclei-level and gene-level NMF weights are interpreted as proportions of a given pattern. We used transfer learning to predict spot-level weights for the 100 patterns in the SRT data to allow for spatial visualization and analysis of patterns (**Methods 4.9, Extended Data Fig. 38**). We limited further investigation to patterns that were found in >1050 spots and thus removed 32 patterns based on limited expression in the SRT dataset (**Extended Data Fig. 40C**). We checked for donor-specific effects in the remaining patterns, identified one pattern that encapsulated the donor origin of CP tissue, and removed an additional two that corresponded to donor sex (**Methods 4.9**, **Extended Data Fig. 41**, **Extended Data Fig. 42**, **Extended Data Fig. 43**)

We found 47 patterns that corresponded strongly to specific cell types and spatial domains, which is unsurprising given that they are built from snRNA-seq gene expression (**Extended Data Fig. 40A,D**). We examined an astrocyte-dominant pattern (nmf81, **Figure 4C**) and an oligodendrocyte- and WM-dominant pattern (nmf44, **Figure 4D**). Among the several NMF patterns corresponding to oligodendrocyte snRNA-seq clusters (**Extended Data Fig. 40A**), we highlighted nmf44 due to the specific increase in weights in observations annotated to WM (**Extended Data Fig. 44A,C,E**). In contrast, nmf81 was specific to the astrocyte snRNA-seq clusters but spot-level weights were distributed throughout spatial domains, consistent with the ubiquitous presence of astrocytes throughout the HPC (**Extended Data Fig. 44B,D,F**). We compared the spot-level NMF weights with orthogonally-validated spot-level deconvolution results and found strong agreement in spatial organization that corresponded to strong correlations for between `RCTD`-predicted cell type and NMF pattern weights (nmf81-Astro = 0.904, nmf44-Oligo = 0.876) (**Figure 4A-D, Extended Data Fig. 36**). This confirms that both NMF and spot-level deconvolution algorithms can be useful to identify cell type identity across data modalities.

An advantage to NMF over spot-level deconvolution is the ability to identify individual genes that contributed to the construction of each pattern without DE analysis, which first requires clustering to define cell populations. For a given NMF pattern, the gene-level weight is representative of the amount of transcriptional variation that is attributable to each gene. While the top ten genes weighted to each NMF pattern are informative (**Supplementary Table 10**), the number of top-weighted genes required to explain a substantial proportion of the transcriptional variation captured by the NMF patterns is variable and difficult to define for the majority of patterns (**Extended Data Fig. 45**). Therefore, we performed gene set enrichment analysis (GSEA) which utilizes the ordinal rank of gene weights to examine whether genes contributing to the transcription patterns captured in nmf81 and nmf44 were consistent with known biology. Genes including *NRXN1* (70), *NCAN* (71), and *PLEC* (72) contributed to the significant enrichment of “Extracellular matrix organization” in the top astrocyte-related nmf81-weighted genes (**Figure 4G**). In oligodendrocyte-related pattern nmf44, top-weighted genes include *ANK3* (73) and *DLG1* (74), which contributed to the significant enrichment of “L1CAM interactions” and “Axon guidance” terms (**Figure 4H**). These results provide evidence that NMF patterns capture gene expression programs that are attributable to known biological function.

Another advantage of NMF is the ability to capture transcriptional variation beyond cell type definition. We observed 19 patterns that were present across cell types and spatial domains (**Extended Data Fig. 40A,D**). We examined two of these “general” patterns (nmf13 and nmf7), which both exhibited increased spot-level weights in multiple neuronal cell types and hippocampal spatial domains (**Figure 4E-F, Extended Data Fig. 46A-F**). These two patterns were anti-correlated in most cell types and spatial domains (**Extended Data Fig. 46G-H**), yet GSEA identified that both nmf13 and nmf7 were significantly enriched for genes participating in “Transmission across Chemical Synapses” and “Neurotransmitter receptors and postsynaptic signal transmission” (**Figure 4I-J**). However, the individual genes that contributed to the identification of these neuronal pathways represent distinct neuronal functions. The biological function of top nmf13-weighted genes, like *CAMKK1* (75), *LRFN2* (76), and *DLGAP3* (77), suggests that nmf13 represents gene expression patterns highly relevant to excitatory postsynaptic response. In contrast, the biological function of top nmf7-weighted genes, like *GABRA1*, *KIF5A* (78–80), and *DYNLL2* (81), led us to conclude that nmf7 represents gene expression patterns highly relevant to the structure and maintenance of inhibitory postsynaptic specializations. Given the domain-restricted increase in nmf13 (GCL, CA2-4, CA1 domains and associated cell types **Extended Data Fig. 46A,C,E**) and nmf7 (SUB, RHP, GABA, and associated cell types **Extended Data Fig. 46B,D,F**), these data indicate that NMF can identify subfield-specific differences in neuronal structure and function, in addition to cell type composition.

### 2.5 NMF captures activity-dependent transcription programs

We identified two NMF patterns (nmf91 and nmf20) that captured stimulus-dependent transcriptional programs (**Methods 4.9**, **Extended Data Fig. 47**). Many of the genes highly weighted to nmf91 were immediate early genes that are transiently and robustly expressed immediately following neuronal activity (e.g., *FOS*, *JUN*, *NR4A1* (82)) (**Extended Data Fig. 47B**). For nmf20, three genes stand out as being most highly weighted: *SORCS3*, *HOMER1*, and *PDE10A,* which is highly expressed in medium spiny neurons of the nucleus accumbens (83) (**Extended Data Fig. 47D**). Notably, while a previous study was unable to detect *PDE10A* RNA expression in the human HPC, our snRNA-seq data is consistent with rodent data that *Pde10a* transcripts and PDE10A protein are expressed in the HPC (84,85) (**Extended Data Fig. 47I-K**). In addition to known roles in modulating synaptic response to activity (86–89), these highly-weighted nmf20 genes exhibit stimulus-dependent expression in response to neuronal activity (*SORCS3* (90)) and dopamine signaling (*PDE10A* (91)), while *HOMER1* expression can be constitutive or activity-dependent for different splicing isoforms (92) (**Extended Data Fig. 47D**). To further investigate the ability of NMF patterns to capture gene expression programs relevant to neuronal activity, we transferred the NMF patterns from our human snRNA-seq dataset onto a mouse snRNA-seq dataset of HPC neurons activated by electroconvulsive seizures (ECS) or HPC neurons under control conditions (Sham) (93). As we observed when transferring human snRNA-seq nuclei-level weights into human SRT data, NMF patterns mathematically projected onto mouse snRNA-seq nuclei were both “general” and cell type-specific (**Extended Data Fig. 48B**). Although some patterns exhibited differential nuclei weights based on ECS status (**Extended Data Fig. 48C**), all patterns exhibited cell type specificity, and we observed no NMF patterns that were more highly weighted to the ECS or sham group across all cell types.

We found that nmf91 and nmf20 were more highly weighted onto nuclei from GCs in the ECS condition than the sham-activated group, supporting our hypothesis that these patterns capture activity-dependent transcriptional programs (**Figure 5A**). We further found that many of the top-weighted genes to nmf91 and nmf20 were significantly increased in ECS GCs compared to sham. For nmf91, these genes included many canonical activity-dependent transcripts (**Figure 5B**). Although the genes highly weighted in nmf20 are less clear, the high weight and increased expression of genes including *BDNF* (94) and *SORCS3* and *SORCS1,* which are mediators of intracellular BDNF signaling (*90,95*), indicate that this pattern may represent genes involved in secondary activity response (**Figure 5C**). We investigated if this pattern was more specifically enriched in synapse-rich domains and found that 98.0% of spots with non-zero nmf20 weights were found in the Neuron (91.0%) or Neuropil (7.0%) broad domains, while 68.9% of spots with non-zero nmf91 weights occurred in these regions (42.8% in Neuron, 26.1% in Neuropil).

**Figure 5:**
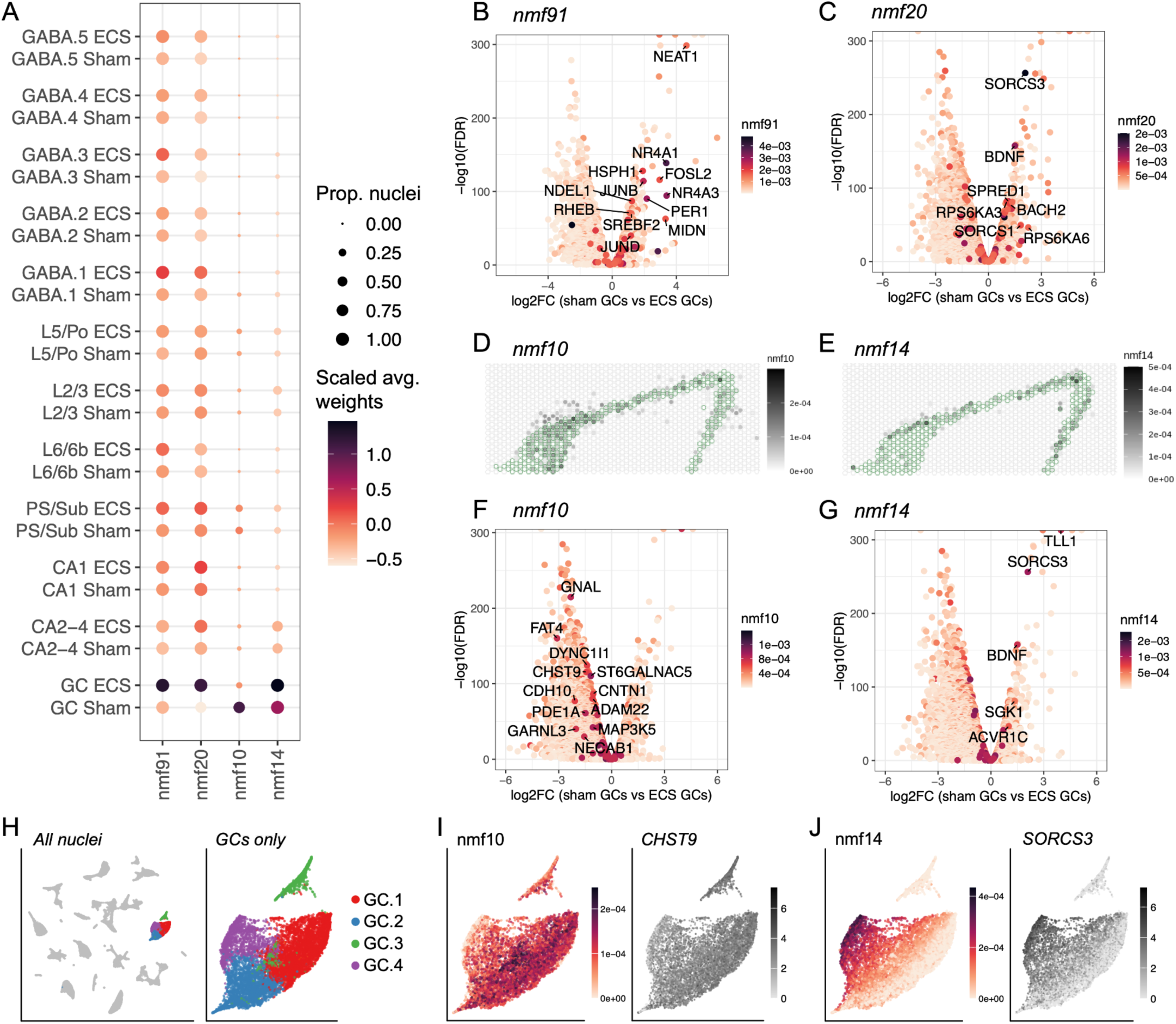
Non-negative matrix factorization (NMF) captures transcriptional programs relevant to neuronal activity. (**A**) Select NMF patterns projected onto a mouse snRNA-seq dataset of hippocampal neurons activated by electroconvulsive stimulation (ECS) or hippocampal neurons under control conditions (Sham) (*y*-axis, by cell type). These patterns (*x*-axis) exhibit altered dentate gyrus granule cell (GC) nuclei weights (dot color, scaled and averaged) between ECS and sham conditions. Dot size indicates the proportion of *y*-axis group with non-zero pattern weight for the given *x*-axis value. CA2-4: *cornu ammonis* (CA) regions 2 through 4 (CA2, CA3, CA4), PS/Sub: prosubiculum and subiculum neurons, L5/Po: layer 5 and polymorphic layer. Differential expression (DE) analysis was performed on mouse GC nuclei, testing for differences in the expression by activity condition. For all volcano plots (**B-C, F-G**), the *y*-axis presents the -log_10_ false discovery rate (FDR)-adjusted *p* value. For all volcano plots (**B-C, F-G**), the *x*-axis presents log_2_ fold change (FC), where negative values indicate greater expression in sham-activated GCs and positive values indicate greater expression in ECS GCs. (**B**) Volcano plot of DE results tested on genes with non-zero nmf91 weights. Points are colored by gene-level nmf91 weight. Gene names are shown for genes with nmf91 weight >0.0015, log_2_FC>1, and -log_10_(FDR)>30. (**C**) Volcano plot of DE results tested on genes with non-zero nmf20 weights. Points are colored by gene-level nmf20 weight. Gene names are shown for genes with nmf20 weight >0.00065, log_2_FC>1, and -log_10_(FDR)>30). Spot plots isolating the dentate gyrus granule cell layer spatial domain (GCL, green outlined spots) demonstrate the differing spatial organization of (**D**) nmf10 and (**E**) nmf14 weights in an example capture area from donor Br3942. Spot fill indicates spot-level NMF pattern weight. (**F**) Volcano plot of sham vs. ECS DE results from mouse snRNA-seq tested on genes with non-zero nmf10 weights. Points are colored by gene-level nmf10 weight. Text is shown for genes with nmf10 weight >0.00065, log_2_FC< -1, and -log_10_(FDR)>30. (**G**) Volcano plot of sham vs. ECS DE results from mouse snRNA-seq tested on genes with non-zero nmf14 weights. Points are colored by gene-level nmf14 weight. Text is shown for genes with nmf14 weight >0.0005, log_2_FC> 1, and -log_10_(FDR)>30. (**H**) Uniform manifold approximation and projection (UMAP) plot of (left) all nuclei present in our human snRNA-seq highlighting the GC clusters. Right: Zoomed UMAP plot of only GC nuclei from our human snRNA-seq dataset with color indicating cluster identity. (**I**) UMAP plot of human GC nuclei showing (left) nmf10 nuclei-level weights and (right) log_2_ normalized counts of highly-weighted nmf10 gene *CHST9*. (**J**) UMAP plot of human GC nuclei showing (left) nmf14 nuclei-level weights and (right) log_2_ normalized counts of highly-weighted nmf14 gene *SORCS3*.

In further examining the mouse ECS dataset, we found two additional NMF patterns of interest. Mouse GC nuclei exhibited specific increases in nmf14 and nmf10 weights, and these patterns were also associated with GC clusters and the GCL in our human datasets (**Extended Data Fig. 40A,D**). While nmf14 weights were mildly increased in ECS GCs, nmf10 weights were robustly decreased in ECS GCs compared to sham-activated GCs (**Figure 5A**). In our human data, nmf10 was weighted to spots throughout the GCL while spots highly weighted to nmf14 exhibited a more restricted localization to the superficial GCL (**Figure 5D,E**).

Examination of genes that are both highly weighted to nmf10 and decreased in ECS GCs revealed that this pattern likely represents transcriptional programs contributing to synaptic adhesion and the cementing of established synapses (cell adhesion molecules *CNTN1*, *ADAM22*; calcium-dependent cell adhesion molecules *FAT4*, *CDH10*; production of cell adhesion molecules *ST6GALNAC5*, *CHST9*) (**Figure 5F**). The decrease in expression of these genes in ECS GCs likely facilitates synaptic remodeling following neuronal activity. In contrast, we identified GC.4 cluster markers *BDNF* and *ACVR1C* (**Extended Data Fig. 22G**), as well as *SORCS3* and *SGK1* as key genes with higher nmf14 weights that were increased in ECS GCs (**Figure 5G**). Given the functional importance of *BDNF* (96), *ACVR1C* (*97*), *SORCS3* (86), and *SGK1* (98,99) in neuronal response to activity and synaptic plasticity, we hypothesize that nmf14 represents gene expression patterns that promote synaptic scaling. The elevated weight of nmf14 in ECS GCs and the superficial GCL suggests that a subset of GCs may be uniquely poised to promote activity-dependent synaptic scaling. Examining the nuclei-level weights of nmf10 and nmf14 and the expression of top-weighted nmf10 and nmf14 genes indicated that the GC.4 cluster fits the criteria for poised GCs (**Figure 5H-J**).

These data indicate that NMF can identify activity-regulated gene expression in the context of cell type-specific recruitment. Activity-regulated transcription is intimately related to physiological function, recruitment of cellular ensembles, and synaptic connectivity, all of which fundamentally contribute to cellular behavior. However, these properties are not often considered when annotating cell types in transcriptomic studies. Here, we used NMF to translate information from animal models into human brain datasets to make predictions across species about functional properties of human cell types. This approach can also be extended to understand cell type function in the context of circuit connectivity in the HPC.

### 2.6 NMF and transfer learning enable spatial mapping and inference of circuit connectivity of pyramidal neurons in the hippocampal formation

Both the HPC and RHP contain multiple classes of pyramidal neurons with distinct molecular profiles (18,19,100), physiological properties (101), axonal projection targets (102), and spatial organization (103–105). Although the RHP has been anatomically classified as a transition zone between the three layer allocortex and five layer agranular cortex, it has historically been difficult to delineate distinctions between neuronal populations from RHP subregions (e.g. subiculum vs entorhinal cortex (ENT)) with snRNA-seq alone. Indeed, our snRNA-seq analysis identifies 7 pyramidal neuron clusters from the HPC and 12 from the RHP. We aimed to better understand these clusters by examining the spatial organization of SRT spots weighted to specific NMF patterns. We find that Sub.1 cluster-specific nmf40 and Sub.2 cluster-specific nmf54 exhibit distinct laminar organization *in situ* (**Figure 6A**). Recent studies indicate that the diversity of excitatory neuron types within the rodent SUB and RHP correspond to multiple efferent target regions (13,14). We asked if performing NMF pattern transfer onto another dataset where the axonal projection targets of HPC and RHP neurons are known could elucidate whether these different subicular layers corresponded to different circuits. We utilized data from a recent study in mice that coupled retrograde viral circuit tracing with single-cell methylation sequencing (snmC-seq) (106). We mapped NMF patterns from the human snRNA-seq data onto mouse snmC-seq data after filtering to excitatory neurons obtained from the HPC and RHP (*n*=2004 nuclei) (**Extended Data Fig. 38, Methods 4.9**). We removed patterns that corresponded to <45 nuclei, which included nmf54/Sub.2 (**Extended Data Fig. 49A**), and were thus unable to investigate if these two cell types corresponded to SUB populations that targeted distinct brain regions. However, we found that patterns specific to CA1, SUB, and RHP cell types/clusters mapped to mouse HPC neurons with distinct efferent targets (**Figure 6B**). These results recapitulated known projections from the SUB to the thalamus (107) and hypothalamus (108), and from the ENT to the HPC and prefrontal cortex (109). We thus further investigated whether the NMF patterns corresponding to distinct snRNA-seq RHP clusters exhibited spatially restricted organization, and if the spatial information could provide insight into the cell type identity of RHP clusters.

**Figure 6.**
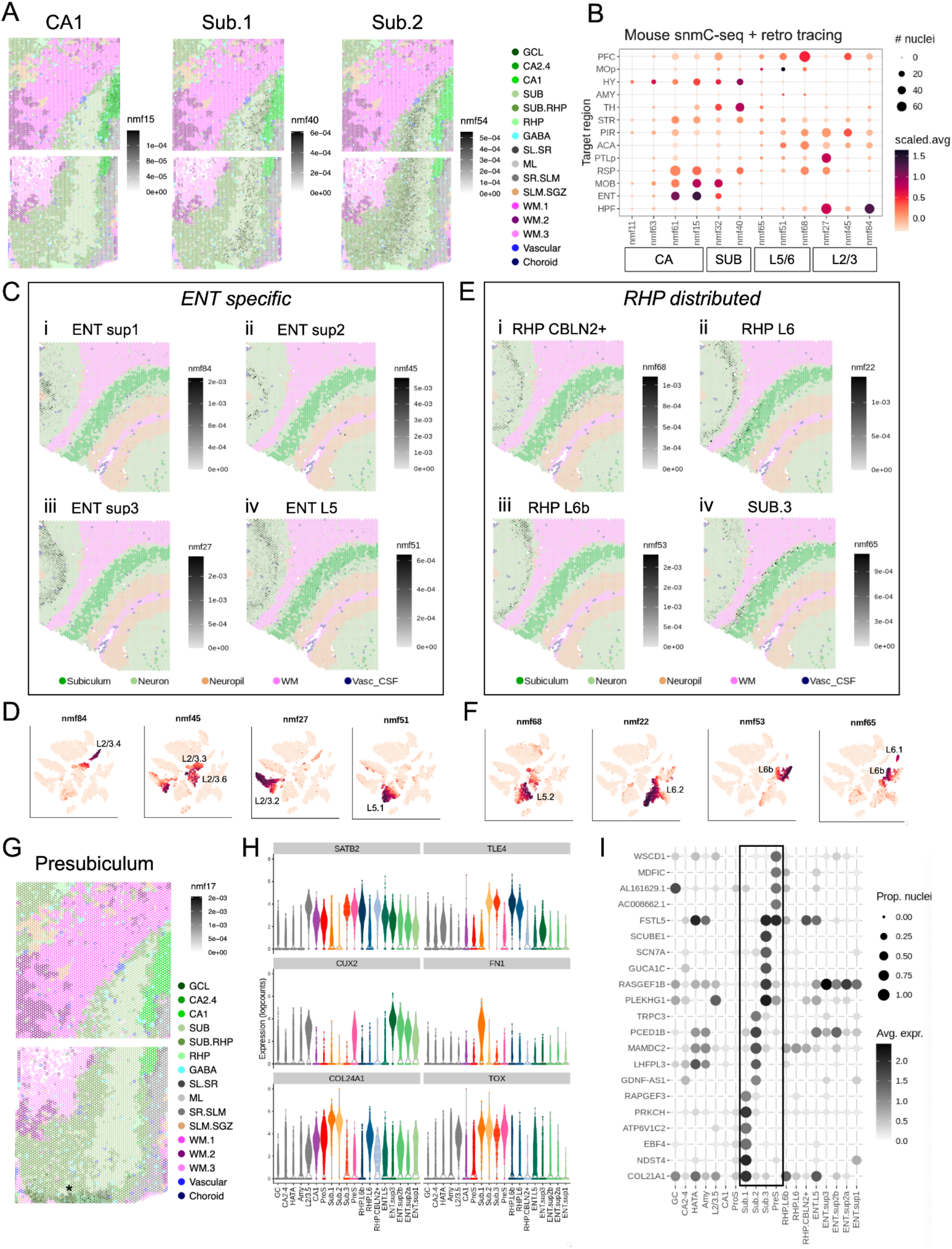
Non-negative matrix factorization (NMF) reveals a continuum of pyramidal cell types across the retrohippocampus (RHP). (**A**) Spot plots for example donor Br3942 highlight the laminar organization of nmf40 (specific to the subiculum Sub.1 snRNA-seq cluster) and nmf54 (specific to the subiculum Sub.2 snRNA-seq cluster), as well as the clear distinction from the CA1. Spots are filled by the spot-level weights for the indicated NMF pattern, with scales corresponding to the maximum spot-level weight of any spot in the SRT dataset. Spot border color indicates spatial domain. See **Supplementary Table 2** for abbreviations. (**B**) Dot plot of mouse single-cell methylation sequencing (snmC-seq) with retroviral tracing (*n*= 2004 nuclei) after label transfer of NMF patterns (106). Rows indicate nuclei axonal projection target region obtained from retroviral tracing experiments, NMF patterns corresponding to HPC and RHP cell types are present as columns. Dot size indicates the number of nuclei with non-zero pattern weights and dot color indicates the scaled, average weight for the given pattern. HPF: hippocampal field, ENT: entorhinal cortex, MOB: main olfactory bulb, RSP: retrosplenial cortex, PTLp: posterior parietal cortex, ACA: anterior cingulate cortex, PIR: piriform cortex, STR: striatum, TH: thalamus, AMY: amygdala, HY: hypothalamus, MOp: primary motor cortex, PFC: prefrontal cortex. For spot plots **C** and **E** an example capture area from donor Br2743 was used. The capture area is mirrored from what is present in **Extended Data Fig. 9** to have the superficial-to-deep organization presented in a left-to-right manner. Spot border color indicates broad domain, with the addition of the subiculum to illustrate the lack of overlap with the subiculum. The subiculum was labeled by thresholding nmf40 and nmf54 weights (**Extended Data Fig. 49C-D**). Spots are filled by spot-level weights for the indicated NMF pattern, with scales corresponding to the maximum spot-level weight of any spot in the SRT dataset. (**C**) Spot plots of entorhinal cortex (ENT) specific NMF patterns. Patterns are shown from the most superficial to the least, with patterns specific to L2/3 snRNA-seq clusters presented in i-iii. Panel iv shows the ENT-specific L5 pattern. (**D**) *t*-distributed stochastic neighbor embedding (TSNE) plots of pyramidal nuclei from the snRNA-seq dataset colored by the NMF patterns in (**C**). Patterns are shown from the most superficial to the least (left to right). The snRNA-seq cluster(s) enriched for specific NMF patterns are labeled. (**E**) Spot plots of retrohippocampus (RHP) NMF patterns that exhibit spot-level weights distributed across the ENT and subicular complex. Patterns are shown from the most superficial to least. Panel iv exhibits low weights in the deep ENT with enriched nmf65 spot-level weights immediately adjacent to subiculum-labeled spots. The specificity of nmf65 to the deep subiculum is explored in **Extended Data Fig. 49E-F**., lending to the classification of this pattern as a third subiculum pattern (SUB.3). (**F**) t-distributed stochastic neighbor embedding (TSNE) plots of pyramidal nuclei from the snRNA-seq dataset colored by the NMF patterns in (**E**). Patterns are shown from the most superficial to the least (left to right). The snRNA-seq cluster(s) enriched for specific NMF patterns are labeled. (**G**) Spot plots for example donor Br3942 exemplify the anatomical location of nmf17 to the presubiculum, indicated by the asterisk. Spot border color indicates spatial domain. Spots are filled by the indicated NMF pattern, with scales corresponding to the maximum nmf17 spot-level weight of any spot in the SRT dataset. An alternative visualization of nmf17 mapping to the presubiculum for all donors is presented in **Extended Data Fig. 51**. (**H**) Violin plots show snRNA-seq log_2_ normalized counts (*y*-axis) across HPC and RHP clusters (*x*-axis) for traditional cortical layer markers *SATB2*, *TLE4*, and *CUX2*. Also shown is canonical subiculum marker *FN1*, and *COL24A1* and *TOX*, new subiculum markers explored in this manuscript. Select clusters have been renamed following SRT-based verification of spatial organization: Sub.3 (formerly L6.1), PreS (presubiculum, formerly L2/3.1), RHP.L6 (formerly L6b), RHP.L6 (formerly L6.2), RHP.CBLN2+ (formerly L5.2), ENT.L5 (formerly L5.1), ENT.sup3 (formerly L2/3.2), ENT.sup2b (formerly L2/3.3), ENT.sup2a (formerly L2/3.6), ENT.sup1 (formerly L2/3.4). The revised cell type abbreviations are also presented in **Supplementary Table 2**. (**I**) A dot plot detailing the results of focused differential expression analysis that was performed on snRNA-seq data to elucidate novel genes that distinguish between the superficial subiculum (Sub.1), middle subiculum (Sub.2), deep subiculum (Sub.3), and the PreS. Dot size indicates the proportion of nuclei in each cluster (column) with non-zero expression for each gene (row). Dot color indicates average log_2_ normalized gene counts. The box highlights the novel differentially expressed genes. Violin plots of these results are shown in **Extended Data Fig. 52**.

We discovered that NMF patterns corresponding to three superficial pyramidal neuron clusters and one L5 pyramidal neuron cluster (nmf84, nmf45, nmf27, and nmf51) were spatially restricted to the distal RHP, suggesting these populations are ENT-specific (**Figure 6C-D**). By leveraging NMF to represent transcriptional patterns shared across the snRNA-seq dataset and the SRT dataset, we observed that pyramidal neuron clusters were labeled by a series of non-overlapping patterns that exhibit stereotyped laminar organization throughout the transition of ENT to SUB (**Figure 6C-F**). Unlike more superficial patterns, those corresponding to deeper layer neurons (nmf68, nmf22, nmf53, and nmf65) were not restricted along the transverse axis (CA1-distal to CA1-proximal) but were present throughout the SUB-ENT transition (**Figure 6E**). One pattern (nmf65) ran along the border of WM and SUB, adjacent to the middle layer of SUB spots highlighted by nmf54 (**Figure 6E**). Given that nmf65 labeled two snRNA-seq clusters (L6.1 and half of L6b) (**Figure 6F**), we hypothesized that one of those cellular populations may represent the deep SUB rather than ENT. We compared the expression of SUB marker genes and top NMF pattern genes in L6.1 nuclei and in L6b nuclei with non-zero nmf65 weights (**Extended Data Fig. 49E-F**). The consistent expression of SUB DEGs like *TOX*, *TSHZ2*, and *ZNF385D* in L6.1 suggests this snRNA-seq cluster comprises the deep SUB layer *in situ*, and we thus re-labeled this cluster as Sub.3.

We were especially surprised to find that the L2/3.1 cluster-specific pattern nmf17 exhibited a dramatically different spatial location than the ENT-specific L2/3 patterns across all donors (**Figure 6G**, **Extended Data Fig. 50**, **Extended Data Fig. 51**). The differential distribution of this pattern in samples from more anterior donors (including Br6423 and Br2743), in which nmf17 expresses alongside the SUB, and more posterior donors (including Br3942 and Br8325), in which nmf17 is expressed at the curve of the SUB, suggests that nmf17 expression follows previous anatomical description of the presubiculum (109,110). Supporting this hypothesis, nmf17 is expressed in an island-like pattern in some donors, again consistent with anatomical studies. However, to our knowledge, there are no molecular markers that have been identified for the presubiculum, highlighting the ability of our approach to uncover novel biology even in the well-studied HPC.

Based on the spatial information, we refined our snRNA-seq cluster annotations to reflect the discrete populations of superficial ENT pyramidal neurons, the three layers of SUB pyramidal neurons, a population of presubiculum neurons, and a group of deep pyramidal neurons present across the RHP continuum. The expression of common cortical layer markers may or may not be sufficient to identify these populations in future experiments (**Figure 6H**). The use of the canonical SUB marker gene *FN1* may be helpful to isolate superficial SUB neurons (Sub.1), but our data indicate that *COL24A1*, which was identified as a SUB DEG in both SRT and snRNA-seq datasets (**Figure 3C**), identifies superficial and middle SUB neurons. *TOX*, another DEG for the SUB in both SRT and snRNA-seq data, appears to be a more general subicular complex indicator, labeling the presubiculum as well (**Figure 6H**). The spatial organization of cluster L2/3.5 was not examined in this study, as the corresponding specific NMF pattern (nmf78) mapped to fewer than 1050 spots. However, it appears likely this cluster also represents SUB complex neurons based on the expression of *TOX* and *TSHZ2* (**Extended Data Fig. 22B**). We performed additional differential expression analysis focused on discovering new gene markers that could be useful for future studies looking to annotate subicular complex neurons (**Methods 4.9**). We set a very robust threshold and identify multiple genes that, in combination with traditional regional and laminar markers, will be useful for future research (**Figure 6I**, **Extended Data Fig. 52**).

### 2.7 Data Access and Visualization

This atlas of integrated single cell and spatial transcriptomic data is the most comprehensive map of the human HPC to date. To enable exploration of these rich data resources, we created multiple interactive data portals, available at research.libd.org/spatial_hpc/. The pseudobulked snRNA-seq and SRT data are freely available through *iSEE* apps (111) to allow users to visualize expression for genes of interest across cell types and spatial domains. To facilitate exploration of SRT data in anatomical context, we merged all Visium capture areas from each donor for visualization of high resolution images and corresponding gene expression data, NMF patterns, and spot-level deconvolution results at the donor level using a web-based interactive tool, `Samui Browser` (112) (10 donors, *n*=36 Visium-H&E capture areas; 2 donors, *n*=8 Visium-SPG capture areas. Raw and processed data are available through Gene Expression Omnibus (GEO) (113) under accession GSE264624 to facilitate access for methods development and in depth analysis. Finally, to make the data accessible for broader neuroscience community, we created an ExperimentHub Data package (https://bioconductor.org/packages/humanHippocampus2024) within the Bioconductor framework such that processed snRNA-seq and SRT data can be downloaded conveniently.

## 3 Discussion

By integrating SRT and snRNA-seq data, we generated a comprehensive transcriptomic atlas of the adult human hippocampus (HPC). We characterized the molecular organization of the human HPC with both spatial and cellular resolution, identifying discrete spatial domains and a rich repertoire of HPC cell types. Spot level deconvolution and non-negative matrix factorization (NMF) uncovered gene expression patterns representing cell type-specific molecular signatures. NMF enabled us to also capture gene expression profiles shared across biological processes, such as synaptic signaling (**Figure 4**). Integration of these patterns with cross-species functional genomics data (**Figure 5**) provides spatial context to behaviorally-relevant cell types, cell states, and molecular pathways. To molecularly define the organization of the subicular complex, we leverage cross-species neuronal circuit data and anatomical insight from our SRT dataset (**Figure 6**).

Measuring transcriptomic changes in response to experimental manipulations is not possible in human postmortem brain tissue, while functional studies in rodents that can test causality of cellular and molecular associations may lack direct relevance to human brain function and behavior. Therefore, integrating human transcriptomic data with functional genomics data from animal models has the potential to facilitate interpretation of cellular responses to various stimuli. To illustrate the biological insights that can be gleaned using this approach, we mapped NMF patterns from the human HPC onto snRNA-seq data from a mouse model of induced electroconvulsive seizures (ECS) (93) to identify gene expression patterns that are putatively associated with neural activity in the human brain (**Figure 5**). Identifying and localizing expression patterns associated with activity-regulated genes in the human HPC is important because expression of these genes is critical for recruitment of HPC neuronal ensembles controlling learning, memory and other cognitive processes (114). NMF patterns that reflected processes known to be regulated by activity, including immediate early gene expression and synaptic signaling, were enriched in activated mouse neurons (**Figure 5**). In the rodent, the majority of GCs are relatively silent under both baseline conditions (115) and during exploration of novel environments (116,117). However, a small fraction of GCs rapidly increase their firing activity during behavior, supporting a sparse coding scheme that facilitates precision in the detection of novelty and the animal’s location during learning (118). NMF patterns that differentially map to control versus activated neurons may represent the population of GCs that are more quiescent versus those that are either actively firing or intrinsically primed for activity (119). Indeed, active and quiescent mouse GCs have substantially different transcriptomic profiles (93). Differences in GC activity levels may reflect differences in their ability to be recruited into neuronal ensembles in response to behavioral experiences (120–125). These analyses provide insight into the putative spatial and molecular organization of cellular activity states in the human DG, which is critical to better understand circuit function in the HPC. Circuit activity in the HPC flows from the DG, extending through the pyramidal, subicular and retrohippocampal regions, which have not been well-characterized in the human at molecular scale. Since annotations for the retrohippocampal transition region rely on a priori knowledge of spatial organization and no molecular profiles are available, we were unable to manually annotate subdivisions in this region. While analysis of the snRNA-seq data enabled identification of many individual cell types along this transition region, unbiased clustering strategies were also unable to fully identify and differentiate across subdivisions. However, application of NMF afforded a more biologically meaningful interpretation of these cell types by mapping their spatial organization in the SRT dataset. With this approach we were able to identify novel gene signatures attributable to specific subdivisions of the ENT and subicular complex (**Figure 6**). These findings highlight the future potential of these approaches for incorporating human transcriptomic data with rapidly emerging viral circuit labeling tools, which have recently enabled the profiling of neuron populations in rodent models based on their innervation patterns (126,127).

We demonstrate that NMF is an effective approach to integrate transcriptomics data with other datasets, particularly across species and data modalities. This approach has some limitations. Namely, NMF can be sensitive to initialization values, patterns corresponding with noise may be incorrectly attributed to distinct biological processes, and NMF weights can be scaled differently across different factorizations. Despite these limitations, the illustrated strategy can be iterated upon to map existing and forthcoming datasets to our atlas. NMF is able to transfer continuous patterns of expression, such as cell types (**Figure 4**, **Figure 6**) and transcriptional activity (**Figure 5**), while other tools, including PCA and spot-level deconvolution, are not able to find these gradients of expression. NMF also enabled the integration of our snRNA-seq and SRT data, which leveraged the spatial information provided by the SRT dataset and the cellular resolution of snRNA-seq to enable the discovery of molecular signatures in human tissue. Ultimately, these signatures can be extended with targeted gene panels at single cell resolution for validation, or reverse-translated to test causality in rodent or cellular models. Importantly, snRNA-seq and SRT data from dorsolateral prefrontal cortex (dlPFC) is available from the same donors as those used in this study (5), and emerging computational methods may enable modeling molecular connectivity patterns across this clinically-relevant circuit in the future. This is an important endeavor with significant potential to improve clinical outcomes given strong scientific rationale and precedence for normalizing circuit-level dysfunction to improve symptoms of neuropsychiatric disorders (128–130). Understanding functional dynamics in HPC circuits is important because their dysregulation is implicated in various neuropsychiatric and neurodevelopmental disorders (131,132). Defining the human HPC at cellular resolution with spatial fidelity is necessary for designing molecular approaches that can facilitate more precise circuit manipulation. Ultimately, the identification of unique molecular identities for spatially-organized HPC cell types based on their innervation targets in the human brain is necessary for the development of circuit-specific therapeutics.

In summary, this highly integrated, well-annotated single cell and spatial transcriptomics dataset of the human HPC provides unique biological insight into the molecular neuroanatomy of the hippocampal formation, including activity-dependent transcriptomic profiles in the GCL and molecular definition of the retrohippocampal transition zone between the ENT and SUB. Our innovative approach can be effectively used to map external data to this and other transcriptomic atlases as illustrated. To facilitate wide access to our data, we provided the human HPC snRNA-seq and SRT atlas as a resource to the scientific community through multiple avenues, including interactive web applications for visualization and exploration.

## 4 Methods

### 4.1 Postmortem human tissue samples

Postmortem human brain tissue from neurotypical adult donors of European ancestry were obtained at the time of autopsy following informed consent from legal next-of-kin, through the Maryland Department of Health IRB protocol #12–24, and from the Western Michigan University Homer Stryker MD School of Medicine, Department of Pathology, the Department of Pathology, University of North Dakota School of Medicine and Health Sciences, and the County of Santa Clara Medical Examiner-Coroner Office in San Jose, CA, all under the WCG protocol #20111080. Using a standardized strategy, all donors underwent a comprehensive retrospective clinical diagnostic review to exclude for any lifetime history of psychiatric or substance use disorders. Macro- and microscopic neuropathological examinations were performed, and subjects with evidence of neuropathological abnormalities were excluded. Additional details regarding tissue acquisition, processing, dissection, clinical characterization, diagnose, neuropathological examination, RNA extraction and quality control (QC) measures have been previously published (133). Demographic information for all donors in this study is listed in **Supplementary Table 1**.

### 4.2 Tissue processing and quality control

All hippocampus dissections in this study were performed by the same neuroanatomist (co-author TMH). The hippocampus was dissected as uniformly as possible using anatomical landmarks to include the anterior portion of the hippocampus proper plus the subicular complex (42). Frozen samples were mounted in OCT (TissueTek Sakura) and cryosectioned at -12°C (Leica CM3050s). To ensure high-quality samples, we measured RNA Integrity Number (RIN) in the sample blocks from a subset of donors and ensured that RIN in our dissected tissue samples was comparable to RIN calculated at time of brain donation (**Supplementary Table 1**). 10μm sections were mounted on standard microscopy slides for staining with hematoxylin and eosin (H&E) for orientation and quality control. Following QC, 2-4 100μm cryosections, totalling approximately 50mg of tissue, were transferred into a low-adhesion tube (Eppendorf) kept on dry ice and reserved from the anterior portion of the HPC for snRNA-seq. Then, two 10μm sections were mounted on standard microscopy slides and reserved. The tissue block was then scored using a razor blade, so that tissue sections approximately the size of a Visium capture area could be positioned and placed onto chilled Visium Spatial Gene Expression slides (part number 2000233, 10x Genomics). Following the successful completion of Visium experiments, additional 100μm cryosections were collected for single-nucleus RNAseq. In some cases, multiple Visium experiments were performed to ensure inclusion of relevant subfields. In these cases, distance along the anterior-posterior axis between experiments did not exceed 500μm. To assist in the reassembly of scored tissue sections, H&E staining was performed on one slide adjacent to the scored Visium sections. RNAscope (ACD) was performed on the other to guide histological annotation of canonical subfields, using previously defined markers of dentate gyrus (*PROX1*), CA3 (*NECAB1*), CA1 (*MPPED1*) and SUB (*SLC17A6*).

### 4.3 Spatially resolved transcriptomics (SRT) data generation

Visium Spatial Gene Expression slides were processed as previously described (4,134). To ensure the optimal time duration of exposure to permeabilization enzyme, tissue optimization experiments were performed according to the manufacturer’s protocols (protocol CG000160, revision B, 10x Genomics). Tissue sections were scored to include the dentate gyrus to facilitate orientation, and exposed to permeabilization enzyme for differing time durations. cDNA synthesis was performed using a fluorescently-labeled nucleotide (CG000238, revision D, 10x Genomics). The slide was then coverslipped and fluorescent images were acquired at 10x magnification with a TRITC filter (ex 550nm/em 600nm) on a Cytation C10 Confocal Imaging Reader (Agilent). Following this experiment, 18 minutes was selected as the optimal permeabilization time. For each Visium slide, H&E staining was performed (protocol CG000160, revision B, 10x Genomics), after which slides were coverslipped and high-resolution, brightfield images were acquired on a Leica CS2 slide scanner equipped with a 20x/0.75NA objective and a 2x doubler. Following removal of the coverslips, tissue was permeabilized, cDNA synthesis was performed, and sequencing libraries were generated for all Visium samples following the manufacturer’s protocol (CG000239, revision C, 10x Genomics). Libraries were loaded at 300 pM and sequenced on a NovaSeq System (Illumina) at the Johns Hopkins Single Cell Transcriptomics core according to manufacturer’s instructions at a minimum depth of 60000 reads per spot.

### 4.4 SRT data processing and analysis

#### Visium raw data processing

FASTQ and image data were pre-processed with the 10x `SpaceRanger` pipeline (version 1.3.1) (135). Reads were aligned to reference genome GRCh38 2020-A. For analysis steps, we used R (version 4.3.2, unless otherwise noted) and Bioconductor (version 3.17) for analysis of genomics data (136). Outputs from the `SpaceRanger` pipeline were read into R and stored in a `SpatialExperiment` (137) (SPE) object using the `read10xVisiumWrapper()` from the `spatialLIBD` package (138). For each donor, we first rearranged the order of capture areas. This was done such that overlapping or adjacent capture areas were adjacent to one another when plotted on a grid (2×2 grid for donors with 3 or 4 capture areas, 1×2 or 2×1 grid for donors with 2 capture areas, or 3×2 grid for donors with 5 capture areas, **Extended Data Fig. 1**). After rearrangement, images from each capture area and spatial coordinates in the SPE were rotated to ensure that neighboring regions from each donor were spatially adjacent and correctly oriented to recapitulate the organization of the initial, unscored cryosection as well as possible. We performed these rotations with the `imgData()` function from the `SpatialExperiment` package. For a given H&E, `imgData()` can transform the raster representation of the image data by some user-specified rotation values (in degrees). In our SPE object, we also independently rotated both the pixel coordinates for each spot (stored in the `spatialCoords` slot) and the capture area array coordinates for each spot (stored in `colData` slot as `array_row` and `array_col`). These rotations were performed to correspond with image rotations.

#### H&E Image Processing

The high-resolution images obtained as part of the 10x Genomics Visium protocol were processed using *VistoSeg*, a *MATLAB*-based pipeline that integrates gene expression data with histological data (139). First, the software splits the high-resolution H&E image of the entire slide into individual capture areas (.tif file format) and the capture areas of a single brain sample are rotated accordingly to orient to each other forming the hippocampal structure. The individual tif files are used as input to 1) `Loupe Browser` (10x Genomics) for fiducial frame alignment, and 2) `SpaceRanger` (version 1.3.0) to extract the spot metrics/coordinates. We then used the `VNS()` function from `VistoSeg` to obtain the initial nuclei segmentations. The `refineVNS()` function with adaptive thresholding was used to segment out the more accurate nuclei from background signals. Segmented regions within the size range of 40 to 4,000 pixels were only retained and a watershed was applied to split the nuclei clusters. These final segmentations were used to obtain the nuclei count per Visium spot. We took two approaches for spot level counting with different stringencies: 1) we counted all segmented nuclei whose centroids were within a given Visium spot and 2) we calculated the proportion of pixels within a given Visium spot in which a segmented nucleus was present. A function in *VistoSeg* then uses the spot metrics and coordinates from `SpaceRanger` to integrate the segmented nuclei counts with gene expression data to obtain the number of nuclei per spot.

#### Visium quality control and count normalization

We filtered the data to remove all undetected genes and spots with zero counts. We used the `addPerCellQC()` function from the `scuttle` Bioconductor package to compute and store standard quality control metrics, such as mitochondrial expression rate, library size, and number of detected genes (140,141). We applied a 3x median absolute deviation (MAD) threshold to discard spots with low library sizes and/or low numbers of detected genes (**Extended Data Fig. 2**). Mitochondrial expression rate was not used to define low-quality spots as this metric appeared to correlate with true biological variation (**Extended Data Fig. 3**). Specifically, hippocampal neuropil-enriched layers, such as dentate gyrus molecular layer, stratum radiatum, stratum lucidum, and stratum lacunosum-moleculare, showed enrichment of mitochondrial genes. These regions are characterized by large numbers of synapses and a paucity of neuronal cell bodies. In neurons, mitochondria are known to be enriched at both presynaptic and postsynaptic sites (142,143). After applying these QC steps, the number of genes x the number of spots was 31,483 x 150,917. Counts were normalized by library size and log_2_-transformed using `computeSumFactors()` from `scran` (140) and `logNormCounts()` from `scuttle` (141).

#### Feature selection, spatially variable genes

Highly variable genes do not necessarily capture spatial expression patterns. Thus, for feature selection prior to unsupervised clustering of SRT data, we used the `nnSVG` Bioconductor package (29) to select the top 2,000 spatially variable genes (SVGs) that exhibited expression patterns that varied across the 2D space of the tissue sample (144). The `nnSVG` package uses a nearest-neighbor Gaussian process model and measures up against other SVG detection methods in improving spatial domain detection in RNAseq data, compared to non-spatial feature selection approaches (31). `nnSVG` aims to identify genes with varying expression patterns across different regions within the tissue by estimating gene-specific spatial ranges and retaining detailed spatial correlation information. The model offers linear scalability and efficient handling of large datasets.

We first filtered out genes with fewer than 100 counts across all spots across all capture areas. For each capture area, we also filtered genes with less than 3 counts in at least 0.5% of spots in that capture area. We did not filter out the mitochondrial genome. We then ran `nnSVG` separately on each capture area using the default parameters. After running nnSVG for each sample, we ranked all genes within each capture area by spatial variance. We averaged gene ranks across all capture areas and ranked genes by mean rank. We retained the top 2,000 genes for clustering analysis (∼10% of all genes retained after filtering steps), constraining to genes ranked in the top 1,000 genes in at least 2 different capture areas (Supplementary Table 3, Figure 1D).

#### Unsupervised clustering of spatial transcriptomics data

For spatially-aware unsupervised clustering of SRT data, we used `PRECAST` v1.5 (30). `PRECAST` was chosen based on its ability to leverage spatial information when performing clustering, its use of joint embeddings that allow for integration across samples and batches, and its relatively fast runtime which allows for optimization of the number of desired clusters (32) (**Extended Data Fig. 6A**). We ran `PRECAST` across all donors using *k*=5 through *k*=20. We used the top 2,000 SVGs calculated using `nnSVG`. Following `PRECAST`, we extracted BIC values from each `PRECASTObj` and prioritized *k*=15 through *k*=20 as these had the lowest BIC values (**Extended Data Fig. 6B**). We visualized spatial domain membership for *k*=15 through *k*=20 and found similar results. However, *k*=18 included a spatial domain that appeared to map to SLM and SGZ, which other clusterings lacked (**Extended Data Fig. 6C-E**).

To ensure robustness of spatial domain predictions across computational algorithms, we compared `PRECAST` results with those generated from two other leading clustering methods designed specifically for spatial transcriptomics. We looked at a domain detection algorithm using graph-based autoencoders, `GraphST` (145). Following the `GraphST` tutorial, features were selected by the highly variable gene method and `GraphST` was performed for each individual slide. Samples were integrated with `Harmony` (146) and we generated *k*=16 clusters. The `GraphST` results were somewhat consistent with the `PRECAST` assignments, but generated a very large cluster corresponding to many distinct HPC regions containing pyramidal neurons (**Extended Data Fig. 11**). We believe this could be due to the use of highly variable genes as recommended in the tutorial, rather than an inherent limitation of the `GraphST` model itself. We also compared `BayesSpace` (147) which is similar to `PRECAST` in that it implements a hidden Markov random field but does so within a Bayesian model framework. For `BayesSpace` implementation, we utilized the same set of SVGs input to `PRECAST` to generate PCs that were MNN-corrected for donor identity, and used *k*=18 clusters. The spatial organization of the `BayesSpace` clusters were very similar to that of `PRECAST` (**Extended Data Fig. 12**). Given the similarities and lack of improved ability to distinguish thalamus and amygdala spots (**Extended Data Fig. 13, introduced below**), we used `PRECAST` clusters from *k*=18 to define our HPC spatial domains.

#### Histologically-defined annotations

All histologically-defined annotations were performed by experienced neurobiologists (co-authors EDN, SCP). To annotate SRT data, we built a temporary `shiny` application using `spatialLIBD::run_app()` (4,138). We annotated the following anatomical regions: granule cell layer (GCL), subgranular zone (SGZ), molecular layer (ML), CA4 pyramidal cell layer (PCL-CA4), PCL-CA3, PCL-CA1, subiculum (SUB), stratum oriens (SO), stratum radiatum (SR), stratum lucidum (SL), stratum lacunosum-moleculare (SLM), white matter (WM), choroid plexus (CP), thalamus (THAL), and cortex (CTX) (**Extended Data Fig. 7**). We used marker genes from snRNA-seq studies in both human HPC (18,19) and mouse HPC (13,100). We also used H&E images for histological reference. To identify GCL, we used *PROX1* and *SEMA5A* and were also guided by high levels of hematoxylin staining due to densely packed granule cell bodies. To identify SGZ, we used *SEMA5A*, but not *PROX1*, expression. We also used *GAD1/GAD2* due to the presence of GABAergic neurons in SGZ (148). To identify ML, we noted low UMIs and detected genes in areas between GCL and SLM. To identify PCL-CA4, we used *NECTIN3*, *AMPH*, *SEMA5A*, *SLC17A7*, and the presence of higher UMIs and detected genes compared with adjacent SGZ. For PCL-CA3, we used *NECTIN3*, *NECAB1*, *AMPH*, and *MPPED1*. For PCL-CA1, we used *FIBCD1*, *MPPED1*, *CLMP*, and *SLC17A7*. For SUB, we used *FN1*, *NTS*, and *SLC17A6*. For SO, we used *PVALB*, *SST*, *GAD1/GAD2*. For SR, we looked for regions between SLM and PCL-CA1/PCL-CA3 with high levels of *GFAP*, but lower levels of *MBP*/*MOBP* compared with SLM and WM. For SL, we looked for regions between SR and PCL-CA3 with low UMIs/detected genes and higher levels of *GFAP* and *MBP*/*MOBP* than adjacent PCL-CA3. For SLM, we noted a strip of darker staining compared with adjacent ML and SR along with expression of *MBP*/*MOBP* and *GFAP*. To identify WM regions, we looked for very high levels of oligodendrocyte markers such as *MBP*, *MOBP*, and *PLP1*. To identify CP, we used *TTR*, *PRLR*, and *MSX1*. CP tissue also had a distinctive sponge-like appearance, making visual identification easy. For thalamus, we used *TCF7L2*. Finally, for CTX, we used *CUX2*, *RORB* and *BCL11B*.

#### Spatial domain annotation

We annotated spatial domains based on anatomical location and expression of canonical marker genes (**Figure 1E**). We found two domains each mapping to CA2-4 (CA2-4.1 and CA2-4.2) and CA1 (CA1.1 and CA1.2). Gene expression profiles for each of these domains were similar (**Extended Data Fig. 8A**), however, the number of nuclei per spot was higher in CA1.1 compared with CA1.2 and CA2-4.1 compared with CA2-4.2 (**Extended Data Fig. 8B**). Therefore, we collapsed CA2-4.1/CA2-4.2 to CA2-4 and CA1.1/CA1.2 to CA1 spatial domains for further analysis. Domains mapping to GCL, Cornu Ammonis pyramidal cell layers (CA2-4.1, CA2-4.2, CA1.1, CA1.2), subiculum (SUB), subiculum and retrohippocampal region (SUB.RHP), retrohippocampal region (RHP), and GABAergic neuron-rich spots (GABA) were grouped as neuron cell body-rich domains at the broad domain level. To assist in annotation of neuropil-rich domains, we referenced the manual annotations (**Extended Data Fig. 8C**). We also examined the expression of genes enriched in the CA1 (MPPED1, FIBCD1), CA3 (TSPAN18, NECTIN3, AMPH), DG (PROX1, SEMA5A), and astrocytic genes since we had a cluster with SGZ-proximal spatial organization (**Extended Data Fig. 8D**). With these aids we annotated domains mapping to dentate gyrus molecular layer (ML), stratum lucidum/stratum radiatum (SL/SR), stratum radiatum/stratum lacunosum-moleculare (SR/SLM), and stratum lacunosum-moleculare/dentate gyrus subgranular zone (SLM/SGZ). Domains mapping to white matter (WM.1, WM.2, and WM.3) were grouped as white matter domains at the broad domain level. Domains mapping to vascular and choroid plexus tissue were grouped as vascular/cerebrospinal fluid (CSF) domains at the broad domain level. The final HPC domain annotations used throughout the manuscript are present in **Extended Data Fig. 9**.

#### Identification of thalamus and amygdala spots

During manual annotation we identified the presence of a small amount of thalamus tissue in one capture area using thalamus-specific marker *TCF7L2* (33,34) (**Extended Data Fig. 10A-C**). In this capture area, thalamus tissue was incorrectly annotated to the SUB and SUB.RHP `PRECAST` domains. The thalamic SUB/SUB.RHP spots were the only spots from SUB or SUB.RHP spatial domains in this capture area. These spots were therefore excluded from differential expression (DE) analyses. In another donor we identified a large, homogenous expanse of tissue that comprised an entire capture area and was annotated to RHP. Based on the anatomy of the region, we believed this region was in fact amygdalar tissue based on the expression of several genes known to be enriched in amygdala: *SLC17A6*, *CDH22* (21), *OPRM1* (36), and *CACNG4* (35) (**Extended Data Fig. 10E-H**). We identified 3 total capture areas from two donors containing tissue from adjacent amygdala based on expression of these genes, all of which were annotated to the RHP spatial domain. These spots were not included in DE analyses.

#### Pseudobulk processing

Following unsupervised clustering, Visium spots were pseudo-bulked by their spatial domain and capture area, as previously described (4). Briefly, we summed the raw gene expression counts for across all spots in a given capture area in a given spatial domain. We performed this aggregation using the `aggregateAcrossCells()` function from `scran` (140). We refer to these aggregated samples as pseudobulk samples. We discarded pseudobulk samples that were composed of less than 50 spots and spots identified as the thalamus or amygdala. We then identified and removed lowly expressed genes using the `filterByExpr()` function from the `edgeR` package (149,150). Counts were normalized by library size, converted to counts per million (CPM), and log2 transformed using the `calcNormFactors()` function from `edgeR`. QC metrics were computed for each pseudobulk sample, again using `addPerCellQC()` from `scuttle` (141). We performed principal component analysis (PCA), retaining the first 100 principal components (PCs) to confirm that pseudobulked samples primarily captured biological variation rather than technical or experimental variables. We used the `getExplanatoryPCs()` function from the `scater` package (141) to compute, for each PC, the percent of variance explained by different metadata variables, including spatial domain, donor, capture area, slide, age, sex, and QC metrics (**Extended Data Fig. 14**, **Extended Data Fig. 15**).

#### Pseudobulk differential expression analysis

We performed differential expression (DE) analysis using pseudobulk log_2_-transformed CPM values to model differences in gene expression across domains or broad domains using `spatialLIBD` functions (138). These functions are wrappers for various functions from the `limma` package (151). We followed a modeling approach similar to Maynard et al. (4). We accounted for age, sex, and slide as covariates. After model matrix construction, we ran `registration_block_cor()` from `spatialLIBD`, blocking by capture area. This function wraps `duplicateCorrelation()` from `limma` to calculate correlation between pseudobulk samples within each block. This approach accounted for capture area-specific variation. For the enrichment model, we used `registration_stats_enrichment()` from `spatialLIBD`, wrapping the same `limma` functions to test for gene expression differences between each domain and all other domains. For each gene and domain, this model computed fold change (FC), Student’s *t*-test statistics, and two-tailed *p-*values. *P*-values were corrected for the false discovery rate (FDR) (152). Genes were considered significant if log_2_FC > 1 or log_2_FC < -1 and FDR < 0.01 (**Extended Data Fig. 16**, **Extended Data Fig. 17**, **Supplementary Table 4**, **Supplementary Table 5**).

### 4.5 snRNA-seq data generation

#### snRNA-seq data collection and sequencing

Using previously mentioned 100μm cryosections collected from each donor, we conducted single-nucleus RNA-sequencing (snRNA-seq) using 10x Genomics Chromium Single Cell Gene Expression V3 technology. Cryosections for each donor were first pooled with chilled Nuclei EZ Lysis Buffer (MilliporeSigma #NUC101) into a glass dounce. Sections were homogenized using 10-15 strokes with both pestles (loose and tight-fit). Homogenates were filtered through 70 μm mesh strainers and before centrifugation at 500 x g for 5 minutes at 4°C using a benchtop centrifuge. Nuclei were resuspended in fresh EZ lysis buffer, centrifuged again, and equilibrated in wash/resuspension buffer (1x PBS, 1% BSA, 0.2U/μL RNase Inhibitor). Nuclei were washed in wash/resuspension buffer and centrifuged 3 times. We labeled nuclei with Alexa Fluor 488-conjugated anti-NeuN (MilliporeSigma cat. #MAB377X), diluted 1:1000 in nuclei stain buffer (1x PBS, 3% BSA, 0.2U/μL RNase Inhibitor) by incubating at 4°C with continuous rotation. Following NeuN labeling, nuclei were washed once in stain buffer, centrifuged, and resuspended in wash/resuspension buffer. Nuclei were labeled with propidium iodide (PI) at 1:500 in wash/resuspension buffer before being filtered through a 35um cell strainer.

We then performed fluorescent activated nuclear sorting (FANS) for using a Bio-Rad S3e Cell Sorter. Gating criteria were selected for whole, singlet nuclei (by forward/side scatter), G0/G1 nuclei (by PI fluorescence), and neuronal nuclei (by Alexa Fluor 488 fluorescence). Nuclei from each donor were split into two equal samples. The first sample was sorted based on PI+ fluorescence, thereby including both neuronal and non-neuronal nuclei. The second sample was sorted based on both PI+ and NeuN+ fluorescence to facilitate enrichment of neurons.

This approach initially gave *n*=20 for snRNA-seq (1 PI+ and 1 PI+NeuN+ sample for all 10 donors) with a total of 18,000 sorted nuclei per donor (9000 per Chromium sample). Samples were collected over multiple rounds, each containing 1-3 donors for 2-6 samples per round. We had poor nuclei yields from one sequencing round, comprising both PI+ and PI+NeuN+ samples from 3 donors. We collected additional PI+ and NeuN+ samples from these 3 donors and performed nuclei sorting steps again, this time with better final nuclei yields. Therefore, we had a final *n*=26 for snRNA-seq (1 P1+ and 1 PI+NeuN+ sample for 7 donors, 2 PI+ and 2 PI+NeuN+ samples for 3 donors).

All samples were sorted into reverse transcription reagents from the 10x Genomics Single Cell 3′ Reagents kit (without enzyme). Enzyme and water were added to bring the reaction to full volume. cDNA synthesis and subsequent sequencing library generation was performed according to the manufacturer’s instructions for the Chromium Next GEM Single Cell 3’ v3.1 (dual-index) kit (CG000315, revision E, 10x Genomics). Samples were sequenced on a Nova-seq (Illumina) at the Johns Hopkins University Single Cell and Transcriptomics Sequencing Core at a minimum read depth of 50000 reads per nucleus.

### 4.6 snRNA-seq data analysis

#### snRNA-seq data processing and quality control

Following sequencing, we mapped reads to Genome Reference Consortium Human Build 38 (GRCh38 2020-A) using `cellranger count` (version 7.0.0). The raw feature matrices were used to generate a `SingleCellExperiment` (136) object that was then filtered with the `emptyDrops()` function from the `dropletUtils` (153) Bioconductor package. We computed quality control metrics using the `addPerCellQC()` function from the `scuttle` Bioconductor package (141).

To identify poor-quality nuclei, we applied a 3x median absolute deviation (MAD) threshold for key QC metrics (percentage of UMIs mapping to mitochondrial genes, library size, and number of detected genes) (141). Because neurons have larger library sizes and numbers of expressed genes compared with non-neuronal cells in human brain tissue (154), and because our PI+NeuN+ samples are enriched for neurons, these initial thresholds were computed on a per-sample basis (**Extended Data Fig. 4A-C**). We recognized that this approach resulted in an inconsistent threshold for the rate of mitochondrial expression that resulted in several samples retaining nuclei with >10% of reads originating from the mitochondrial genome (**Extended Data Fig. 4A**) and that these samples with the highest proportion of mitochondrial reads had the lowest number of nuclei removed (**Extended Data Fig. 4D**). Instead of pooling nuclei for MAD calculation on a per-sample basis, we calculated a new 3MAD threshold by pooling the higher quality samples with original thresholds of <5% expression from the mitochondrial genome (**Extended Data Fig. 4G**). This increased the number of nuclei excluded based on mitochondrial fraction, particularly in samples from sequencing round 3 that exhibited low cDNA abundance and were re-run (**Extended Data Fig. 4J**). The initial 3MAD thresholds for the other two QC metrics resulted in no nuclei being excluded from almost all PI+ sorted samples and many PI+NeuN+ sorted samples (**Extended Data Fig. 4E-F**). For these samples, we manually set the library size and number of genes detected threshold to 1,000 counts and 1,000 detected genes (**Extended Data Fig. 4H-I**), increasing the number of nuclei excluded (**Extended Data Fig. 4K-L**). Following nuclei quality control, the dimensions of our dataset were 36,601 genes x 86,905 nuclei.

We observed that one sample of PI+NeuN+ nuclei remaining after QC filters possessed noticeably lower library sizes and fewer detected genes than all other PI+NeuN+ samples (**Extended Data Fig. 5B-C**). To see if this discrepancy was likely the result of a neuronal population unique to that sample (17c-scp), we performed rudimentary clustering with the `quickCluster()` function from `scran` and estimated the number of neurons present in each cluster based on raw expression of SYT1 >1 count (**Extended Data Fig. 5D**). We found that across neuronal clusters, sample 17c-scp displayed an increased number of nuclei with low library size and few detected genes, but that there remained 17c-scp nuclei that matched the distribution of these QC metrics present in the other PI+NeuN+ samples (**Extended Data Fig. 5E**). We reasoned that many of the nuclei in sample 17c-scp were low-quality neurons. To prevent detrimental impact on downstream analyses, we took a conservative approach and removed all nuclei with fewer than 5000 detected genes in this sample only. Following this removal step, the dimensions of our dataset were 36,601 genes x 80,594 nuclei (**Extended Data Fig. 5F**).

#### snRNA-seq feature selection, dimensionality reduction, and clustering

For feature selection and dimension reduction, we used the methods developed in `scry` (155) which fit a Poisson model to the raw counts of each gene and then rank genes by deviance from the null hypothesis that the modeled expression is equal across all cells. The ranks were used to select highly deviant genes (HDGs) via `devianceFeatureSelection()`, and the Pearson residuals from the Poisson model for these 2000 HDGs (obtained with `nullResiduals()`) were used as input for principal component analysis (PCA) that was implemented with `scater` (141). Utilizing the Pearson residuals rather than log-normalized count data as input for PCA avoids biases introduced by log-normalization of UMI count data (155). Mutual nearest neighbors (MNN) correction was implemented with `batchelor` (156) and was applied to the top 50 PCs to correct for batch effects introduced by donor and sequencing rounds. Visualizing these results with UMAP embedding shows this approach was highly effective (157) (**Extended Data Fig. 18**). Gene expression count data was then normalized using `scran` (140). We computed cluster-specific size factors by generating rough cluster assignments using the `quickCluster()` function followed by the `calcSumFactors()` function from `scran` (158). These size factors were used to normalize the count matrix before implementing log transformation.

We implemented graph-based clustering to identify cell types present in the snRNA-seq dataset. A nearest neighbor graph was generated with `scran` (140) based on the 50 MNN-corrected PCs using *k*=5 nearest neighbors and Jaccard weights followed by `igraph` (159) implementation of Louvain clustering (160). This resulted in 59 clusters. We annotated these clusters as neuronal or non-neuronal based on expression of neuron-specific genes, such as *SYT1* (**Extended Data Fig. 19A**). We identified three low-quality neuron clusters based on fewer detected genes (**Extended Data Fig. 19B**). We also identified one cluster of likely doublets, which were marked by coexpression of oligodendrocyte, astrocyte, OPC, and microglia markers in the same nuclei (**Extended Data Fig. 19C**). These 4 clusters were removed, together accounting for 5% of all nuclei.

After removing low quality neurons and likely doublets, we repeated feature selection, dimensionality reduction, batch correction, and graph-based clustering using the same parameters as before. This resulted in 62 clusters which we classified as neuronal and non-neuronal based on *SYT1* expression (**Extended Data Fig. 19D**). Using the number of detected genes and co-expression of distinct glial markers, we identified one low-quality neuron cluster (**Extended Data Fig. 19E**) and one low-quality non-neuronal cluster, respectively (**Extended Data Fig. 19F**). These two clusters contained 1.6% of all nuclei, and so we did not reprocess the dataset after removal. Our final dataset contained 75,411 nuclei classified into 60 clusters.

#### Preliminary snRNA-seq cluster annotation

We utilized markers from published snRNA-seq data in humans (18) to label broad cell types including glia cell types (**Extended Data Fig. 19F**), inhibitory/GABAergic neurons, and excitatory/glutamatergic neurons (**Extended Data Fig. 20A**). Within excitatory neurons, we were able to identify distinct clusters of *cornu ammonis* pyramidal cells, dentate gyrus granule cells, and subiculum pyramidal cells based on marker genes like *PROX1*, *CALB1*, *FNDC1*, *TSPAN18*, *CARTPT*, and *FN1*. We knew that we should have a small thalamus population and identified this population by looking for a cluster with elevated *TCF7L2* expression based on our SRT data (**Extended Data Fig. 10A-C**, **Extended Data Fig. 20B**). Most of the remaining excitatory cell clusters likely comprise the retrohippocampal regions which exhibit laminar organization akin to cortical layers and are often annotated based on the expression of *CUX2* (superficial marker), *SATB2* (deep callosal projection marker), *TLE4* (deep subcortical projection marker) (**Extended Data Fig. 20A**). We also knew that there should be some amygdalar neurons present, but were unable to identify these clusters based on the expression of genes used to classify amygdalar domains in the SRT dataset (*SLC17A6*, *CDH22*, *OPRM1*, and *CACNG4*) (**Extended Data Fig. 10D-H**). To identify amygdala neuron clusters in the snRNA-seq data we isolated the RHP spots from SRT data and used `scran::findMarkers()` to implement a binomial test for gene markers that distinguished the spots annotated to the amygdala (*n*= 4519 spots) from the true RHP domain (*n*= 3940 spots). Evaluating markers increased in either population at an FDR<.05 and log_2_FC>.3, we identified 73 genes increased in amygdala spots and 86 genes increased in RHP spots. We examined the expression of these genes in all excitatory cells annotated to RHP layers and identified 5 clusters that we relabeled as amygdalar neurons (*n*= 5524 nuclei) (**Extended Data Fig. 20C**). Importantly, these nuclei overwhelmingly came from the donors with amygdala domains identified in SRT (Br6432 *n*= 3380 nuclei, Br6423 *n*= 1338 nuclei), although we did annotate 721 nuclei from Br8667 to these amygdala clusters.

#### Pseudobulk processing

Following unsupervised clustering, nuclei were pseudo-bulked by cluster (*n*=60) and sample. We dropped all genes with zero counts across all nuclei and summed the raw gene expression counts for across all nuclei in a given sample in a given cluster using the `aggregateAcrossCells()` function from `scran` (140). We discarded pseudobulk samples that were composed of less than 10 nuclei (864 remaining pseudobulk samples). This resulted in the complete removal of one cluster (cluster 58) that had already been identified as Cajal Retzius cells by the expression of *RELN*. We then identified and removed lowly expressed genes using the `filterByExpr()` function from the `edgeR` package (149,150) (21,104 genes remaining). Counts were normalized by library size, converted to counts per million (CPM), and log_2_ transformed using the `calcNormFactors()` function from `edgeR`. QC metrics were computed for each pseudobulk sample, again using `addPerCellQC()` from `scuttle` (141).

#### Pseudobulk differential expression analysis

We performed differential expression (DE) analysis on the *n*=60 clusters using pseudobulk log_2_-transformed CPM values. As with SRT DE analysis, we used `spatialLIBD` functions (138). We accounted for age, sex, and sequencing round as covariates. After model matrix construction, we ran `registration_block_cor()` blocking by sample to account for sample-specific variation. We then fit an enrichment model as described in Maynard et al. (4), taking the block correlation into account. For each gene and cluster, this model computed fold change (FC), Student’s *t*-test statistics, and two-tailed *p*-values. *P*-values were corrected for the false discovery rate (FDR) (152) (**Supplementary Table 6).**

#### Detailed snRNA-seq cluster annotation

We used the significant markers (log_2_FC > 2 and FDR < 0.0001) from pseudobulk DE analysis to provide greater detail to the cluster labels and identified differentially expressed genes (DEGs) that distinguished nearly all 60 clusters as unique in some way (**Extended Data Fig. 22**, **Figure 3D**). With this approach we were able to annotate small populations of distinct non-neuronal cell types through the cluster-specific expression of *LAMA2* (fibroblast population of vascular lepto-meningeal cells (VLMCs)) (161), *DLC1* (pericytes/ smooth muscle cells (PC/SMC)) (162), *MECOM* (endothelial cells) (163), and *GRP17* (committed oligodendrocyte precursors (COPs)) (37) (**Extended Data Fig. 22A**). DEGs across the six superficial retrohippocampal (RHP) pyramidal neuron clusters indicated the grouping of two clusters (L2/3.1 and L2/3.5) based on the expression of *TESPA1* and *TSHZ2* (**Extended Data Fig. 22B**, **Figure 3D**). Even across the amygdala neurons we identified genes distinguishing the clusters from one another, even though the number of donors contributing to this population was small as the amygdala was not systematically targeted for dissection (**Extended Data Fig. 22C**). We were able to annotate two subiculum clusters based on *CNTN6* expression (13) and all HPC pyramidal neurons exhibited a gradient of *NUP93* expression that may serve to drive HPC identity (164) (**Extended Data Fig. 22D**).

GABAergic neurons aggregated into cell types based on the expression of cell type-specific gene expression (**Extended Data Fig. 22E**) and gene expression that indicated broader GABAergic families (**Extended Data Fig. 22F**). Using these DEGs, we annotated GABAergic families corresponding to central ganglionic eminence (CGE) origin (consisting of VIP and HTR3A cell types) based on *ADARB2* expression (154) and lack of *NXPH1*, a *LAMP5*+ family of *ADARB2* expressing interneurons (165) (consisting of CXCL14, LAMP5.CGE and LAMP5.MGE cell types), a group of GABAergic neurons from the medial ganglionic eminence (MGE) (consisting of CRABP1, C1QL1, PV.FS, CORT, and SST cell types) based on *LHX6* expression (154,165), and a population of *PENK*+ interneurons that originates prior to the major GABAergic/ glutamatergic split (165). We noted that the CXCL14 cluster exhibited high *RELN* expression but confirmed these were not mis-labeled Cajal Retzius neurons based on the lack of *TP73* expression (**Extended Data Fig. 22F**) and likely represent a population of *LAMP5*+ interneurons that have been shown to continue to express *RELN* into adulthood (166).

Even the smallest clusters (like GC.5 *n*=96 nuclei, thalamus *n*=35 nuclei, COP *n*=57 nuclei, AHi.4 *n*=73 nuclei) exhibited clear expression of marker genes unique from similar clusters. Some clusters were difficult to distinguish even with DEG expression (e.g., Micro.1 vs Micro.2, CP.1 vs CP.2 vs CP.3, **Extended Data Fig. 22A**). However, our ability to discern gene markers unique to each of the 60 clusters suggested that the high resolution with which we determined our cell clusters did not result in over-clustering.

It is often useful to classify nuclei at a broader resolution, so we generated several different classification levels based on the *n*=60 superfine cluster annotations (**Extended Data Fig. 28**). The “fine” classification level is used for visualization throughout the manuscript, and the “broad”, “mid”, and “fine” classifications were used in cell type deconvolution. We verified that across these resolutions we did not see donor-specific enrichment that was unexpected (e.g., amygdala enrichment expected) (**Extended Data Fig. 21**).

#### Validation of snRNA-seq analysis approach

Given the importance of cluster identity for downstream analysis, we performed two additional checks to increase confidence in the cell types identified in the snRNA-seq results. First, we correlated the gene-level enrichment model *t*-statistics for each of the 60 snRNA-seq clusters with the *t*-statistics from the enrichment model of the SRT domains (**Extended Data Fig. 23**). Second, we performed a re-analysis of the snRNA-seq data from the initial QC through to clustering. Our rationale with this re-analysis is that, if we identify similar cell clusters using an orthogonal framework with different computational tools, then the conclusions generated from our *n*=60 clusters are generalizable. Re-analysis was performed while blinded to whether individual nuclei were included or excluded in the original analysis.

In our re-analysis, instead of identifying empty droplets with Bioconductor tools, we started with the filtered feature matrix from which empty droplets were already removed during `cellranger` processing. We removed the samples from sequencing round 3 that had increased debris during sorting and exhibited high mitochondrial fraction (starting nuclei = 118,415) (**Extended Data Fig. 24A**). Because the samples from sequencing round 3 had been re-collected, there were still N=10 donors represented. After evaluating a maximum initial mitochondrial fraction of 20%, we further filtered out all nuclei with >10% reads from the mitochondrial genome (remaining nuclei = 116,494) (**Extended Data Fig. 24B-C**). Minimum thresholds for library size and the number of detected genes were evaluated separately for PI+ and PI+NeuN+ samples due enrichment of neurons in the PI+/NeuN+ population, the increased number of reads in the larger neuronal nuclei, and the distinct distribution of these metrics across the two sorting methods (**Extended Data Fig. 24D,H**). For the PI+NeuN+ data, one sample in sequencing round 2 (sample 17c-scp) exhibited a substantially shifted distribution and was therefore excluded when determining QC thresholds (**Extended Data Fig. 24F,J**). Our final thresholds for re-analysis were a maximum of 10% mitochondrial fraction for any sample, a minimum of 750 detected genes for PI samples (**Extended Data Fig. 24E**), a minimum of 1000 detected genes for PI+NeuN+ samples (**Extended Data Fig. 24G**), a minimum of 2000 reads for PI samples (**Extended Data Fig. 24I**), and a minimum of 3500 reads for PI+NeuN+ samples (**Extended Data Fig. 24K**). The total number of nuclei after QC filters was 81,838, with 22.5% of the 36,577 discarded nuclei originating from sample 17c-scp with reduced quality round 2 (all remaining samples constituting 2.85% to 6.80% of excluded nuclei, consistent with **Extended Data Fig. 5F**).

Feature selection for dimensionality reduction for re-analysis was conducted using the same binomial deviance approach as before, given the ability of this method to handle raw count data and account for batch effects. For dimensionality reduction we performed `scVI` (38) analysis in python to generate latent representations that replaced PCA followed by MNN batch correction. Our `scVI` model included the covariates of sequencing round, donor ID, mitochondrial fraction, age, and postmortem interval. `Scanpy` (167) was then used to construct a nearest neighbors map of *k*=15 neighbors based on the scVI latent representations, and then to generate cell clusters utilizing the Leiden algorithm at a resolution of 0.5 (**Extended Data Fig. 25A**). `scVI` latent representations were also used to calculate UMAP embeddings for visualization. Clusters were then refined based on the expression of *MALAT1*, which has been shown to indicate the quality of nuclei in snRNA-seq data, and based on mitochondrial fraction (168) (**Extended Data Fig. 25B-F**). After filtering out low quality clusters, new neighbors (*k*=10) and clusters were found, and new UMAP embeddings computed (**Extended Data Fig. 25G-H**). Two small clusters (less than 100 nuclei) were removed due to size, and one additional small cluster was removed because it comprised nuclei from mostly one sample (110 of 138 nuclei from one sample).

Despite the very different approach to QC, re-analysis within a python framework produced a highly similar set of nuclei: 64,251 were nuclei present in both the new set (74,216 total) and our original set (75,411 total) (**Extended Data Fig. 25I**). To examine the similarity of the cell clusters generated, we performed the pseudobulk enrichment model on the re-analysis clusters. Specifically, we found that the clusters identified with `scVI` re-analysis correlated strongly with our original *n*=60 clusters when comparing the gene expression patterns captured by the nuclei groupings (**Extended Data Fig. 25J**). This shows that the same cell types were found in the re-analysis and that the nuclei that were present in only one of the two datasets were spread across all clusters, suggesting we were not “missing” any cell types in either analysis.

### 4.7 Stratified Linkage disequilibrium score regression (S-LDSC)

Prior to S-LDSC, we defined gene sets for each set of labels. For snRNA-seq superfine cell types and SRT spatial domains, we used *t*-statistics from our pseudobulk DE analysis enrichment model (each domain/superfine cell type vs all other domains/cell types). For each unique domain or superfine cell type, genes were ranked by enrichment model *t*-statistics. The top 10% of genes by t-statistic were selected as the gene set for each unique domain or superfine cell type. For NMF patterns, we first calculated *z*-scores for each gene across all patterns not mapping to batch effects or technical variables. We then ranked genes by *z*-score for each pattern. The top 10% of genes by *z*-score were selected as the gene set for each unique NMF pattern. To construct a genome annotation for each unique label (spatial domain, snRNA-seq superfine cell type, or NMF pattern), we added a 100Kb window upstream and downstream of the transcribed region of each gene in that label’s corresponding gene set.

We performed stratified LD score regression (`S-LDSC`) to evaluate the enrichment of heritability of brain-related traits for gene sets defined for different domains. We also included two non-brain traits, human height and type 2 diabetes, as negative controls to examine whether our findings are specific to brain-related traits. We downloaded GWAS summary statistics of each trait (46–61). Following recommendations from the LDSC resource website (https://alkesgroup.broadinstitute.org/LDSCORE), `S-LDSC` was run for each gene set with the baseline LD model v2.2 that included 97 annotations to control for the LD between variants with other functional annotations in the genome. We used HapMap Project Phase 3 SNPs as regression SNPs, and 1000 Genomes SNPs of European ancestry samples as reference SNPs, which were all downloaded from the LDSC resource website. To evaluate the unique contribution of gene sets to trait heritability, we utilized the metric from `S-LDSC`: the *z*-score of per-SNP heritability. This metric allows us to discern the unique contributions of candidate annotations while accounting for contributions from other functional annotations in the baseline model. The *p*-values are derived from the *z*-score assuming a normal distribution and FDR was computed from the *p*-values based on Benjamini & Hochberg procedure.

### 4.8 Spot-level deconvolution algorithm benchmarking

### Spot deconvolution benchmarking

Given that individual Visium spots often contain multiple cell types (**Extended Data Fig. 29**), we performed spot deconvolution (**Extended Data Fig. 29**) to better understand the cellular composition of spots mapping to unsupervised spatial domains (**Extended Data Fig. 29**). While several algorithms have been developed to predict cell type proportions within individual Visium spots using single cell reference data, they have not yet been comprehensively benchmarked across various brain regions due to limited availability of complete reference datasets which contain 1) paired SRT and snRNA-seq data from the same donors, 2) known cell type abundances in each gene expression spot, 3) anatomical regions enriched for specific cell types. Given that the morphological organization of the hippocampus presents a unique computational challenge for these algorithms, we present the first gold standard spot deconvolution dataset for 4 broad cell types in postmortem human anterior HPC using the Visium Spatial Proteogenomics assay (Visium-SPG), which replaces H&E histology with immunofluorescence staining to label proteins of interest with fluorescent dyes (**Extended Data Fig. 27**) similar to previous work in the dlPFC (134). We selected 2 out of the 10 brain samples, one male and one female, based on exceptional morphology and inclusion of all spatial domains. We performed immunofluorescent staining for established cell type-specific proteins, including NEUN (marking neurons), OLIG2 (marking oligodendrocytes), GFAP (marking astrocytes), and TMEM119 (marking microglia) (**Extended Data Fig. 27**). Following multispectral fluorescence imaging, we proceeded with the standard Visium protocol to generate corresponding gene expression libraries from the same tissue section. This provided an orthogonal measure of broad cellular identity for each spot.

#### Visium-SPG data generation

To enable labeling with four cell-type marker proteins (NeuN for neurons, TMEM119 for microglia, GFAP for astrocytes, and OLIG2 for oligodendrocytes) (**Extended Data Fig. 27A**), we performed combinatorial immunofluorescence staining paired with spatial gene expression using the Visium-SPG protocol (protocol CG000312, revision D, 10x Genomics) with some modifications. Briefly, gene expression slides were fixed in methanol, incubated in blocking buffer (3X SSC supplemented with 2% BSA, 0.1% Triton X100 and 2U/ul RNAse inhibitor (RiboLock, Thermofisher, Cat# EO0384)), then incubated with primary antibodies for 30 minutes at room temperature: mouse anti-NeuN antibody conjugated to Alexa 488 (Sigma Aldrich, Cat# MAB377X, 1:100), rabbit anti-TMEM119 antibody (Sigma Aldrich, Cat# HPA051870, 1:20), rat anti-GFAP antibody (Thermofisher, Cat# 13-0300, 1:100), and goat anti-OLIG2 antibody (R&D systems, Cat# AF2418, 1:20). Slides were washed with wash buffer (3X SSC with 2% BSA, 0.1% Triton X100, 2U/ul RNAse inhibitor and supplemented with ribonucleoside vanadyl complex (Concentration, NEB S1402S), then incubated for 30 minutes with secondary antibodies at RT: donkey anti-rabbit IgG conjugated to Alexa 555 (Thermofisher, Cat# A-31572, 1:300), donkey anti-rat IgG conjugated to Alexa 594 (Thermofisher, Cat# A-21209, 1:600), and donkey anti-goat IgG conjugated to Alexa 647 (Thermofisher, Cat# A-21447, 1:400). DAPI (Thermofisher, Cat# D1306, 1:3000, Final 1.67 μg/ml) was included as a nuclear counterstain. The slide was coverslipped with glycerol containing RNase Inhibitor (2 U/μl) and scanned on a Vectra Polaris slide scanner (Akoya Biosciences) at 20x magnification with the following exposure time per given channel: 2.70 msec for DAPI; 185 msec for Opal 520; 900 msec for Opal 570; 160 msec for Opal 620; 1300 msec for Opal 690; 100 msec for autofluorescence. Following imaging, samples were processed as above for library generation and sequencing (**Methods 4.3**).

#### Visium-SPG quality control and count normalization

FASTQ and image data were pre-processed with the 10x `SpaceRanger` pipeline and aligned to reference genome GRCh38 2020-A (version 2.0.0) (135). Visium-SPG spot-level data was subject to the same quality control metrics and count normalization steps performed above (**Methods 4.4**, **Extended Data Fig. 27B-D**). Following QC, we retained 37,845 spots in *n*=8 capture areas for the Visium-SPG data (**Extended Data Fig. 27E**).

#### Spatial domain projection into Visium-SPG samples

We used the `RcppML` (67) R package to approximate spatial domain membership for Visium-SPG samples, which were not included in the `PRECAST` clustering analysis. We used the `aggregateAcrossCells()` function from `scran` (140) to pseudobulk the logcounts matrix from Visium-H&E samples only, summing all logcounts for all genes across all spots in each domain. We pseudobulked the logcounts by domain such that our pseudobulked SPE object had 16 columns, one for each domain. We filtered lowly expressed genes using the `filterByExpr()` function from the `edgeR` package (149,150). We rescaled columns of the pseudobulked matrix such that each column summed to 1, akin to the scaling diagonal used in NMF calculations in `RcppML` (67). We then used the `project()` function from `RcppML` to estimate `PRECAST` spatial domain membership of each spot in the Visium-SPG samples. This function projects a linear factor model by solving this equation for *H* when given *A* and *W*:

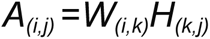

where *A* is a matrix with dimensions *i x j* and *W* and *H* are orthogonal, lower-rank matrices with dimensions *i x k* and *k x j*, respectively. In the context of `RcppML`, this function is generally used to project an NMF model of rank *k* trained in one dataset into a new dataset. We detail this process further in **Methods 4.8**. Here, we instead used this function to predict `PRECAST` domain membership of each spot in the Visium-SPG samples (*H* matrix), given the Visium-SPG logcounts matrix (*A* matrix) and the pseudobulked domain-level matrix described above (*W* matrix). In this context, the *A* matrix was the Visium-SPG logcounts matrix, the *W* matrix was the pseudobulked logcounts matrix described above, and the *H* matrix was a matrix of spatial domain membership predictions for Visium-SPG spots. These predicted domain membership values were continuous. Therefore, for each spot, we assigned domain membership based on the highest predicted domain value (**Extended Data Fig. 27F**).

#### Image-based cell type classification

We leveraged the immunofluorescence (IF) images in the Visium-SPG assay to directly quantify cell type abundance and provide an imaging-based calculation of cell counts for comparison with outputs from spot deconvolution tools. Starting from the `cyto` model provided by *Cellpose* 2.0 (169), nuclear segmentations of Br3942 and Br8325 were iteratively improved in the *Cellpose* GUI, and a new model was trained until DAPI segmentations visually satisfied our accuracy standards (**Extended Data Fig. 30A**). The final refined model was used to segment all 8 capture areas to produce masks. Segmentation masks were read into python 3.8.12, and mean fluorescence intensity for the NeuN, OLIG2, GFAP and TMEM119-marked channels was quantified using `skimage.measure.regionprops_table()` from the `scikit-image` (170) v0.19.2 library. For 2 capture areas (V12D07-332_D1 and V12D07-335_D1) which represented all HPC regions, an expert experimenter (co-author SCP) used `Samui Browser` (112) to randomly select and annotate a spatially diverse set of 200 cells of each immunolabeled cell type – e.g. neuron, oligodendrocyte, astrocyte, microglia, or other (i.e. presence of DAPI, but absence of other immunofluorescent signals). The resulting dataset included ∼1000 cells with a cell type label, and the corresponding mean fluorescence intensities from the respective channels (**Extended Data Fig. 30A**). We then calculated the actual number of neurons, astrocytes, microglia, and oligodendrocytes per spot by segmenting individual nuclei and implementing a classification and regression tree (CART) approach in `scikit-learn` (171) to categorize all nuclei into either one of the 4 immunolabeled cell types or “other” (**Extended Data Fig. 29**, **Extended Data Fig. 30B**).

#### Input marker gene detection

We considered both the “fine” and “mid” cell type classifications from the snRNA-seq data (**Figure 3B-C**, **Extended Data Fig. 28**). The `get_mean_ratio2()` function from `DeconvoBuddies` (version 0.99.0) (172) was applied to rank each gene in the snRNA-seq data in terms of its suitability as a marker for each possible target cell type at both resolutions (**Extended Data Fig. 29**). This method, termed “mean ratio”, determines the ratio of expression between a target cell type and the next-highest-expressing cell type (**Extended Data Fig. 31**) (172). Mitochondrial genes and genes not present in the spatial data were excluded from the ranking process. Three snRNA-seq clusters from the fine grouping (HATA, Amy, and GABA.PENK) were also excluded as these were not represented in any Visium-SPG samples and could occlude marker gene identification for other clusters. We retained the thalamus cluster because one of the Visium-SPG samples included a very small amount of thalamus (**Extended Data Fig. 1B, Extended Data Fig. 10**) which was transcriptionally distinguishable from hippocampal cells. The top-25-ranked markers by mean ratio were taken for each cell type, and, to ensure greater expression in the target cell type than all others, we verified that the mean ratio exceeded 1 for each gene (**Extended Data Fig. 31**, **Extended Data Fig. 32**, **Extended Data Fig. 33**). This resulted in a total of 200 mid-resolution markers and 525 fine-resolution markers used downstream for spot deconvolution (**Supplementary Table 7**).

To confirm the utility of the snRNA-seq-derived marker genes for spot deconvolution, we evaluated their proportion of nonzero expression in Visium-SPG data (**Extended Data Fig. 32**) and confirmed that 25 genes per cell type were sufficient to identify the expected cell type distribution patterns (**Extended Data Fig. 33**). We confirmed that detected marker genes were specific to each cell type by comparing expression of the top genes for each cell type to canonical gene markers in the expected spatial domains (**Extended Data Fig. 32**, **Extended Data Fig. 33**, **Supplementary Table 7**). `RCTD` (64) dropped a high number of spots using the Mean Ratio marker genes compared to `cell2location` (63) and `Tangram` (65) , thus, the default `RCTD` marker gene detection strategy was employed to run RCTD (**Extended Data Fig. 34**).

#### Applying spot deconvolution softwares to Visium-SPG data

Having verified the selection of robust marker genes for each cell type at both fine and mid resolutions, we ran `cell2location` (63), `RCTD` (64) and `Tangram` (65) and calculated the predicted cell type counts and proportions per spot (**Extended Data Fig. 29**). `Cell2location` v0.8a0, `RCTD` v2.2.0 and `Tangram` v1.0.2 were applied with default parameters following the respective tutorials provided by their authors, with a few exceptions. The marker genes derived from the `Deconvobuddies` package were used as training genes. This strategy was effective for `cell2location` and `Tangram`. However, in `RCTD` it failed to predict cell types in spots with fewer UMIs (∼<100) after filtering the genes to include only the user derived marker genes (**Extended Data Fig. 34**). We observed that `RCTD` dropped a high number of spots when user-defined marker genes were used (**Extended Data Fig. 34A**), but performance improved when `RCTD`’s default marker gene detection strategy was employed (**Extended Data Fig. 34B**). `Tangram` requires an accurate cell count per spot, and thus was provided total cell counts per spot derived through nuclei segmentation with `Cellpose` v2.0 (Visium-SPG images) and through `VistoSeg` (Visium-H&E images). `Cell2location` and `RCTD` require an average count of nuclei per spot, which was calculated as 5 across all 8 Visium-SPG capture areas. While `Tangram` and `Cell2location` predict cell type abundances in each spot, `RCTD` predicts the cell type proportion per spot. Therefore, to more directly compare across algorithms, we multiplied the cell type proportions obtained from `RCTD` by the nuclei counts to obtain abundances for each spot.

#### Evaluating performance of spot deconvolution methods

The immunofluorescence values provided an transcriptomic-independent metric to classify cell type identity within each spot (**Extended Data Fig. 29**). The cell type counts estimated by each algorithm in the Visium-SPG data at fine and mid resolutions were “collapsed” down to the broad categories (neuron, astrocytes, microglia, oligodendrocytes, and other (comprised of OPCs, vasculature, and CSF-related cells)) to enable direct comparison with the CART-calculated counts (**Extended Data Fig. 29**). We then visually compared the predicted cell type (of mid cell types collapsed down to broad cell types) proportion per spot from the 3 softwares to the CART-estimated proportions of immunolabeled cells (**Extended Data Fig. 35**). The collapsed counts from mid and fine resolutions were compared against each other which should theoretically match (**Extended Data Fig. 358A**). To quantify the comparisons, the “other” cell type was excluded from the benchmarking analyses and counts for each software tool and CART-quantified cell type were summed across each Visium-SPG section. Totals for each software method (gene expression) were compared against the CART-predicted (immunofluorescence) cell count totals using Pearson correlation and root mean squared error (RMSE) at mid (**Extended Data Fig. 36**) and fine (**Extended Data Fig. 36**) levels (**Supplementary Table 8**). Counts for each software tool and `CART` predictions were normalized to add to 1 across the four cell types which allowed the calculation of the Kulback-Lieber divergence (KL divergence) from each software tool’s predictions to the `CART` predictions. This treats the CART-predicted cell type composition as a ground-truth probability distribution that each software tool is attempting to estimate (**Extended Data Fig. 36**).

Given the challenge of deconvolving transcriptionally fine cell types, our Visium-SPG benchmark analyses confirmed `RCTD` showed the most consistent performance at both mid and fine resolution across all cell types and samples (**Extended Data Fig. 36**) closely aligning with biological expectations and revealing spatial variations in cell types (**Extended Data Fig. 37**). `Tangram` predicted a similar cellular composition in each domain and predicted the presence of all cell types in all spots. `cell2location` showed a similar pattern but with less pronounced consistency (**Extended Data Fig. 29**). In summary, these analyses confirmed `RCTD` is most suitable for spot deconvolution of Visium HPC data because it accounts for the cellular heterogeneity seen across the HPC.

#### Applying spot deconvolution software to Visium-H&E data

Having evaluated the performance on the Visium-SPG data, we ran `RCTD`, the best performing method, on the whole SRT data for which orthogonal measures of spot-level cellular composition were not available (**Extended Data Fig. 29**, **Figure 4**, **Extended Data Fig. 37**). The `RCTD` deconvolutions results for Visium-SPG and the SRT data are available on the Samui web application for visualization (see **Results 2.7**).

### 4.9 Non-negative matrix factorization (NMF)

#### NMF factorization

We used non-negative matrix factorization (NMF) to identify continuous patterns of gene expression within our snRNA-seq data (**Extended Data Fig. 38**). We refer to the snRNA-seq data as our “source” data. NMF is a dimensionality reduction technique for pattern recognition (173) used across various disciplines, including transcriptomics (66,69). NMF decomposes a matrix *A* into two lower-rank matrices *W* and *H* corresponding to gene-level and nucleus-level weights matrices, respectively:

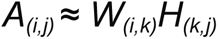

where *k* is the rank of the factorization and both *w* and *h* are constrained to have only non-zero elements. The decomposition is performed iteratively until the difference between *A* and the product of *w* and *h.* Iterations continue until the change in mean squared error (MSE) between successive iterations falls below a specified level (tolerance of fit). We used the `RcppML` package for NMF (67), which also features a scaling diagonal *D* such that:

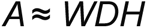

where *D* diagonalizes columns in *W* and rows in *H* to sum to 1. In our case, *k*=100 patterns and *A* was the normalized snRNA-seq counts matrix with dimensions 36,601 genes x 75,411 nuclei. Therefore, *W* had dimensions 36,601 genes x 100 patterns and *H* had dimensions 100 patterns x 75,411 nuclei.

We used the `singlet` and `RcppML` packages for NMF analysis. These packages leverage the `RcppML` implementation of NMF for use with snRNA-seq/scRNA-seq data. `RcppML` is much faster and more memory efficient than other NMF methods (67). Additionally, its diagonal scaling approach allows for L1/L2 regularization and reproducible NMF loadings.

We used the `cross_validate_nmf()` function from `singlet` to perform cross-validation of different NMF ranks (174). For a given number of replicate runs, this function randomly splits the data into training and test sets of a user-specified size. For each replicate, NMF is performed for a given number of ranks in the training set until the tolerance threshold is met. Model overfitting is also measured by a tolerance of overfit metric. After 5 iterations, the factorization’s performance is evaluated in the test set and test set MSE is calculated. At each successive iteration thereafter, the test set MSE is calculated compared to the test set MSE at 5 iterations. If the test set MSE for a given iteration is greater than test set MSE at iteration 5 by a specified value, the model is considered to be overfit. We cross-validated using 3 replicates with parameters tolerance of fit=10^-3^ (`singlet`-recommended threshold for cross-validation), tolerance of overfit=10^-4^, and L1=0.1 across ranks *k*=5, 10, 50, 100, 125, 150, and 200. Test set MSE at tolerance of fit decreased dramatically until k=100, and did not clearly decrease at *k*=125, *k*=150, or *k*=200 (**Extended Data Fig. 39**). Additionally, model overfitting was observed at *k*=150 and *k*=200 for all 3 replicates. Because models with *k*=100 and k=125 performed similarly and did not overfit the test data, we *k*=100 for the sake of interpretability, reasoning that 100 patterns would be more easily annotated than 125. We performed the final factorization using k=100 and tolerance of fit=10^-6^ (`singlet`-recommended threshold for high-quality factorization results >= 10^-5^). The final feature matrix *W* is presented in **Supplementary Table 9**.

#### NMF label transfer approach

We took a transfer learning approach to predict pattern weights for the NMF patterns identified in human snRNA-seq data across other target datasets (**Extended Data Fig. 38**). For every observation in the target dataset, we generated an observation-level weight for each *k*=100 NMF patterns learned from the snRNA-seq data that we used to investigate transcriptional variation in different data modalities and experimental designs. We used the `project()` function from the `RcppML` (67) package for this purpose, which makes use of a fast implementation of non-negative least squares (175). Using the snRNA-seq data as the source, we mathematically project the *W* matrix with *k*=100 NMF patterns into a second, target dataset based on the target matrix (*A’*) with dimensions *i’* genes and *j’* observations to generate the target coefficient matrix of *H’*:

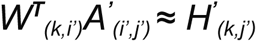

Note that *i’* is the number of genes present in both source snRNA-seq (*A*) and target (*A’*) gene expression matrices. To assess the validity of NMF pattern transfer, we examined the frequency of non-zero weighted observations for each NMF pattern in the target coefficient matrix (*H’*) and used the empirical cumulative distribution function to determine a minimum threshold for NMF patterns.

#### NMF transfer to SRT

Since the goal of NMF in this manuscript is to integrate SRT and snRNA-seq data, after NMF transfer to SRT we excluded patterns with non-zero weights in <1050 spots (**Extended Data Fig. 40C**). We looked for NMF patterns that might represent donor sex by searching the top 50 genes weighted to each pattern for *XIST* and for genes located on the Y chromosome. We only found one pattern with *XIST* in the top 50 genes (nmf37) and one pattern with more than one chrY gene in the top 50 (nmf27) (**Extended Data Fig. 41**). These patterns were removed from downstream analysis.

We utilized average nuclei weights across the *n*=60 cell types and found that NMF patterns could be classified as either “general”, equally distributed across many cell types, or “specific” to a particular cluster or cell type (**Extended Data Fig. 40A**). We found that patterns excluded from downstream analysis due to low mapping to SRT spots were both general and specific. The specific patterns removed from analysis corresponded to small snRNA-seq clusters and after removal of patterns with <1050 spots the only cell type groupings without a corresponding specific pattern were the thalamus and Cajal Retzius neurons, which were similarly enriched for nmf23. We observed that that after transfer to SRT general patterns stayed general and that specific patterns corresponded with appropriate cell types (**Extended Data Fig. 40D**), with one exception. We found that nmf84 was labeled as “general” based on nuclei-level weights across the snRNA-seq clusters, but, when projected to the SRT dataset was highly weighted in spots annotated to the choroid plexus spatial domain (**Extended Data Fig. 42A**). We determined that nmf94 encapsulated a *TTR*-dominant transcriptional program that recapitulated the donor-bias of CP nuclei and was distinct from the *TTR*-dominant CP-specific transcriptional program captured by nmf48 (**Extended Data Fig. 42B-E**). When nmf94 was projected onto the SRT dataset, the nuanced donor-specific effect was not conserved due to the strong importance of *TTR* expression in determining observation-level weights, and nmf94-weighted spots were nearly identical to nmf48-weighted spots (**Extended Data Fig. 42F-K**). This highlights the importance of close examination of NMF patterns to determine the biological or technical significance of the gene expression patterns they represent and whether they can be interpreted the same way in a dataset they are projected into.

To ascertain if additional NMF patterns captured donor-specific effects that could confound biological interpretation in downstream analysis, we examined the donor-level representation in non-zero weighted observations across all general and specific NMF patterns (**Extended Data Fig. 43**). We identified two patterns (nmf91 and nmf20) that were equally represented across snRNA-seq samples but exhibited substantial enrichment for two donors when projected to the SRT dataset (**Extended Data Fig. 47A and C**). These patterns were characterized by stimulus- and activity-dependent genes (**Extended Data Fig. 47B and D**). The increased abundance of these genes and the increased weight of nmf91 and nmf20 in nuclei across donors is likely due to the increased sensitivity of snRNA-seq to detect these sparsely expressed transcripts (**Extended Data Fig. 47E-F**). Thus, the apparent donor bias seemingly introduced when nmf91 and nmf20 are projected onto the SRT dataset (**Extended Data Fig. 47G-H**) is likely biologically meaningful and highlights specific donors in whom stimulus-dependent transcripts were enriched.

#### Gene set enrichment analysis

To validate the biological relevance of genes highly weighted to individual NMF patterns, we performed gene set enrichment analysis (GSEA) on patterns explored in **Figure 4**. GSEA was implemented with `fgsea` (176) utilizing `reactome.db` pathways (177), limited to pathways annotated to *Homo sapiens*. We used the `fgseaMultilevel` function to perform our analysis, specifying a minimum gene set size of 15 and maximum of 500. Due to the non-negative nature of pattern weights, enrichment scores were calculated for one-tailed tests by setting the score type parameter to `“pos”`. Non-zero gene-level weights were used for the gene score, resulting in 4661 genes for oligodendrocyte-specific nmf44, 9030 genes for astrocyte-specific nmf81, 7616 genes for trans-neuronal nmf13, and 7675 genes for trans-neuronal nmf7. Results were evaluated as significant at a FDR<.05. The FDR-adjusted *p*-value and normalized enrichment score (NES) for these tests are reported in **Figure 4G-J**.

#### NMF transfer to mouse electroconvulsive stimulation snRNA-seq dataset

For pattern transfer to the mouse electroconvulsive stimulation (ECS) snRNA-seq data (93), we downloaded the processed data object from the GitHub repository for the original mouse study and utilized the full dataset (*n*= 15,990 nuclei) (https://github.com/Erik-D-Nelson/ARG_HPC_snRNAseq/blob/main/processed_data/sce_subset.rda.xz). Information on how these data were processed prior to download can be found in the same GitHub repository (https://github.com/Erik-D-Nelson/ARG_HPC_snRNAseq/tree/main/code). To assist in interpretability with our human datasets, we identified human orthologs for mouse genes in the ECS dataset and removed all genes without orthologs. We matched these orthologs with genes in the *W* matrix and discarded any genes not included in both datasets. Following removal of genes without orthologs and genes not included in both datasets, we retained 17,557 genes. We used normalized, log_2_ transformed counts from the mouse ECS snRNA-seq dataset as the *A’* matrix. Following label transfer, patterns were normalized to sum to 1 in each target dataset. We removed patterns with non-zero weights in <1000 nuclei in the mouse ECS snRNA-seq dataset (**Extended Data Fig. 48A-B**). To ease interpretation within the context of our human findings, we further subset to only NMF patterns that also mapped to >1050 SRT spots (**Extended Data Fig. 48C**). We found that the appropriate patterns corresponded to the relevant cell types (**Extended Data Fig. 48B-C**).

To investigate if the genes contributing to the differential weights of select NMF patterns between ECS GCs and sham GCs were associated with neuronal activation, we performed DE analysis using `scran` to find markers with a binomial test. The ECS snRNA-seq data was subset to GCs and limited to the genes with non-zero weights for the NMF pattern being tested (tests run for nmf91, nmf20, nmf10, and nmf14). Highly significant (-log_10_(FDR)>30 and absolute value of the log_2_ fold change >1) differences in gene expression were examined for genes that were highly weighted to the NMF pattern being tested.

We observed that nmf55 nuclei weights were higher in ECS GCs compared to sham GCs (**Extended Data Fig. 48C carrot**). However, 8 of the top 10 genes weighted to nmf55 encode ribosomal subunits (*RPS24*, *RPL26*, *RPS27A*, *RPL32*, *RPS12*, *RPLP1*, *RPL34*, *RPS8*), and the remaining two genes are strongly associated with translation (*TPT1* and *EEF1A1*). We performed DE analysis on non-zero weighted nmf55 genes and found that although these genes were significantly increased in ECS GCs, the magnitude of log_2_FC and adjusted *p*-values were much attenuated compared to nmf91 and nmf20 that captured activity-dependent transcriptional programs (**Extended Data Fig. 48D**, **Figure 5F,G**). *De novo* protein production in response to neuronal activity is critical for subsequent changes in synaptic and circuit connectivity (178) and production of translational machinery like ribosomes likely facilitates such changes (179). Thus, although ribosome transcript-dominant nmf55 is associated with activated mouse GCs, it is unlikely that nuclei or spots highly weighted for nmf55 in our human snRNA-seq or SRT datasets were likely to have been recently activated. This is supported by the ubiquitous weighting of nmf55 across SRT spatial domains (**Extended Data Fig. 48E**). This example highlights the need for biological context in the interpretation of NMF patterns.

#### NMF transfer to mouse single-nucleus methylation sequencing (snmC-seq) dataset

We leveraged a mouse model linking single-nucleus methylation sequencing (snmC-seq) in neurons with their axon projection targets through retroviral tracing (106) to see if any NMF patterns generated from our human snRNA-seq data corresponded to HPC neurons with specific axonal projection targets. We downloaded the data from GEO (accession code GSE230782). As only the raw data was available, we reproduced the original authors’ pipeline by adapting code found in the GitHub repository for the snmC-seq study (https://github.com/zhoujt1994/EpiRetroSeq2023/blob/main/02.integration_mc_rna/03.process_RS2.ipynb). This required extracting CH gene body methylation counts to approximate gene expression, based on the premise that non-CpG cytosine methylation exhibits a strong inverse relationship with gene expression in neurons (180). Following the original authors’ specifications, data was QC processed and cell type clusters were generated by integrating with mouse single-cell RNA-seq data. We then subset the snmC-seq dataset to excitatory HPC and RHP neurons only (*n*= 2,004 nuclei). Then, following the initial study design, CH gene body methylation counts were extracted, log-scaled, and negated. This matrix was used for the *A’* matrix for pattern transfer. We identified human orthologs for mouse genes in this dataset and removed all genes without orthologs. We matched these orthologs with genes in the *W* matrix and discarded any genes not included in both datasets. Following removal of genes without orthologs and genes not included in both datasets, we retained 14,452 genes. Following label transfer, patterns were normalized to sum to 1 in each target dataset. We isolated the NMF patterns that we previously identified as corresponding to the tissue source of the snmC-seq dataset (CA patterns: nmf11, nmf63, nmf61, nmf15; prosubiculum/ subiculum patterns: nmf32, nmf40, nmf54; RHP patterns: nmf65, nmf22, nmf53, nmf68, nmf51, nmf45, nmf84, nmf27, nmf17) (**Extended Data Fig. 40A**). We then removed those patterns with non-zero weights in <45 snmC-seq nuclei (**Extended Data Fig. 49A**). We found that the remaining patterns correctly corresponded to nuclei collected from relevant brain regions (**Extended Data Fig. 49B**).

#### Thresholding of NMF patterns for SRT visualization

Binarization thresholds for individual NMF patterns were determined by taking one fifth of the 95% max spot-level weight. After determining the laminar organization Sub.1-specific nmf40 and Sub.2-specific nmf54 (**Figure 6A**), these patterns were thresholded (**Extended Data Fig. 49C, D**) and the union of the spots labeled were classified as Subiculum for visualization in **Figure 6C, E** along with the broad domains of Neuron, Neuropil, WM, and Vasc/CSF. The same approach was applied to additional NMF patterns to generate the following additional labels: CA1 (nmf15), CA3 (union of nmf11 and nmf63), ENT_sup (union of nmf84, nmf45, nmf27), and ENT_L5 (nmf51). To compile these 5 thresholds, only spots that passed a single one of the thresholds were kept for the combined annotation (**Extended Data Fig. 50**). This NMF-driven, combined, thresholded annotation was used for visualization and classification of the presubiculum in **Extended Data Fig. 51**.

#### Use of NMF to refine snRNA-seq cluster annotations and generate novel subicular complex marker genes

In SRT data, nmf65 was present in the deep RHP but was enriched in the deep subiculum (**Figure 6E**). In snRNA-seq data we identified two clusters (L6b and L6.1) that were labeled by nmf65; the entirety of the L6.1 cluster was highly weighted to nmf65, while only a portion of the L6b cluster was highly weighted to this pattern (**Figure 6F**). To ascertain if the two clusters corresponded to the separate populations of RHP and subiculum-adjacent spots labeled by nmf65, we split L6b into two portions by thresholding nmf65 weights (L6b_nmf65 with nmf65 weights >.00035). This grouping was used to determine that L6.1 expressed subiculum genes and this cluster was re-labeled to Sub.3 (**Extended Data Fig. 49E, F**).

Following NMF-driven assessment of the spatial localization of the clusters corresponding to subicular and retrohippocampal pyramidal neurons, the following snRNA-seq clusters were re-labeled: L6.1 to Sub.3, L2/3.1 to PreS, L2/3.4 to ENT.sup1, L2/3.6 to ENT.sup2a, L2/3.3 to ENT.sup2b, L2/3.2 to ENT.sup3, L5.1 to ENT.L5, L5.2 to RHP.CBLN2+ (based on **Extended Data Fig. 22B**), L6.2 to RHP.L6, and L6b to RHP.L6b.

Following the identification of the superficial subiculum (Sub.1), middle subiculum (Sub.2), deep subiculum (Sub.3), and presubiuculum (PreS), we performed DE analysis (`scran`-implented binomial test) in our snRNA-seq dataset focused on identifying genes that could specifically identify these regions (**Figure 6I**, **Extended Data Fig. 52**). To test for genes that distinguished the Sub.1 and Sub.2 clusters, we subset to regions with similar gene expression and that are spatially adjacent (CA1, ProS, Sub.3, PreS), testing for enrichment across 6 groups in total and only for genes with counts in >100 nuclei (22572 genes x 8783 nuclei). New Sub.1 markers were considered significant at -log_10_(FDR)>30 and have a log_2_FC of >1 vs each of the CA1, ProS, Sub.2, Sub.3, and PreS (146 genes). After filtering to genes with an average logcount expression of >1 in the Sub.1 nuclei, we identified 10 superficial subiculum marker genes: *AC007368.1*, *AL138694.1*, *ATP6V1C2*, *COL21A1*, *EBF4*, *FN1*, *NDST4*, *PARD3B*, *PRKCH*, *RAPGEF3* (**Extended Data Fig. 52A**). New Sub.2 markers were required to be significant at -log_10_(FDR)>30 and have a log_2_FC of >1 vs each of the CA1, ProS, Sub.1, Sub.3, and PreS (184 genes). After filtering to genes with an average logcount expression of >1 in the Sub.2 nuclei, we identified 7 middle subiculum marker genes: *GDNF-AS1*, *LHFPL3*, *MAMDC2*, *PCED1B*, *RORB*, *SULF1*, and *TRPC3* (**Extended Data Fig. 52B**). New ProS markers were required to be significant at -log_10_(FDR)>30 and have a log_2_FC of >1 vs each of the CA1, Sub.1, Sub.2, Sub.3, and PreS (17 genes). After filtering to genes with an average logcount expression of >1 in the ProS nuclei, we identified no unique prosubiculum marker genes.

To test for genes that distinguished the Sub.3 cluster from other subiculum layers and other deep RHP clusters, we subset to regions with similar gene expression and that are spatially adjacent (Sub.1, Sub.2, RHP.L6b, RHP.L6), testing for enrichment across 5 groups in total and only for genes with counts in >100 nuclei (21037 genes x 5647 nuclei). New Sub.3 markers were required to be significant at -log_10_(FDR)>30 and have a log_2_FC of >1 vs each of the Sub.1, Sub.2, RHP.L6b, and RHP.L6 (68 genes). After filtering to genes with an average logcount expression of >1 in the Sub.3 nuclei, we identified 21 deep subiculum marker genes: *AC007100.1*, *AC010967.1*, *AC023503.1*, *AC046195.2*, *AL356295.1*, *CD36*, *COL4A1*, *COL4A2*, *DISC1*, *FSTL5*, *GUCA1C*, *LINC01194*, *LINC01239*, *LINC01821*, *NR2F2-AS1*, *PLEKHG1*, *RASGEF1B*, *SCN7A*, *SCUBE1*, *SNCAIP*, and *VEGFC* (**Extended Data Fig. 52C**).

To test for genes that distinguished the PreS cluster from other subiculum layers and other superficial RHP clusters, we subset to regions with similar gene expression and that are spatially adjacent (Sub.1, Sub.2, ProS, ENT.sup1, ENT.sup2a, ENT.sup2b, ENT.sup3), testing for enrichment across 8 groups in total and only for genes with counts in >100 nuclei (22708 genes x 9835 nuclei). New PreS markers were required to be significant at -log_10_(FDR)>30 and have a log_2_FC of >1 vs each of the Sub.1, Sub.2, ProS, ENT.sup1, ENT.sup2a, ENT.sup2b, and ENT.sup3 (38 genes). After filtering to genes an average logcount expression of >1 in the PreS nuclei, we identified 5 presubiculum marker genes: *AC008662.1*, *AL161629.1*, *FSTL5*, *MDFIC*, and *WSCD1* (**Extended Data Fig. 52D**).

### 4.10 Data Visualization

#### Multi-array integration for Visium-SPG sample visualization

To enhance interactive visualization incorporating IF images alongside transcriptomic and spatial domain clustering, we utilized a robust, web-based data visualization tool tailored for spatially-resolved transcriptomics data, `Samui Browser` (112). We integrated all capture areas from each donor to enable visualization of the intact hippocampal structure by performing the following steps, based on a previously described strategy (181). First, we imported the high-resolution single-channel (DAPI) images from the `SpaceRanger` (version 2.0.0) outputs in `ImageJ` (182) to derive approximate transformations for all capture areas per donor. Due to image size constraints in `ImageJ`, high-resolution images featuring visible tissue landmarks were employed instead of full-resolution images or low resolution counterparts which lacked sufficient landmarks for accurate alignment. The "Tracing and Evaluation of Neural Anatomy using MipMaps" (`TrakEM2`) plugin (182–184) facilitated capture area alignment to reconstruct the hippocampal structure. Final transformations were exported in XML format. These transformations were extracted from the XML file of all capture areas per donor, then scaled to full resolution by applying scaling factors from the spaceranger JSON file using custom Python (v3.10.12) code, in particular `PIL` v10.0.0 and `numpy` v1.24.4. Then, the scaled transformations were applied to the full-resolution capture areas for each channel, and the transformed capture areas were placed onto a blank image encompassing the entire hippocampal structure from all capture areas from each donor. This composite image served as the basis for visualization in the `Samui Browser`. We then applied the same transformations to the Visium spot coordinates, where spot coordinates from all capture areas for each donor were concatenated into a single file. Overlapping spots from capture areas with wrinkled or curled tissue sections were disregarded. A `SpatialExperiment` object per donor was converted into an `AnnData` object in R, ensuring compatibility with the `Samui Browser`. Specifically, the R function `zellkonverter::SCE2AnnData()` (185) is used to convert the existing `SpatialExperiment` into an `AnnData` object, which allows gene expression and phenotype data to be accessible within python and therefore usable with the `Samui` API. We then used the combined image, spot coordinates and AnnData object to create the final `Samui` directory for each donor.

#### Multi-array integration for Visium-H&E sample visualization

A similar approach was also used for the Visium-H&E samples. The `Samui Browser` (112) was used as the final visualization tool. Using the `Fiji` distribution of `ImageJ,` images of the individual Visium slide capture areas were loaded into `ImageJ` and manually aligned. The transformations were saved as XML files. The `Spaceranger` JSON files were used to scale transformations to full resolution. The `zellkonverter::SCE2AnnData()` function was used to convert the existing `SpatialExperiment` into an `AnnData` object for use by `Samui Browser`. As in the Visium-SPG example, above, we used the combined image, spot coordinates and AnnData object to create the final product that is ready to be imported into `Samui.` All SRT data is available in a joint `Samui` browser.

#### iSEE apps

To enable exploration of the data, we created *iSEE* websites for pseudobulked snRNA-seq and spatial data as previously described (111,134). All interactive websites are available at research.libd.org/spatial_hpc/.

## 4.12 Data and Code Availability

Raw and processed data are available through Gene Expression Omnibus (GEO) under accession GSE264624 (113) and via ExperimentHub (https://bioconductor.org/packages/humanHippocampus2024). The code for this project is publicly available through GitHub at https://github.com/LieberInstitute/spatial_hpc. Analyses were performed using R version 4.3.2 with Bioconductor version 3.17 unless otherwise noted. Image alignment and transformation was performed with `ImageJ` (version 2.14.0).

## Acknowledgements

Portions of some figures were created with BioRender.com. We thank the LIBD neuropathology team, particularly James Tooke and Amy Deep-Soboslay, for curation of the brain samples and assistance with tissue dissections. We thank the staff and physicians at the brain donation sites, and the generosity of the brain donors and their families, without whom this work would not be possible. We thank Daniel R Weinberger, members of the Hansen-Hicks working group and the Spatial LIBD team for feedback on the manuscript. Finally, we thank the families of Connie and Steve Lieber and Milton and Tamar Maltz for their generous support.

## Author Contributions

Conceptualization: KM, SCP, SCH

Methodology: JRT, EDN, MT, JY, SCH

Software: JRT, EDN, MT, ADR, RAM, NJE, LCT, JY, SH, SCH

Validation: JRT, EDN, ADR, SCP

Formal Analysis: JRT, EDN, MT, ADR, JY, SCH

Investigation:JRT, EDN, MT, ADR, HRD, EAP, SHK, SVB, UMK, SCP

Resources: JEK, TMH

Data Curation: JRT, EDN, MT, ADR, SCH

Writing – original draft: JRT, EDN, MT, KRM, KM, SCP, SCH

Writing – review & editing: JRT, EDN, MT, ADR, HRD, RAM, SVB, SH, KRM, KM, SCP, SCH

Visualization: JRT, EDN, MT, ADR, SCP

Supervision: KRM, KM, SCP, SCH

Funding acquisition: KRM, KM, SCP, SCH

Project Administration: KRM, KM, SCP, SCH

