## Extended Data Figures and Tables for "An integrated single-nucleus and spatial transcriptomics atlas reveals the molecular landscape of the human hippocampus"

**Extended Data Fig. 1. Expression of *SNAP25* and *MBP* reveal neuron-rich and oligodendrocyte-rich regions across hippocampus (HPC) samples.**

H&E histology images (first column), $\log_2$ transformed normalized counts for *SNAP25* (pan-neuronal marker, second column), and $\log_2$ transformed normalized counts for *MBP* (oligodendrocyte marker, third column) for each donor (rows).

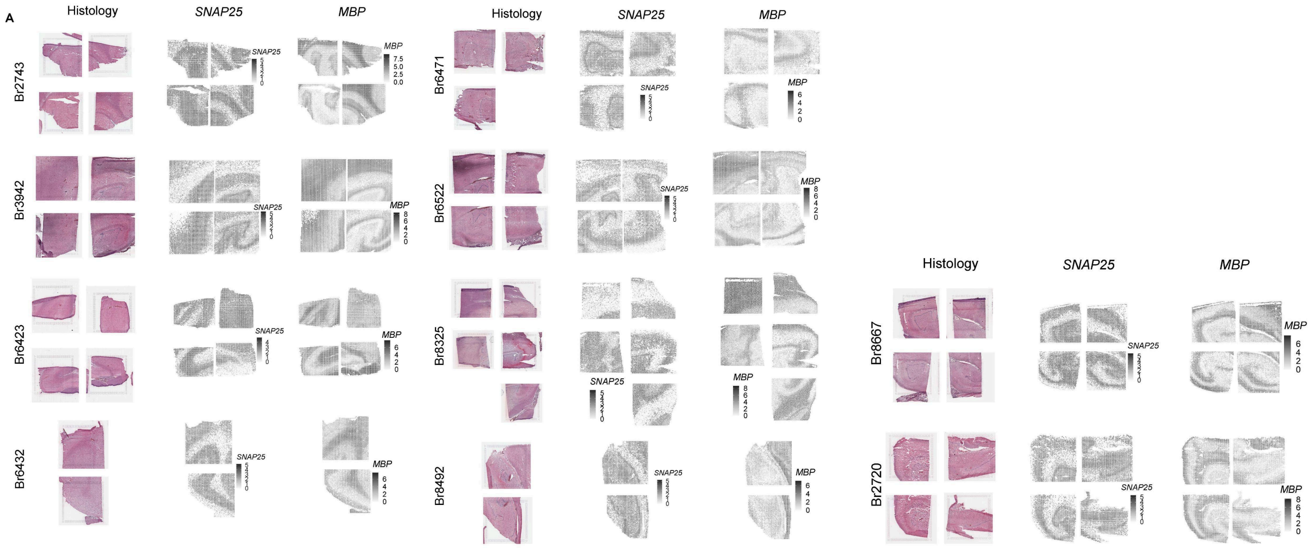

**Extended Data Fig. 2. Quality control (QC) metrics used to filter out low-quality spots.**

(**A**) Boxplots of library size (*y*-axis, total number of reads in a single spot) based on three median absolute deviation (3MAD) threshold calculated for each capture area independently. The spots are grouped by donor (*x*-axis) and are stratified by whether the spots were kept (black) or removed (red).

(**B**) Boxplots of number of detected genes (*y*-axis, total number of genes with at least one read in a single spot) based on three median absolute deviation (3MAD) threshold calculated for each capture area independently. The spots are grouped by donor (*x*-axis) and are stratified by whether the spots were kept (black) or removed (red).

(**C**) Bar plots demonstrate the number of spots (*y*-axis, log scale) per sample (individual bar) for each donor (*x*-axis). The plot is faceted by whether the spots were removed based on QC filter (top) or kept in the final dataset (bottom).

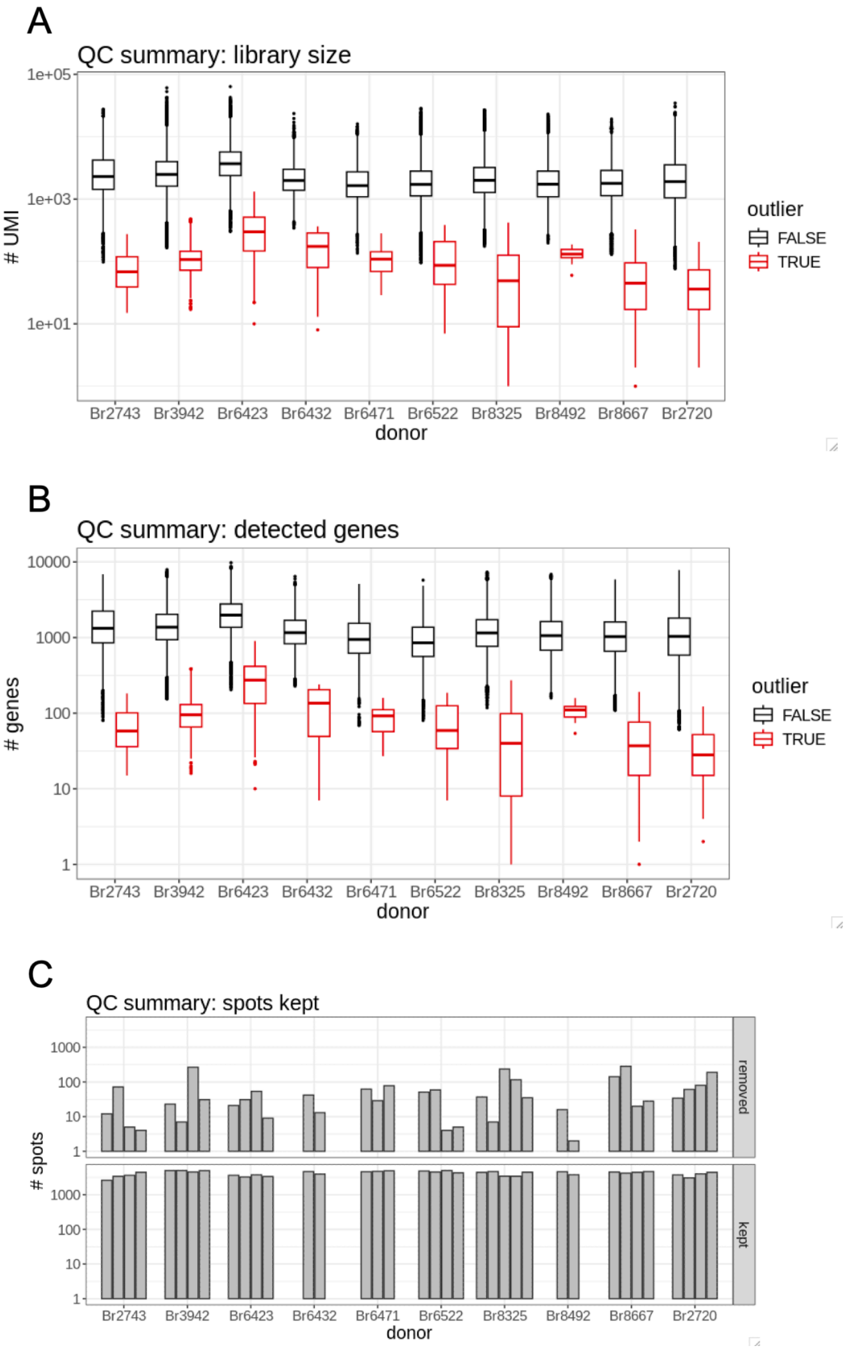

**Extended Data Fig. 3. Spot plot of proportion of reads mapping to mitochondrial genes per spot for each capture area.**

Within each capture area, each spot is colored by the proportion of reads mapping to mitochondrial genes (legend bottom left), which are enriched in hippocampal neuropil-enriched layers. Capture areas are grouped by donor number.

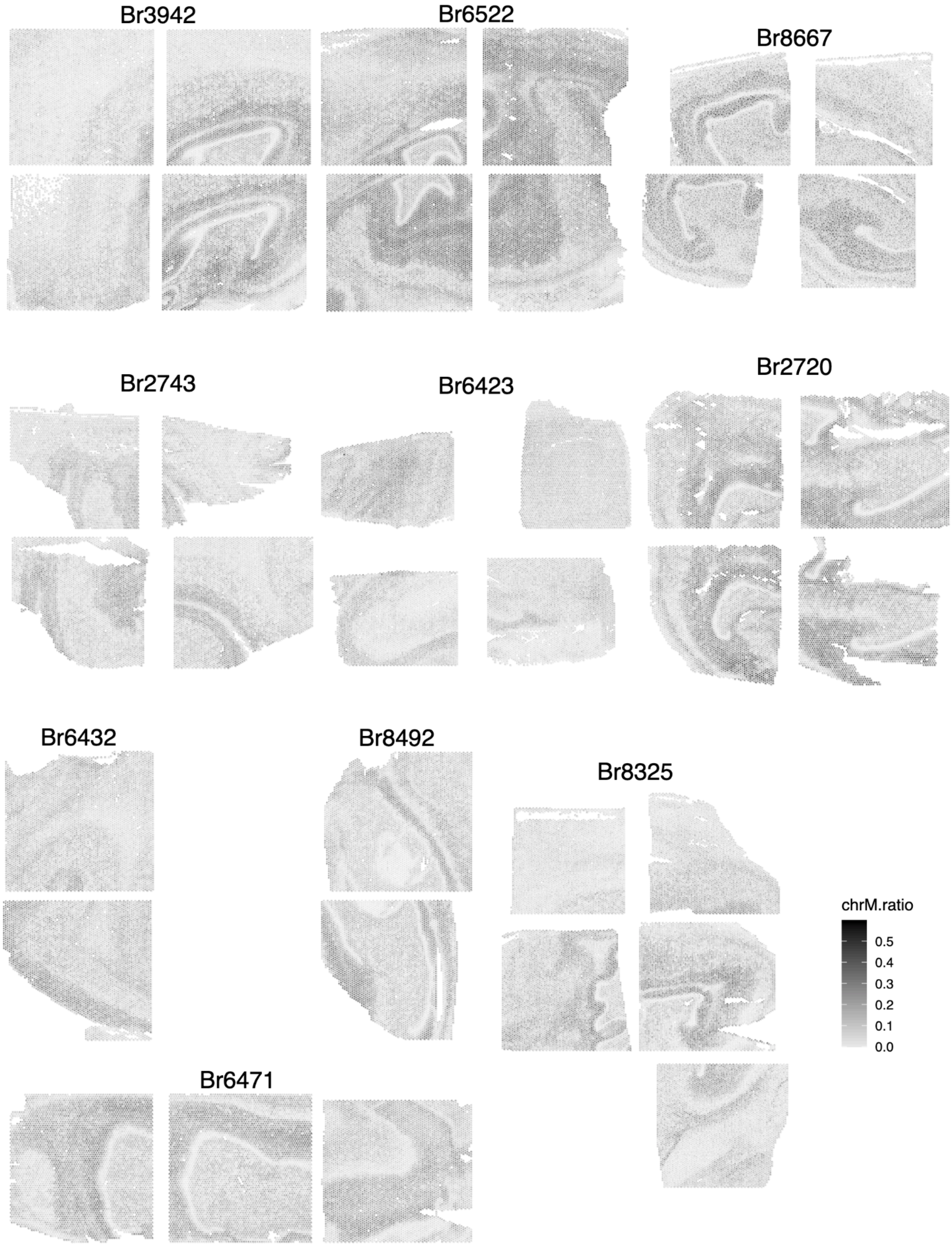
Br3942
Br6522
Br8667
Br2743
Br6423
Br2720
Br6432
Br8492
Br8325
Br6471
chrM.ratio
0.5
0.4
0.3
0.2
0.1
0.0

**Extended Data Fig. 4. Quality control (QC) metrics used to filter out low-quality nuclei.**

Boxplots of QC threshold (*y*-axis) calculated by three median absolute deviations (MAD) for the (**A**) maximum percent of reads originating from the mitochondrial genome (also known as mitochondrial fraction), (**B**) the minimum library size (total number of reads), and (**C**) the minimum number of genes with detected reads. Boxplots are grouped by sorting protocol (*x*-axis, propidium iodide (PI)+ or PI+NeuN+) and the individual values are the thresholds generated for each sample, colored by the round in which the sample was sequenced.

Boxplots of the number of nuclei excluded from each sample based on the raw 3MAD threshold for (**D**) maximum mitochondrial fraction, (**E**) minimum library size, and (**F**) minimum number of detected genes. Boxplots are grouped by sorting protocol (*x*-axis) and the individual values are the total number of nuclei removed from each sample based on the corresponding threshold in **A-C**, colored by the round in which the sample was sequenced.

(**G**) Rather than calculating the 3MAD maximum mitochondrial fraction threshold on a per-sample basis, the threshold for mitochondrial fraction was re-calculated based on the 3MAD mitochondrial fraction of high quality samples that had an original mitochondrial fraction threshold of <5%. Boxplots depict the new maximum percentage of reads coming from the mitochondrial genome for each sample with the revised threshold derived from high quality samples. Boxplots are grouped by sorting protocol (*x*-axis) and the individual values are colored by the round in which the sample was sequenced.

(**H**) All samples with zero nuclei removed with the original 3MAD threshold for minimum library size were subject to a revised minimum library size threshold of 1000 reads. Boxplots depict the updated library size threshold for all samples. Boxplots are grouped by sorting protocol (*x*-axis) and the individual values are colored by the round in which the sample was sequenced.

(**I**) All samples with zero nuclei removed with the original 3MAD threshold for minimum number of detected genes were subject to a revised minimum detected genes threshold of 1000 genes. Boxplots depict the updated detected genes threshold for all samples. Boxplots are grouped by sorting protocol (*x*-axis) and the individual values are colored by the round in which the sample was sequenced.

Boxplots of the number of nuclei excluded from each sample based on the revised 3MAD threshold for (**J**) maximum mitochondrial fraction, (**K**) minimum library size, and (**L**) minimum number of detected genes. Boxplots are grouped by sorting protocol (*x*-axis) and the individual values are the total number of nuclei removed from each sample based on the corresponding threshold in **G-I**, colored by the round in which the sample was sequenced.

### Original 3MAD QC thresholds (per-sample)

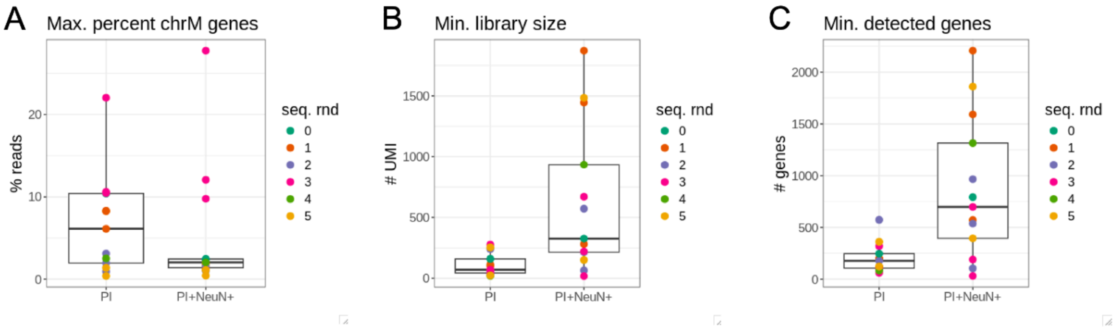


### Nuclei excluded with original 3MAD thresholds

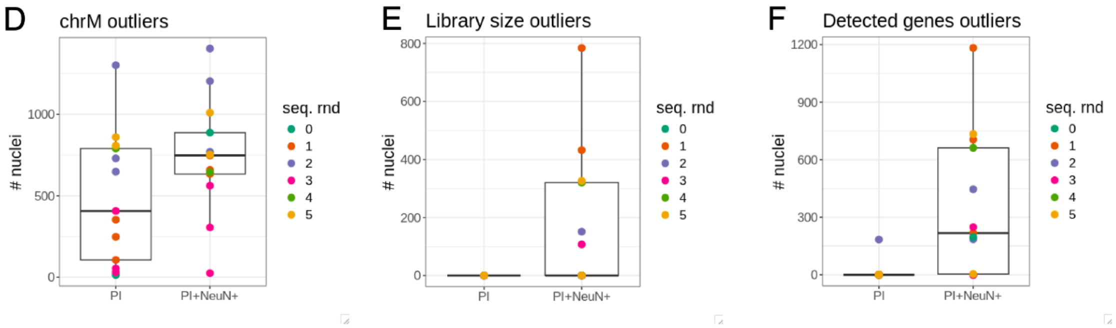


### Revised 3MAD QC thresholds

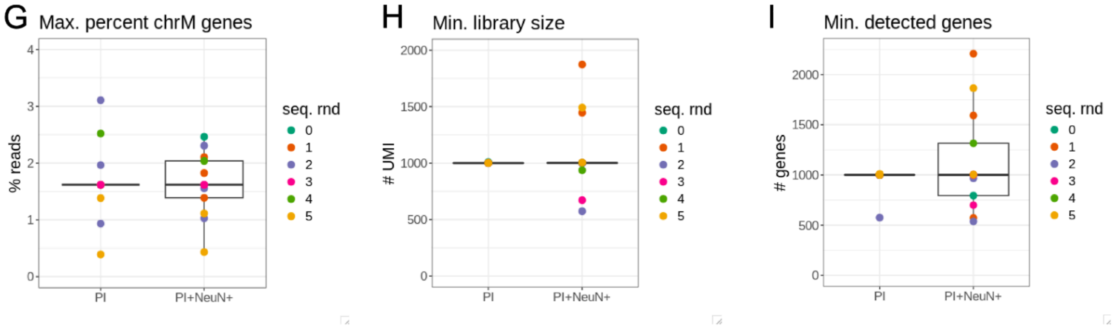


### Nuclei excluded with revised 3MAD thresholds

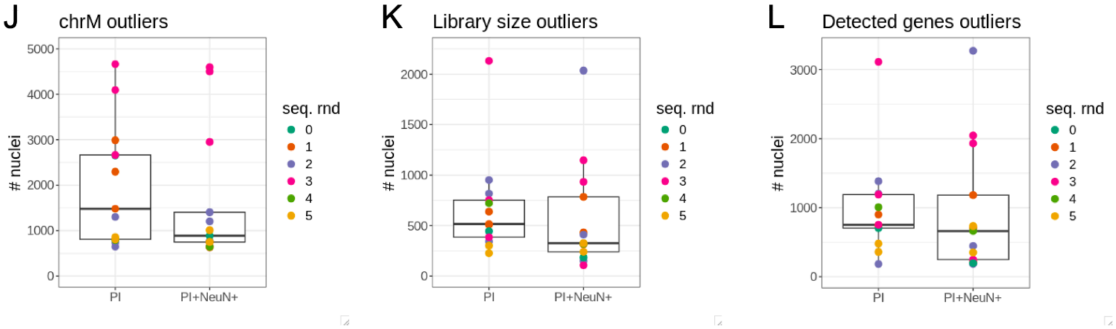

**Extended Data Fig. 5. Summary of quality control (QC) metrics in discarded and retained nuclei.**

(**A**) Violin plots show the distribution of mitochondrial fraction (*y*-axis; proportion of reads originating from mitochondrial genome) in populations of nuclei removed (left) or kept (right) based on QC thresholds, faceted by sorting protocol (propidium iodide (PI)+ samples (top) and PI+NeuN+ samples (bottom)). Each sample occupies a single position on the *x*-axis. The color of the violin plot indicates the round in which each sample was sequenced.

(**B**) Violin plots show the distribution of library size (*y*-axis; total number of reads) in populations of nuclei removed (left) or kept (right) based on QC thresholds, faceted by sorting protocol (PI+ samples (top) and PI+NeuN+ samples (bottom)). Each sample occupies a single position on the *x*-axis. The color of the violin plot indicates the round in which each sample was sequenced. The asterisk indicates sample 17c-scp for which the retained nuclei exhibit abnormally low library sizes.

(**C**) Violin plots show the distribution of detected genes (*y*-axis) in populations of nuclei removed (left) or kept (right) based on QC thresholds, faceted by sorting protocol (PI+ samples (top) and PI+NeuN+ samples (bottom)). Each sample occupies a single position on the x-axis. The color of the violin plot indicates the round in which each sample was sequenced. The asterisk indicates sample 17c-scp for which the retained nuclei exhibit abnormally few detected genes.

(**D**) Temporary cell clusters were easily identified as neuronal or non-neuronal based on the proportion of nuclei within the cluster that possessed at least one read of *SYT1* gene (raw counts).

(**E**) Violin plots present the distribution of the number of detected reads in each temporary neuronal cell cluster (*x*-axis), colored and stratified by in the neuronal nuclei originated from sample 17c-scp (red) or any other sample (grey). The dotted line indicates 5000 detected genes, highlighting the disproportionate number of 17c-scp nuclei with fewer than 5000 detected genes across all temporary neuronal clusters.

(**F**) Bar plots show the number of nuclei (*y*-axis) that were removed (left) or kept (right) in the snRNA-seq dataset after all QC filters were applied. The plots are faceted by sorting protocol (PI+ samples (top) and PI+NeuN+ samples (bottom)) and the color of the bar indicated the round in which the sample was sequenced. Each sample occupies a single position on the *x*-axis, and the outlier purple bar is sample 17c-scp for which all nuclei with fewer than 5000 detected genes were removed.

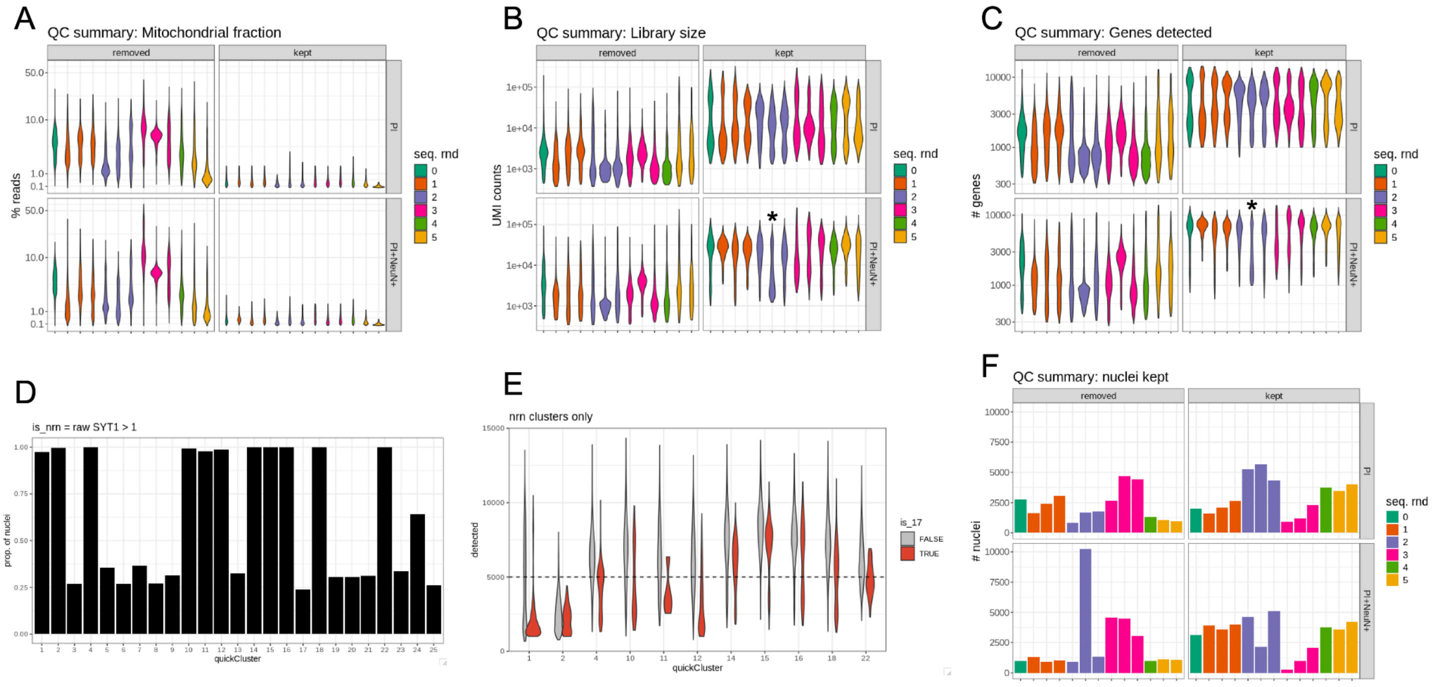
A
QC summary: Mitochondrial fraction
removed
kept
% reads
PI
PI+NeuN+
seq. rnd
B
QC summary: Library size
UMI counts
C
QC summary: Genes detected
### genes
D
is_nrn = raw SYT1 > 1
prop. of nuclei
quickCluster
E
nrn clusters only
detected
is_17
FALSE
TRUE
F
QC summary: nuclei kept
### nuclei

**Extended Data Fig. 6. Comparing `PRECAST` clustering results at *k*=16, *k*=17, and *k*=18.**

(**A**) Schematic of the `PRECAST` method (reproduced from Liu et al 2023 (30) Fig.1B) demonstrating its ability to integrate multiple tissue samples and reduce batch and technical effects when generating spatially-informed clusters.

(**B**) Plot showing Akaike Information Criterion (AIC, *y*-axis) by value of *k* (number of groups, *x*-axis) for *k*=2 through *k*=20. Dotted vertical lines mark breakpoints determined in segmented regression.

(**C**) Heatmaps for `PRECAST` clusters generated at (left) *k*=16, (middle) *k*=17, and (right) *k*=18 depict the expression ($log_2$ normalized counts) for genes (rows) averaged across each cluster (columns). On the far left, rows are annotated by the HPC region for which they are a marker. The additional cluster gained at *k*=17 vs *k*=16 is outlined in orange and appears to be a neuropil cluster, which exhibits more refined marker gene expression at *k*=18 (dotted outline). The additional cluster gained at *k*=18 vs *k*=17 is outlined in pink and appears to be a WM cluster.

(**D**) Pie charts of `PRECAST` cluster (facets) colored by manually annotated HPC region for clusters generated at (left) *k*=16, (middle) *k*=17, and (right) *k*=18. The additional cluster gained at *k*=17 vs *k*=16 is outlined in orange and appears to be a neuropil cluster, which is maintained at *k*=18 (dotted outline). The additional cluster gained at *k*=18 vs *k*=17 is outlined in pink and appears to be a WM cluster. THAL: thalamus, CTX: cortex, SUB: subiculum, PCL: pyramidal cell layer, GCL: granule cell layer, SGZ: dentate gyrus, subgranular zone, ML: dentate gyrus molecular layer, SL: stratum lucidum, SR: stratum radiatum, SLM: stratum lacunosum-moleculare, SO: stratum oriens, WM: white matter, CP: choroid plexus. Manual annotation abbreviations are also presented in **Supplementary Table 2**.

(**E**) Example spot plots (donor Br3942) of `PRECAST` clusters at (left) *k*=16, (middle) *k*=17, and (right) *k*=18 are colored by cluster. The new neuropil cluster at *k*=17 (tan) does not exhibit a spatial organization consistent with any known neuropil region, but the same cluster at *k*=18 exhibits a more stereotyped location to known neuropil-rich regions.

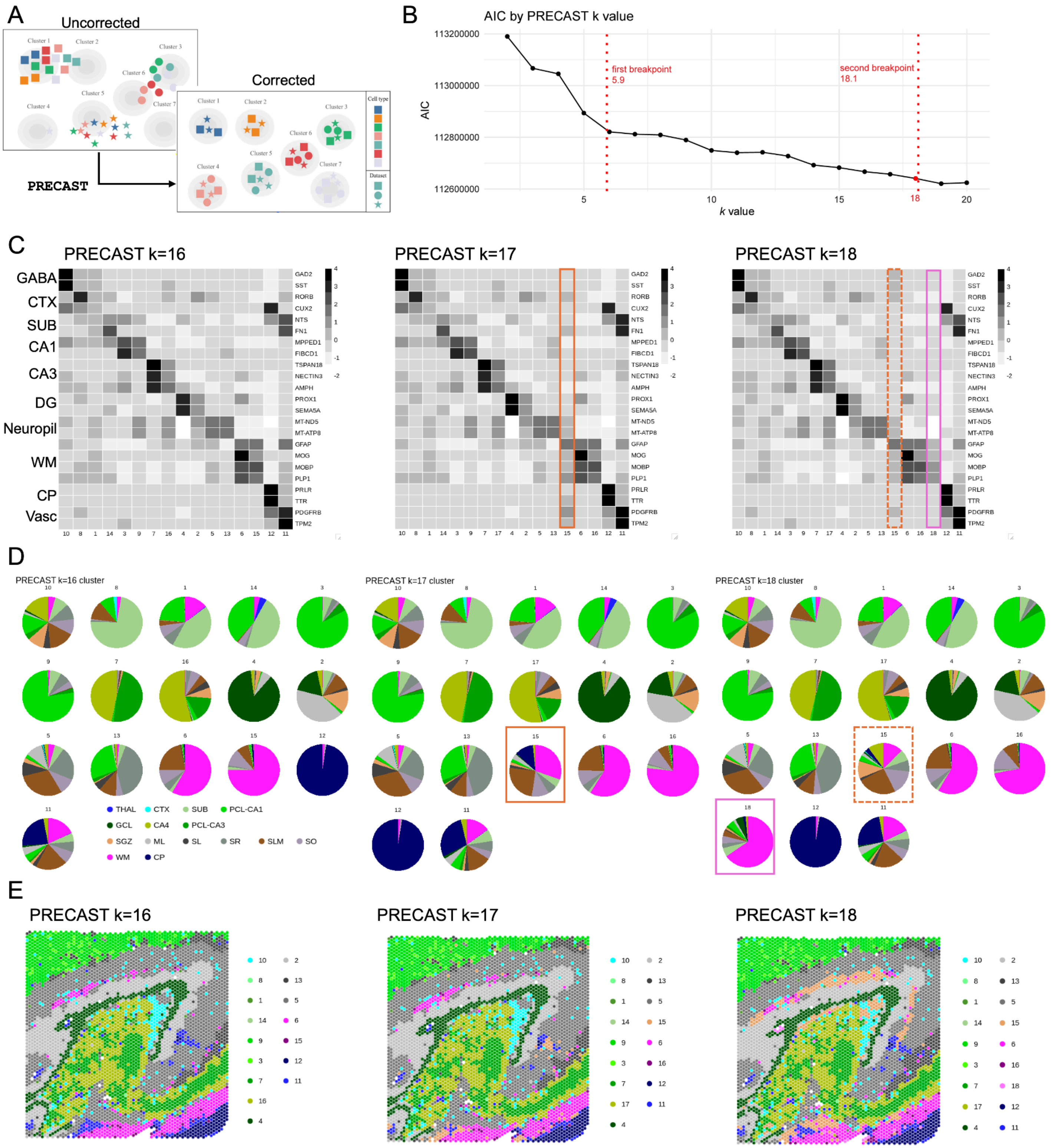
A
Uncorrected
Corrected
PRECAST
Cell type
Dataset
B
AIC by PRECAST k value
AIC
k value
first breakpoint
5.9
second breakpoint
18.1
113200000
113000000
112800000
112600000
5
10
15
18
20
C
PRECAST k=16
PRECAST k=17
PRECAST k=18
GABA
CTX
SUB
CA1
CA3
DG
Neuropil
WM
CP
Vasc
GAD2
SST
RORB
CUX2
NTS
FN1
MPPED1
FIBCD1
TSPAN18
NECTIN3
AMPH
PROX1
SEMA5A
MT-ND5
MT-ATP8
GFAP
MOG
MOBP
PLP1
PRLR
TTR
PDGFRB
TPM2
D
PRECAST k=16 cluster
PRECAST k=17 cluster
PRECAST k=18 cluster
THAL
CTX
SUB
PCL-CA1
GCL
CA4
PCL-CA3
SGZ
ML
SL
SR
SLM
SO
WM
CP
E
PRECAST k=16
PRECAST k=17
PRECAST k=18

**Extended Data Fig. 7. Histologically guided manual annotations of spatial domains.**

Using the H&E histology and cytoarchitecture and canonical marker gene expression, each spot was manually annotated to a spatial domain across the *n*=36 Visium-H&E capture areas. Spot color indicates the manual annotation (legend in bottom right). See **Supplementary Table 2** for abbreviations. Capture areas are grouped by donor number.

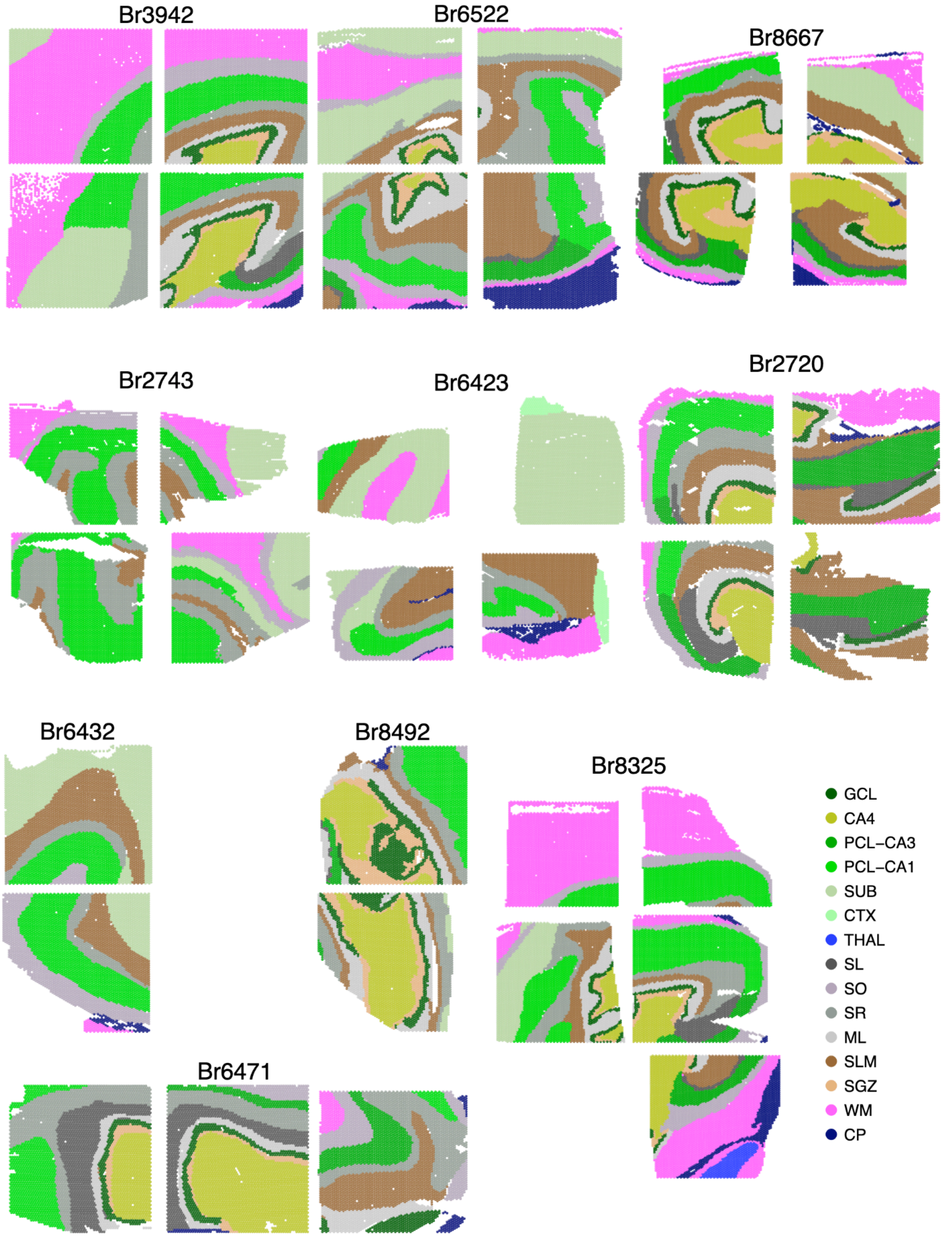
Br3942
Br6522
Br8667
Br2743
Br6423
Br2720
Br6432
Br8492
Br8325
Br6471
GCL
CA4
PCL−CA3
PCL−CA1
SUB
CTX
THAL
SL
SO
SR
ML
SLM
SGZ
WM
CP

**Extended Data Fig. 8. PRECAST cluster annotation and spatial domain definitions.**

(**A**) Heatmap showing expression of CA1 marker genes (*MPPED1, FIBCD1*, *CMLP*), CA2 marker gene *RGS14*, CA3 marker genes (*TSPAN18*, *NECTIN3*, *AMPH*), and mossy cell/ hilus/ CA4 marker gene *CARTPT* across *k*=18 annotated PRECAST clusters. See **Supplementary Table 2** for abbreviations.

(**B**) Boxplots showing number of nuclei per spot and library size across CA2-4.1, CA2-4.2, CA1.1, and CA1.2. Outliers were removed prior to plotting.

(**C**) Spot plot using example capture area from donor Br3942 highlighting manually annotated neuropil locations in comparison to annotated *k*=18 PRECAST clusters.

(**D**) Heatmap showing expression of HPC marker genes (CA1: *MPED1*, *FIBCD1*; CA3: *TSPAN18*, *NECTIN3*, *AMPH*; DG: *PROX1*, *SEMA5A*) and astrocyte-specific genes (*GFAP*, *AQP4*, *APOE*, *ID4*, *SOX2*) across neuropil spatial domains.

(**E**) Boxplots showing distribution of nuclei count per spot (*y*-axis) across final 16 spatial domains (*x*-axis).

Violin plots across final spatial domains (*x*-axis) showing (**F**) library size (total number of UMI counts) per spot, (**G**) number of detected genes per spot and (**H**) number of reads mapping to mitochondrial genes per spot.

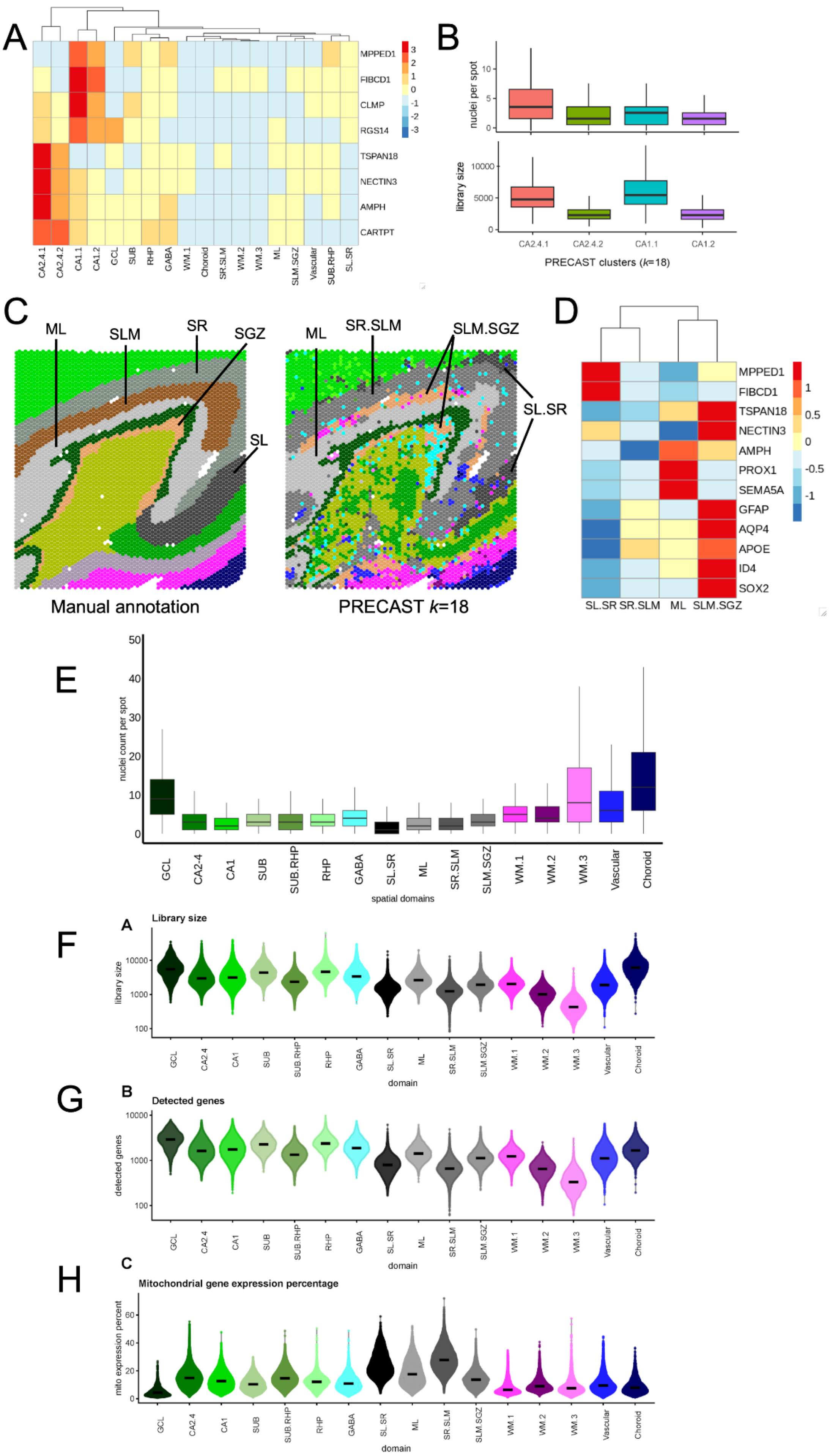

A
MPPED1
FIBCD1
CLMP
RGS14
TSPAN18
NECTIN3
AMPH
CARTPT
CA2.4.1
CA2.4.2
CA1.1
CA1.2
GCL
SUB
RHP
GABA
WM.1
Choroid
SR.SLM
WM.2
WM.3
ML
SLM.SGZ
Vascular
SUB.RHP
SL.SR
B
nuclei per spot
library size
PRECAST clusters (k=18)
C
ML
SLM
SR
SGZ
SL
SR.SLM
SLM.SGZ
SL.SR
Manual annotation
PRECAST k=18
D
MPPED1
FIBCD1
TSPAN18
NECTIN3
AMPH
PROX1
SEMA5A
GFAP
AQP4
APOE
ID4
SOX2
SL.SR SR.SLM ML SLM.SGZ
E
nuclei count per spot
spatial domains
GCL
CA2-4
CA1
SUB
SUB.RHP
RHP
GABA
SL.SR
ML
SR.SLM
SLM.SGZ
WM.1
WM.2
WM.3
Vascular
Choroid
F
A
Library size
library size
domain
G
B
Detected genes
detected genes
H
C
Mitochondrial gene expression percentage
mito expression percent

**Extended Data Fig. 9. Data-driven HPC domain annotations derived from spatial clustering (`PRECAST` *k*=18) across all *n*=36 Visium-H&E samples.**

Spatial clustering results from `PRECAST` k=18 were annotated to HPC spatial domains by examining marker gene expression and anatomical location. From the 18 `PRECAST` clusters we generated 16 spatial domains by merging clusters that were annotated to the same HPC subfields. Spot color indicates the final spatial domain annotation (legend in bottom right). See **Supplementary Table 2** for abbreviations. Capture areas are grouped by donor number.

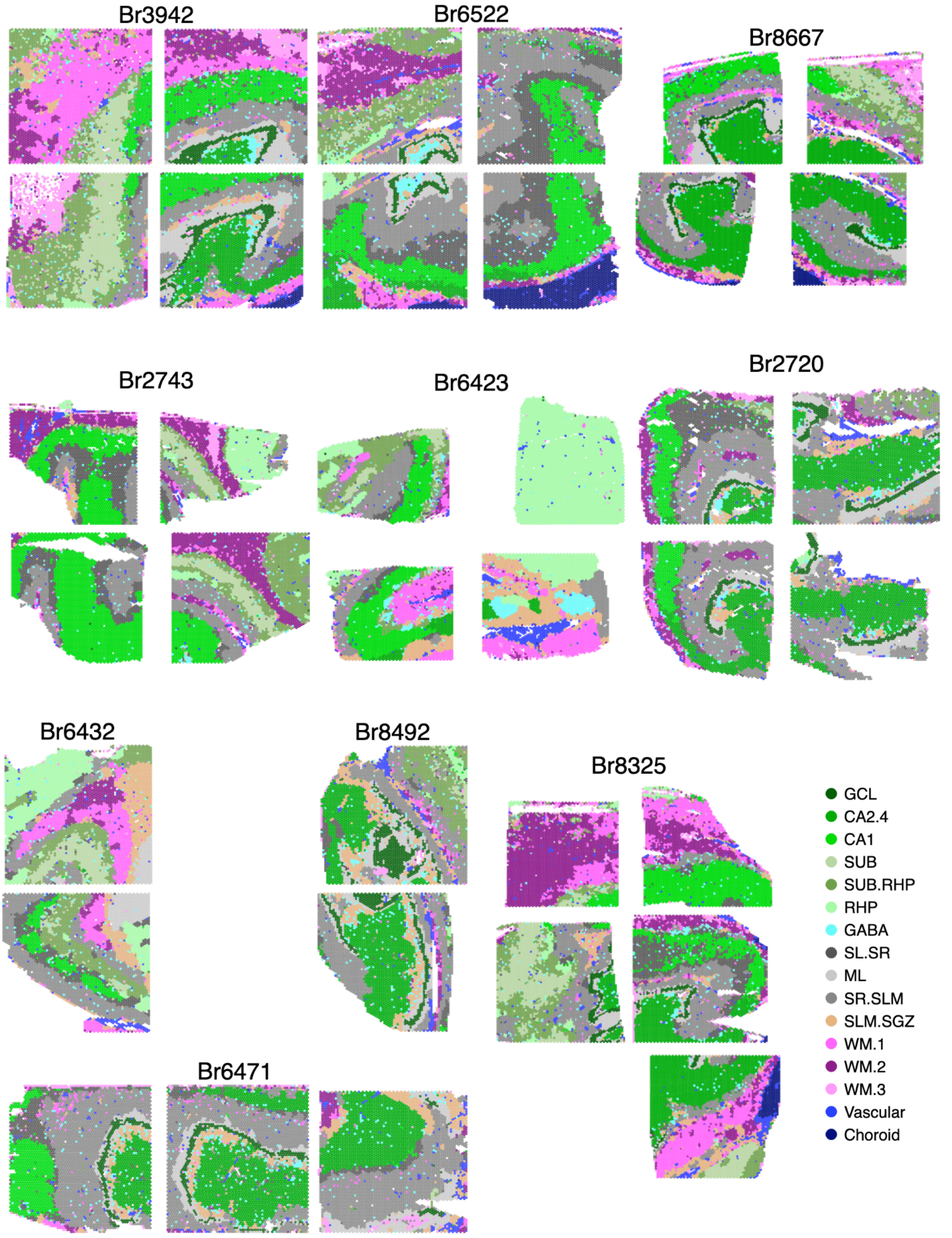
Br3942
Br6522
Br8667
Br2743
Br6423
Br2720
Br6432
Br8492
Br8325
Br6471
GCL
CA2.4
CA1
SUB
SUB.RHP
RHP
GABA
SL.SR
ML
SR.SLM
SLM.SGZ
WM.1
WM.2
WM.3
Vascular
Choroid

**Extended Data Fig. 10. Identification of thalamus and amygdala tissue in select SRT capture areas.**

(**A**) H&E histology images of Br8325 capture areas. Putative thalamic tissue is circled.

(**B**) Spot plot showing expression of *TCF7L2*, a thalamus-specific transcription factor (33). Spots are filled by $\log_2$ normalized counts. Spot borders are colored by spatial domain. See **Supplementary Table 2** for abbreviations.

(**C**) Spot plot showing expression of *SHOX2*, a thalamus-specific transcription factor (34). Spots are filled by $\log_2$ normalized counts. Spot borders are colored by spatial domain.

(**D**) Top: H&E histology image of example cryosection from Br6423 prior to scoring; approximate capture areas are outlined. Bottom: H&E histology images of Br6423 capture areas. Putative amygdala capture area (top right) is labeled.

(**E**) Spot plot showing expression of *SLC17A6*, which is enriched in human amygdala excitatory neurons (21). Spots are filled by $\log_2$ normalized counts. Spot borders are colored by spatial domain.

(**F**) Spot plot showing expression of *OPRM1*, which has been shown to be more highly enriched in amygdala than hippocampus (36). Spots are filled by $\log_2$ normalized counts. Spot borders are colored by spatial domain.

(**G**) Spot plot showing expression of *CDH22*, which is enriched in human amygdala excitatory neurons (21). Spots are filled by $\log_2$ normalized counts. Spot borders are colored by spatial domain.

(**H**) Spot plot showing expression of *CACNG4*, which has been shown to be more highly enriched in amygdala than hippocampus (35). Spots are filled by $\log_2$ normalized counts. Spot borders are colored by spatial domain.

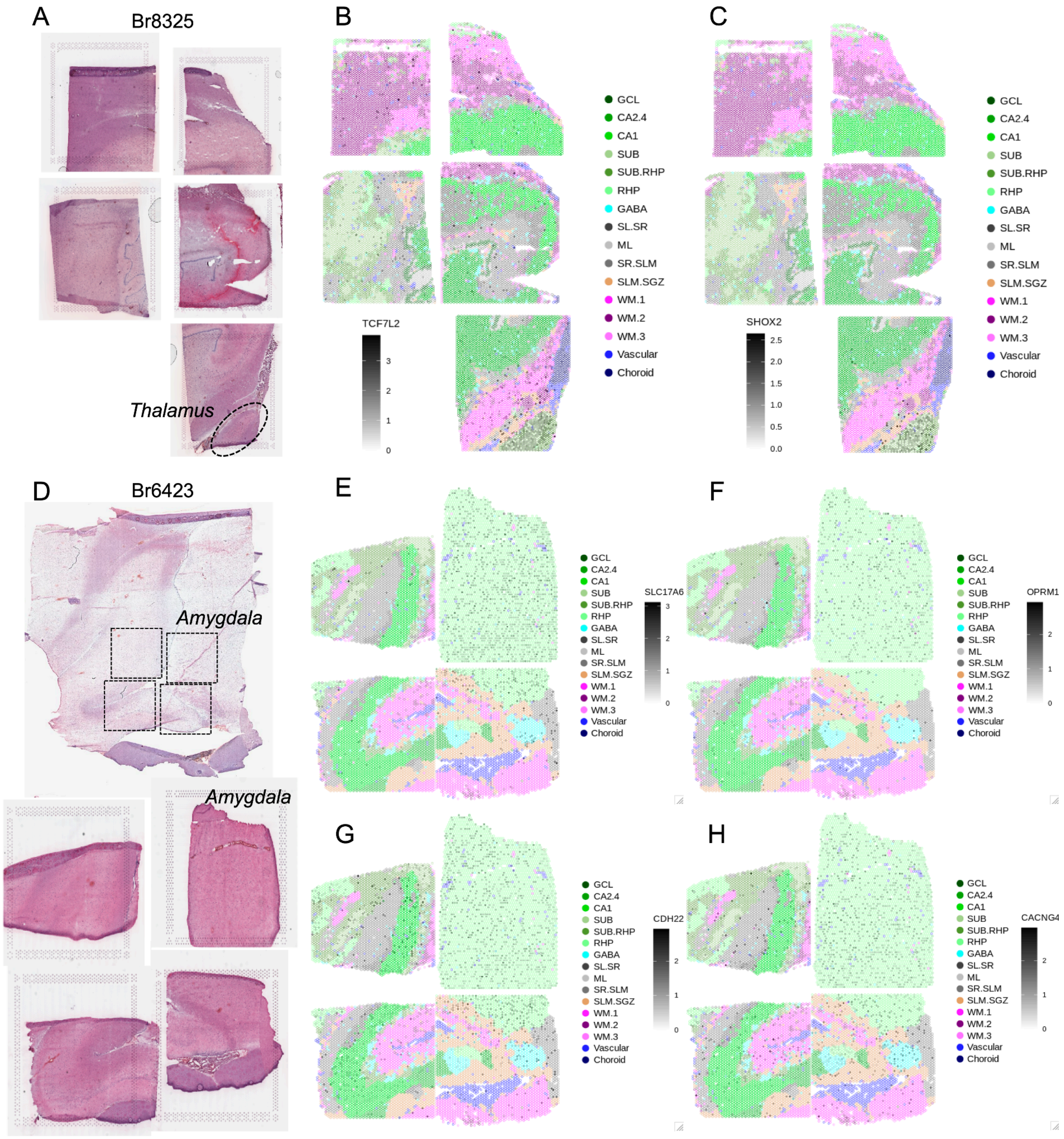

A
Br8325
Thalamus
B
C
GCL
CA2.4
CA1
SUB
SUB.RHP
RHP
GABA
SL.SR
ML
SR.SLM
SLM.SGZ
WM.1
WM.2
WM.3
Vascular
Choroid
TCF7L2
SHOX2
D
Br6423
Amygdala
Amygdala
E
F
G
H
SLC17A6
OPRM1
CDH22
CACNG4

**Extended Data Fig. 11. Spatial clustering results using `GraphST` (145) for *k*=16.**

`GraphST` clusters (color, legend in bottom right) are depicted across *n*=36 capture areas grouped by donor number.

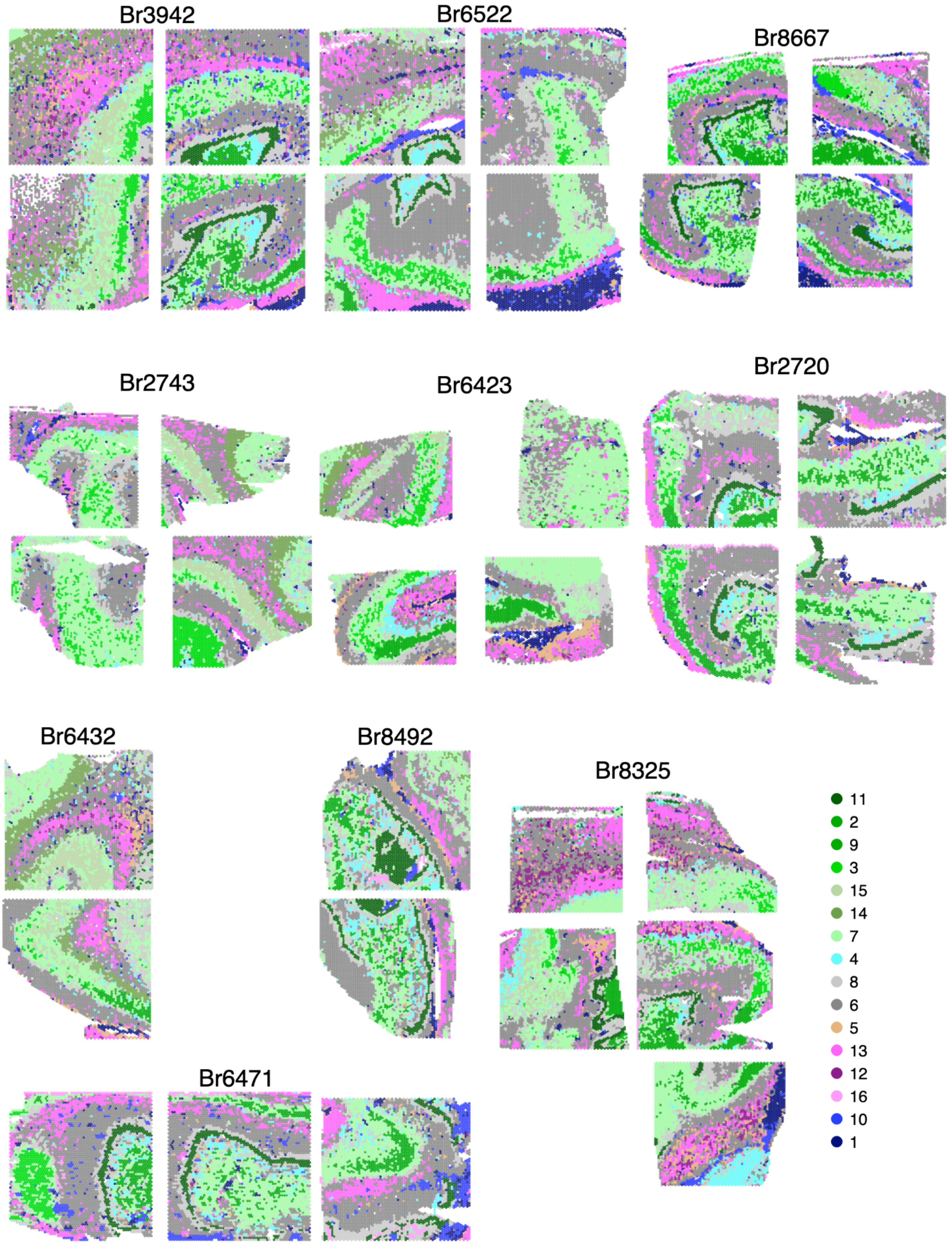

Br3942
Br6522
Br8667
Br2743
Br6423
Br2720
Br6432
Br8492
Br8325
Br6471
11
2
9
3
15
14
7
4
8
6
5
13
12
16
10
1

**Extended Data Fig. 12. Spatial clustering results using `BayesSpace` (147) for *k*=18.**

`BayesSpace` clusters (color, legend in bottom right) are depicted across by *n*=36 capture areas grouped by donor number.

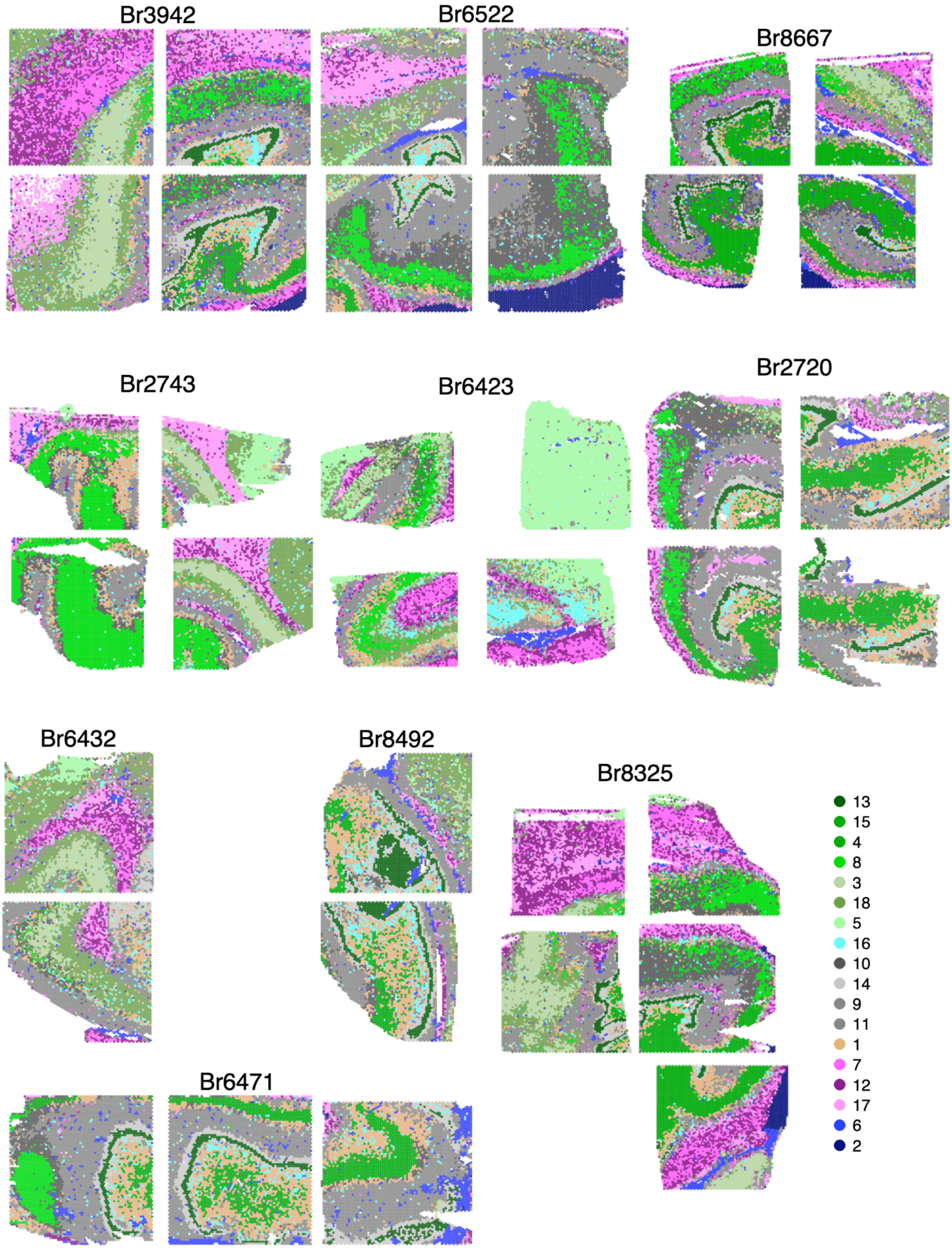

Br3942
Br6522
Br8667
Br2743
Br6423
Br2720
Br6432
Br8492
Br8325
Br6471
13
15
4
8
3
18
5
16
10
14
9
11
1
7
12
17
6
2

**Extended Data Fig. 13. Comparison of spot-level assignment based on different annotation techniques.**

Comparing `PRECAST` *k*=18 clusters (rows, amended to distinguish the limited amygdala and thalamus regions) to the manual annotations (left), `BayesSpace` clusters (middle), and `GraphST` clusters (right). In each case the color of the tile indicates the number of spots ($\log_{10}$ transformed) that were present in both the `PRECAST`-derived groups (rows) and the groups in the technique being compared (columns). See **Supplementary Table 2** for abbreviations.

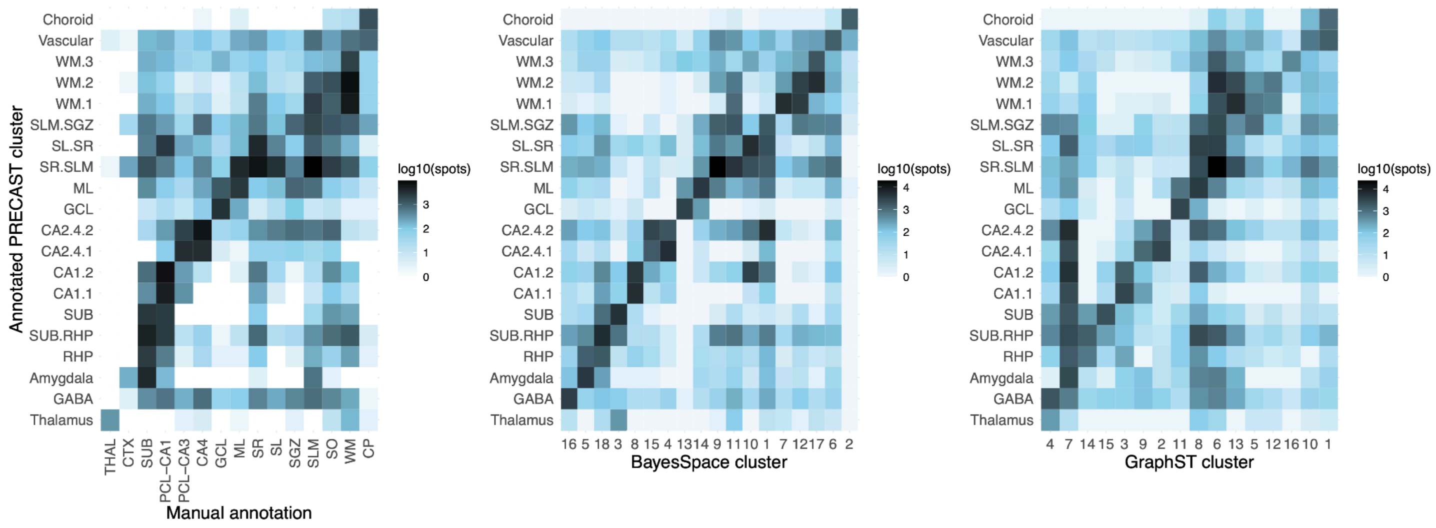

**Extended Data Fig. 14. Principal components plot using pseudobulked spatial data colored by domain.**

Principal component (PC) plots for the top 5 PCs (facets) after pseudo-bulking Visium spots across spatial domains (color) and 10 donors (total of 317 pseudo-bulked tissue samples). Facet labels include the percent variance explained for each PC. See **Supplementary Table 2** for abbreviations.

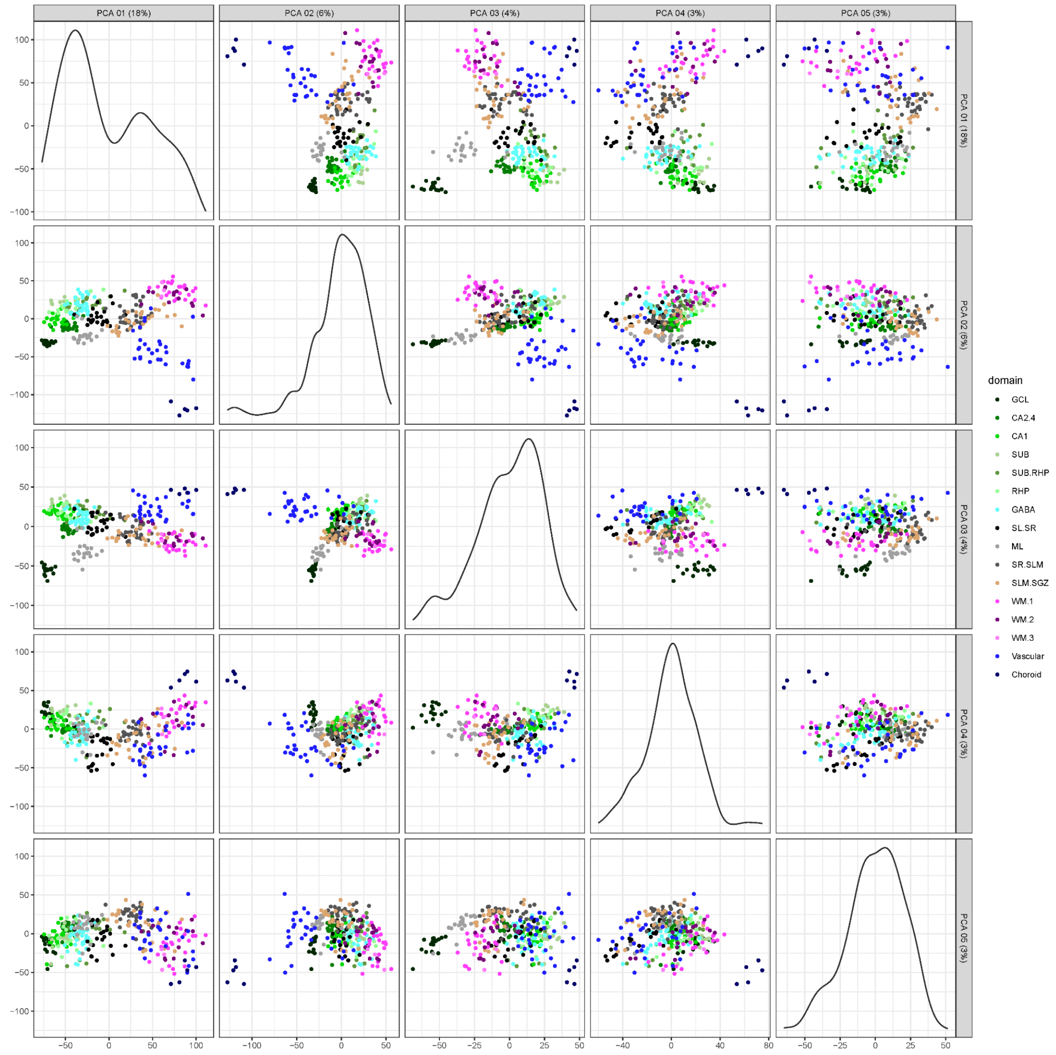

**Extended Data Fig. 15. Summary of gene-level variance explained by different variables.**

Density plot for the percent of gene expression variance (*x*-axis) explained for several experimental variables in the pseudobulked spatial data across the *n*=36 SRT samples. The spatial domains (red line) contribute the largest percent of gene expression variance, importantly more than capture area (green) or donor (purple).

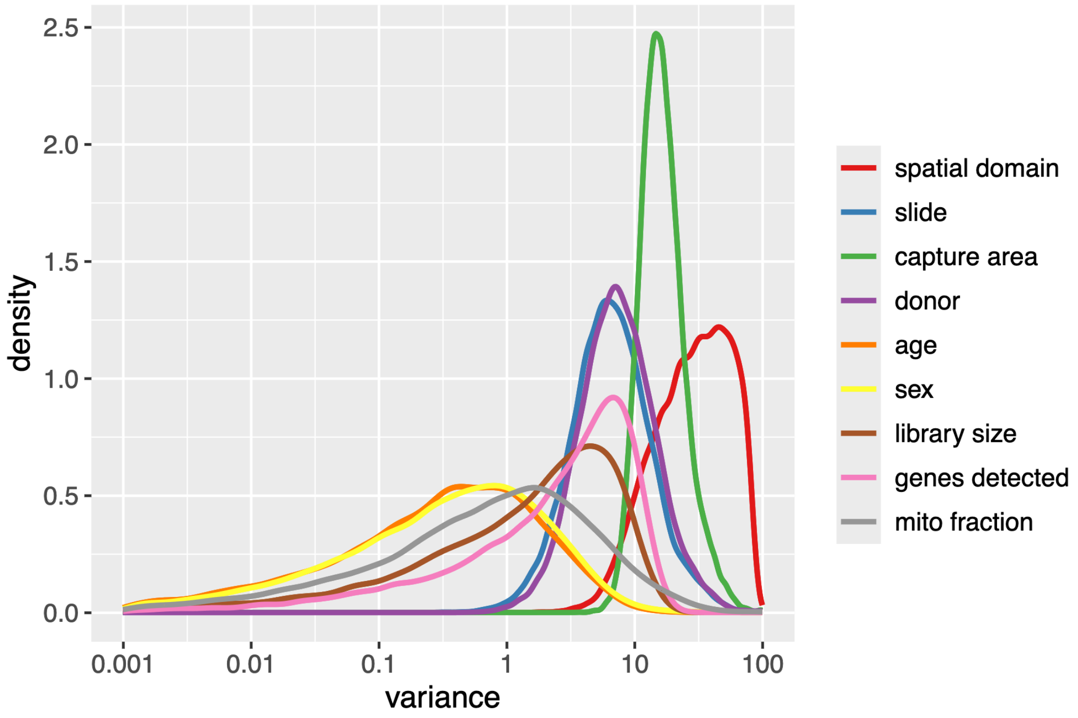

**Extended Data Fig. 16. Spatial domain-level differential expression analysis of spatially-resolved transcriptomics (SRT) data.**

Volcano plots illustrate results from pseudobulked differential expression (DE) analysis for each spatial domain, with $\log_2$ fold change on the *x*-axis and FDR adjusted, $-\log_{10}$ transformed *p*-values on the *y*-axis. Genes colored red pass both FDR and $\log_2$ fold change thresholds (FDR adjusted $p$-value < 0.01 and $\log_2$ fold change > 1). Top DE genes are labeled. See **Supplementary Table 2** for spatial domain (plot title) abbreviations.

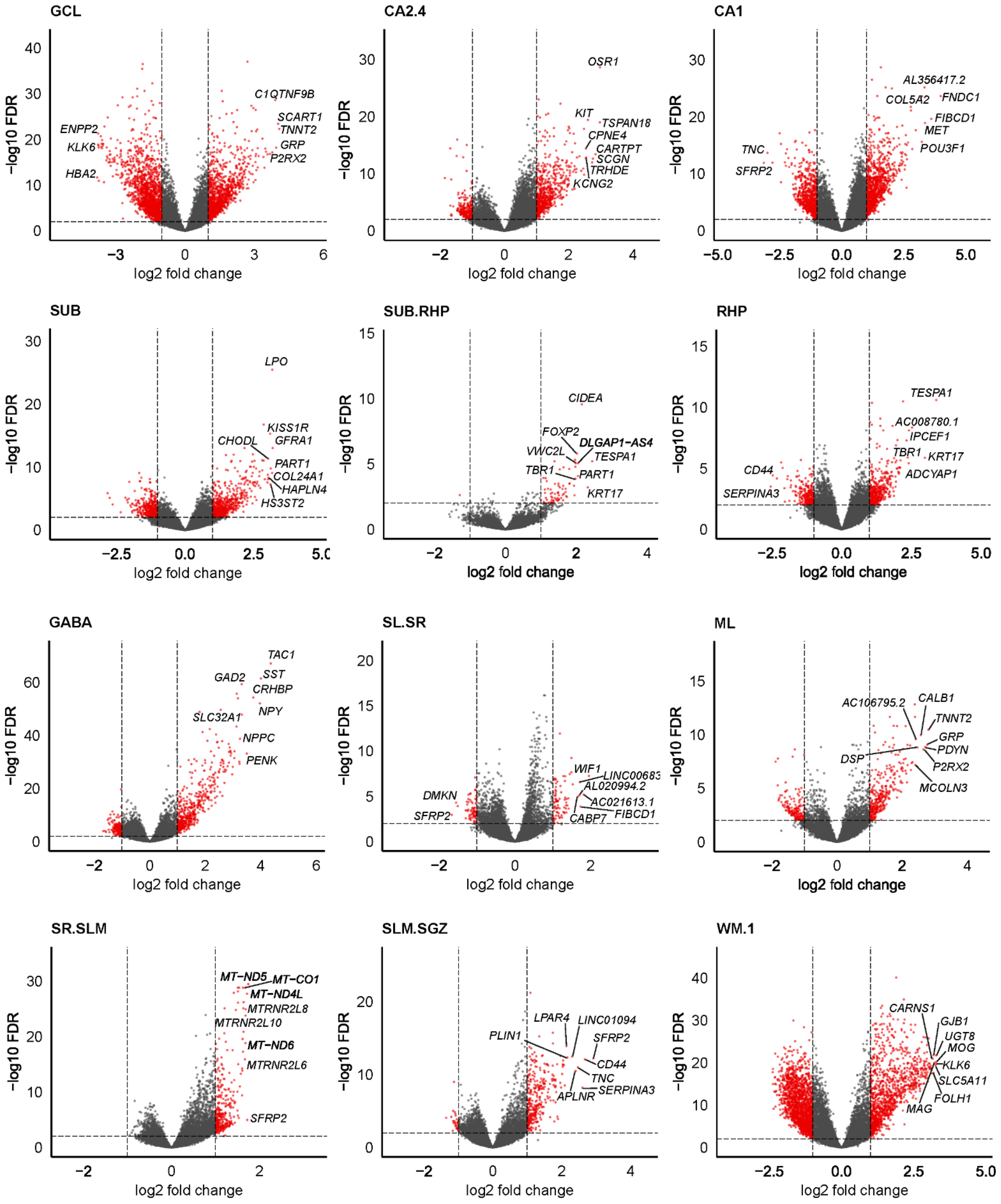
GCL
C1QTNF9B
SCART1
TNNT2
GRP
P2RX2
ENPP2
KLK6
HBA2
CA2.4
OSR1
KIT
TSPAN18
CPNE4
CARTPT
SCGN
TRHDE
KCNG2
CA1
AL356417.2
COL5A2
FNDC1
FIBCD1
MET
POU3F1
TNC
SFRP2
SUB
LPO
KISS1R
CHODL
GFRA1
PART1
COL24A1
HAPLN4
HS3ST2
SUB.RHP
CIDEA
FOXP2
DLGAP1−AS4
VWC2L
TESPA1
TBR1
PART1
KRT17
RHP
TESPA1
AC008780.1
IPCEF1
TBR1
KRT17
ADCYAP1
CD44
SERPINA3
GABA
TAC1
GAD2
SST
CRHBP
NPY
SLC32A1
NPPC
PENK
SL.SR
WIF1
LINC00683
AL020994.2
AC021613.1
FIBCD1
CABP7
DMKN
SFRP2
ML
CALB1
AC106795.2
TNNT2
GRP
PDYN
DSP
P2RX2
MCOLN3
SR.SLM
MT−ND5
MT−CO1
MT−ND4L
MTRNR2L8
MTRNR2L10
MT−ND6
MTRNR2L6
SFRP2
SLM.SGZ
LPAR4
LINC01094
PLIN1
SFRP2
CD44
TNC
APLNR
SERPINA3
WM.1
CARNS1
GJB1
UGT8
MOG
KLK6
SLC5A11
FOLH1
MAG
−log10 FDR
log2 fold change

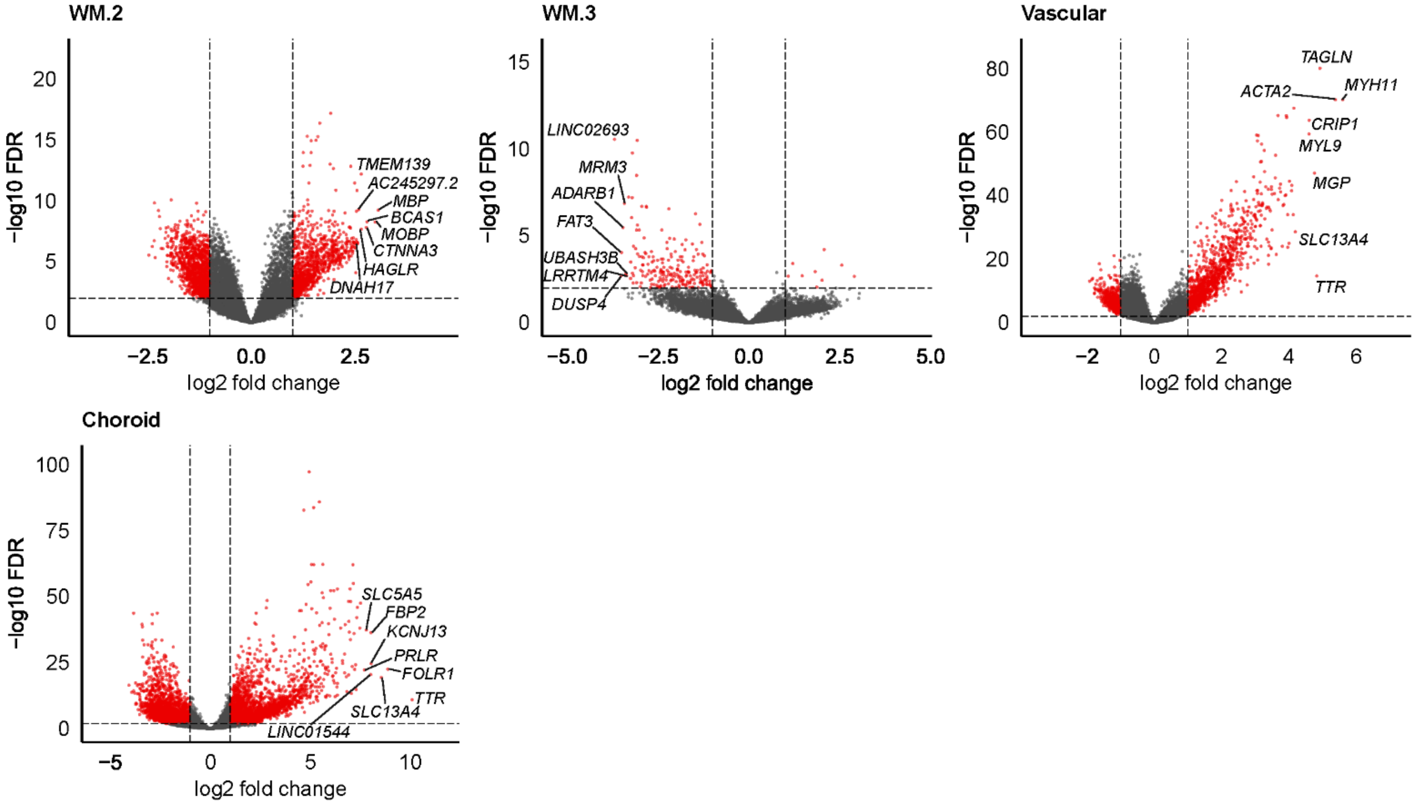

WM.2
−log10 FDR
20
15
10
5
0
TMEM139
AC245297.2
MBP
BCAS1
MOBP
CTNNA3
HAGLR
DNAH17
−2.5
0.0
2.5
log2 fold change
WM.3
−log10 FDR
15
10
5
0
LINC02693
MRM3
ADARB1
FAT3
UBASH3B
LRRTM4
DUSP4
−5.0
−2.5
0.0
2.5
5.0
log2 fold change
Vascular
−log10 FDR
80
60
40
20
0
TAGLN
ACTA2
MYH11
CRIP1
MYL9
MGP
SLC13A4
TTR
−2
0
2
4
6
log2 fold change
Choroid
−log10 FDR
100
75
50
25
0
SLC5A5
FBP2
KCNJ13
PRLR
FOLR1
TTR
SLC13A4
LINC01544
−5
0
5
10
log2 fold change

**Extended Data Fig. 17. Number of differentially expressed genes (DEGs) for each `PRECAST` spatial domain using enrichment model results.**

Bar plot showing the number of DEGs (*y*-axis, FDR adjusted *p*-value < 0.01) in each domain (*x*-axis). Upregulated genes are shown in red ($\log_2$ fold change > 1), while downregulated genes are shown in blue ($\log_2$ fold change < -1). See **Supplementary Table 2** for abbreviations.

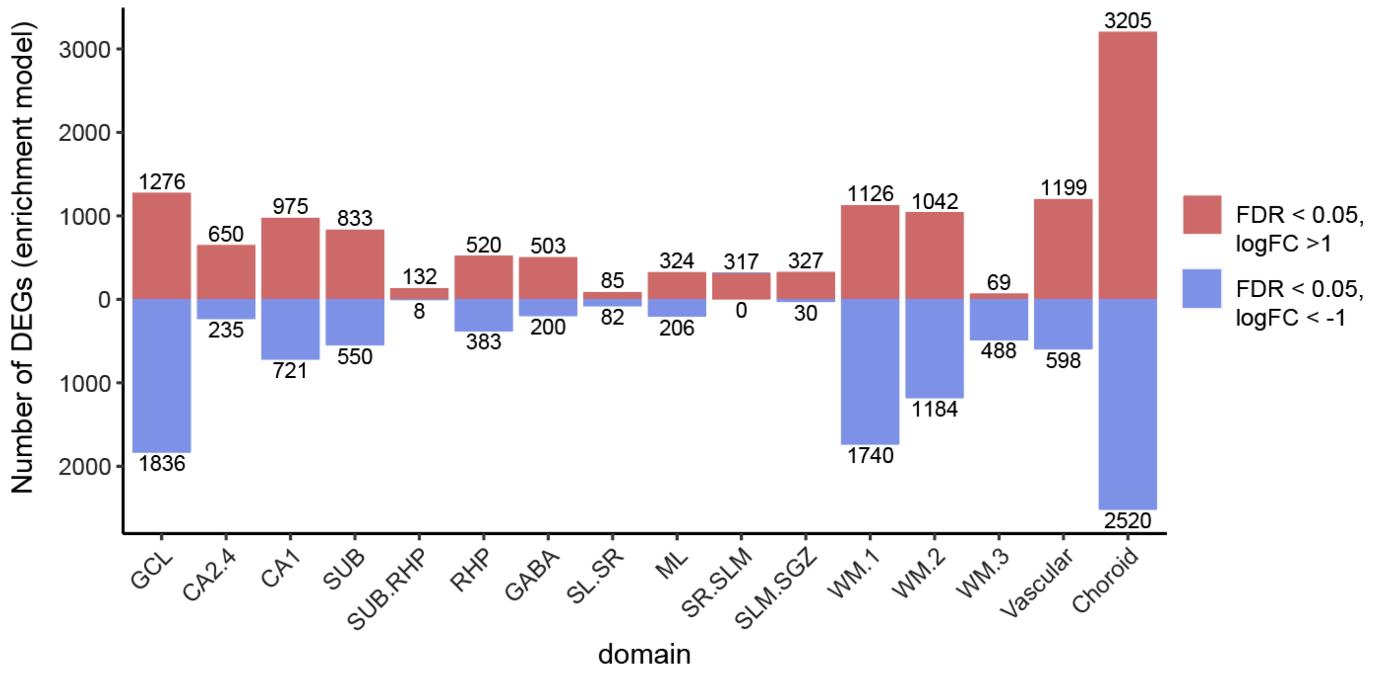

**Extended Data Fig. 18. Final batch correction for the hippocampus (HPC) single-nucleus RNA-sequencing (snRNA-seq) data.**

**(A)** Uniform manifold approximation (UMAP) representation of snRNA-seq data before mutual nearest neighbors (MNN) batch correction. Points are colored by donor and labeled by cell types. See **Supplementary Table 2** for abbreviations.

**(B)** UMAP representation of snRNA-seq data before MNN batch correction. Each facet shows all nuclei from a given donor. Points are colored by donor.

**(C)** UMAP representation of snRNA-seq data after MNN batch correction. Points are colored by donor and labeled by cell types.

**(D)** UMAP representation of snRNA-seq data after MNN batch correction. Each facet shows all nuclei from a given donor. Points are colored by donor.

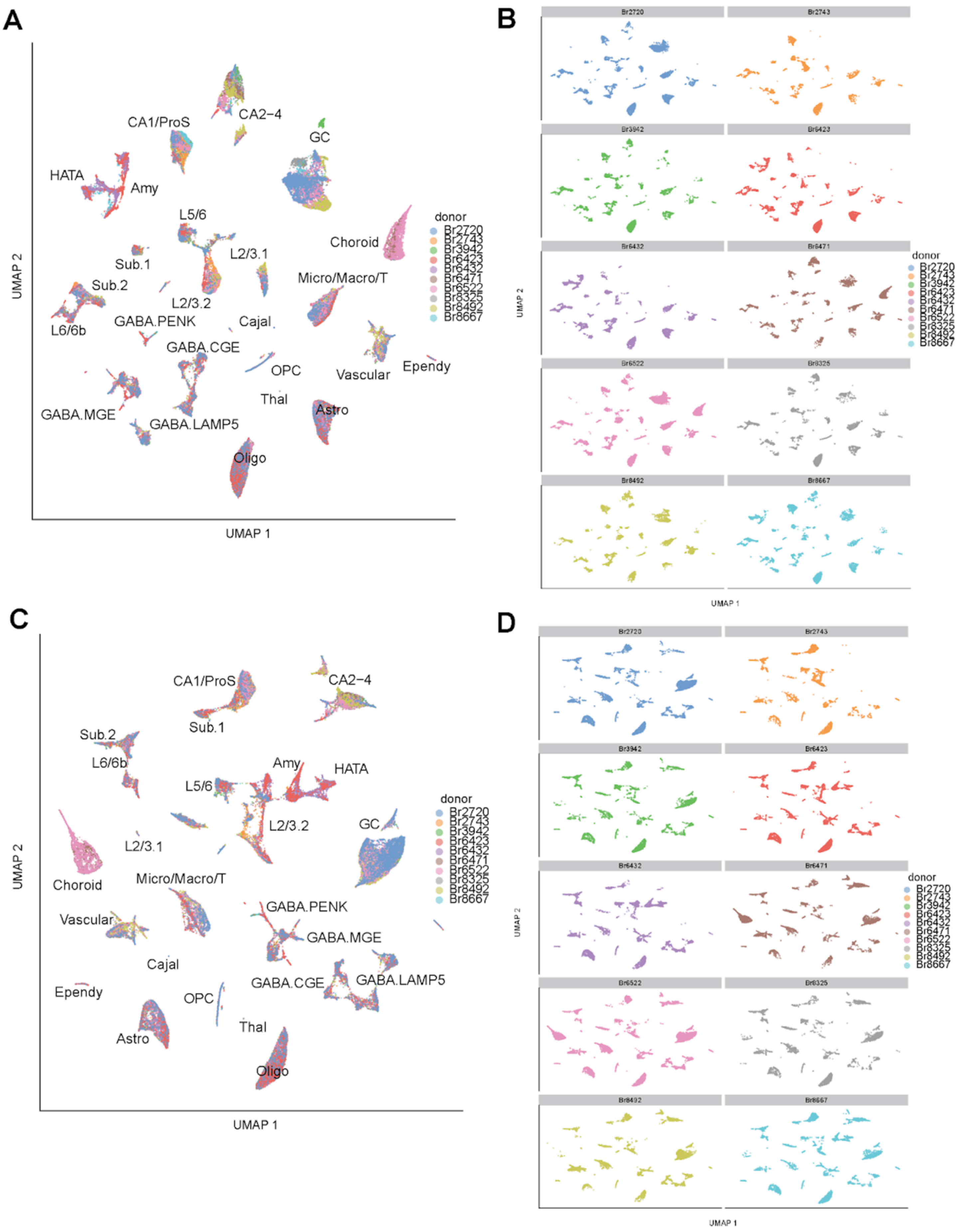
A
CA1/ProS
CA2−4
GC
HATA
Amy
L5/6
Choroid
L2/3.1
Sub.1
Sub.2
Micro/Macro/T
L2/3.2
GABA.PENK
Cajal
L6/6b
GABA.CGE
OPC
Vascular
Ependy
Thal
Astro
GABA.MGE
GABA.LAMP5
Oligo
UMAP 2
UMAP 1
donor
Br2720
Br2743
Br3942
Br6423
Br6432
Br6471
Br6522
Br8325
Br8492
Br8667
B
UMAP 2
UMAP 1
C
CA1/ProS
CA2−4
Sub.1
Sub.2
L6/6b
Amy
HATA
L5/6
L2/3.2
GC
L2/3.1
Choroid
Micro/Macro/T
GABA.PENK
Vascular
GABA.MGE
Cajal
GABA.CGE
GABA.LAMP5
Ependy
OPC
Thal
Astro
Oligo
UMAP 2
UMAP 1
D
UMAP 2
UMAP 1

**Extended Data Fig. 19. Determination of final *n*=60 clusters used for snRNA-seq dataset**

(**A**) Boxplots depict the $\log_2$ normalized counts of *SYT1* (*y*-axis) in the initial Louvain clusters (*x*-axis) clearly segregating neuronal clusters from non-neuronal clusters.

(**B**) Boxplots depict the number of detected genes (*y*-axis) in the initial Louvain neuronal clusters (*x*-axis), with boxplot color indicating three clusters that exhibited abnormally few detected genes and were therefore removed prior to re-clustering.

(**C**) A dot plot displaying the proportion of nuclei (size) and the average $\log_2$ normalized counts within those nuclei (color) of select glial marker genes (*y*-axis) across the initial non-neuronal clusters (*x*-axis). The asterisk indicates a single cluster that equally expressed markers from distinct classes of glial cells and was therefore removed prior to re-clustering. See **Supplementary Table 2** for abbreviations.

(**D**) Boxplots depict the $\log_2$ normalized counts of *SYT1* (*y*-axis) in the final Louvain clusters (*x*-axis) clearly segregating neuronal clusters from non-neuronal clusters.

(**E**) Boxplots depict the number of detected genes (*y*-axis) in the final Louvain neuronal clusters (x-axis), with boxplot color indicating one cluster that exhibited abnormally few detected genes and was therefore removed from the final dataset.

(**F**) A dot plot displaying the proportion of nuclei (size) and the average $\log_2$ normalized counts within those nuclei (color) of select glial marker genes (*y*-axis) across the final non-neuronal clusters (*x*-axis). The asterisk indicates a single cluster that equally expressed markers from distinct classes of glial cells and was therefore removed from the final dataset.

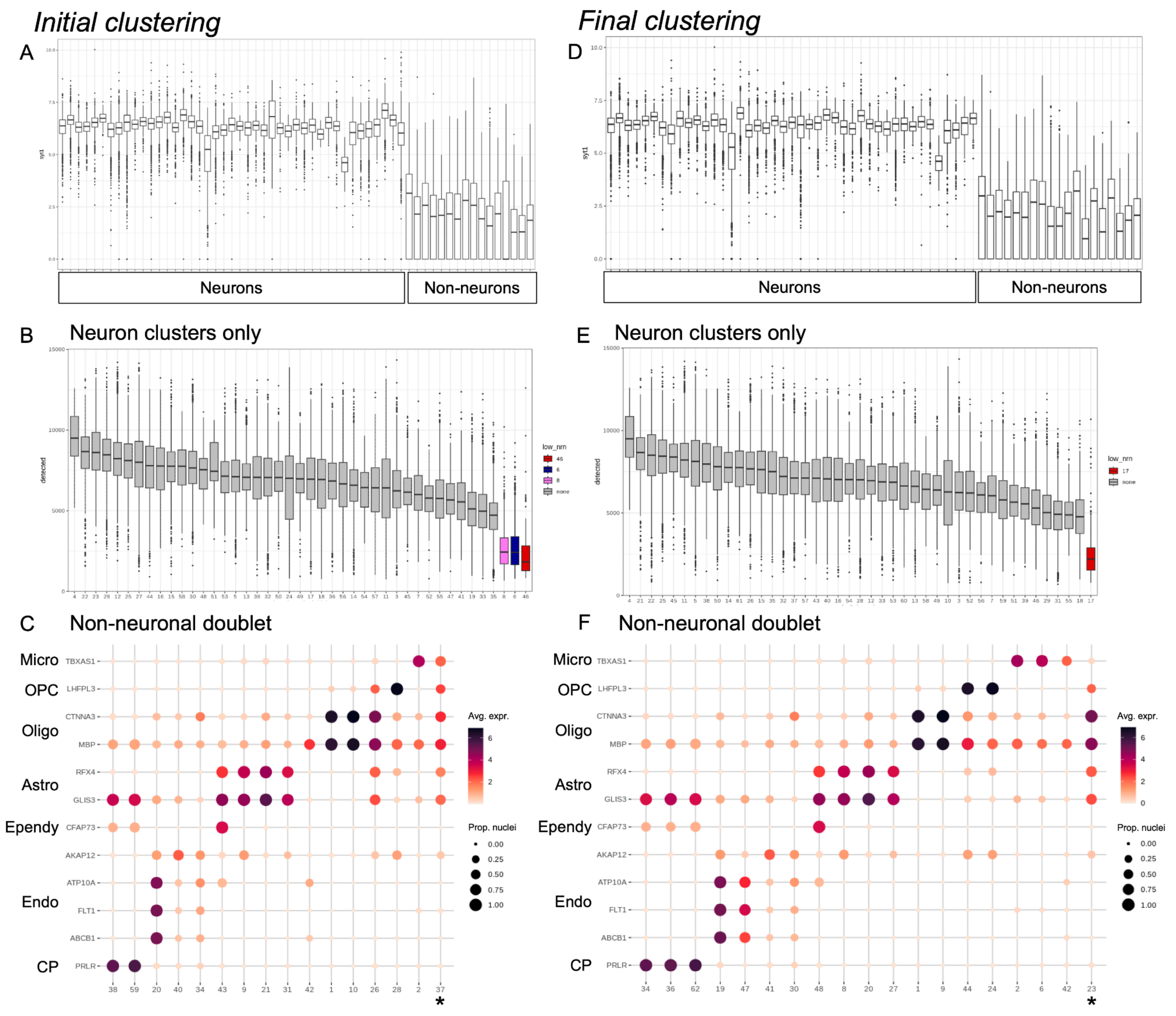
Initial clustering
Final clustering
A
D
syt1
Neurons
Non-neurons
Neurons
Non-neurons
B
Neuron clusters only
E
Neuron clusters only
detected
low_nrn
none
C
Non-neuronal doublet
F
Non-neuronal doublet
Micro
OPC
Oligo
Astro
Ependy
Endo
CP
TBXAS1
LHFPL3
CTNNA3
MBP
RFX4
GLIS3
CFAP73
AKAP12
ATP10A
FLT1
ABCB1
PRLR
Avg. expr.
Prop. nuclei
0.00
0.25
0.50
0.75
1.00
*

**Extended Data Fig. 20. Preliminary annotation of neuronal nuclei.**

(**A**) Heatmap depicting cluster average $log_2$ normalized counts (color, scaled and centered) of known marker genes for inhibitory neurons (*GAD2*, *ADARB2*, *LHX6*) and Cajal Retzius cells (*RELN*) readily distinguish several clusters from the remaining excitatory neuron clusters. Excitatory neurons clusters were identified as hippocampal/ subiculum neurons (*PROX1*, *CALB1*, *FNDC1*, *TSPAN18*, *CARTPT*, *FN1*), with most of the remaining excitatory neuron clusters likely belong to laminar structures of the greater retrohippocampal formation based on the expression of cortical layer markers *TLE4*, *SATB2*, and *CUX2*. Few clusters did not express any of these markers in abundance.

(**B**) Boxplots showing average $log_2$ normalized counts of *TCF7L2* (y-axis) clearly identifies cluster 4 as a cluster of thalamic neurons.

(**C**) Heatmap shows the average $log_2$ normalized counts (color, scaled and centered) for genes (rows) found to be differentially expressed in amygdalar SRT spots compared to retrohippocampal (RHP) spots. Five of the excitatory neuron clusters in (**A**) exhibited increased expression of the amygdala genes and decreased expression of the RHP genes. These clusters (boxed) were subsequently annotated to the amygdala.

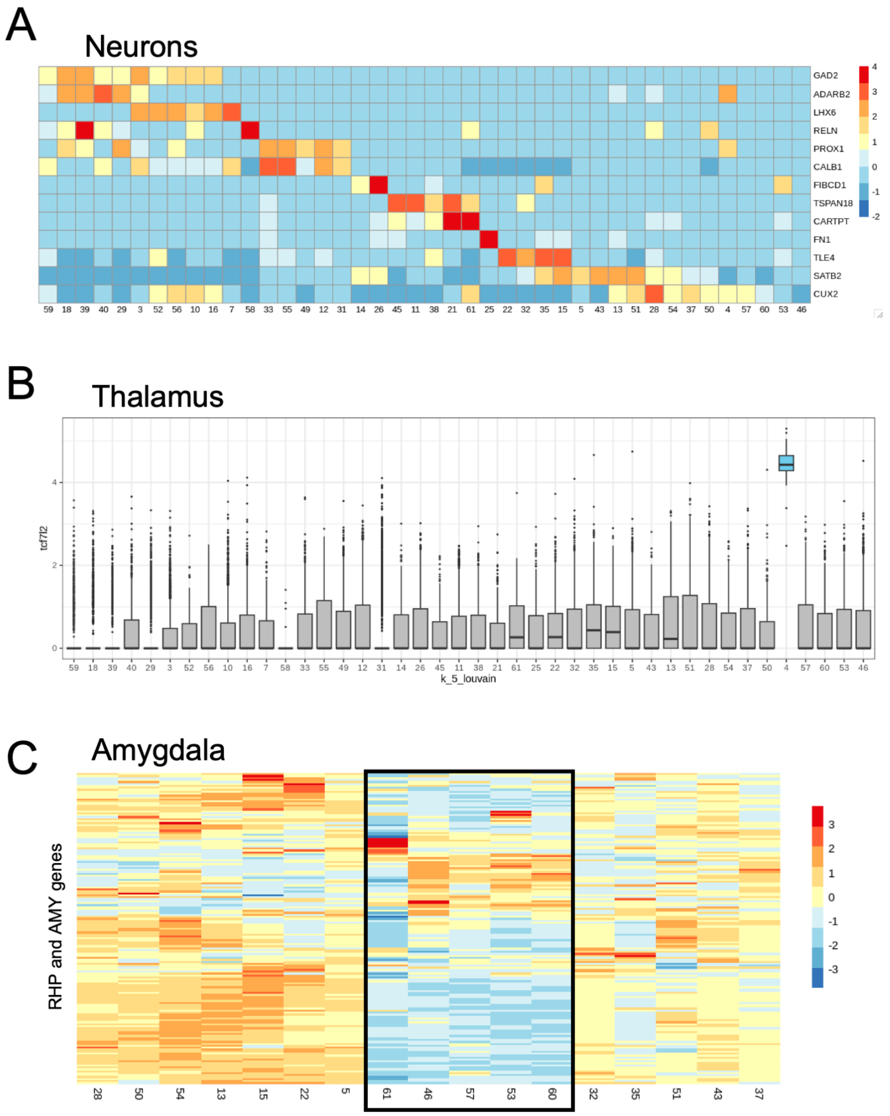

**Extended Data Fig. 21. Proportions of nuclei across fine, mid, and broad cell types.**

(**A**) Bar plots show the relative proportion of nuclei across cell types for each donor, each sorting strategy, and in total. See **Supplementary Table 2** for abbreviations. PI: propidium iodide.

(**B**) Bar plots show the relative proportion of nuclei across cell types when excitatory neuron (ExcN) and inhibitory neuron (InhN) clusters are collapsed for each donor, each sorting strategy, and in total.

(**C**) Bar plots show the relative proportion of nuclei across cell types collapsed to basic annotations for each donor, each sorting strategy, and in total.

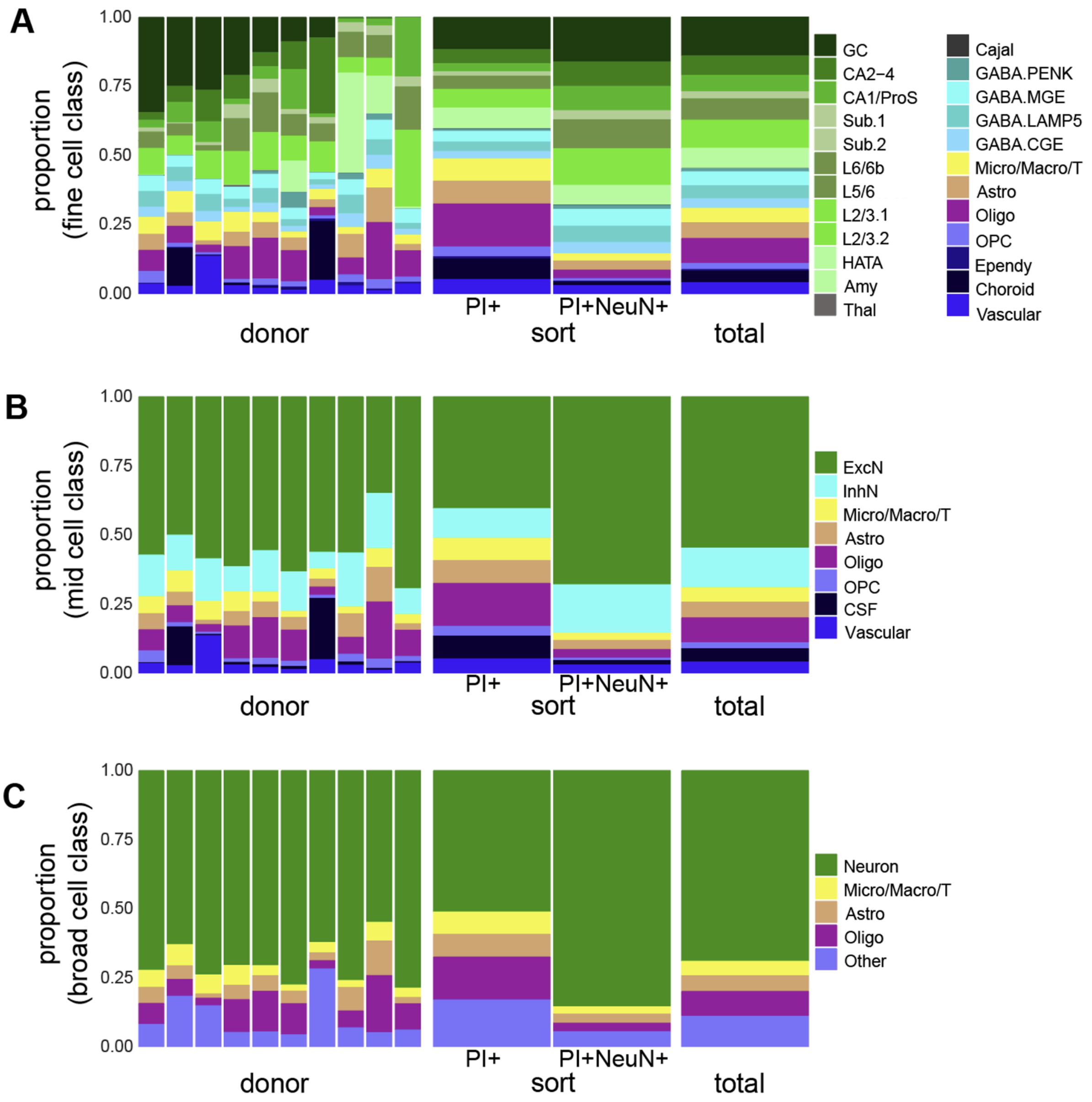

**Extended Data Fig. 22. Differentially expressed genes and the annotation of *n*=60 snRNA-seq clusters.**

Dot plots for select differentially expressed genes (*y*-axis) for *n*=60 cell types (*x*-axis). The color represents the $\log_2$ normalized counts averaged across all nuclei in that cell type and the size of the dot represents the proportion of nuclei that express those genes. Heatmaps are included for (**A**) glia, (**B**) retrohippocampus (RHP), (**C**) amygdala, (**D**) hippocampus (HPC), (**E**) GABAergic neurons, (**F**) GABAergic neuron markers with higher expression range (including Cajal Retzius neurons for *RELN* expression comparison), (**G**) and dentate gyrus granule cells. See **Supplementary Table 2** for cluster abbreviations.

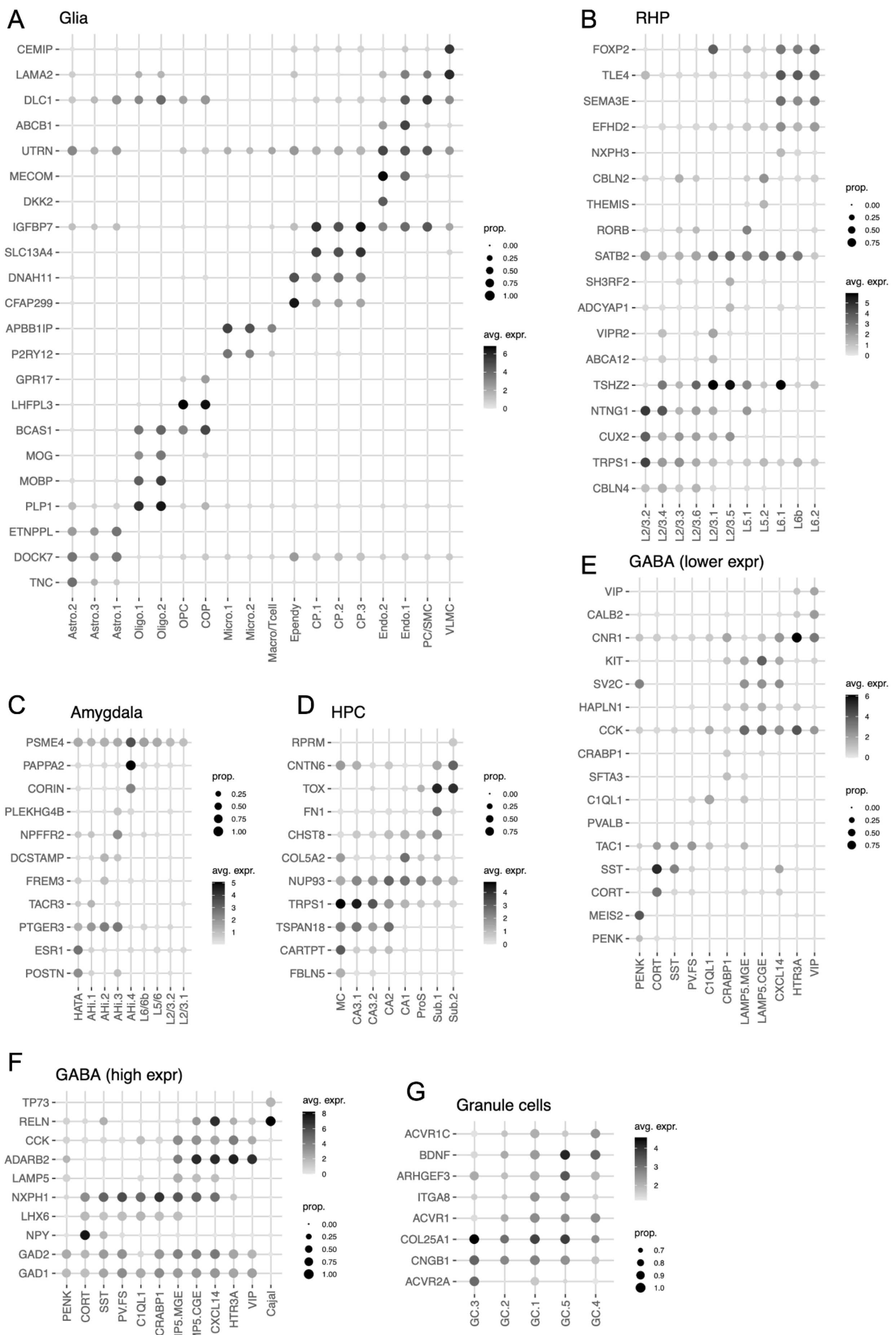
A Glia
CEMIP
LAMA2
DLC1
ABCB1
UTRN
MECOM
DKK2
IGFBP7
SLC13A4
DNAH11
CFAP299
APBB1IP
P2RY12
GPR17
LHFPL3
BCAS1
MOG
MOBP
PLP1
ETNPPL
DOCK7
TNC
Astro.2
Astro.3
Astro.1
Oligo.1
Oligo.2
OPC
COP
Micro.1
Micro.2
Macro/Tcell
Ependy
CP.1
CP.2
CP.3
Endo.2
Endo.1
PC/SMC
VLMC
prop.
0.00
0.25
0.50
0.75
1.00
avg. expr.
6
4
2
0
B RHP
FOXP2
TLE4
SEMA3E
EFHD2
NXPH3
CBLN2
THEMIS
RORB
SATB2
SH3RF2
ADCYAP1
VIPR2
ABCA12
TSHZ2
NTNG1
CUX2
TRPS1
CBLN4
L2/3.2
L2/3.4
L2/3.3
L2/3.6
L2/3.1
L2/3.5
L5.1
L5.2
L6.1
L6b
L6.2
prop.
0.00
0.25
0.50
0.75
avg. expr.
5
4
3
2
1
0
C Amygdala
PSME4
PAPPA2
CORIN
PLEKHG4B
NPFFR2
DCSTAMP
FREM3
TACR3
PTGER3
ESR1
POSTN
HATA
AHi.1
AHi.2
AHi.3
AHi.4
L6/6b
L5/6
L2/3.2
L2/3.1
prop.
0.25
0.50
0.75
1.00
avg. expr.
5
4
3
2
1
D HPC
RPRM
CNTN6
TOX
FN1
CHST8
COL5A2
NUP93
TRPS1
TSPAN18
CARTPT
FBLN5
MC
CA3.1
CA3.2
CA2
CA1
ProS
Sub.1
Sub.2
prop.
0.00
0.25
0.50
0.75
avg. expr.
4
3
2
1
0
E GABA (lower expr)
VIP
CALB2
CNR1
KIT
SV2C
HAPLN1
CCK
CRABP1
SFTA3
C1QL1
PVALB
TAC1
SST
CORT
MEIS2
PENK
PENK
CORT
SST
PV.FS
C1QL1
CRABP1
LAMP5.MGE
LAMP5.CGE
CXCL14
HTR3A
VIP
avg. expr.
6
4
2
0
prop.
0.00
0.25
0.50
0.75
F GABA (high expr)
TP73
RELN
CCK
ADARB2
LAMP5
NXPH1
LHX6
NPY
GAD2
GAD1
PENK
CORT
SST
PV.FS
C1QL1
CRABP1
LAMP5.MGE
LAMP5.CGE
CXCL14
HTR3A
VIP
Cajal
avg. expr.
8
6
4
2
0
prop.
0.00
0.25
0.50
0.75
1.00
G Granule cells
ACVR1C
BDNF
ARHGEF3
ITGA8
ACVR1
COL25A1
CNGB1
ACVR2A
GC.3
GC.2
GC.1
GC.5
GC.4
avg. expr.
4
3
2
prop.
0.7
0.8
0.9
1.0

**Extended Data Fig. 23. Correlation of snRNA-seq clusters to SRT spatial domains.**

Heatmap of Pearson correlation coefficients (color) showing the relationship between pseudobulked differential expression analysis enrichment model *t*-statistics for the SRT spatial domains (rows) and those of *n*=60 snRNAseq clusters (columns). See **Supplementary Table 2** for cluster abbreviations.

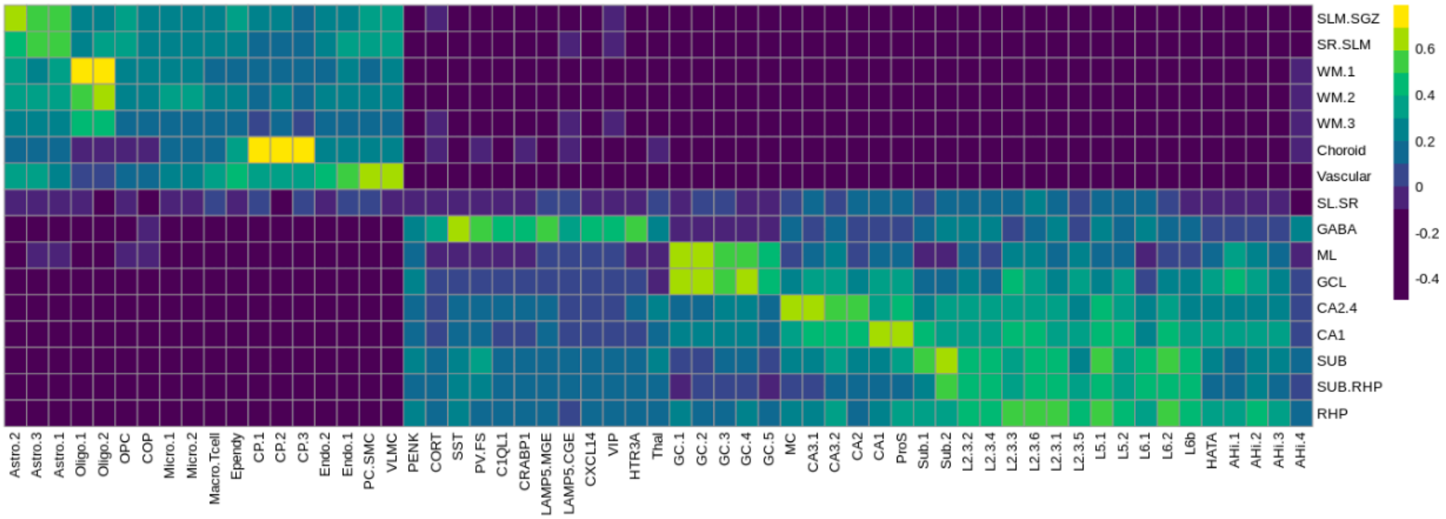

**Extended Data Fig. 24. Preliminary annotation of neuronal nuclei in validation re-analysis.**

(**A**) Violin plots of nuclei mitochondrial fraction, separated by the sequencing round. An initial mitochondrial fraction threshold of 20% indicates that a stricter limit of 10% is suitable for (**B**) propidium iodide (PI)+ samples and (**C**) PI+NeuN+ samples.

(**D**) Nuclei from PI+ and PI+NeuN+ samples exhibit distinct distributions of the number of detected genes.

(**E**) Setting the minimum number of detected genes to 750 is a reasonable limit across all sequencing rounds for PI+ samples.

(**F**) For PI+NeuN+ samples, sequencing round 2 has an abnormally high number of nuclei with few detected genes. This effect is driven by one sample (17c-scp).

(**G**) After removing the errant round 2 sample, the distribution of PI+NeuN+ nuclei based on the number of detected genes is similar across sequencing rounds and indicates keeping nuclei with a minimum of 1000 detected genes.

(**H**) Nuclei from PI+ and PI+NeuN+ samples exhibit distinct distributions of the number of reads per nucleus.

(**I**) Setting the minimum number of reads to 2000 is a reasonable limit across all sequencing rounds for PI+ samples.

(**J**) For PI+NeuN+ samples, sequencing round 2 has an abnormally high number of nuclei with few reads. This effect is driven by one sample (17c-scp).

(**K**) After removing the errant round 2 sample (17c-scp), the distribution of PI+NeuN+ nuclei based on the number of reads is similar across sequencing rounds and indicates keeping nuclei with a minimum of 3500 reads.

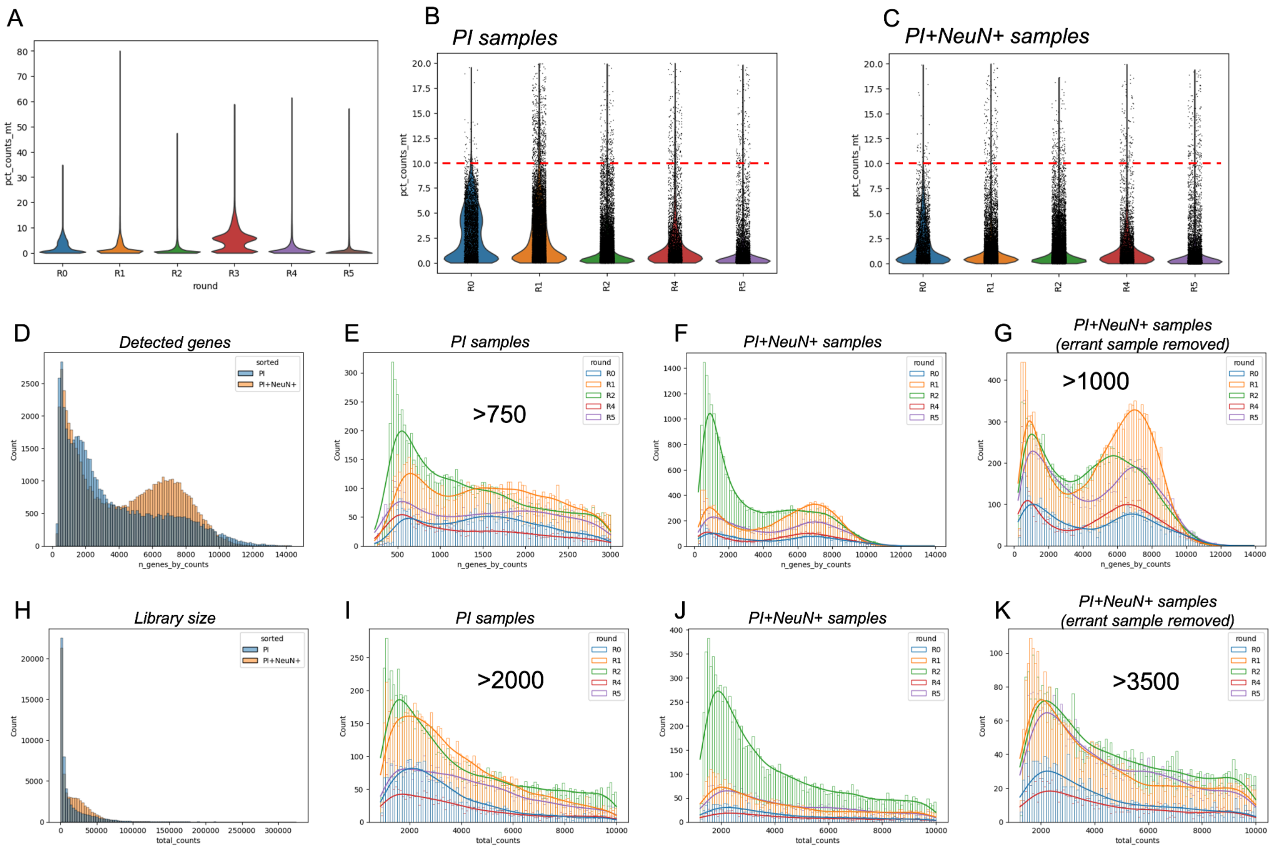
A
pct_counts_mt
round
R0
R1
R2
R3
R4
R5
B
PI samples
C
PI+NeuN+ samples
D
Detected genes
sorted
PI
PI+NeuN+
n_genes_by_counts
Count
E
PI samples
>750
F
PI+NeuN+ samples
G
PI+NeuN+ samples
(errant sample removed)
>1000
H
Library size
total_counts
I
PI samples
>2000
J
PI+NeuN+ samples
K
PI+NeuN+ samples
(errant sample removed)
>3500

**Extended Data Fig. 25. Clustering approach adopted in validation re-analysis.**

(**A**) Uniform manifold approximation (UMAP) plot of initial Leiden clustering results based on `scVI` representations, annotated with cluster number.

(**B**) UMAP plot of initial Leiden clustering results based on `scVI` representations, colored based on the percent of reads originating from the mitochondrial genome (chrM) (also known as mitochondrial fraction).

(**C**) UMAP plot of initial Leiden clustering results based on `scVI` representations, colored based on the $\log_2$ normalized counts of *MALAT1*, a gene used to indicate nuclei quality.

(**D**) Violin plot depicts mitochondrial fraction (*y*-axis) across initial Leiden clusters (*x*-axis). Asterisks indicate clusters that were removed prior to re-clustering due to high mitochondrial fraction.

(**E**) Violin plot depicts *MALAT1* logcount expression (*y*-axis) across initial Leiden clusters (*x*-axis). Asterisks indicate clusters that were removed prior to re-clustering due to low *MALAT1* expression.

(**F**) Cluster 8 exhibited borderline exclusionary mitochondrial fraction that seemed to be due to nuclei from a single donor. This violin plot of mitochondrial fraction (*y*-axis) across all clusters from this sample only (*x*-axis) demonstrates that the increased mitochondrial fraction of cluster 8 is abnormal amongst this sample's nuclei. Nuclei belonging to the cluster labeled with an asterisk were removed prior to re-clustering.

(**G**) UMAP plot after re-clustering with *n*=10 neighbors following removal of low quality initial clusters. Color and number indicate final cluster annotations.

(**H**) UMAP plot demonstrating improved nuclei quality based on mitochondrial fraction (color) after removal of low quality initial clusters.

(**I**) The 74,216 nuclei obtained from the python-based re-analysis, with the 64,251 nuclei jointly present in the manuscript labeled in orange and the 9,965 nuclei unique to the re-analysis labeled in blue.

(**J**) Heatmap depicts the correlation coefficient (color) of the *t*-statistics generated from pseudobulked cluster enrichment differential expression analysis in the re-analyzed clusters (rows) and the original *n*=60 clusters (columns). See **Supplementary Table 2** for cluster abbreviations.

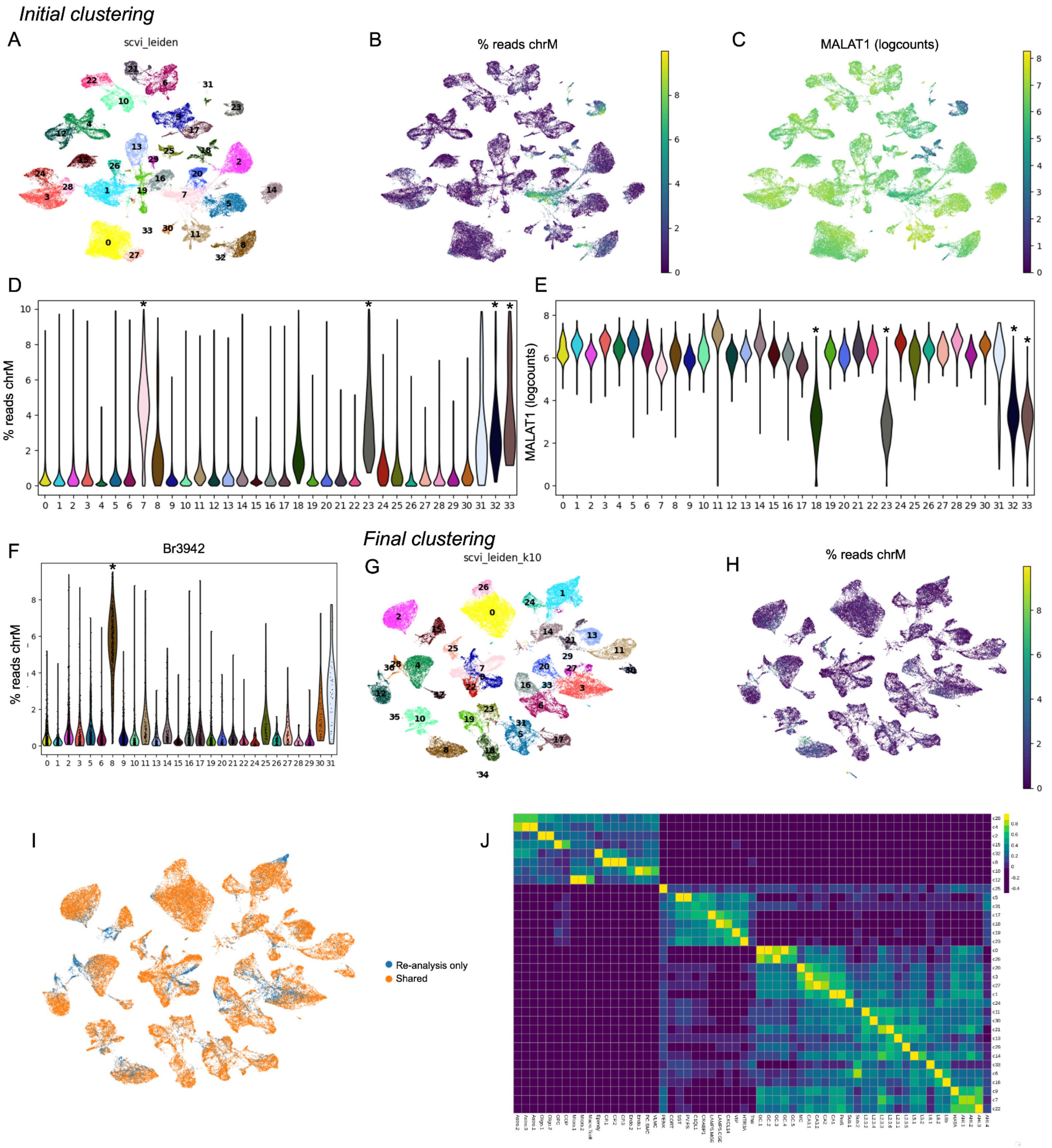
Initial clustering
A
scvi_leiden
B
% reads chrM
C
MALAT1 (logcounts)
D
% reads chrM
E
MALAT1 (logcounts)
F
Br3942
% reads chrM
Final clustering
G
scvi_leiden_k10
H
% reads chrM
I
Re-analysis only
Shared
J

**Extended Data Fig. 26. Linkage disequilibrium score regression identifies differential disease risk heritability across cell types and spatial domains.**

(**A**) Plot showing linkage disequilibrium score regression (LDSC) coefficient *z*-scores for heritability of different polygenic traits (*y*-axis) across snRNA-seq cell types (*x*-axis). Coefficients found to be significantly different from the baseline model (43) following multiple testing correction (FDR < 0.05) are shown. Circles are colored by LDSC regression coefficient *z*-score and sized by -$\log_{10}$ FDR-adjusted *p*-value. Positive coefficient *z*-scores denote a positive contribution of a given cluster (*x*-axis) to trait heritability. Negative coefficient *z*-score denotes depletion of trait heritability within a given cluster (*x*-axis). See **Supplementary Table 2** for abbreviations.

(**B**) Same as A, but for SRT spatial domains (*x*-axis). See **Supplementary Table 2** for abbreviations.

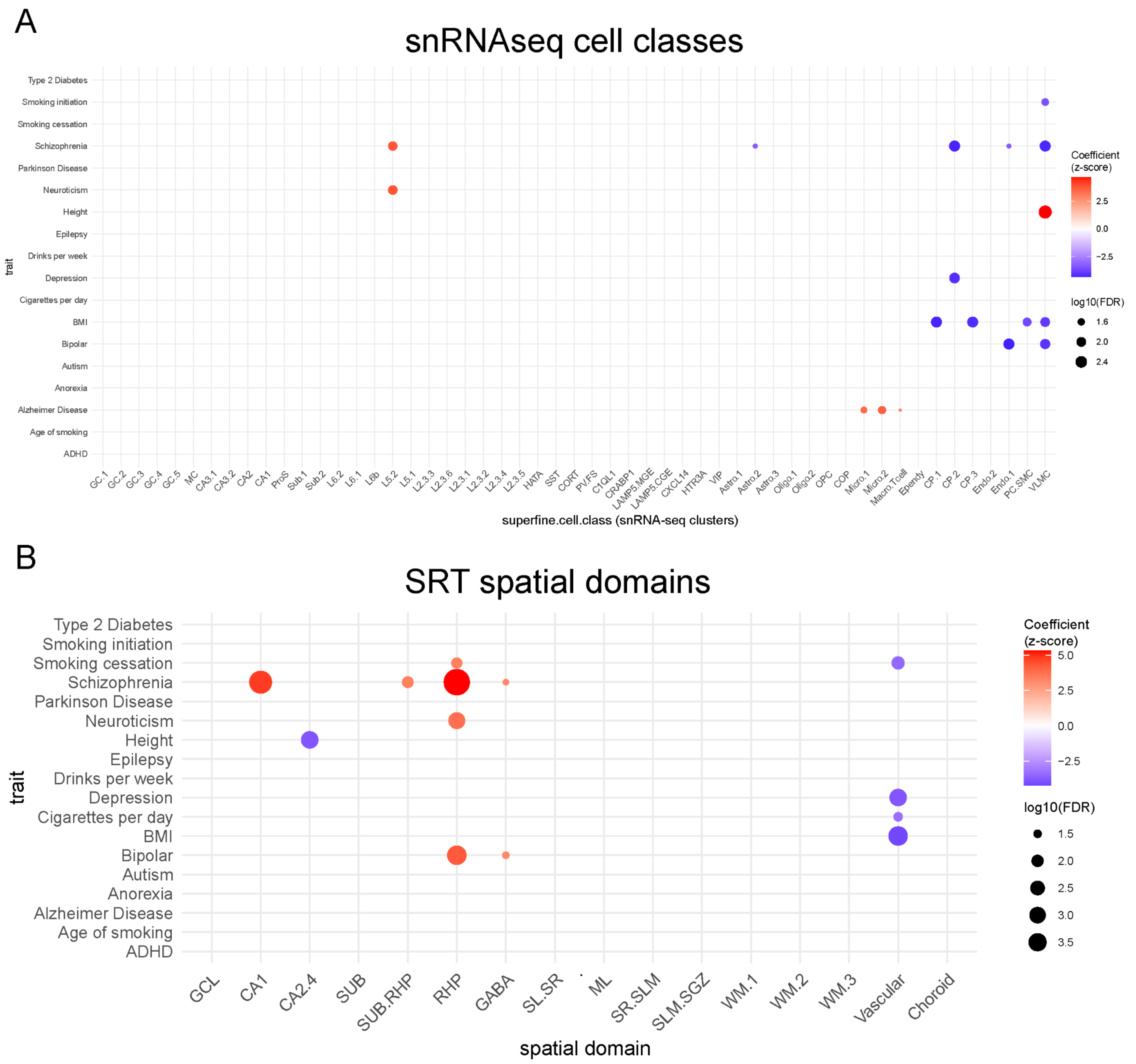

**Extended Data Fig. 27**. **Visium Spatial Proteogenomics (Visium-SPG) quality control and spatial domain label transfer.**

(**A**) Left: Merged immunofluorescence (IF) images from Visium Spatial Proteogenomics (SPG) data from donors Br3942 and Br8325. IF images represented DAPI (blue, all nuclei), NeuN (green, neuronal nuclei), TMEM119 (yellow, microglia), GFAP (red, astrocytes), and OLIG2 (magenta, oligodendrocytes). For each donor, spot plots of $\log_2$ transformed normalized counts for (middle) *SNAP25* and (right) *MBP*.

(**B**) Boxplots of library size (*y*-axis, total number of reads in a single spot) based on three median absolute deviation (3MAD) threshold calculated for each capture area independently. The spots are grouped by donor (*x*-axis) and are stratified by whether the spots were kept (black) or removed (red).

(**C**) Boxplots of number of detected genes (*y*-axis, total number of genes with at least one read in a single spot) based on three median absolute deviation (3MAD) threshold calculated for each capture area independently. The spots are grouped by donor (*x*-axis) and are stratified by whether the spots were kept (black) or removed (red).

(**D**) Bar plots demonstrate the number of spots (*y*-axis, log scale) per sample (individual bar) for each donor (*x*-axis). The plot is faceted by whether the spots were removed based on QC filter (top) or kept in the final dataset (bottom).

(**E**) Spot plot visualizing spots excluded for each Visium-SPG sample.

(**F**) Spot plots of Visium-SPG captures areas colored by spatial domain. Spatial domain labels annotated in the SRT dataset were projected to Visium-SPG dataset using `RcppML` (**Methods 4.8**). See **Supplementary Table 2** for abbreviations.

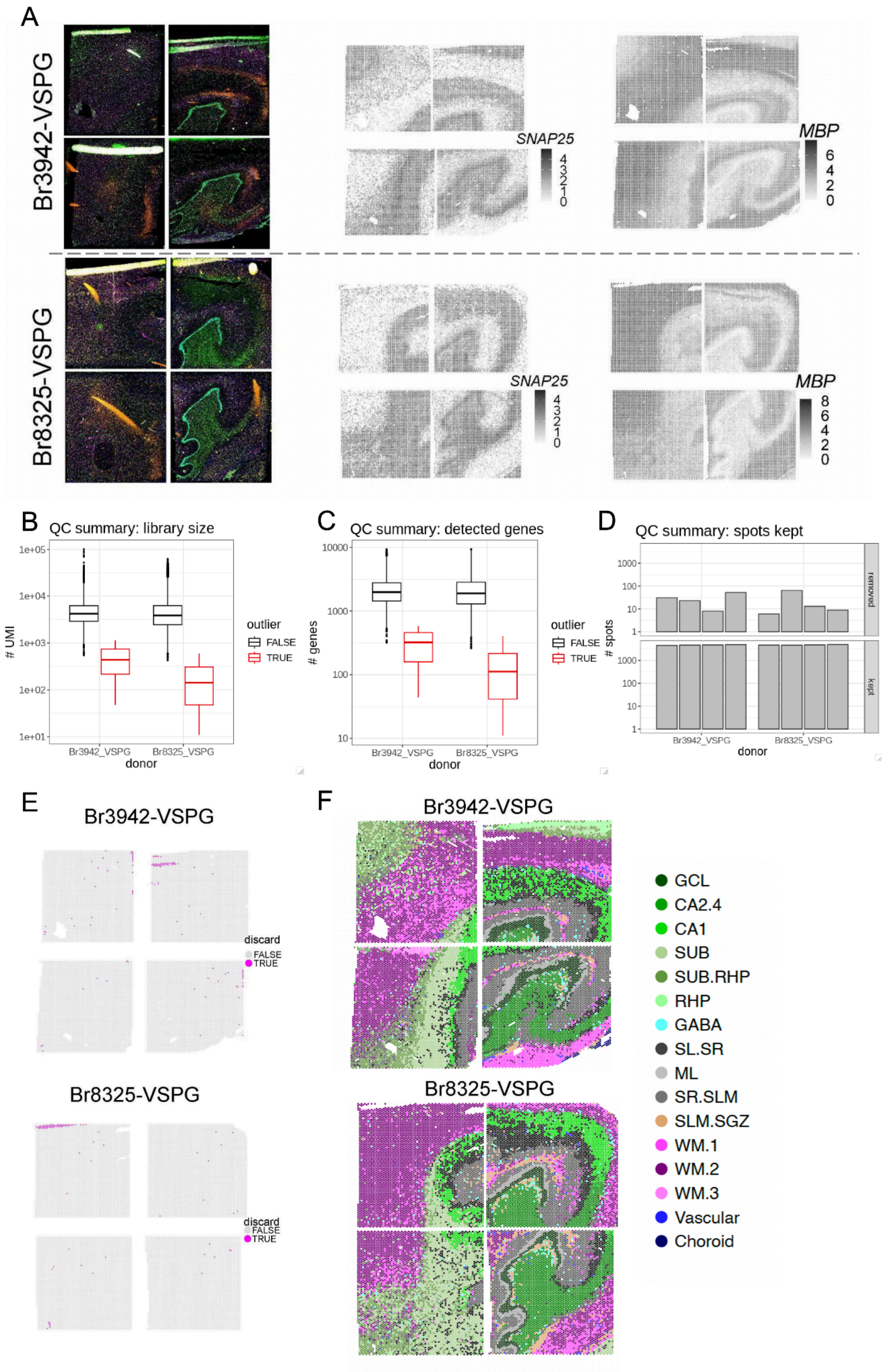

A
Br3942-VSPG
Br8325-VSPG
SNAP25
MBP
B
QC summary: library size
### UMI
donor
Br3942_VSPG
Br8325_VSPG
outlier
FALSE
TRUE
C
QC summary: detected genes
### genes
D
QC summary: spots kept
### spots
removed
kept
E
discard
F
GCL
CA2.4
CA1
SUB
SUB.RHP
RHP
GABA
SL.SR
ML
SR.SLM
SLM.SGZ
WM.1
WM.2
WM.3
Vascular
Choroid

**Extended Data Fig. 28. Hierarchy of cell type labels across superfine, fine, mid, and broad classifications.**

Dendrogram shows the hierarchical classification of cell type labels across cluster resolutions utilized for deconvolution. See **Supplementary Table 2** for abbreviations.

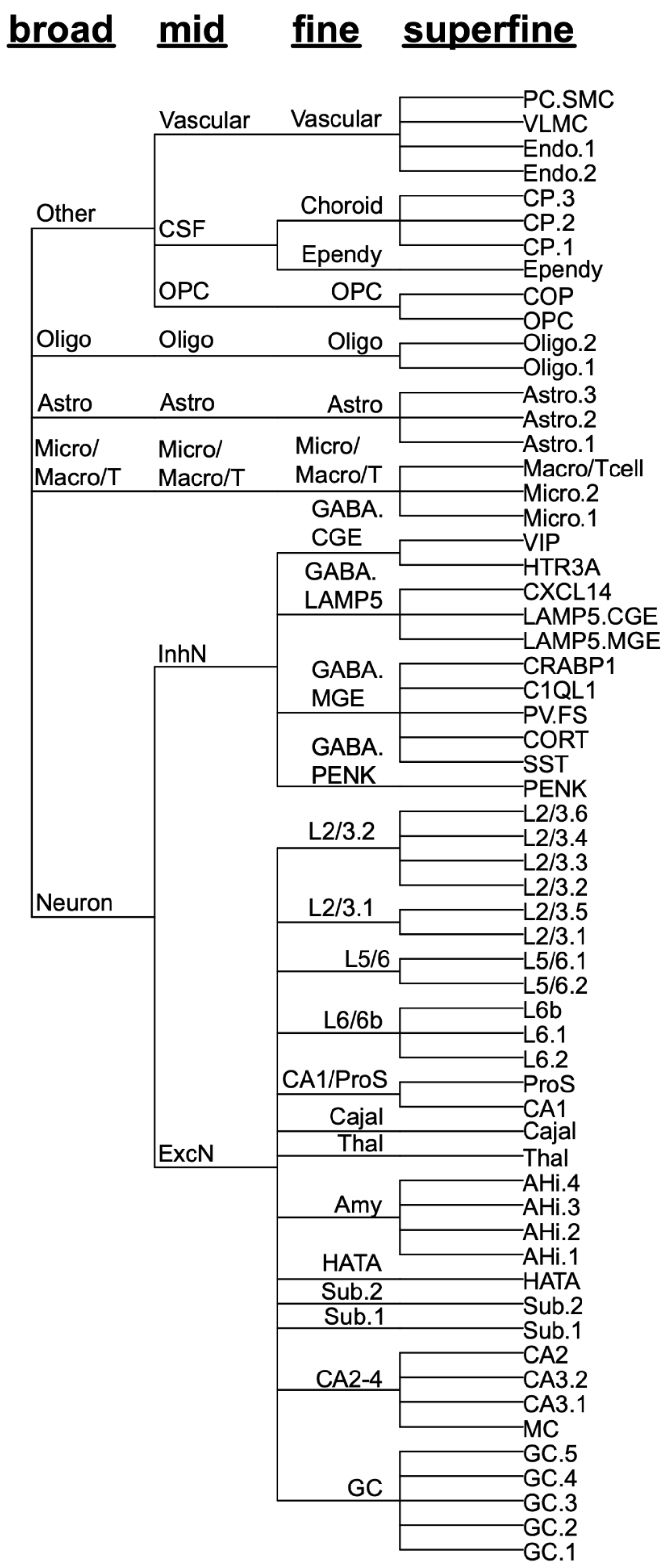

**Extended Data Fig. 29. Schematic for deconvolution workflow.**

(**A**) Schematic showing the complexity of cell types in Visium spots.

(**B**) Marker gene detection strategy, for mid and fine level cell types. See **Supplementary Table 2** for abbreviations.

(**C**) Performance of spot-level deconvolution algorithms was benchmarked by comparison to orthogonal assignment of cell type identity based on Classification And Regression Tree (CART) results from Visium Spatial Proteogenomics (Visium-SPG) data (top). Scatterpie plots illustrate the performance of deconvolution tools (from top: `RCTD`, `cell2location`, `Tangram`). Inset regions in the GCL and WM are denoted by white squares.

(**D**) `RCTD` was applied to the Visium-H&E data, where the orthogonal cell type information is not present. Stacked bar plots illustrate the proportion of each cell type within each spatial domain.

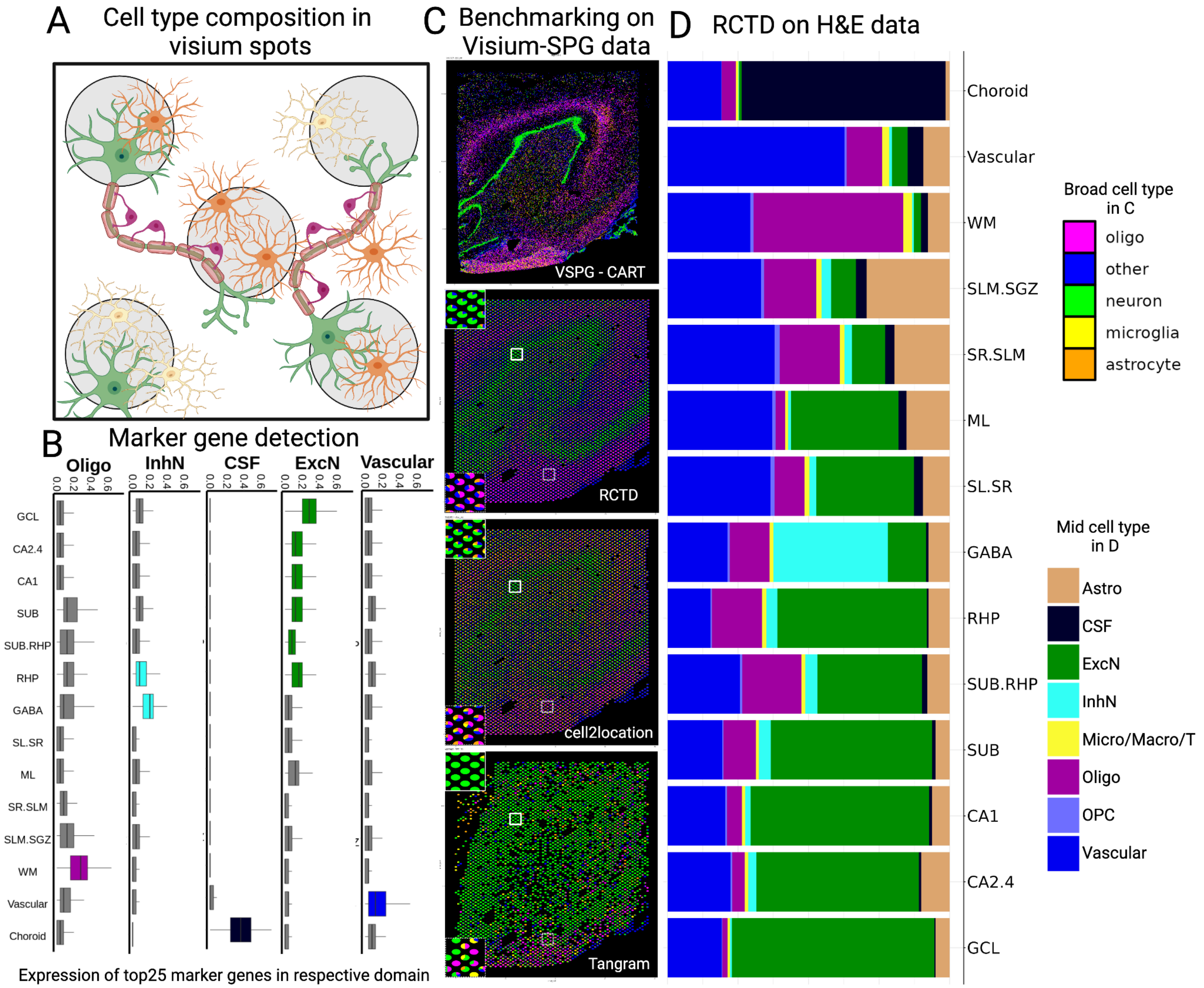

**Extended Data Fig. 30. Decision-tree framework for classification and regression tree (`CART`) - cell types using immunofluorescence (IF) intensities.**

(**A**) For each randomly chosen segmented nucleus, shown in the leftmost column (cell mask), fluorescence intensity for the DAPI channel and other channels marking each cell type is shown in the remaining columns. The decision tree uses the mean intensity values within the cell mask from each channel for classification.

(**B**) The decision-tree structure, which is traversed downwards. Each colored box represents a decision, made by comparing the mean fluorescence of a single channel for a given segmented cell against a learned threshold (indicated within each box). The red and green arrows indicate the direction of decision depending on the threshold indicated in the box, if mean intensity is below threshold tree follows red arrow otherwise it follows a green arrow. For example, for a cell to be classified as microglia, the mean pixel intensity of its cell mask should be less than 15 in the NeuN channel, less than 21 in the OLIG2 channel, less than 16 in the GFAP channel and exceeding 17 in the TMEM119 channel.

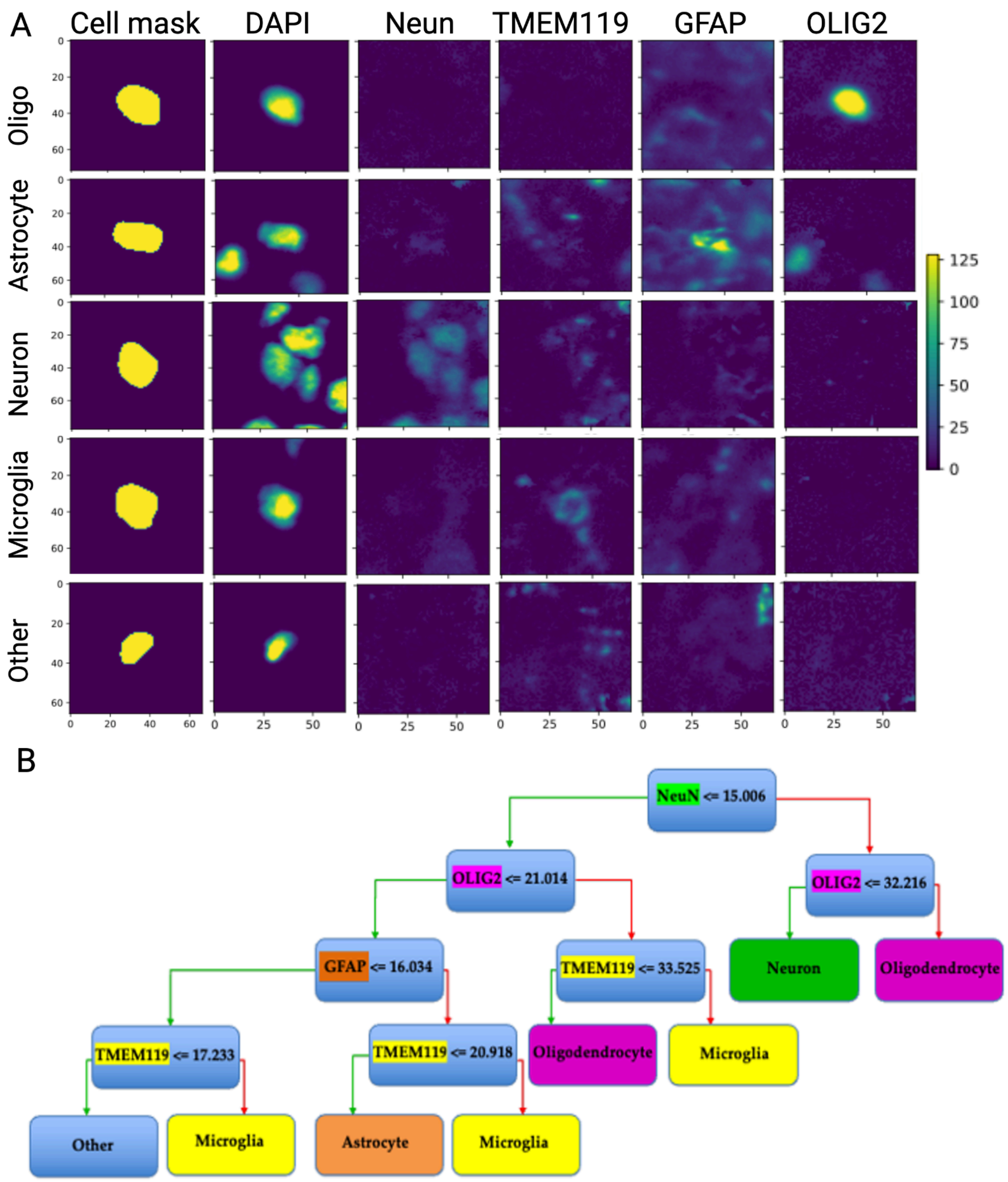

**Extended Data Fig. 31. Distribution of standard log-fold change vs. mean ratio of all genes for fine cell types.**

For each cell type, blue points are the 25 genes with the largest Mean Ratios (172), which were chosen as marker genes and used as input into spot-deconvolution methods. Red points are the remaining genes, which are not considered cell type markers and therefore not used in downstream analyses. See **Supplementary Table 2** for abbreviations.

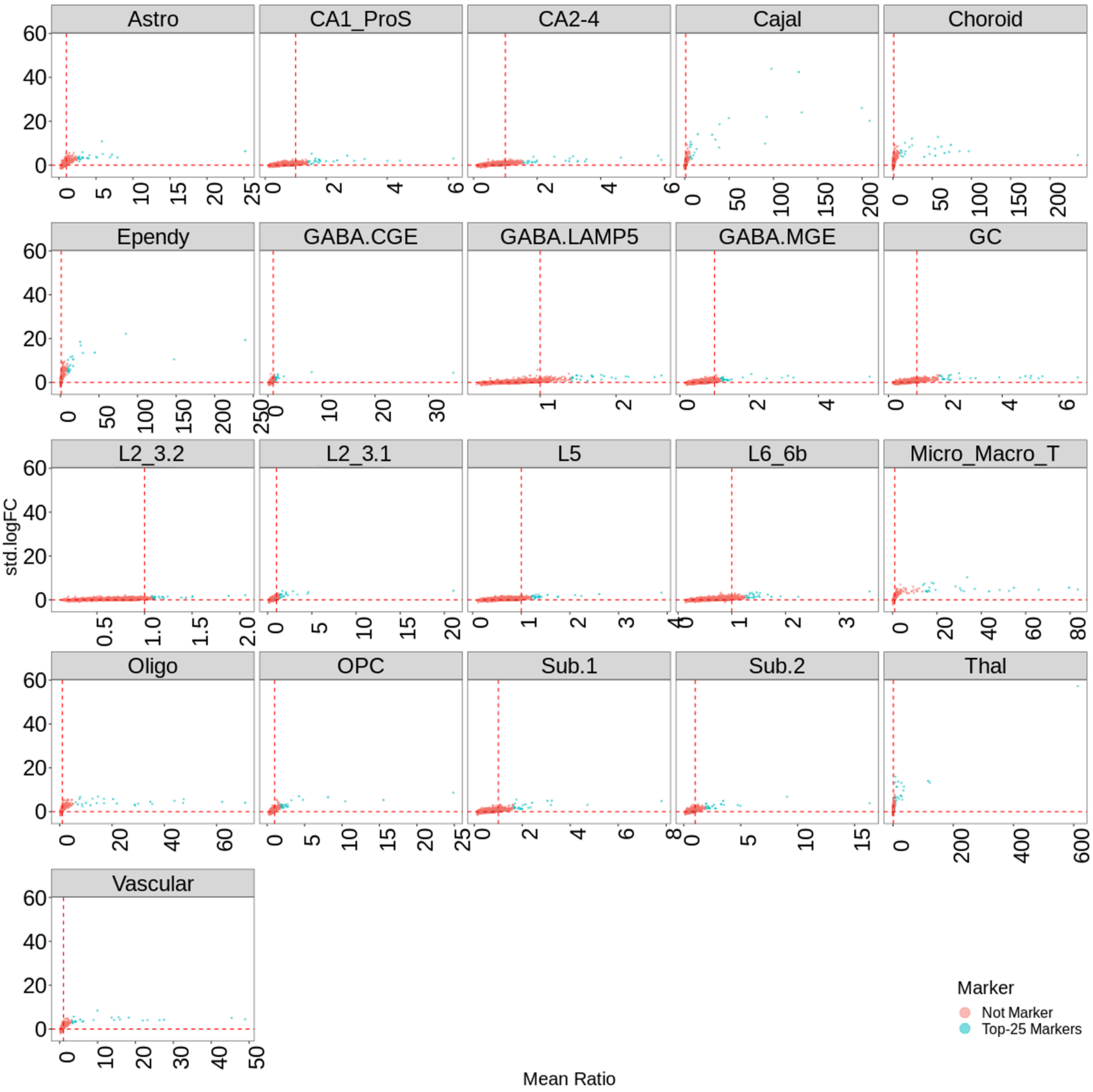

**Extended Data Fig. 32. Determining the optimal number of gene markers for spot deconvolution.** Comparison of *PPFIA2* counts to GC excitatory neuron marker gene expression with increasing number of marker genes. The top row shows spot plots for raw counts of *PPFIA2*, a marker for GCL, for all four Visium-SPG sections from donor Br3942. Remaining rows show spot plots depicting the proportion of GC marker genes with non-zero counts at each spot, when selecting the top 15, 25, or 50 marker genes. The choice of 25 marker genes per cell type results in an informative signal without significant expression outside of the expected layer.

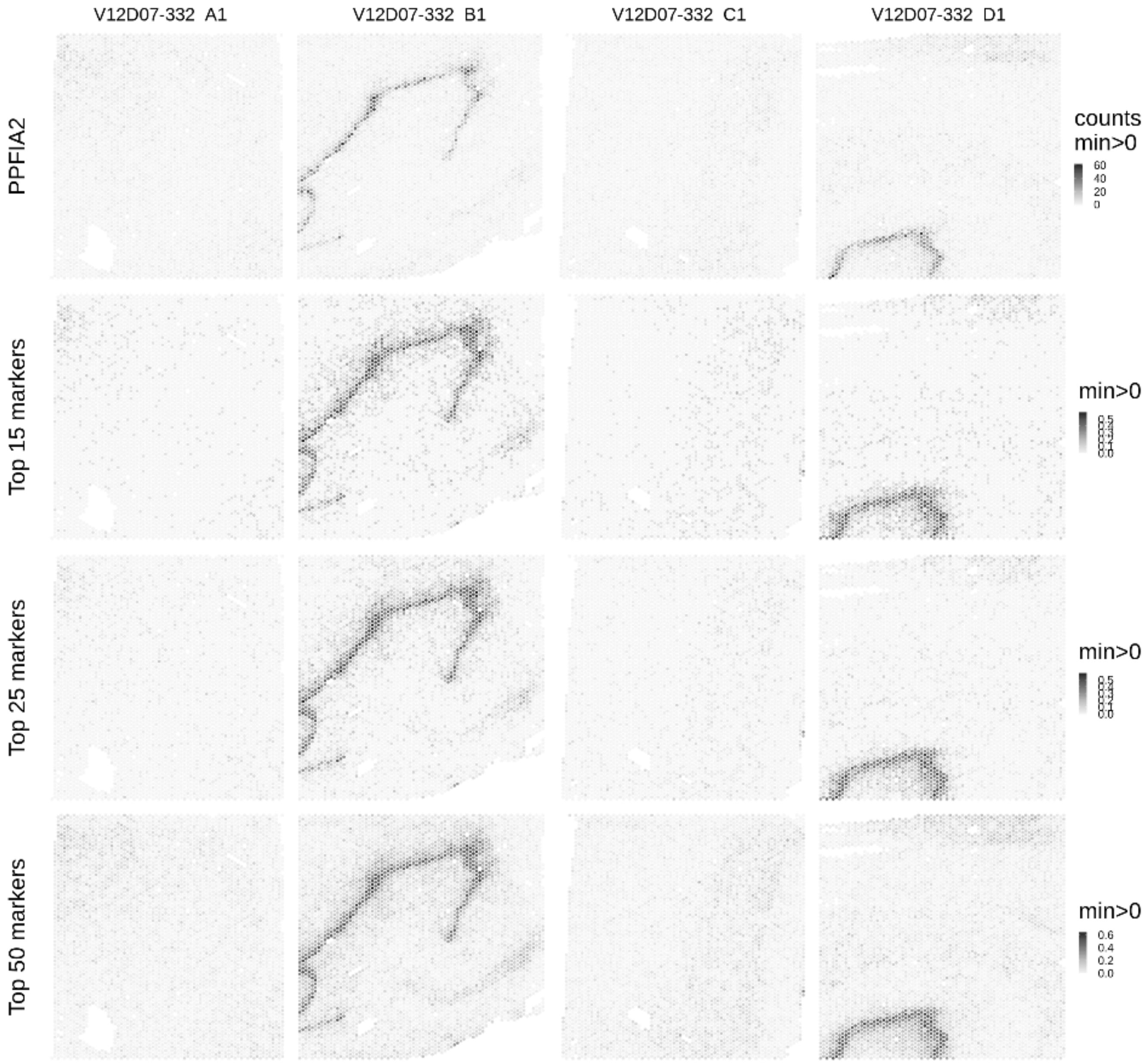

**Extended Data Fig. 33. Spatial validation of fine cell type marker gene (top 25) expression.**
The boxplots show the expression of top 25 marker genes (in a given spot, the proportion of the top 25 marker genes for each cell type having at least one count was computed) at fine level cell type, across the projected spatial domains for the 8 Visium-SPG sections. This analysis was performed to validate the spatial integrity of marker genes. For example, as expected, the largest proportion of oligodendrocyte marker genes is in white matter (WM) domain. See **Supplementary Table 2** for abbreviations.

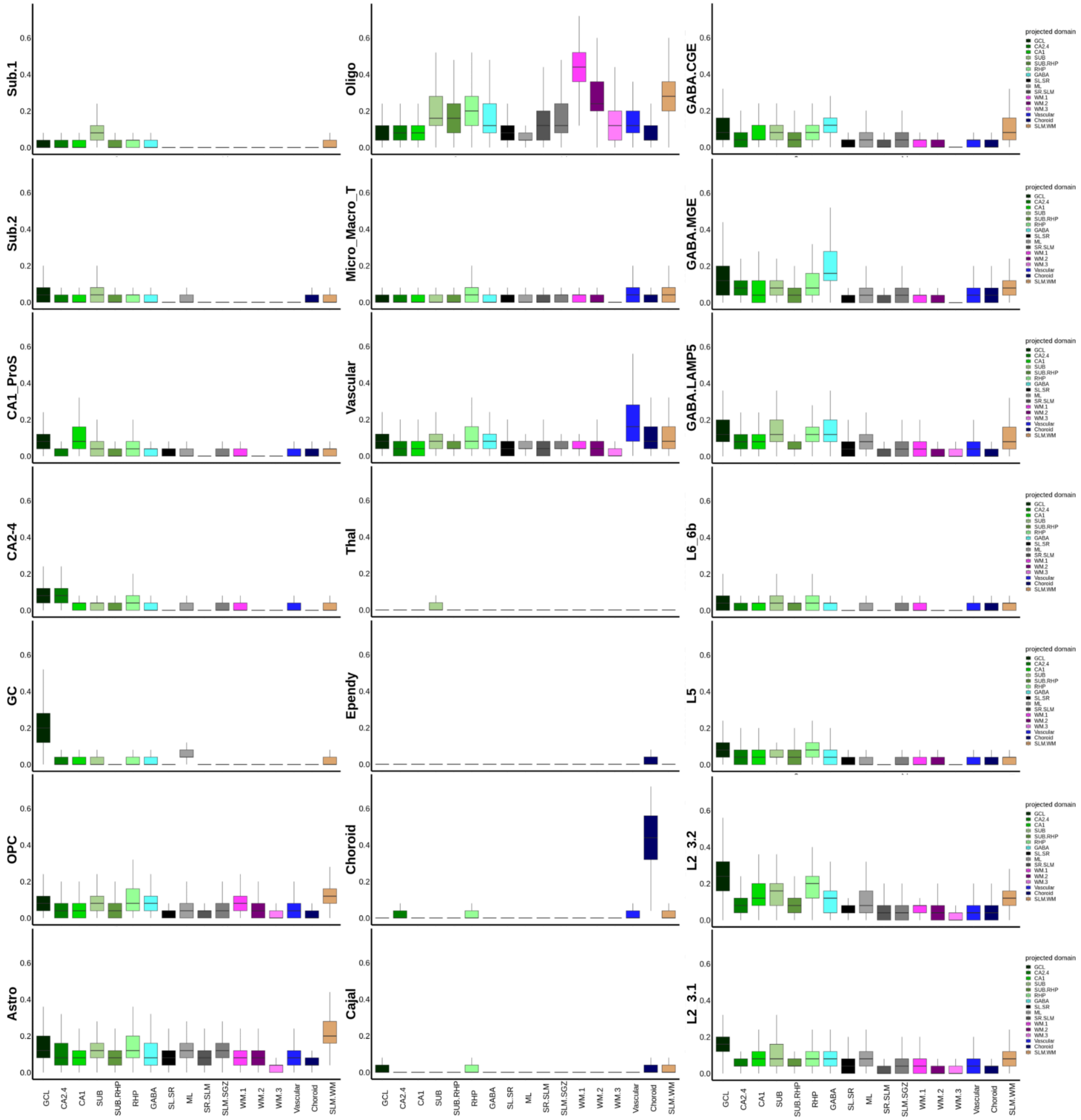

**Extended Data Fig. 34. `RCTD` performance with user derived marker genes vs default marker genes.**

(**A**) Bar plots on left show the number spots dropped by `RCTD` when using user-derived marker genes from `DeconvoBuddies` compared with other tools (`cell2location` and `Tangram`). Bar plots on the left show that `RCTD` does not drop additional spots when using `RCTD` derived marker genes.

(**B**) `RCTD` predictions for mid cell type proportions with the user derived marker genes from `DeconvoBuddies` package (172). Red spots indicate the `RCTD` dropped spots when using user defined marker genes. See **Supplementary Table 2** for abbreviations.

(**C**) `RCTD` predictions for mid cell type proportions with default marker genes.

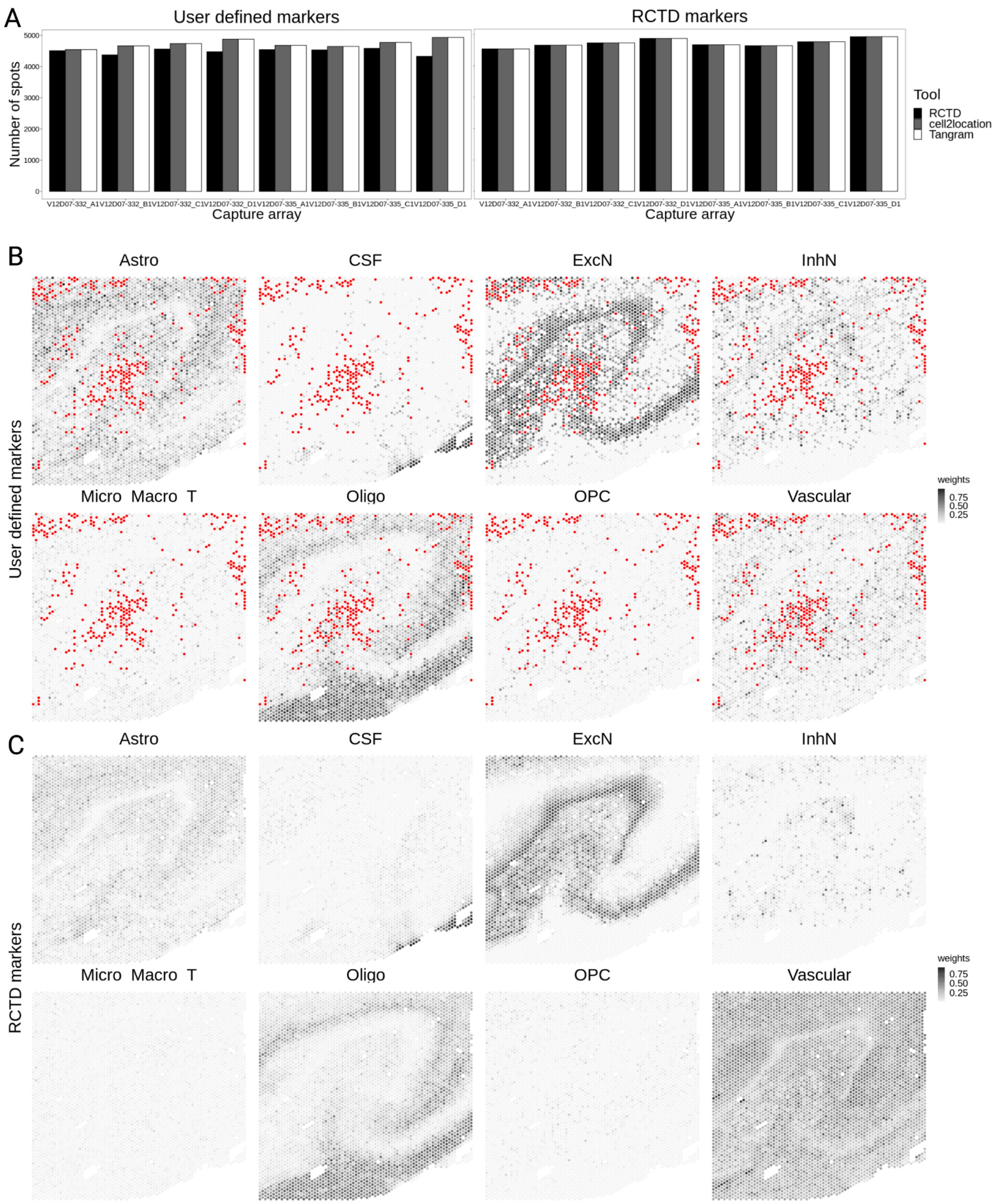

**Extended Data Fig. 35. Spatial comparison of broad cell type proportions in classification and Regression Tree (`CART`) vs. spot deconvolution tool.**

The top row shows spot plots of broad cell type counts from `CART`-predictions, for one Visium-SPG capture area V12D07-332_D1 from donor Br3942. The bottom three rows show collapsed mid cell type counts as estimated by `cell2location`, `RCTD` and `Tangram`.

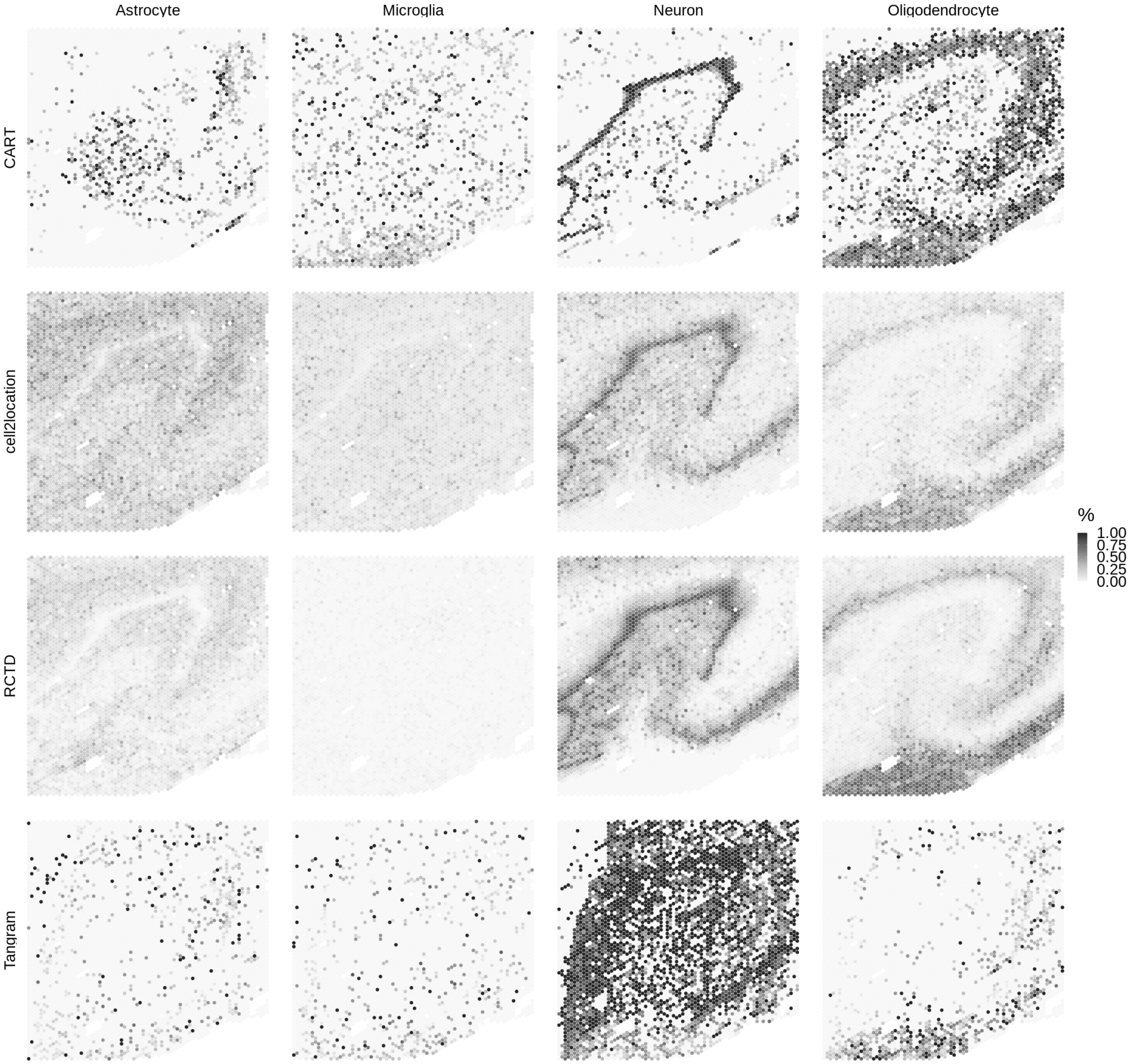

**Extended Data Fig. 36. Benchmarking of spot deconvolution results.**

(**A**) Comparison of cell type counts from mid and fine cell type classifications (both collapsed into broad cell types) from all three deconvolution tools.

(**B**) (i) Classification and Regression Tree (`CART`)-calculated vs. deconvolution-estimated counts for four immunolabeled cell types (Astro, Micro, Neuron, Oligo) across eight Visium-SPG sections. Each point represents the number of cells of the given cell type present in a tissue section, both as estimated by each deconvolution method (*x*-axis, facets) and as determined by the decision-tree classifier (*y*-axis). Since the CART framework can only predict the four cell types indicated by the immunofluorescence channels present in the Visium-SPG data, the deconvolution results that were predicted at mid cell type resolution are here collapsed down to broad cell types. Kulback-Leibler divergence (KL Div), Pearson correlation (Cor) and root mean squared error (RMSE) are the metrics computed to summarize the relationship between `CART`-calculated and deconvolution-estimated counts for a particular software method across four cell types and eight tissue sections. (ii) Predicted counts for a given method (facets), section (points), and mid cell type classification collapsed to broad (colors) are compared against the corresponding `CART` predictions by computing the Pearson correlation (*x*-axis) and RMSE (*y*-axis). Each of these values is then averaged to generate a single correlation (Avg. Cor) and RMSE (Avg. RMSE) value for each method (inset text). (iii) The predicted proportion (*y*-axis) of the broad cell types (fill color) by the four prediction methods (*x*-axis) for each sample (facets). Totals do not add to 1 because `CART` can classify cells as “other” (negative for Astro, Micro, Neuron, and Oligo), and the Vascular, OPC, CSF cell type predictions from the 3 softwares are grouped as others.

(**C**) The same metrics and plots from (**B**) were repeated for the fine cell types collapsed to broad cell type.

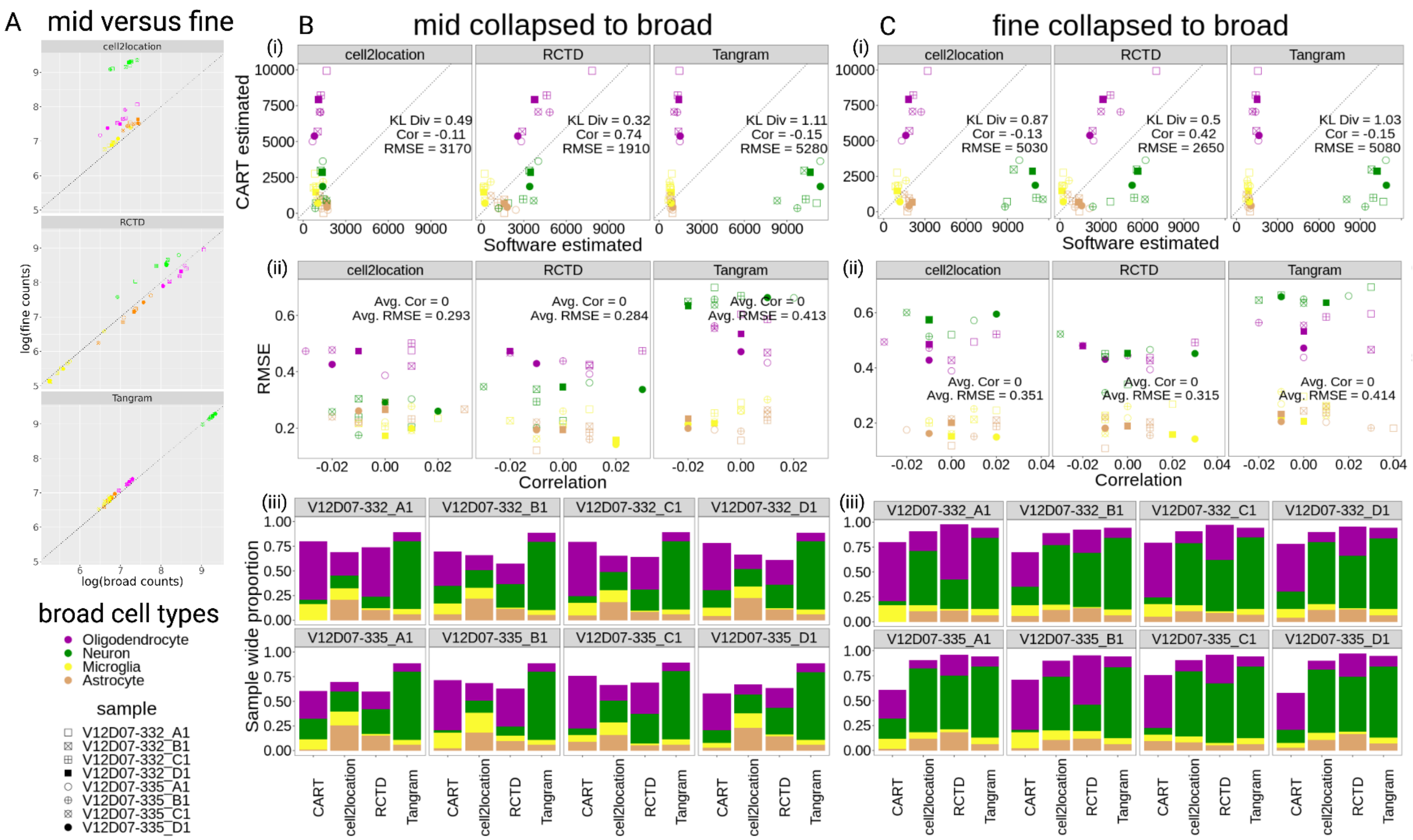


**Extended Data Fig. 37. Distribution of fine versus mid cell type proportions**.

(**A**) Comparison of cell type composition of 8 mid cell types with 21 fine cell types (collapsed down to 8 mid cell types) predictions from all 3 tools benchmarked across `PRECAST` spatial domains in Visium-SPG data. See **Supplementary Table 2** for abbreviations.

(**B**) Cell type composition of 21 fine cell type predictions from `RCTD` in Visium-H&E data.

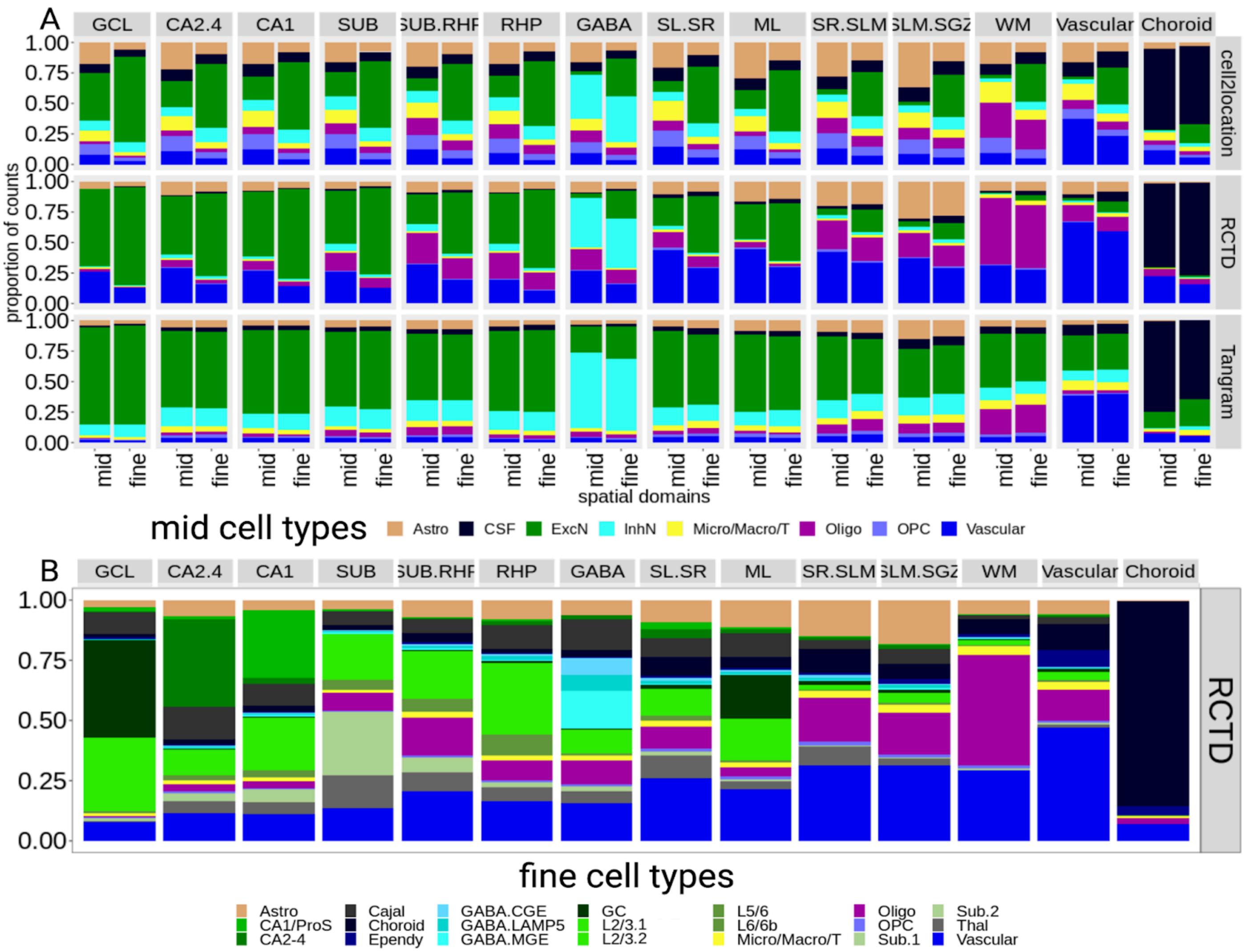
A
GCL
CA2.4
CA1
SUB
SUB.RHP
RHP
GABA
SL.SR
ML
SR.SLM
SLM.SGZ
WM
Vascular
Choroid
cell2location
RCTD
Tangram
proportion of counts
mid
fine
spatial domains
mid cell types
Astro
CSF
ExcN
InhN
Micro/Macro/T
Oligo
OPC
Vascular
B
RCTD
fine cell types
Astro
CA1/ProS
CA2-4
Cajal
Choroid
Ependy
GABA.CGE
GABA.LAMP5
GABA.MGE
GC
L2/3.1
L2/3.2
L5/6
L6/6b
Micro/Macro/T
Oligo
OPC
Sub.1
Sub.2
Thal
Vascular

**Extended Data Fig. 38. Schematic representation of non-negative matrix factorization.**

The top portion represents a schematic for how to identify patterns using non-negative matrix factorization (NMF). Specifically, for a given source matrix *A* with dimensions of *i* genes and *j* observations, such as the snRNA-seq data, we use NMF to decompose *A* into two matrices *W*, representing the feature-level weights matrix with dimensions of *i* genes and *k* NMF patterns, and *H*, representing the observation-level weights matrix for *j* observations and *k* patterns. Gray matrices are known matrices, blue matrices are generated by the analysis step.

The bottom portion represents a schematic for how we transfer the NMF patterns into other datasets, for example the SRT data. Specifically, we use the transposed feature-level weights matrix *W* (learned from the source data, glow) and a target dataset matrix (*A'* with dimensions of *i'* genes and *j'* observations) to obtain a new observation-level weights matrix from the target (*H'* with dimensions of *k* patterns and *j'* observations). Therefore, for every observation (spot or nuclei) in the target dataset, we have a weight for every corresponding NMF pattern learned from the source data (snRNA-seq). Gray matrices are known matrices, blue matrices are generated by the analysis step.

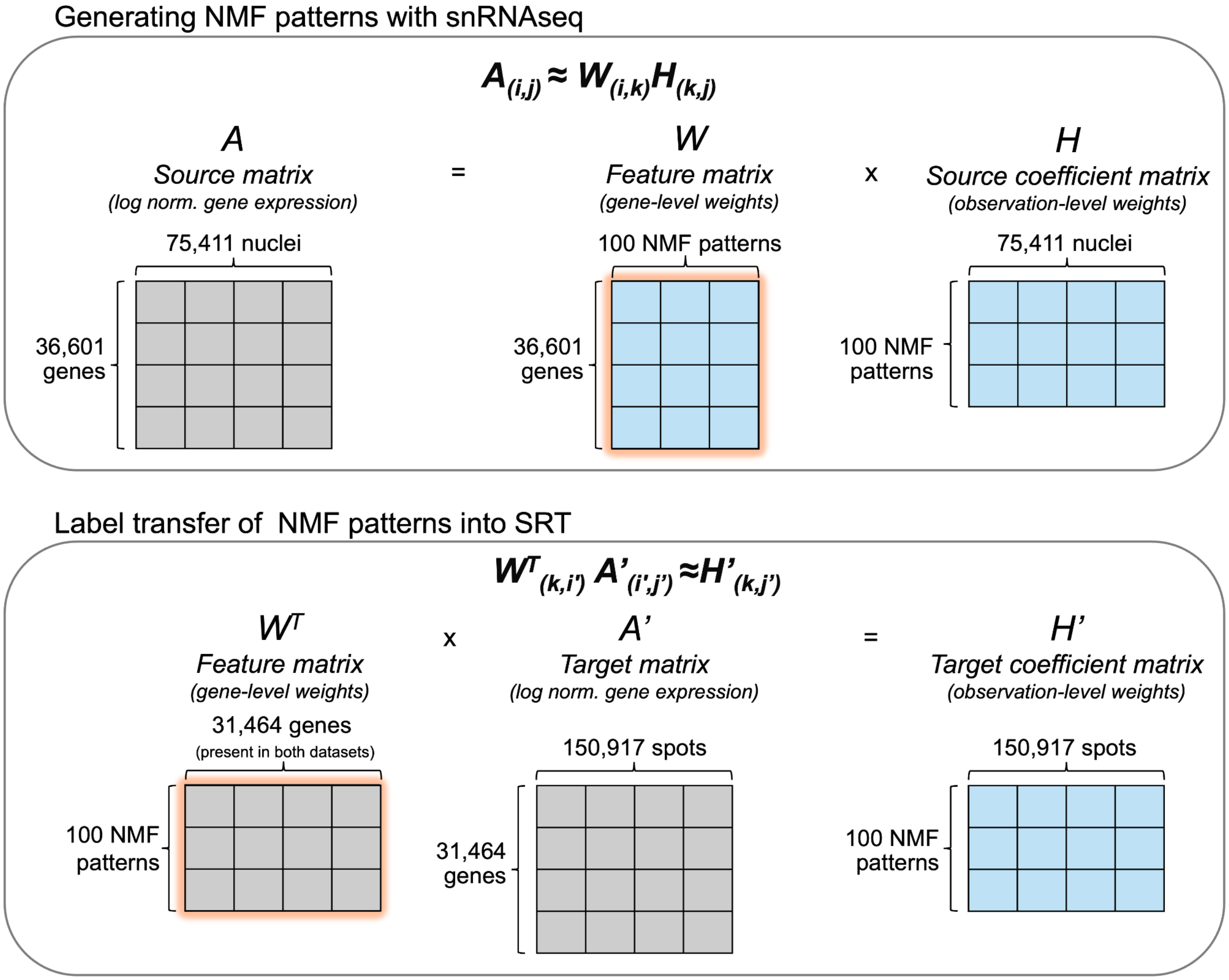

**Extended Data Fig. 39. Results from non-negative matrix factorization (NMF) cross-validation using 3 replicates.**

Line plot showing test set mean squared error (MSE) (*y*-axis) across different NMF ranks (*x*-axis) (**Methods 4.9**). $Log_2$ transformed, normalized single-nucleus RNA-sequencing (snRNA-seq) counts were split into training and test sets comprising 80% and 20% of the data, respectively. NMF was performed on the training set for various *k* values (*x*-axis). Model performance was evaluated in the test set based on MSE. This process was repeated three times (line color). NMF began to overfit to the training set at *k*=150 and *k*=200 for all replicates (**Methods 4.9).**

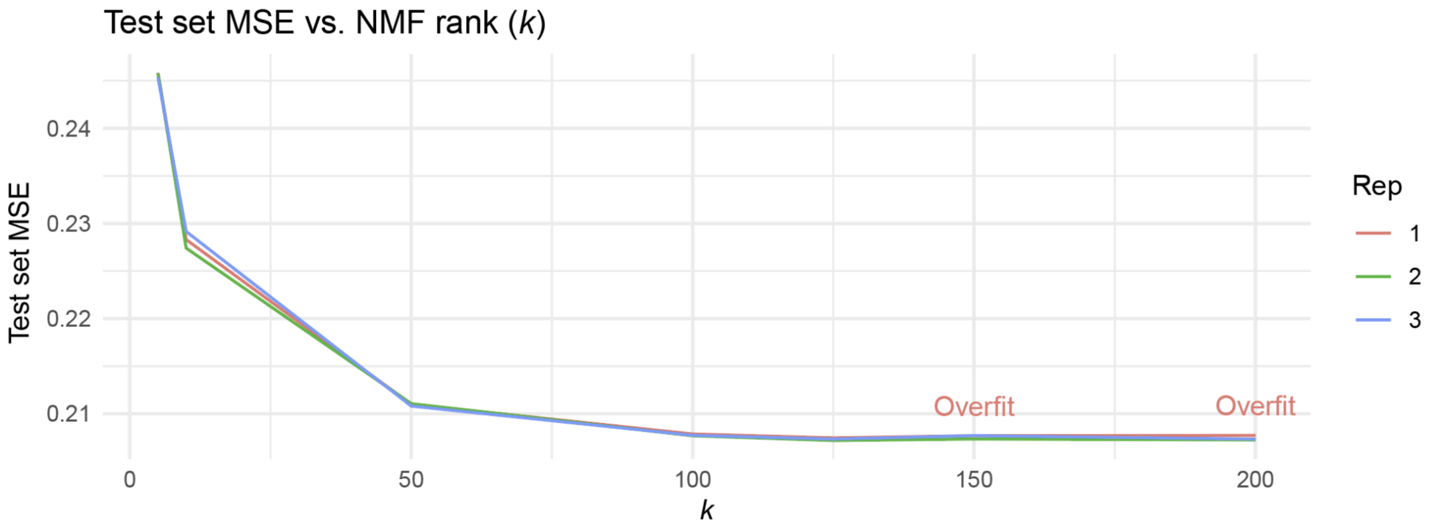

**Extended Data Fig. 40. Non-negative matrix factorization (NMF) patterns recapitulate known transcriptional distinctions by cell type and spatial domain.**

(**A**) NMF patterns (*x*-axis) learned from snRNA-seq $log_2$ normalized gene expression counts correspond well with snRNA-seq clusters (*y*-axis). For dot color, the nuclei-level weights for each NMF pattern were scaled and then averaged across the *y*-axis groups. Many of these patterns mapped to a high number of SRT spots after transfer and can be classified as general NMF patterns and specific NMF patterns based on the abundance of non-zero weighted nuclei across snRNA-seq cell type clusters (dot size). See **Supplementary Table 2** for *y*-axis abbreviations.

(**B**) All NMF patterns (points) learned from snRNA-seq data produce non-zero weights for thousands of nuclei (*x*-axis).

(**C**) When these NMF patterns are transferred to SRT data, the number of NMF patterns (points) with few non-zero weighted spots (*x*-axis) increases. NMF patterns that produced non-zero weights in fewer than 1050 spots (34 patterns, red points) were excluded from downstream analysis, as were the sex-based patterns nmf37 and nmf28 (**Extended Data Fig. 41**).

(**D**) After transfer to SRT data, NMF patterns with non-zero weights in >1050 spots (*x*-axis) correspond well to spatial domains (*y*-axis). With one exception (nmf94, discussed in **Extended Data Fig. 42**), the NMF patterns classified as general or specific in snRNA-seq data (**A**), likewise are abundant across multiple spatial domains or specific spatial domains. For dot color, the spot-level weights for each NMF pattern were scaled and then averaged across the *y*-axis groups. Dot size indicates the proportion of spots belonging to each *y*-axis group that exhibited non-zero weights for the given NMF pattern (*x*-axis). See **Supplementary Table 2** for *y*-axis abbreviations.

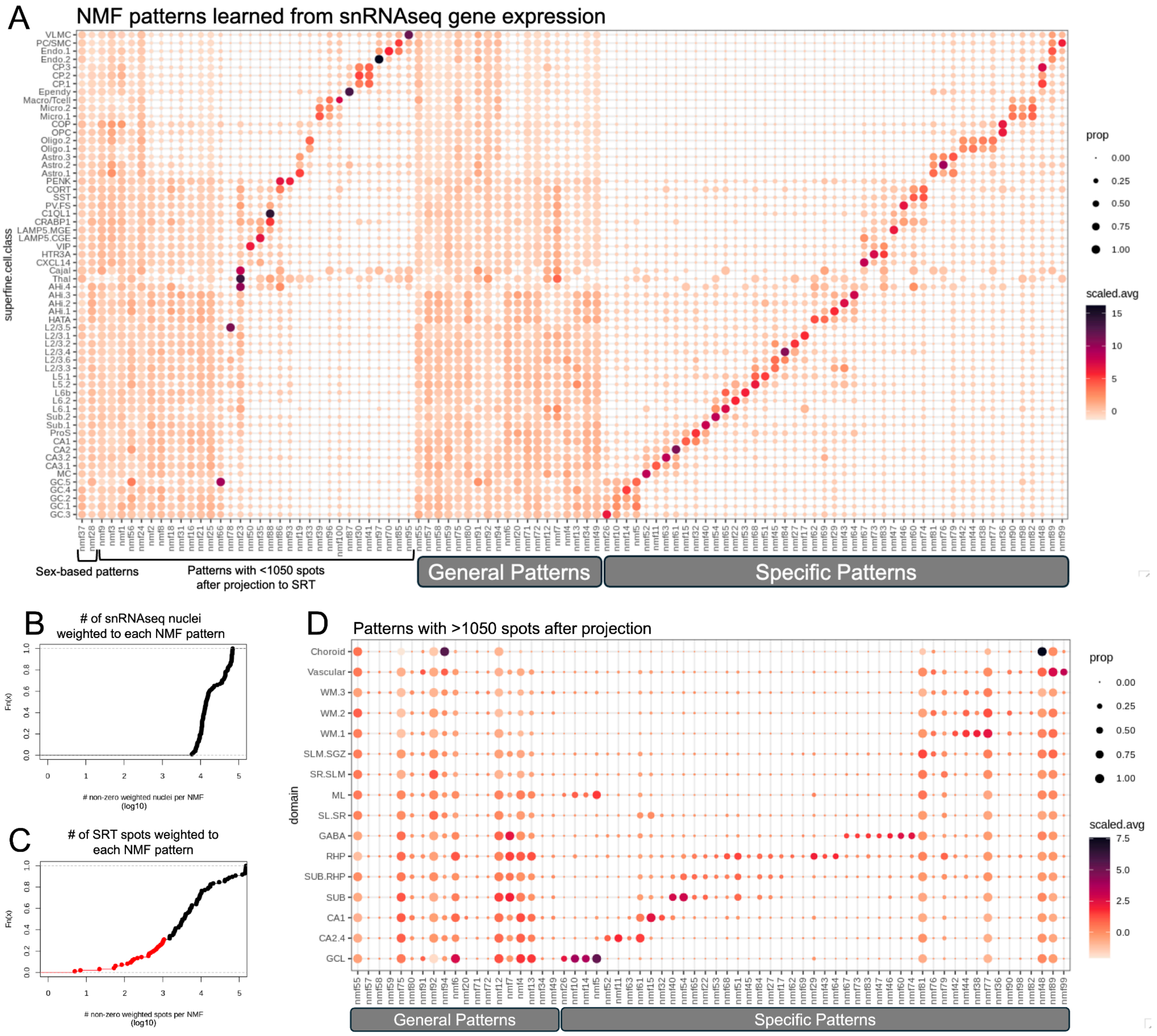

A
NMF patterns learned from snRNAseq gene expression
superfine.cell.class
prop
0.00
0.25
0.50
0.75
1.00
scaled.avg
15
10
5
0
Sex-based patterns
Patterns with <1050 spots after projection to SRT
General Patterns
Specific Patterns
B
### of snRNAseq nuclei weighted to each NMF pattern
Fn(x)
### non-zero weighted nuclei per NMF (log10)
C
### of SRT spots weighted to each NMF pattern
Fn(x)
### non-zero weighted spots per NMF (log10)
D
Patterns with >1050 spots after projection
domain
Choroid
Vascular
WM.3
WM.2
WM.1
SLM.SGZ
SR.SLM
ML
SL.SR
GABA
RHP
SUB.RHP
SUB
CA1
CA2.4
GCL
prop
0.00
0.25
0.50
0.75
1.00
scaled.avg
7.5
5.0
2.5
0.0
General Patterns
Specific Patterns

**Extended Data Fig. 41. Non-negative matrix factorization (NMF) patterns capture sex-based gene expression programs.**

(**A**) Uniform manifold approximation (UMAP) plot of snRNA-seq nuclei colored by nmf37 nuclei weights identified nuclei obtained from female donors and is almost identical to (**B**) the UMAP plot of $\log_2$ normalized expression of *XIST*, the top weighted gene for this pattern.

(**C**) UMAP plot of snRNA-seq nuclei colored by nmf28 nuclei weights identifies nuclei obtained male donors. The 4 genes most highly weighted to nmf28 originate from the Y chromosome: *TTTY14*, *NLGN4Y*, *USP9Y*, and *UTY*.

(**D**) UMAP plot of snRNA-seq nuclei colored by $\log_2$ normalized expression of *USP9Y* is extremely similar to the pattern of nmf28 weights in (**C**).

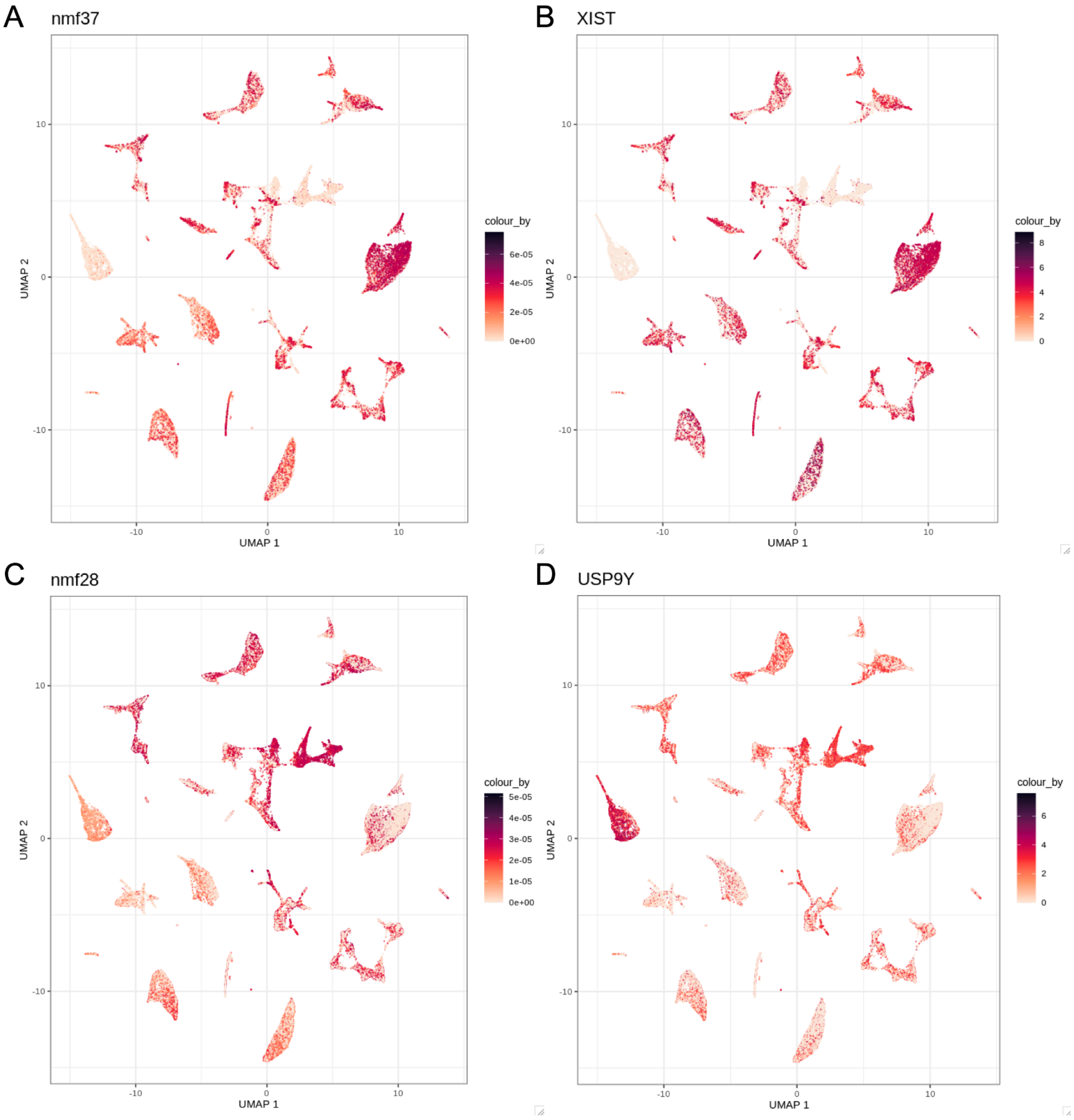

**Extended Data Fig. 42. Non-negative matrix factorization (NMF), donor effect, and projection.**

(**A**) The average nuclei (snRNA-seq, left) or spot (SRT, right) weights for nmf94 and nmf48 (*x*-axis), grouped by broad cell type (left) or domain (right) (*y*-axis). The size of the dots indicates the proportion of nuclei/spots in the y-axis group with non-zero weights for the given NMF pattern. See **Supplementary Table 2** for *y*-axis abbreviations.

(**B**) Uniform manifold approximation (UMAP) plot of snRNA-seq data colored by $\log_2$ normalized *TTR* expression. *TTR*, a choroid plexus marker gene, is the most highly weighted gene for nmf94 (**C**) and nmf48 (**D**).

(**E**) Choroid plexus nuclei (snRNA-seq, top) or spots (SRT, bottom) were isolated from a limited number of donors. Pie charts depict the proportion of choroid plexus-annotated observations from either dataset originating from each donor (fill color). Top: Br6471 (light gray) n= 1613 choroid plexus nuclei; Br6522 (yellow) n= 1549 choroid plexus nuclei. Bottom: Br6522 (yellow) n= 942 choroid plexus spots; Br8325 (aqua) n= 225 choroid plexus spots; Br8667 (dark gray) n= 159 choroid plexus spots; Br3942 (pink) n= 96 choroid plexus spots.

(**F**) nmf94 nuclei weights (top, *y*-axis) are not specifically increased in choroid plexus-annotated nuclei (red) but rather in the the non-choroid plexus nuclei (Not CP; grey) from donors (*x*-axis) with the greatest abundance of choroid plexus in the snRNA-seq dataset. In contrast, when projected to the SRT dataset (bottom), nmf94 spots weights (*y*-axis) are highest in choroid plexus spots and exhibit no trend by donor (*x*-axis).

(**G**) nmf48 weights (*y*-axis) are consistently increased only in choroid plexus nuclei (snRNA-seq, top) and spots (SRT, bottom) and still reflect the origin of choroid plexus observations from a subset of donors (*x*-axis).

Spot plots of spot-level (**H**) logcount expression of *TTR*, (**I**) weights of nmf94, and (**J**) weights of nmf48 in example capture areas from donor Br3942.

(**K**) Scatter plots of nmf94 spot-level weights (*x*-axis) and nmf48 spot-level weights (*y*-axis), faceted by the broad domain spot annotation.

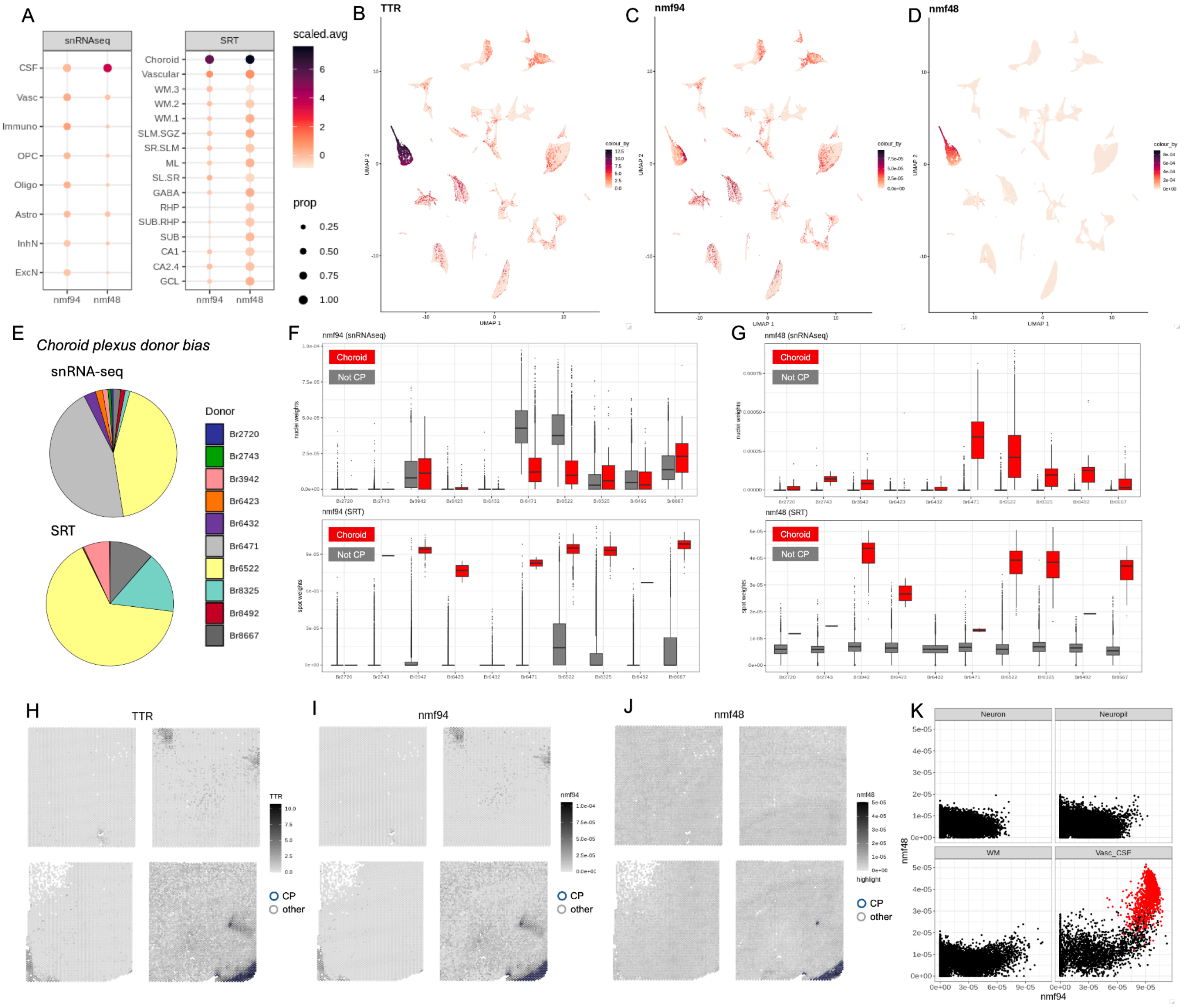

A
snRNAseq
SRT
scaled.avg
prop
CSF
Vasc
Immuno
OPC
Oligo
Astro
InhN
ExcN
Choroid
Vascular
WM.3
WM.2
WM.1
SLM.SGZ
SR.SLM
ML
SL.SR
GABA
RHP
SUB.RHP
SUB
CA1
CA2.4
GCL
nmf94
nmf48
B
TTR
C
nmf94
D
nmf48
UMAP 1
UMAP 2
E
Choroid plexus donor bias
snRNA-seq
SRT
Donor
Br2720
Br2743
Br3942
Br6423
Br6432
Br6471
Br6522
Br8325
Br8492
Br8667
F
nmf94 (snRNAseq)
nmf94 (SRT)
G
nmf48 (snRNAseq)
nmf48 (SRT)
Choroid
Not CP
H
TTR
I
nmf94
J
nmf48
CP
other
K
Neuron
Neuropil
WM
Vasc_CSF
nmf94
nmf48

**Extended Data Fig. 43. Non-negative matrix factorization (NMF) patterns by donor and sample.**

Bar plots show the proportion of snRNA-seq nuclei (*x*-axis) with non-zero weights to (**A**) general and (**B**) specific NMF patterns (*y*-axis) originating from individual samples (fill color, labeled by donor, sequencing round, and sorting protocol). Samples originating from the same donor are depicted in monochromatic palettes. nmf94 (A) and nmf48 (B) (discussed in **Extended Data Fig. 42**) show the largest enrichment from specific donors. nmf12 (A) and nmf89 (B) exhibit an enrichment in nuclei originating from donor Br8492 (salmon, dark red) but consistent contributions from all other donors. Due to differences in tissue composition in the tissue block, samples from some donors, like Br2743 (greens), do not contain much dentate gyrus and therefore have fewer nuclei with non-zero weights in NMF patterns that specifically correspond to these regions (nmf26, nmf10, nmf14, nmf5). Other donors, like Br2720 (blues) have greater enrichment for dentate gyrus tissue and the corresponding NMF patterns.

(**C**) A bar plot showing the proportion of SRT spots (*x*-axis) with non-zero weights to general NMF patterns (*y*-axis) originating from each donor (fill color). Other than nmf94 (discussed in **Extended Data Fig. 42**), nmf91 and nmf20 (indicated by asterisks) exhibit substantial bias in the donor from which the non-zero weighted spots originated. This is explored further in **Extended Data Fig. 47**.

(**D**) A bar plot showing the proportion of SRT spots (*x*-axis) with non-zero weights to cell type- and/or domain-specific NMF patterns (*y*-axis) originating from each donor (fill color). As with the snRNA-seq data, differences in tissue composition are evident in certain NMF patterns that map to specific spatial domains that were enriched in specific donors (e.g. Br6423 (orange) and amygdala-associated NMF patterns nmf62, nmf69, nmf29, nmf43, nmf64).

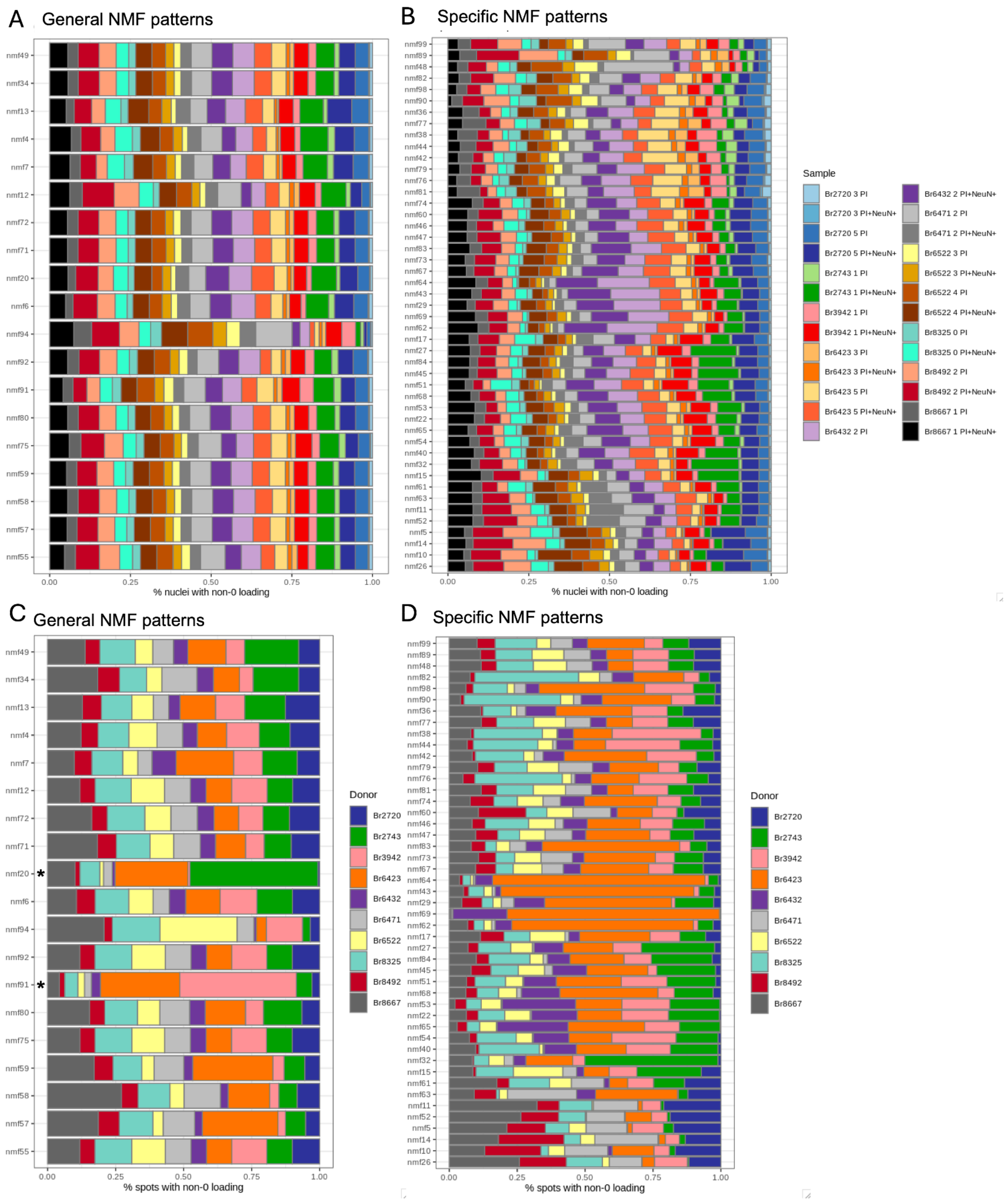
A General NMF patterns
B Specific NMF patterns
C General NMF patterns
D Specific NMF patterns
% nuclei with non-0 loading
% spots with non-0 loading
0.00
0.25
0.50
0.75
1.00
Sample
Br2720 3 PI
Br2720 3 PI+NeuN+
Br2720 5 PI
Br2720 5 PI+NeuN+
Br2743 1 PI
Br2743 1 PI+NeuN+
Br3942 1 PI
Br3942 1 PI+NeuN+
Br6423 3 PI
Br6423 3 PI+NeuN+
Br6423 5 PI
Br6423 5 PI+NeuN+
Br6432 2 PI
Br6432 2 PI+NeuN+
Br6471 2 PI
Br6471 2 PI+NeuN+
Br6522 3 PI
Br6522 3 PI+NeuN+
Br6522 4 PI
Br6522 4 PI+NeuN+
Br8325 0 PI
Br8325 0 PI+NeuN+
Br8492 2 PI
Br8492 2 PI+NeuN+
Br8667 1 PI
Br8667 1 PI+NeuN+
Donor
Br2720
Br2743
Br3942
Br6423
Br6432
Br6471
Br6522
Br8325
Br8492
Br8667
nmf49
nmf34
nmf13
nmf4
nmf7
nmf12
nmf72
nmf71
nmf20
nmf6
nmf94
nmf92
nmf91
nmf80
nmf75
nmf59
nmf58
nmf57
nmf55
nmf99
nmf89
nmf48
nmf82
nmf98
nmf90
nmf36
nmf77
nmf38
nmf44
nmf42
nmf79
nmf76
nmf81
nmf74
nmf60
nmf46
nmf47
nmf83
nmf73
nmf67
nmf64
nmf43
nmf29
nmf69
nmf62
nmf17
nmf27
nmf84
nmf45
nmf51
nmf68
nmf53
nmf22
nmf65
nmf54
nmf40
nmf32
nmf15
nmf61
nmf63
nmf11
nmf52
nmf5
nmf14
nmf10
nmf26
*

**Extended Data Fig. 44. Non-negative matrix factorization (NMF) identifies cell type-specific transcriptional patterns.**

Boxplots of spot-level weights (*y*-axis) grouped by spatial domain (*x*-axis) for (**A**) oligodendrocyte pattern nmf44 and (**B**) by astrocyte pattern nmf81. See **Supplementary Table 2** for *x*-axis abbreviations.

Boxplots of spot-level weights (*y*-axis) grouped by manual annotation (*x*-axis) for (**C**) oligodendrocyte pattern nmf44 and (**D**) by astrocyte pattern nmf81. See **Supplementary Table 2** for *x*-axis abbreviations.

Boxplots of nuclei-level weights (*y*-axis) grouped by snRNA-seq fine cell classes (*x*-axis) for (**E**) oligodendrocyte pattern nmf44 and (**F**) by astrocyte pattern nmf81. See **Supplementary Table 2** for *x*-axis abbreviations.

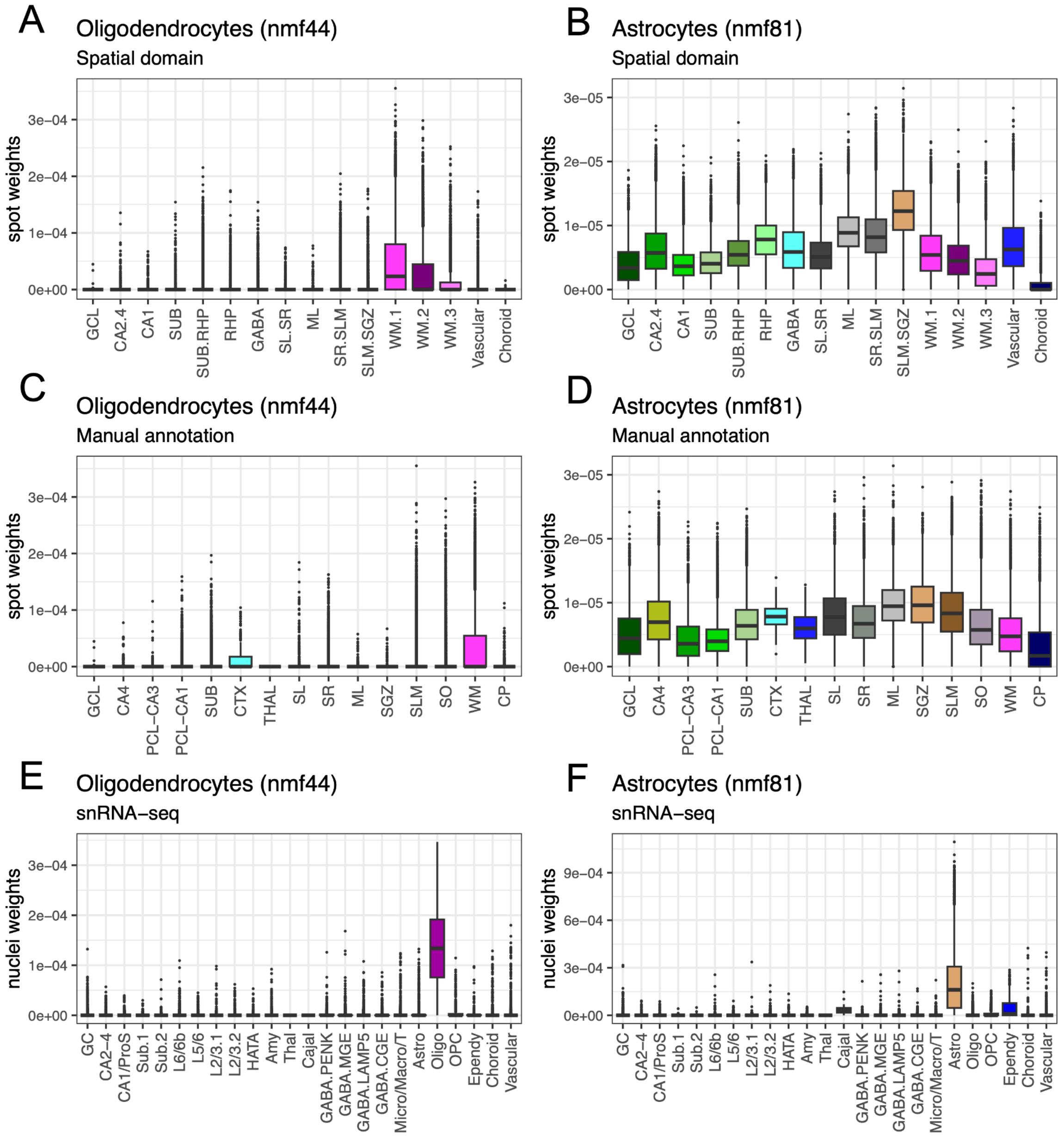

**Extended Data Fig. 45. Gene-level weights for all non-negative matrix factorization (NMF) patterns.**

Scatter plots of gene-level weights (*y*-axis) stratified by the average $\log_2$ normalized expression across all nuclei in our snRNA-seq dataset (*x*-axis). The top 10 genes weighted to each NMF pattern are indicated in red and for ease of readability these points are 1.66x larger than the remaining genes in black.

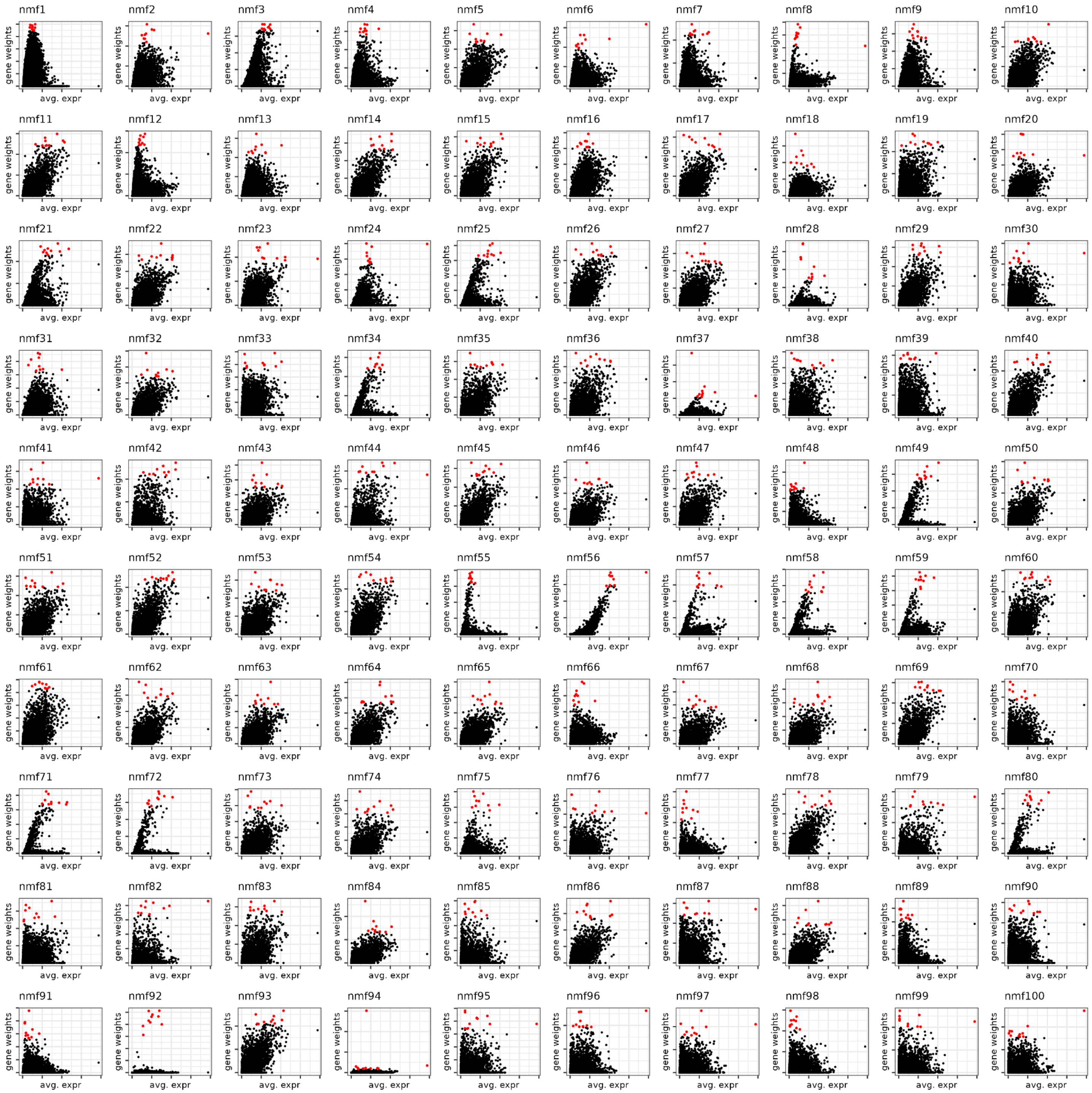

**Extended Data Fig. 46. Non-negative matrix factorization (NMF) identifies trans-neuronal transcriptional patterns.**

Boxplots of spot-level weights (*y*-axis) grouped by spatial domain (*x*-axis) for NMF patterns highlighting (**A**) excitatory postsynaptic specializations and (**B**) inhibitory postsynaptic specializations. See **Supplementary Table 2** for *x*-axis abbreviations.

Boxplots of spot-level weights (*y*-axis) grouped by manual annotation (*x*-axis) for NMF patterns highlighting (**C**) excitatory postsynaptic specializations and (**D**) inhibitory postsynaptic specializations. See **Supplementary Table 2** for *x*-axis abbreviations.

Boxplots of nuclei-level weights (*y*-axis) grouped by snRNA-seq fine cell classes (*x*-axis) for NMF patterns highlighting (**E**) excitatory postsynaptic specializations and (**F**) inhibitory postsynaptic specializations. See **Supplementary Table 2** for *x*-axis abbreviations.

(**G**) Scatter plots of spot-level weights for nmf13 (*x*-axis) and nmf7 (*y*-axis) faceted by SRT spatial domain.

(**H**) Scatter plots of nuclei-level weights for nmf13 (*x*-axis) and nmf7 (*y*-axis) faceted by snRNA-seq fine cell classes.

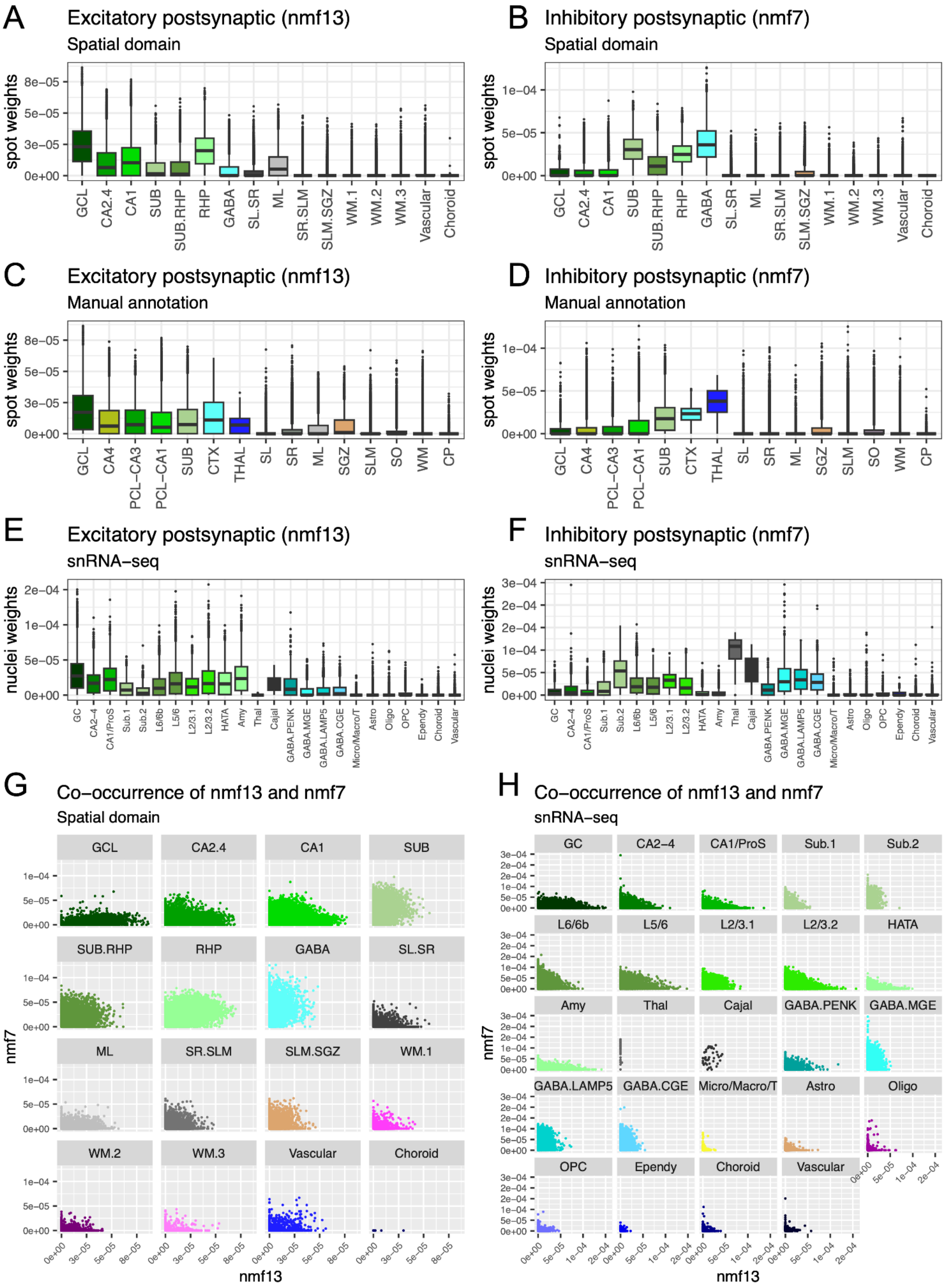
A
Excitatory postsynaptic (nmf13)
Spatial domain
spot weights
8e−05
5e−05
3e−05
0e+00
GCL
CA2.4
CA1
SUB
SUB.RHP
RHP
GABA
SL.SR
ML
SR.SLM
SLM.SGZ
WM.1
WM.2
WM.3
Vascular
Choroid
B
Inhibitory postsynaptic (nmf7)
Spatial domain
spot weights
1e−04
5e−05
0e+00
C
Excitatory postsynaptic (nmf13)
Manual annotation
spot weights
GCL
CA4
PCL−CA3
PCL−CA1
SUB
CTX
THAL
SL
SR
ML
SGZ
SLM
SO
WM
CP
D
Inhibitory postsynaptic (nmf7)
Manual annotation
spot weights
E
Excitatory postsynaptic (nmf13)
snRNA−seq
nuclei weights
2e−04
1e−04
5e−05
0e+00
GC
CA2−4
CA1/ProS
Sub.1
Sub.2
L6/6b
L5/6
L2/3.1
L2/3.2
HATA
Amy
Thal
Cajal
GABA.PENK
GABA.MGE
GABA.LAMP5
GABA.CGE
Micro/Macro/T
Astro
Oligo
OPC
Ependy
Choroid
Vascular
F
Inhibitory postsynaptic (nmf7)
snRNA−seq
nuclei weights
3e−04
2e−04
2e−04
1e−04
5e−05
0e+00
G
Co−occurrence of nmf13 and nmf7
Spatial domain
nmf7
nmf13
0e+00
3e−05
5e−05
8e−05
H
Co−occurrence of nmf13 and nmf7
snRNA−seq
nmf7
nmf13
0e+00
5e−05
1e−04
2e−04

**Extended Data Fig. 47. Non-negative matrix factorization (NMF) identifies donor enrichment due to activity-dependent transcription.**

(**A**) Pie charts for snRNA-seq nuclei (top) and SRT spots (bottom) show the proportion of observations with non-zero nmf91 weights originating from each donor. Top: snRNA-seq n= 39,991 total non-zero nmf91 nuclei. Bottom: SRT n= 9,380 total non-zero nmf91 spots; Br3942 (pink) n= 4020 non-zero nmf91 spots; Br6423 (orange) n= 2736 non-zero nmf91 spots.

(**B**) Nine of the 50 genes with the highest nmf91 weight (*y*-axis) are canonical immediate early genes (red) that are transcribed after strong stimulus in neurons and non-neuronal cells. Average $\log_2$ normalized gene expression in all nuclei on *x*-axis.

(**C**) Pie charts for snRNA-seq nuclei (top) and SRT spots (bottom) show the proportion of observations with non-zero nmf20 weights originating from each donor. Top: snRNA-seq n= 36,679 total non-zero nmf20 nuclei. Bottom: SRT n= 2109 total non-zero nmf20 spots; Br2743 (green) n= 991 non-zero nmf20 spots; Br6423 (orange) n= 565 non-zero nmf20 spots.

(**D**) The three genes (red) with the highest nmf20 weights (*y*-axis) are known regulators of neuronal function in response to activation. Average $\log_2$ normalized gene expression in all nuclei on *x*-axis.

(**E**) *JUN* and (**F**) *HOMER1* expression ($\log_2$ normalized, *y*-axis) is relatively equal in neuronal nuclei from all donors (*x*-axis), recapitulating the equal contribution of all donors to nuclei with non-zero weights from nmf91 and nm20, respectively. Red indicates neuronal nuclei, gray indicates non-neuronal nuclei.

(**G**) In contrast, the expression ($\log_2$ normalized, *y*-axis) of top nmf91 marker gene *JUN* is increased in spots from donors (*x*-axis) Br3942 and Br6423, the two donors that comprise a disproportionate number of spots with non-zero nmf91 weights. Red indicates neuronal spots, gray indicates non-neuronal spots.

(**H**) Similarly, the expression ($\log_2$ normalized, *y*-axis) of top nmf20 marker gene *HOMER1* is increased in neuronal spots from donors (*x*-axis) Br2743 and Br6423, the two donors that comprise a disproportionate number of spots with non-zero nmf20 weights. Red indicates neuronal spots, gray indicates non-neuronal spots.

(**I**) Uniform manifold approximation (UMAP) plot of *PDE1A* $\log_2$ normalized expression in our human snRNA-seq data.

(**J**) Violin plot of *PDE1A* $\log_2$ normalized expression (*y*-axis) in our human snRNA-seq data, grouped by fine cell class (*x*-axis). See **Supplementary Table 2** for *x*-axis abbreviations.

(**K**) Violin plot of *PDE1A* $\log_2$ normalized expression (*y*-axis) in our human SRT data, grouped by spatial domain (*x*-axis). See **Supplementary Table 2** for *x*-axis abbreviations.

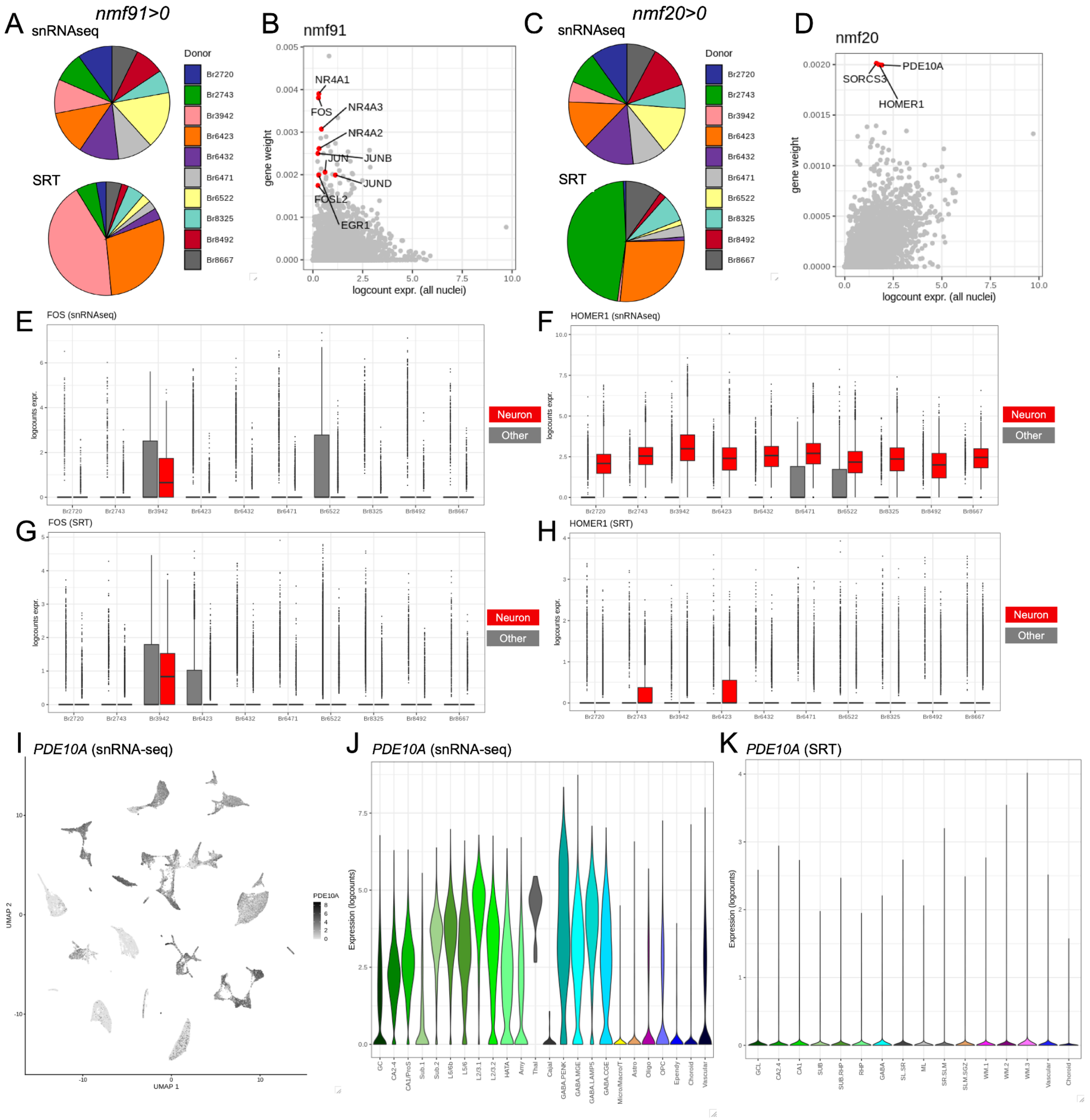

A
nmf91>0
snRNAseq
SRT
Donor
Br2720
Br2743
Br3942
Br6423
Br6432
Br6471
Br6522
Br8325
Br8492
Br8667
B
nmf91
NR4A1
FOS
NR4A3
NR4A2
JUN
JUNB
JUND
FOSL2
EGR1
gene weight
logcount expr. (all nuclei)
C
nmf20>0
snRNAseq
SRT
D
nmf20
PDE10A
SORCS3
HOMER1
E
FOS (snRNAseq)
F
HOMER1 (snRNAseq)
G
FOS (SRT)
H
HOMER1 (SRT)
logcounts expr.
Neuron
Other
I
PDE10A (snRNA-seq)
UMAP 1
UMAP 2
J
PDE10A (snRNA-seq)
Expression (logcounts)
K
PDE10A (SRT)

**Extended Data Fig. 48. Non-negative matrix factorization (NMF) patterns across annotated cell types in mouse neuron electroconvulsive stimulation (ECS) snRNA-seq data.**

(**A**) Following label transfer of NMF patterns to mouse snRNA-seq electroconvulsive stimulation (ECS) dataset with retroviral tracing dataset (93) ($n$=15990 nuclei), NMF patterns were removed (red) that mapped to <1000 nuclei ($x$-axis).

(**B**) Dotplot of mouse nuclei-level weights for NMF patterns ($x$-axis) after filtering, averaged by cluster ($y$-axis). Dot size indicates the proportion of nuclei in each cluster with non-zero pattern weights and dots are colored by the scaled pattern weight. GC: granule cells, CA2-4: *cornu ammonis* (CA) regions 2 through 4 (CA2, CA3, CA4), PS/Sub: prosubiculum and subiculum neurons, L5/Po: layer 5 and polymorphic layer.

(**C**) Dotplot of nuclei-level weights for NMF patterns ($x$-axis) after further filtering to patterns that were present in >1050 SRT spots, averaged by cluster and ECS condition ($y$-axis). Dot size indicates the proportion of nuclei in each cluster with non-zero pattern weights and dots are colored by the scaled pattern weight averaged across nuclei in $y$-axis groups. Asterisks indicate patterns that are highlighted in **Figure 5**. Carrot indicates nmf55 that is described in (**D**,**E**) and **Methods 4.9**.

(**D**) Volcano plot of differential expression results tested on genes with non-zero nmf55 weights. $Y$-axis of $-\log_{10}$(FDR) values are plotted with a $\log_{10}$ scale. $X$-axis is the $\log_2$ fold change (FC), where negative values indicate greater expression in sham-activated GCs and positive values indicate greater expression in ECS GCs. Points are colored by nmf55 weight. Gene names are shown for genes with nmf55 weight>.0025, $\log_2$FC>0.5, and FDR<0.05.

(**E**) Spot plots for example capture area from donor Br3942 are colored by (left) spatial domain and (right) nmf55 weight, demonstrating the ubiquitous presence of nmf55. See **Supplementary Table 2** for spatial domain abbreviations.

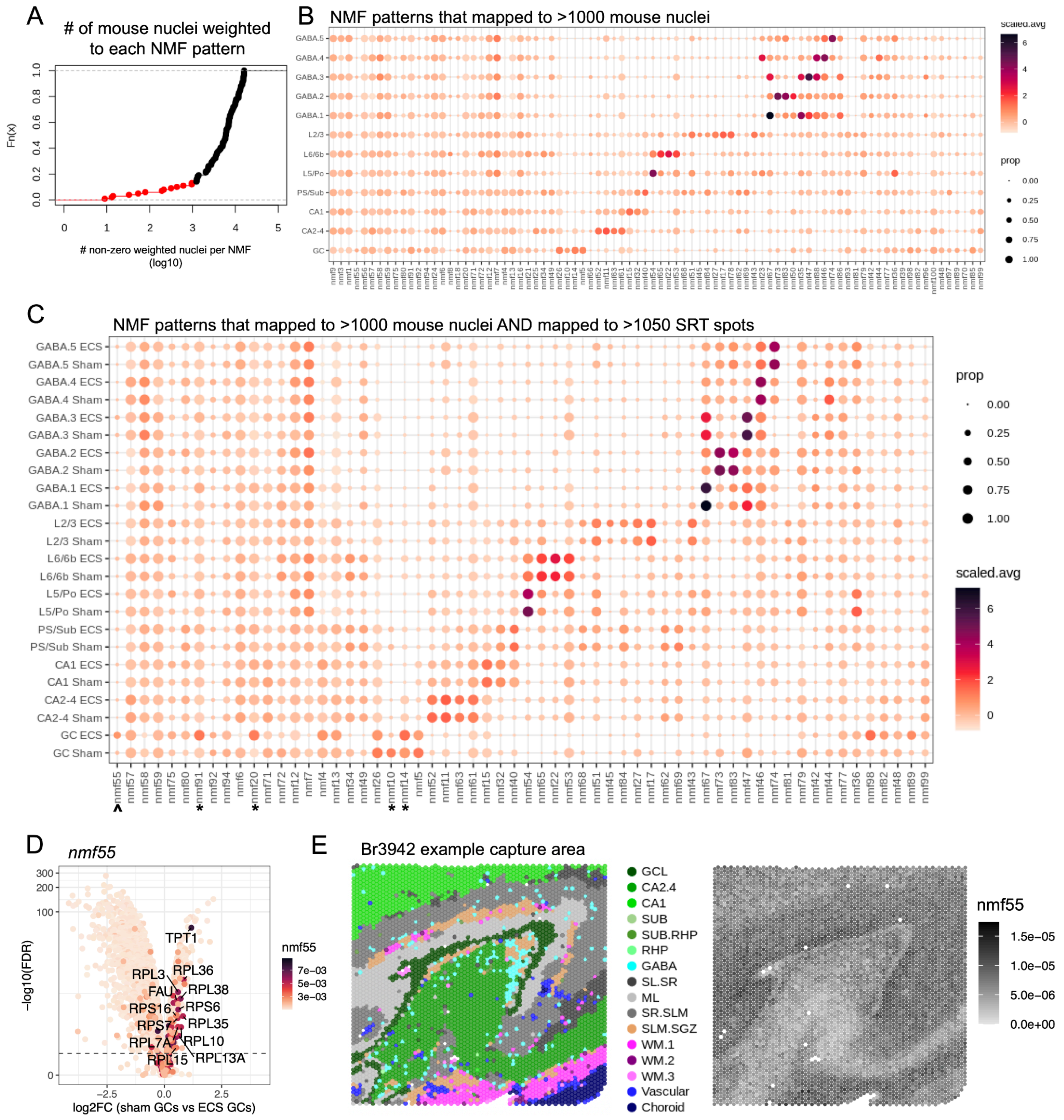
A
### of mouse nuclei weighted to each NMF pattern
Fn(x)
### non-zero weighted nuclei per NMF (log10)
B
NMF patterns that mapped to >1000 mouse nuclei
GABA.5
GABA.4
GABA.3
GABA.2
GABA.1
L2/3
L6/6b
L5/Po
PS/Sub
CA1
CA2-4
GC
scaled.avg
prop
C
NMF patterns that mapped to >1000 mouse nuclei AND mapped to >1050 SRT spots
GABA.5 ECS
GABA.5 Sham
GABA.4 ECS
GABA.4 Sham
GABA.3 ECS
GABA.3 Sham
GABA.2 ECS
GABA.2 Sham
GABA.1 ECS
GABA.1 Sham
L2/3 ECS
L2/3 Sham
L6/6b ECS
L6/6b Sham
L5/Po ECS
L5/Po Sham
PS/Sub ECS
PS/Sub Sham
CA1 ECS
CA1 Sham
CA2-4 ECS
CA2-4 Sham
GC ECS
GC Sham
prop
scaled.avg
D
nmf55
-log10(FDR)
log2FC (sham GCs vs ECS GCs)
TPT1
RPL3
RPL36
FAU
RPL38
RPS16
RPS6
RPS7
RPL35
RPL7A
RPL10
RPL15
RPL13A
E
Br3942 example capture area
GCL
CA2.4
CA1
SUB
SUB.RHP
RHP
GABA
SL.SR
ML
SR.SLM
SLM.SGZ
WM.1
WM.2
WM.3
Vascular
Choroid
nmf55

**Extended Data Fig. 49. Non-negative matrix factorization (NMF) label transfer to mouse single-nucleus methylation sequencing (snmC-seq) dataset, subiculum annotation via binarization of NMF patterns, and elucidation of new deep subiculum cluster snRNA-seq cluster.**

(**A**) Following label transfer of NMF patterns to mouse snmC-seq with retroviral tracing dataset (106) (*n*=2004 nuclei), hippocampal (HPC)- and retrohippocampal (RHP)-specific patterns were removed (red) that mapped to <45 nuclei (*x*-axis).

(**B**) Verification that NMF patterns (*x*-axis) identified as HPC- and RHP-specific in our human SRT and human snRNA-seq datasets corresponded with the nuclei collection source (*y*-axis) for the mouse snmC-seq dataset. Dots are colored by scaled, average pattern weight and dot size indicates the number of mouse nuclei with non-zero pattern weights. CA: *cornu ammonis* regions, SUB: subiculum.

(**C**) To assist in determining whether RHP pyramidal NMF patterns overlapped with the subicular complex, we binarized the two patterns corresponding to the superficial (nmf40) and middle (nmf54) subiculum (**Figure 6A**). Violin plots present the spot-level weights (*y*-axis) across spatial domains (*x*-axis). Thresholds are indicated by the dotted lines, determined by taking one fifth of the 95% max spot-level weight. See **Supplementary Table 2** for spatial domain abbreviations.

(**D**) Spots assigned to the subiculum (SUB) by binarizing NMF patterns were specific to the SUB spatial domain. Bar plots show the number of spots (*x*-axis) for each spatial domain (*y*-axis), colored by the new SUB identity as determined with binary thresholding. Spots not labeled as SUB are present in gray, spots newly labeled as subiculum are present in green, with different shades indicating the largely non-overlapping distribution of nmf40 and nmf54.

(**E**) Dot plot demonstrating the increased expression of SUB marker genes in L6.1 and not the portion of L6b highly weighted to nmf65 (L6b_nmf65). SUB marker genes were isolated by the overlap of significant pseudobulk snRNA-seq results and top nmf40 and nmf54 weighted-genes. Dot size indicates the proportion of nuclei in each cluster (*x*-axis) with non-zero expression of the indicated gene (*y*-axis). Dot color indicates the average $\log_2$ normalized expression. See **Supplementary Table 2** for cluster abbreviations.

(**F**) *t*-distributed stochastic neighbor embedding (TSNE) plots of the expression of select marker genes from (**E**) in the pyramidal nuclei of our human snRNA-seq results. Points are colored by the average $\log_2$ normalized expression. The expression of SUB marker genes *TOX*, *TSHZ2*, and *ZNF385D* and RHP L6 genes *SGCZ* and *NFIA* in the labeled clusters support the reassignment of L6.1 as deep SUB nuclei (Sub.3).

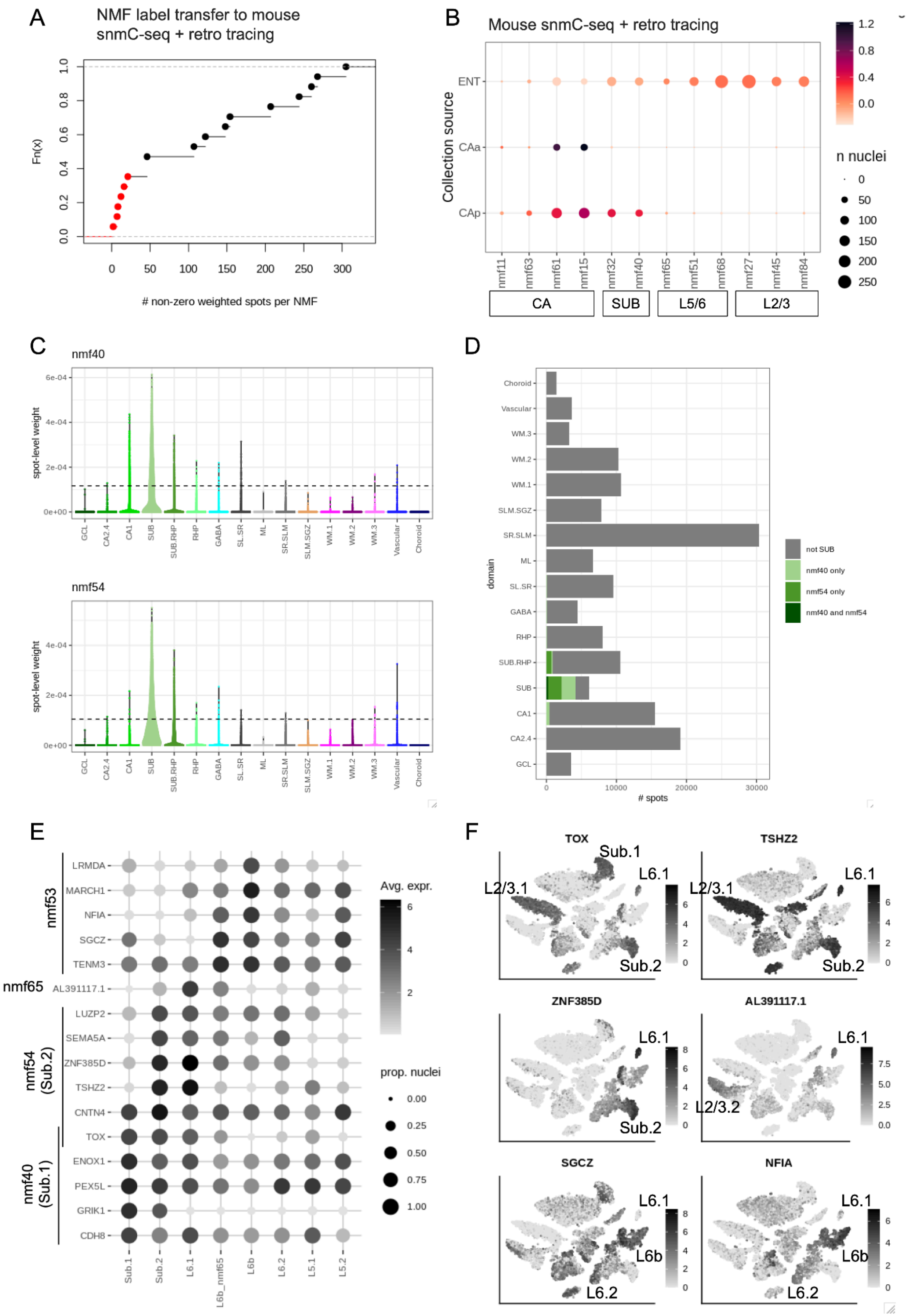
A
NMF label transfer to mouse
snmC-seq + retro tracing
Fn(x)
### non-zero weighted spots per NMF
B
Mouse snmC-seq + retro tracing
Collection source
ENT
CAa
CAp
nmf11
nmf63
nmf61
nmf15
nmf32
nmf40
nmf65
nmf51
nmf68
nmf27
nmf45
nmf84
CA
SUB
L5/6
L2/3
n nuclei
C
nmf40
nmf54
spot-level weight
GCL
CA2.4
CA1
SUB
SUB.RHP
RHP
GABA
SL.SR
ML
SR.SLM
SLM.SGZ
WM.1
WM.2
WM.3
Vascular
Choroid
D
domain
### spots
not SUB
nmf40 only
nmf54 only
nmf40 and nmf54
E
nmf53
nmf65
nmf54 (Sub.2)
nmf40 (Sub.1)
LRMDA
MARCH1
NFIA
SGCZ
TENM3
AL391117.1
LUZP2
SEMA5A
ZNF385D
TSHZ2
CNTN4
TOX
ENOX1
PEX5L
GRIK1
CDH8
Sub.1
Sub.2
L6.1
L6b_nmf65
L6b
L6.2
L5.1
L5.2
Avg. expr.
prop. nuclei
F
TOX
TSHZ2
ZNF385D
AL391117.1
SGCZ
NFIA
Sub.1
Sub.2
L6.1
L2/3.1
L2/3.2
L6b
L6.2

**Extended Data Fig. 50. Binarization of non-negative matrix factorization (NMF) patterns for simultaneous visualization on spot plots.**

(**A**) Table indicating how many spots were classified to each individual threshold, with 1 indicating classification for the column and colored lightly. The total number of spots classified by each threshold are italicized at the bottom of the corresponding column. Spots that were classified to only one of the domains were included in the final combined annotation (bright colored cells). Final combined annotation sample sizes: 5263 subiculum spots, 12640 CA1 spots, 10430 CA3 spots, 1209 superficial entorhinal cortex (ENT_sup) spots, 1462 entorhinal cortex layer 5 (ENT_L5) spots.

(**B**) The number of spots (*x*-axis) labeled by the final combined annotation (color) were distributed across the corresponding domains (*y*-axis). See **Supplementary Table 2** for *y*-axis abbreviations.

A

| subiculum | CA1 | CA3 | ENT_sup | ENT_L5 | n spots |
|---|---|---|---|---|---|
| 1 | 0 | 0 | 0 | 0 | 5263 |
| 1 | 0 | 0 | 0 | 1 | 115 |
| 1 | 0 | 0 | 1 | 0 | 13 |
| 1 | 0 | 0 | 1 | 1 | 1 |
| 1 | 0 | 1 | 0 | 0 | 5 |
| 1 | 1 | 0 | 0 | 0 | 266 |
| 1 | 1 | 1 | 0 | 0 | 9 |
| 1 | 1 | 1 | 0 | 1 | 1 |
| 0 | 1 | 0 | 0 | 0 | 12460 |
| 0 | 1 | 0 | 0 | 1 | 6 |
| 0 | 1 | 0 | 1 | 0 | 1 |
| 0 | 1 | 1 | 0 | 0 | 81 |
| 0 | 0 | 1 | 0 | 0 | 10430 |
| 0 | 0 | 0 | 1 | 0 | 1209 |
| 0 | 0 | 0 | 1 | 1 | 179 |
| 0 | 0 | 0 | 0 | 1 | 1462 |
| 0 | 0 | 0 | 0 | 0 | 119416 |
| *5673* | *12824* | *10526* | *1403* | *1764* | |

B

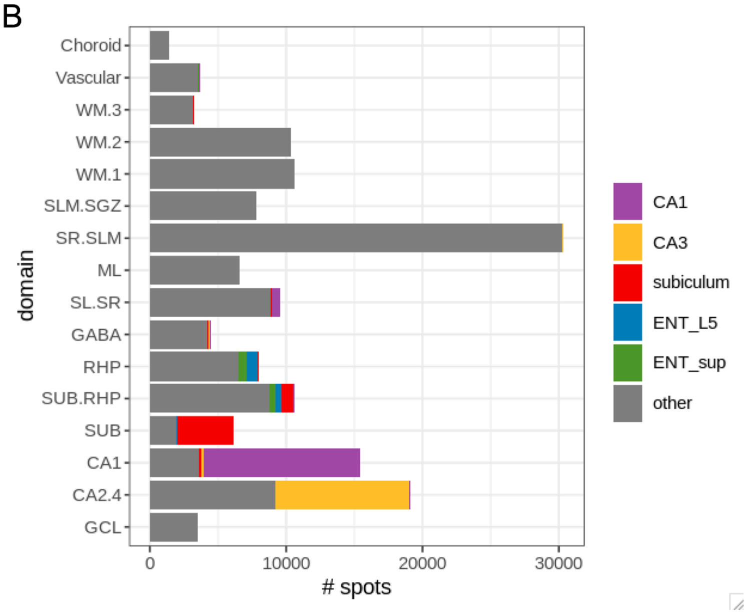

**Extended Data Fig. 51. Spatial organization of nmf17 and identification of the presubiculum.**

Spot outlines are colored by the combined annotation obtained by thresholding spatially-restricted and cell type-specific NMF patterns (legend lower right) (**Extended Data Fig. 50**). Spot fill color indicates nmf17 weight (color bar lower right). The region believed to correspond to the presubiculum is indicated by an asterisk. Capture areas are grouped by donor number. ENT_L5: entorhinal cortex layer 5, ENT_sup: superficial entorhinal cortex.

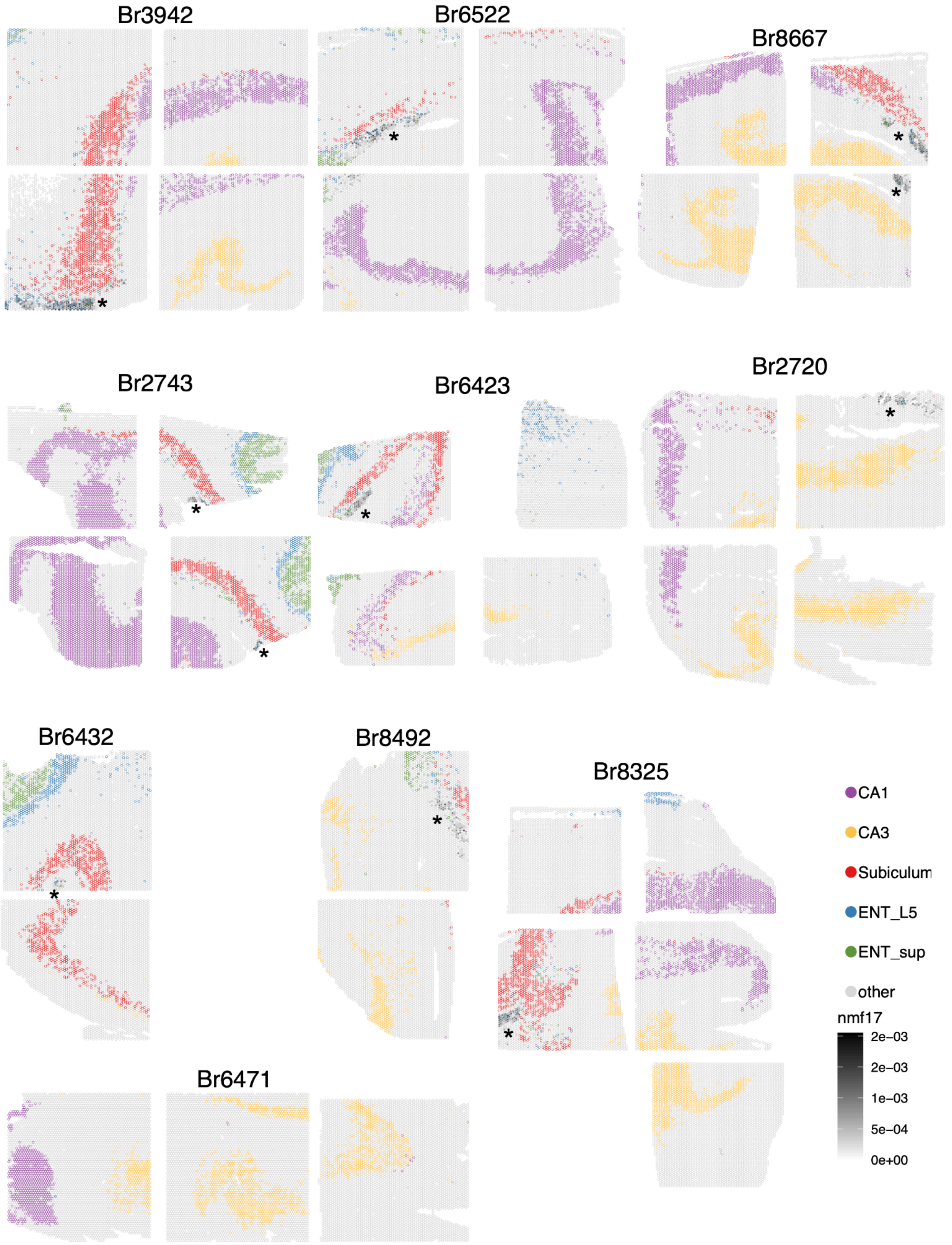

Br3942
Br6522
Br8667
Br2743
Br6423
Br2720
Br6432
Br8492
Br8325
Br6471
CA1
CA3
Subiculum
ENT_L5
ENT_sup
other
nmf17
2e−03
2e−03
1e−03
5e−04
0e+00

**Extended Data Fig. 52. Subiculum-focused differential expression analysis in snRNA-seq.**

Violin plots depict the $log_2$ normalized expression (*y*-axis) across snRNA-seq cell type clusters (*x*-axis) that were tested. See **Supplementary Table 2** for *x*-axis abbreviations. Plots are grouped by the significant cluster: (**A**) superficial subiculum (Sub.1), (**B**) middle subiculum (Sub.2), (**C**) deep subiculum (Sub.3), and (**D**) presubiculum (PreS).

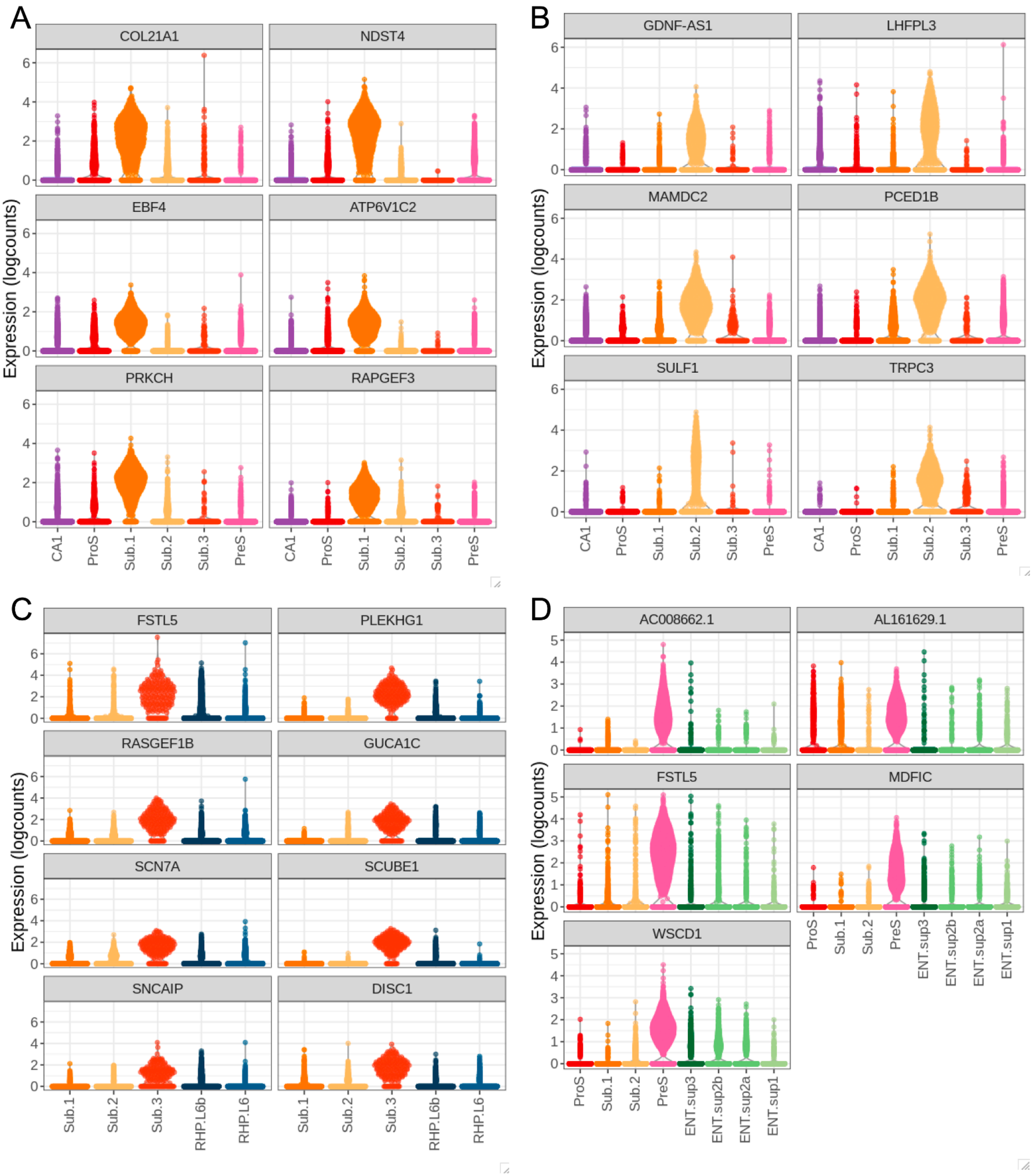

### Supplementary Tables

**Supplementary Table 1. Donor demographic information.** Demographic information on brain donors, including Sample ID (BrNum), age at time of death, sex, psychiatric diagnosis, post-mortem interval (PMI), body mass index (BMI, calculated postmortem), Screening RNA Integrity Number (RIN, calculated at time of brain collection in the prefrontal cortex–PFC), and, for some samples, RIN measured in the tissue blocks included in this study. The assays performed on each tissue sample are included.

**Supplementary Table 2**. **Abbreviations**. Guide to abbreviations used in naming spatial domains and snRNA-seq cell types.

**Supplementary Table 3. nnSVG gene ranks.** Results from nnSVG analysis, used to identify spatially variable genes (SVGs) in the SRT data. Columns include ENSEMBL ID, gene name, overall gene rank, mean gene rank across all capture areas, number of capture areas for which each gene was highly ranked (within top 1000 most variable genes), and whether each gene was used for `PRECAST` clustering.

**Supplementary Table 4. Differential expression results across pseudobulked spatial domains using the layer-enriched model.** For each gene, a linear mixed-effects model was fit with counts aggregated (or pseudobulked) across spots within a spatial domain to identify differences in expression enriched in one domain compared to all other domains using Student's *t*-test statistics.

**Supplementary Table 5. Differential expression results across four pseudobulked broad spatial domains using the layer-enriched model.** Four broad domains: neuron-enriched domains (comprised of pyramidal, granule and GABAergic neuron clusters), neuropil-enriched domains (including ML, SL, SR, SLM.SGZ), WM, and vascular/cerebrospinal fluid (CSF)-related domains (comprised of the two domains mapping to vasculature and choroid plexus, respectively). For each gene, a linear mixed-effects model was fit with counts aggregated (or pseudobulked) across spots within a spatial domain to identify differences in expression enriched in one domain compared to all other domains using Student's *t*-test statistics.

**Supplementary Table 6. Differential expression results across pseudobulked "superfine" cell types using the layer-enriched model.** For each gene, a linear mixed-effects model was fit with counts aggregated (or pseudobulked) across nuclei within each superfine cell type to identify differences in expression enriched in one superfine cell type compared to all other superfine cell types using Student's *t*-test statistics.

**Supplementary Table 7. Spot deconvolution cell type marker gene statistics.** For each cell type at mid and fine resolution, statistics of top 25 marker genes (mean ratio>1) extracted with the *DeconvoBuddies* package using the filtered (genes that overlapped with Visium data only) snRNA-seq data.

**Supplementary Table 8. RMSE and correlation stats.** For each cell type at mid and fine resolution, Pearson Correlation and RMSE for spot-deconvolution software results compared with CART predictions were calculated for each Visum-SPG capture area.

**Supplementary Table 9**. **Gene-level NMF weights.** Table with gene-level weights for NMF patterns identified using `RcppML`. Each row is a gene and each column is an NMF pattern. Patterns were normalized such that each column sums to 1.

**Supplementary Table 10. Top ten weighted genes for each NMF pattern.** The top ten genes weighted to each pattern in Supplementary Table 9 are isolated and presented alongside the gene-level weights.
